# Inhibition of chloroplast translation as a new target for herbicides

**DOI:** 10.1101/2021.09.28.462089

**Authors:** Kirill V. Sukhoverkov, Karen J. Breese, Aleksandra W. Debowski, Monika W. Murcha, Keith A. Stubbs, Joshua S. Mylne

**Affiliations:** The University of Western Australia, School of Molecular Sciences, 35 Stirling Highway, Crawley, Perth 6009, Australia; The ARC Centre of Excellence in Plant Energy Biology, 35 Stirling Highway, Crawley, Perth 6009, Australia; School of Biomedical Sciences, 35 Stirling Highway, Crawley, Perth 6009, Australia; Centre for Crop and Disease Management, School of Molecular and Life Sciences, Curtin University, Bentley, WA 6102, Australia

## Abstract

The rise in herbicide resistance over recent decades threatens global agriculture and food security and so discovery of new modes of action is increasingly important. Here we reveal linezolid, an oxazolidinone antibiotic that inhibits microbial translation, is also herbicidal. To validate the herbicidal mode of action of linezolid we confirmed its micromolar inhibition is specific to chloroplast translation and did not affect photosynthesis directly. To assess the herbicide potential of linezolid, testing against a range of weed and crop species found it effective pre- and post-emergence. Using structure-activity analysis we identified the critical elements for herbicidal activity, but importantly also show, using antimicrobial susceptibility assays, that separation of antibacterial and herbicidal activities was possible. Overall these results validate chloroplast translation as a viable herbicidal target.

## Introduction

Implementing herbicides into agricultural practice in the 1940s greatly improved crop productivity, and although this success continues today, the growth of herbicide resistance in the last few decades puts world food security under threat. The first field case of herbicide resistance, triazine-resistant *Senecio vulgaris*, was documented in 1968 and the number of weed species resistant to one or more herbicides has increased to 263 in 2021.^1, 2^ Although there are strategies to manage herbicide resistance, such as herbicide rotation, the appearance of multiple-resistance weed species highlight the need for new modes of action.^3, 4^ Despite this need, over 30 years passed with no new herbicide mode of action until the very recent introduction of tetflupyrolimet that targets dihydroorotate dehydrogenase and cyclopyrimorate that targets homogentisate solanesyltransferase.^5, 6^

A typical approach for discovering new herbicides is screening large and diverse compound libraries to select the most active compounds for further investigation, including a detailed analysis of its mechanism of action which might be a known or new target.^3^ Another approach suggested as a useful starting point for herbicidal development is to exploit the physico-chemical connections between existing pharmaceuticals and herbicides and to use the principles of drug development to develop compounds towards a biological target of interest.^7^ The similarities between drugs and herbicides,^8, 9^ is highlighted by herbicides with antiprotozoal activity such as glyphosate,^10^ trifluralin^11^ and endothall,^12^ that are toxic to *Plasmodium* spp, and haloxyfop that is active against *T. gondii*, the causative agent of toxoplasmosis.^13^ Recent work has shown the converse to also be true, that many antiprotozoals, especially antimalarials, have herbicidal activity. Herbicidal antimalarial drugs include dihydrofolate reductase inhibitors,^14-16^ the non-mevalonate pathway inhibitor fosmidomycin^17^ and antimicrobial fluoroquinolones (e.g. ciprofloxacin) that act on DNA gyrase.^18^ Overall, these results suggest drug targets might be a source of inspiration for new herbicidal modes of action.

Unlike other eukaryotes, plants contain three genomes: nuclear, mitochondrial and plastid. The latter is present in chloroplasts, which are regarded as originating from symbiotic cyanobacteria and therefore its genome is homologous to prokaryotes.^19^ Although most chloroplast proteins are synthesized in the cytosol and imported by plastids, the chloroplast genome includes genes encoding essential parts of photosynthetic and translational machinery that are translated within the plastid.^20^ Therefore chloroplast translation is an essential process for plant viability, and its interruption interferes with embryo and seed development,^21^ causes chlorophyll deficiency,^22^ impairs cell division, root stem cell and leaf development,^23, 24^ reduces oxygen consumption and decreases fitness,^25^ and eventually leads to death of a plant^21, 22, 26^. Therefore, chloroplast translation machinery might be a viable target for herbicide development.

Despite differences in their regulatory pathways, the molecular mechanisms for chloroplast and bacterial translation are conserved^27^ so chloroplast translation can be inhibited by antibiotics that block microbial protein synthesis, e.g. spectinomycin and lincomycin.^28^ Virtually all information about chloroplast translation inhibitors such as puromycin, spectinomycin, lincomycin, chloramphenicol, streptomycin, erythromycin and anisomycin, and their effects on plants (as herbicides) is derived from studies with agar-grown plants,^21, 22, 25, 29-31^ and few studies report outcomes as foliar applications or pre-emergence with soil-grown plants.^32^ Although chloroplast translation is an essential plant process, it is not targeted by any commercial herbicide.^3^ Therefore, an opportunity exists to peruse this potentially new herbicidal mode of action by starting with antimicrobials that inhibit protein synthesis. To achieve this, we used the antimicrobial linezolid **1**, a known inhibitor of bacterial protein translation. Linezolid **1**, is potent against Gram-positive bacteria and a member of the oxazolidinone family that share an *N*-aryl substituted 2-oxazolidinone structural core and contain a (*S*)-methylene group in the position 2 (Fig. 1). Using **1** we demonstrate that its biological target possesses several advantages including the potential to control a broad spectrum of weeds.

**Fig. 1.**
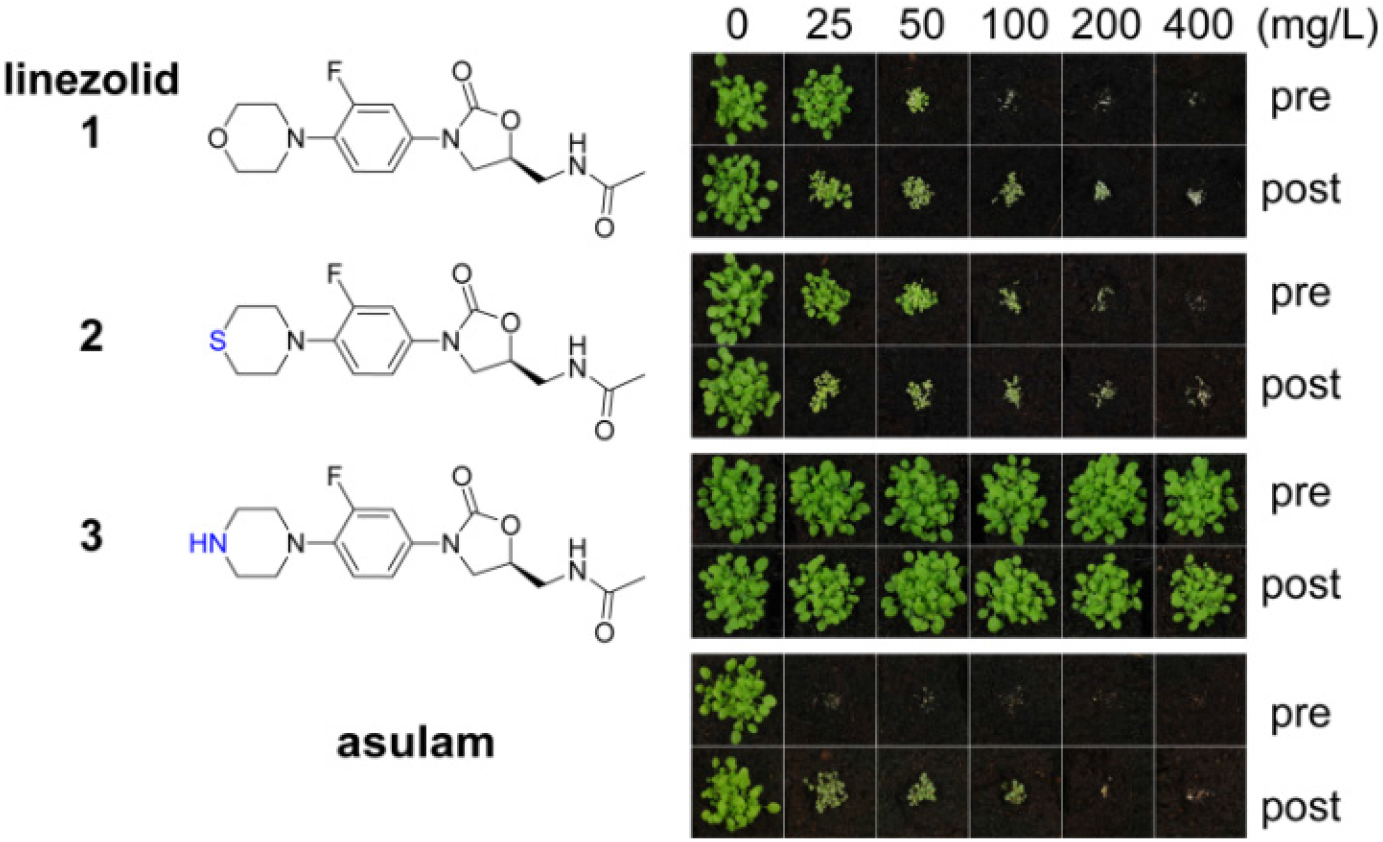
Oxazolidinones are herbicidal. Each compound was applied on *A. thaliana* seeds on soil (pre) or on seedlings 3 and 6 days after germination (post). The commercial herbicide asulam (inhibiting dihydropteroate synthase) was used as a positive control. Differences between linezolid **1**, sutezolid **2** and **3** are shown in blue.

## Results and Discussion

To investigate the herbicidal potential of oxazolidinones, in the first instance we used commercially available **1**. This compound is highly active *in vitro* against *S. aureus* (MIC 1 -8 µg/mL) and *M. tuberculosis* (MIC 0.125 -0.5 µg/mL),^33, 34^ approved for treating diseases caused by Gram-positive bacteria (e.g. *E. faecium* and *S. aureus*),^35^ and has potential for treating multidrug resistant tuberculosis.^36^ Assays against a model plant, *Arabidopsis thaliana*, were conducted by growing plants on soil and treating with a range of concentrations of **1** (25-400 mg/L) either pre-or post-emergence (Fig. 1).

Applying **1** as a post-emergence herbicide inhibited growth at the lowest dose 25 mg/L and was lethal at 100 mg/L. Pre-emergence activity was slightly lower with growth inhibition at 50 mg/L and death at 100 mg/L (Fig. 1). The leaves of plants exposed to **1** appeared chlorotic and desiccated if the compound was applied post-emergence, whereas pre-emergence treatment resulted in plants that emerged bleached. This efficiency was less than, but comparable to the commercial herbicide asulam that was lethal at all concentrations tested pre- and post-emergence. Sutezolid **2**, a thiomorpholine analogue of linezolid with similar antimicrobial effects,^37^ had activity similar to **1**. By contrast for **1** and **2**, the piperazyl analogue **3** of **1** was inactive (Fig. 1). An excellent antimicrobial, **3** displays only slightly lower activity than **1**^38^ so these results show the amino moiety of **3** plays a role either in uptake of the compound into the plant or in target binding. The bacterial mode of action and symptoms of the plants after exposure to **1** (chlorosis, necrosis), similar to photosystem I and photosystem II herbicides,^39^ were consistent with the mode of action of **1**, and indeed oxazolidinones, being disruption of chloroplast function.

To confirm **1** interferes with chloroplast function, its effects during exclusively heterotrophic growth or exclusively autotrophic growth were compared. *A. thaliana* seedlings were grown on **1** (10 µM) in glucose-deficient MS-agar medium under a long day regime to stimulate photosynthesis (autotrophic growth). Exclusively heterotrophic growth was stimulated by growth on glucose-supplemented MS-agar medium in the dark. Ciprofloxacin, an inhibitor of DNA gyrase,^18^ was used as a control as it causes bleaching, but does not directly affect chloroplast translation. Lincomycin, an inhibitor of chloroplast translation, was also used as a control as it affects translation in chloroplasts, but not in the cytosol.^31^ To assess growth inhibition we measured root length. Under autotrophic conditions all compounds reduced plant growth (Fig. 2). A clear difference in phenotype was observed by the cotyledon stage for autotrophic plants with **1** causing chlorosis and a significant decrease of root length (ESI Fig. 1). In contrast, neither seedlings nor roots were affected significantly by **1** for plants grown in the dark on glucose (Fig. 2, ESI Fig. 1). The chloroplast translation inhibitor lincomycin suppressed growth in an autotrophic, but not heterotrophic growth regime. A different pattern was observed for ciprofloxacin, which inhibited growth in both regimes. Therefore, the herbicidal activity of **1** is attributable to disruption of chloroplast function, because it only affects photosynthesising plants.

**Fig. 2.**
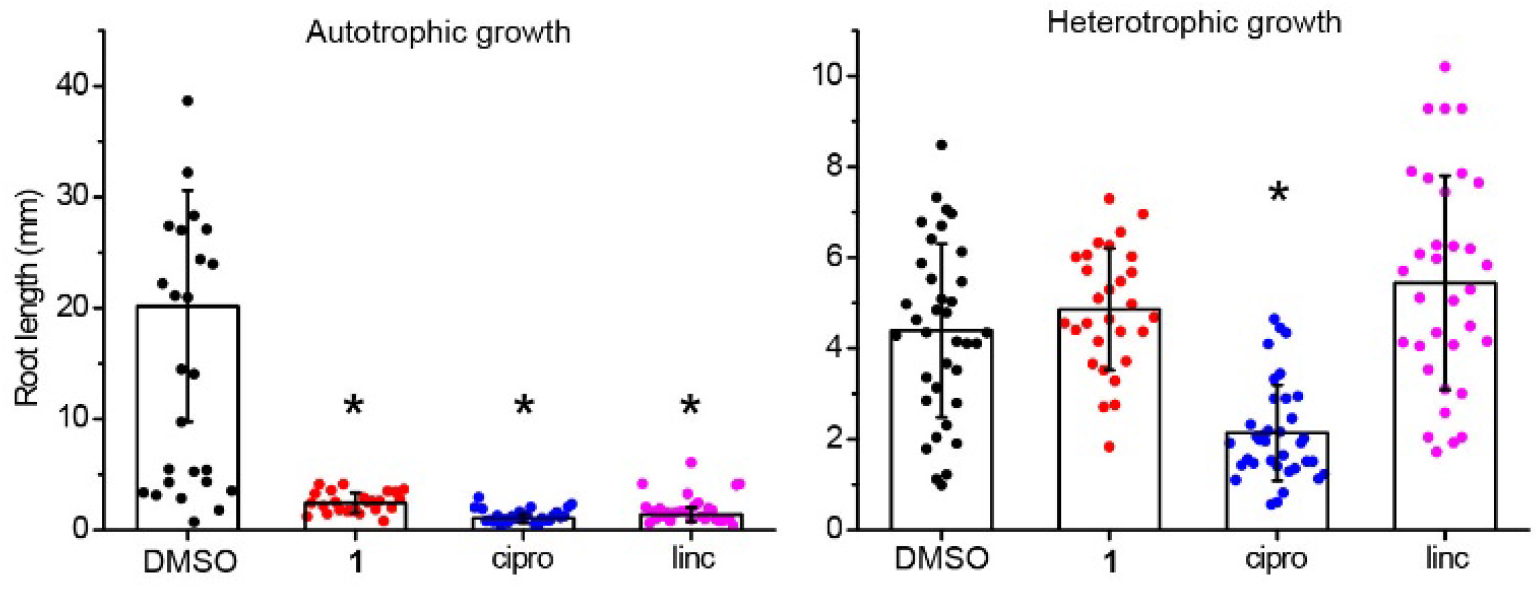
Linezolid 1 is inactive against dark-grown plants. Linezolid **1** was active in autotrophic (“auto”, no glucose in growth media, under light), but not heterotrophic (“hetero”, glucose in growth media, no light) growth regimes; ciprofloxacin (cipro), which does not directly affect chloroplast translation, was used as a positive control and inhibited plant growth in both conditions; lincomycin (linc), which inhibits chloroplast translation, was active only in autotrophic conditions.

As the symptoms of injury, due to **1**, (chlorosis and necrosis) can be caused by photosystem II inhibitors or inhibitors of chloroplast translation,^22, 39^ we investigated how **1** affects electron transfer and translation using *Pisum sativum* (pea) chloroplasts as an *in vitro* model. To assess protein translation we used a [^35^S]-methionine *in organelle* protein translation assay; translation was strongly inhibited by **1** in the concentration range of 25-100 µM with an efficiency exceeding spectinomycin, a known inhibitor of chloroplast translation (Fig. 3A). To assess how linezolid affects electron transfer in thylakoid membranes we used the Hill reaction, a colorimetric assay based on reduction of dichlorophenolindophenol (DCPIP), an artificial electron acceptor used to measure electron transfer during photosynthesis.^40, 41^ We found **1** did not affect the reduction of DCPIP (Fig. 3B), indicating **1** does not block electron transfer in the thylakoid membrane. To confirm DCPIP reduction distinguishes between inhibitors and non-inhibitors of photosynthesis we included controls atrazine (a photosystem II inhibitor, inhibits photosynthesis) and asulam (a dihydropteroate synthase inhibitor, does not inhibit photosynthesis). We found atrazine inhibited DCPIP reduction at 1 µM and completely blocked electron transfer at 100 µM, whereas asulam (like **1**) did not affect the electron transfer even at the highest tested concentration, distinguishing between inhibitors of photosynthesis and those that do not affect photosynthesis. Hence, **1** inhibits protein translation in chloroplasts analogous to its mode of action in bacteria.

**Fig. 3.**
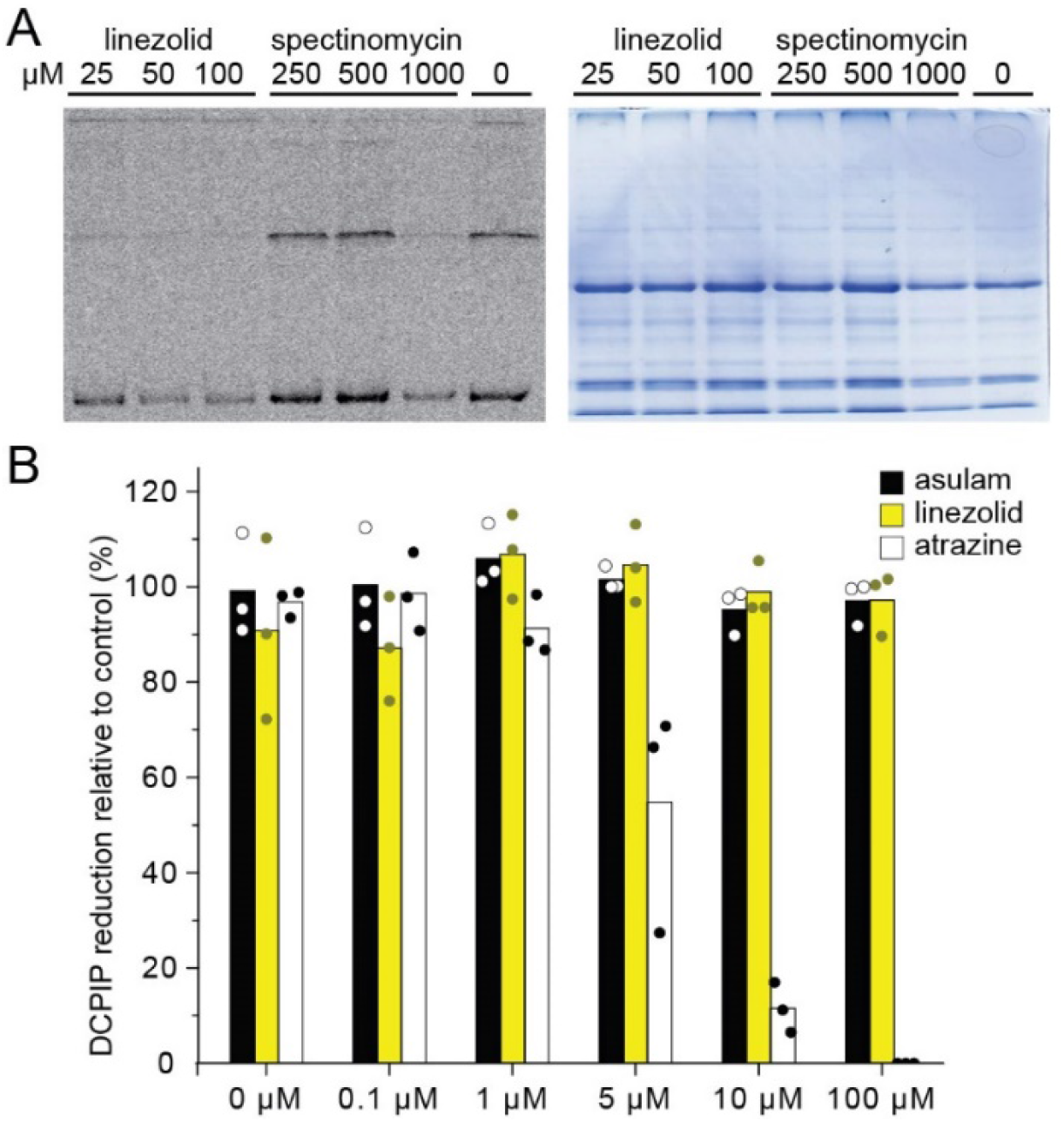
Linezolid 1 inhibits chloroplast translation but not electron transfer. (A) Inhibition of chloroplast translation as determined by [^35^S]-methionine incorporation assay (left) using **1** and spectinomycin. Linezolid **1** inhibited translation at a dose 1000x lower than spectinomycin. Coomassie gel loading control (right) is shown to demonstrate similar loading of each lane. (B) Hill reaction assays show that **1** did not inhibit electron transfer.

To assess the potential of **1** as a herbicidal scaffold we tested it against plant species, including monocots (*Brachypodium distachyon, Cynodon dactylon, Eragrostis tef, Triticum aestivum, Lolium rigidum*) and dicots (*P. sativum, Solanum lycopersicum, Ratibida columnifera*) at doses of 25 – 800 mg/L pre-and post-emergence (Fig. 4). We found **1** was active against all species pre- and post-emergence. Although the effective dose for **1** was ∼2-4 times greater than for *A. thaliana*, it had comparable activity to the glyphosate control against some species. The most susceptible species were *R. columnifera* and *C. dactylon*; these species were sensitive to **1** at 100 mg/L both pre- and post-emergence. The least sensitive plant was *T. aestivum*; visible growth inhibition was observed only at 800 mg/L. For other species the efficient dose was 200-800 mg/L.

**Fig. 4.**
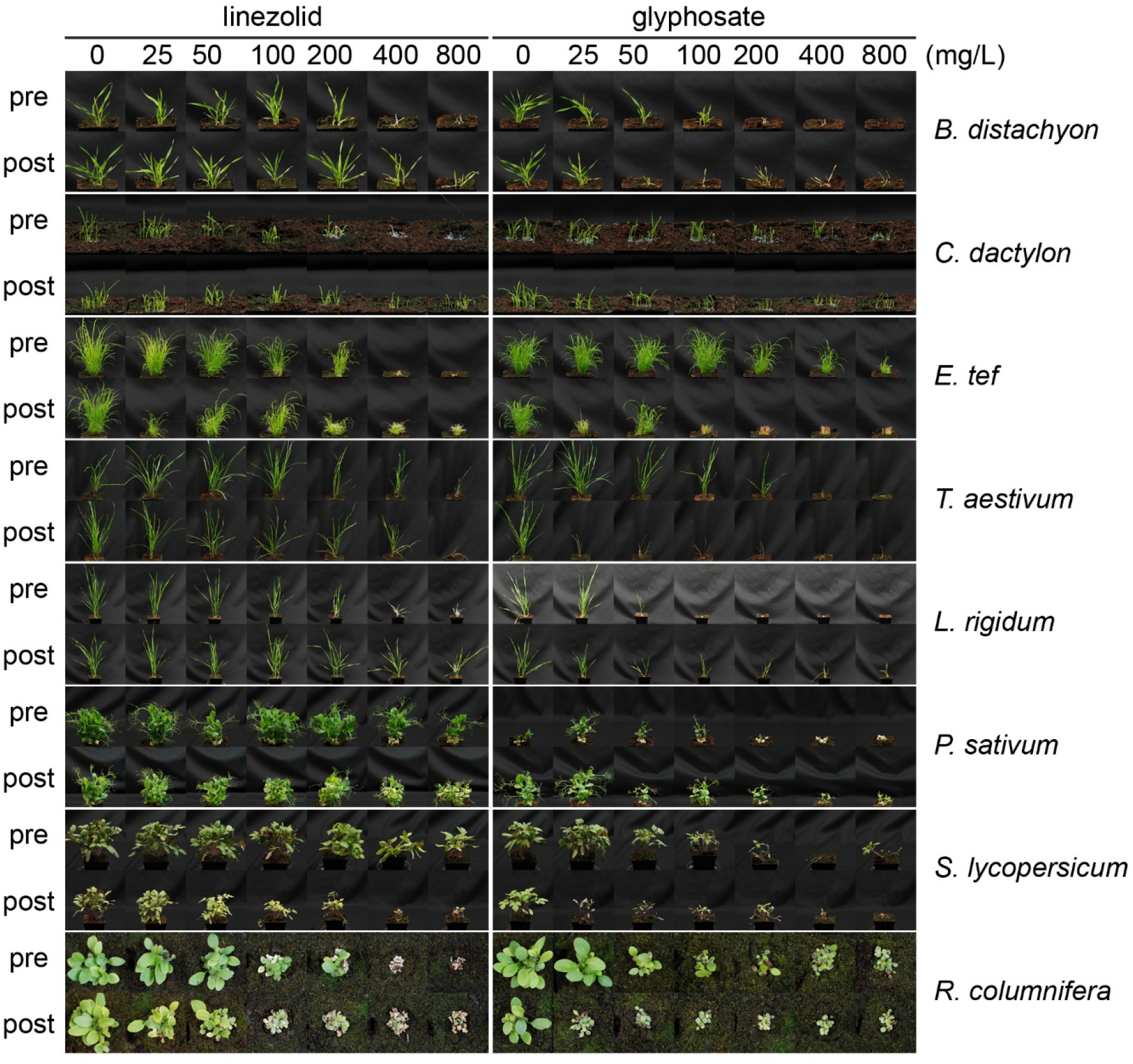
Linezolid is active against a wide range of plant species. Compounds were applied either to seeds on soil (pre) or to the surface of seedlings on the 3^rd^ and the 6^th^ day after emergence (post). Glyphosate was used as a positive control.

With **1** having good and wide-acting activity, we explored which of its moieties were required for activity and which could be modified and retain herbicidal, but not antibacterial activity. This is crucial as herbicides should be plant-specific and **1** is a valuable antibiotic. To make sure analogues of **1** possessed physico-chemical properties suitable for herbicides, we used an interactive database of physico-chemical data for 360 commercial herbicides (used previously for herbicide discovery^15, 16, 42, 43^) to choose, and then prepare analogues **4**-**57** (see ESI) that varied e.g. size, shape and hydrophobicity, but overall retained favourable physico-chemical properties for herbicides (Fig. 5 and ESI Fig. 2).

**Fig. 5.**
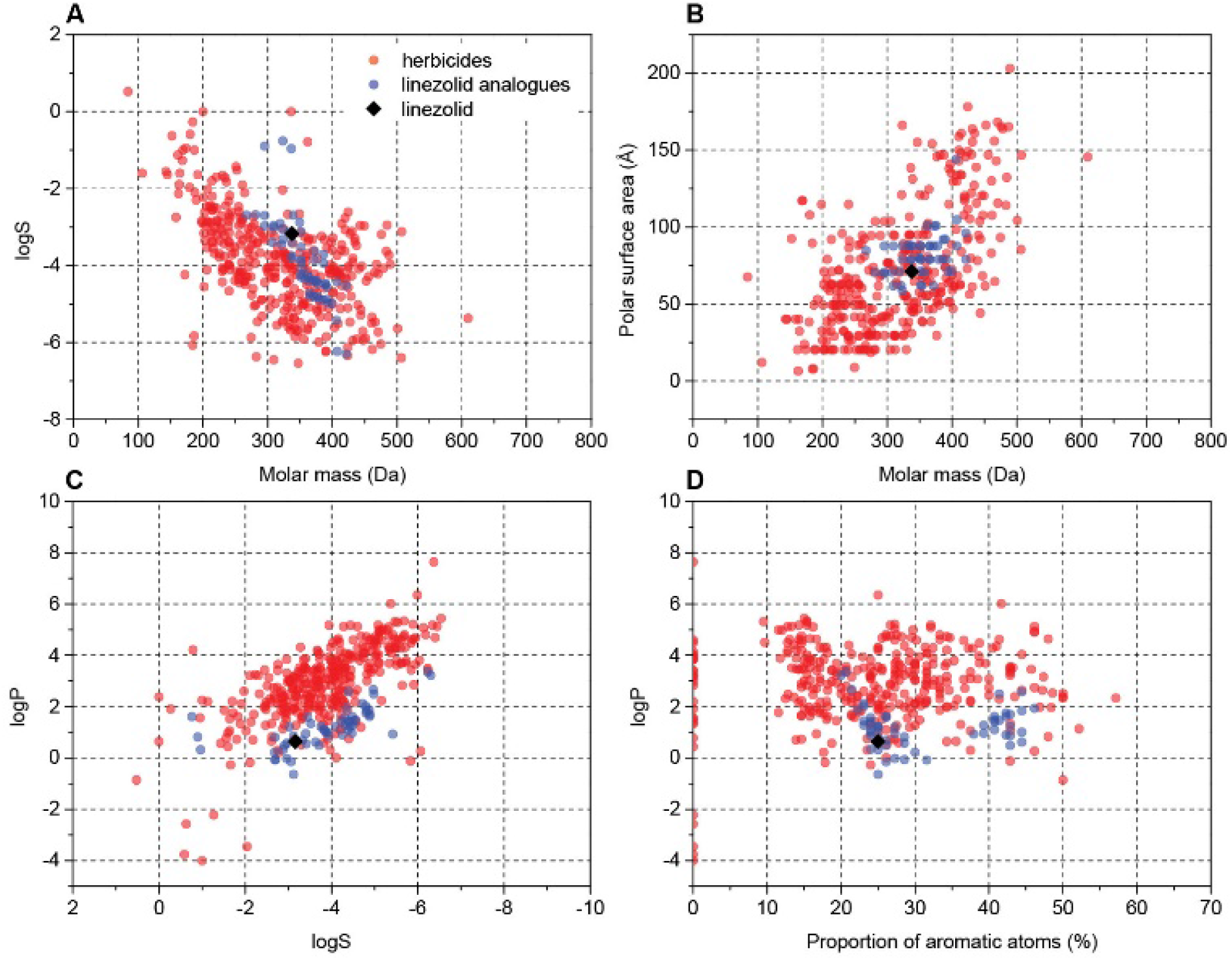
Examples of cluster analysis of physicochemical properties of linezolid 1 and prepared analogues versus known herbicides. Charts were extracted from an interactive database of physicochemical properties for commercial herbicides ^9, 43^ (red dots). Data points for this work were added (blue dots, **1** shown as black rhombus) and plotted with two properties compared (A) logS vs molar mass; (B) polar surface area vs molar mass; (C) logP vs logS; (D) logP vs proportion of aromatic atoms (see also ESI Fig. 2).

The overall structure of **1** contains chemically tractable motifs that were readily modified. Analogues were prepared where the core phenyloxazolidone was retained, but the piperazinyl, acetyl and fluoro groups were modified. A total of 54 analogues of **1** were synthesised (see Supporting Information) and then tested on soil for post-emergence activity (Fig. 6) using *A. thaliana*. Of the 54 analogues tested, many compounds lost herbicidal activity, but some interesting trends were observed. Extending the acetyl group of **1** with longer alkyl chains seemed to be tolerated for shorter chains (as in **5**) but activity was reduced across longer chains, as in **6**. Any other modification to the acetamido unit (as in **4, 7, 8** and **9**) abolished activity. The morpholinyl moiety also plays an important role in activity. Substituting this unit for small groups as in **12** and **21** retained activity at 200 mg/L, but longer and substituted acyl chains were not tolerated. Interestingly the formyl derivative **11** had no activity.

**Fig. 6.**
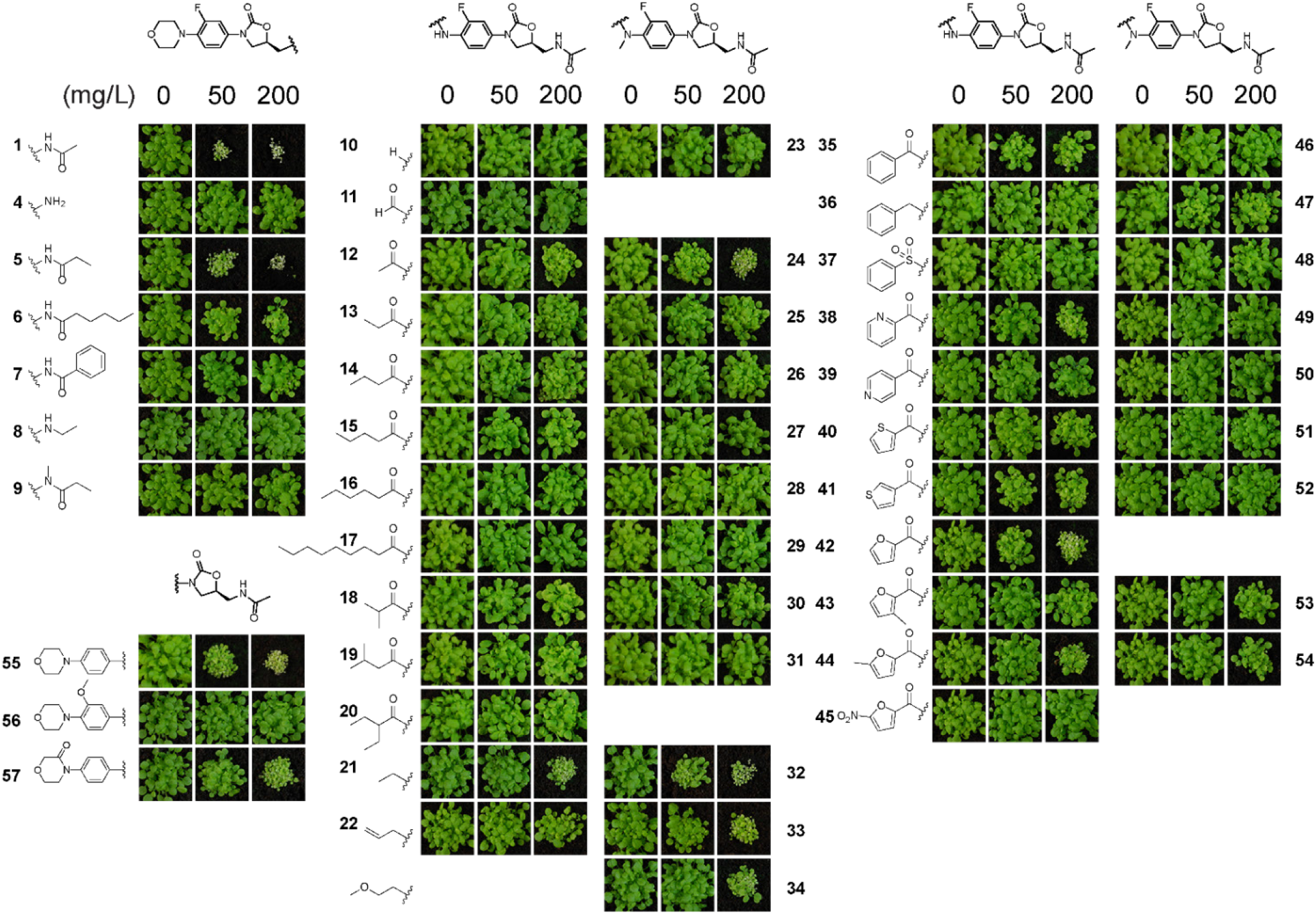
Screening of analogues 4-57 reveals insight into the post-emergent structure activity of linezolid 1. Compounds were prepared that possessed physicochemical properties as similar to commercial herbicides. Assays were performed as described in Fig. 1.

In the *N*-methyl series **23**-**34**, similar trends were observed with compounds **24, 32**-**34** all possessing good activity at 200 mg/L. Examination of the aryl amides, which bring planarity and potential fortuitous pi-stacking interactions, showed these compounds were far less tolerated. The benzoyl **35**, 2-pyridoyl **38**, 2-furoyl **42** and the 5’-methylfuroyl **44** derivatives possessed some activity at 200 mg/L but interestingly all other molecules were inactive at the concentrations tested. Of note is that the corresponding *N*-methyl derivatives lost all activity. Modification of the morpholinyl ring itself as a 3-morpholinone, as in **57**, also retained good activity at 200 mg/L. A simpler structure-activity relationship was observed when modifying the fluoro group. The desfluoro analogue **55** was still quite potent at 50 mg/L, but changing the group to a methoxy unit as in **56** abolished activity.

To assess if there was any improved selectivity gained for the prepared molecules as herbicides compared to antimicrobial activity, antibiotic susceptibility tests against *Bacillus subtilis* were performed for the compounds that retained herbicidal activity (Table 1). Activity against *B. subtilis* is relevant being that it is a common soil-borne bacterium and because **1** is more active against Gram-positive bacteria. All the synthesised analogues had zones of inhibition values less than linezolid **1** and interestingly, some compounds saw a considerable improvement in selectivity towards a herbicidal mode of action. Analogues **5** and **55**, which have similar herbicidal activities compared to **1**, showed a decrease in antimicrobial potency. Additionally, compounds **6, 24** and **34**, despite having a slight reduction in herbicidal potency compared to **1**, more importantly showed no antibacterial activity.

**Table 1.** Susceptibility of *B. subtilis* against the various linezolid analogues. Susceptibility determined in an agar diffusion assay using 6 mm filter disks loaded with 1.25 μg of compound. The zone of clearance was measured after incubation overnight. NE means “no effect”: there was no inhibition zone.

| Compound | Inhibition zone (mm) | Compound | Inhibition zone (mm) |
| --- | --- | --- | --- |
| **Linezolid 1** | 18.6 ± 0.4 | **34** | NE |
| **5** | 15.2 ± 1.0 | **35** | NE |
| **6** | NE | **38** | 14.5 ± 1.0 |
| **12** | NE | **42** | 12.5 ± 0.5 |
| **21** | 13.7 ± 0.5 | **44** | 12.9 ± 0.4 |
| **24** | NE | **55** | 14.1 ± 0.9 |
| **32** | 15.2 ± 1.1 | **57** | NE |
| **33** | 12.7 ± 0.7 |  |  |

## Conclusions

No commercial herbicide targets chloroplast protein translation. To evaluate if chloroplast translation might be a viable target we examined the oxazolidinone molecular class of microbial translation inhibitors as herbicides, using **1** as an exemplar and found this and other oxazolidinones were active against soil-grown plants (Fig. 1). Although translation in the cytosol differs from the chloroplast, some prokaryotic translation inhibitors, such as streptomycin, inhibit both cytosolic and chloroplast translation.^31^ Linezolid **1** and lincomycin specifically inhibit chloroplast translation as plants grow normally in a heterotrophic regime, but growth is reduced in an autotrophic regime (Fig. 2). The ability of **1** to selectively inhibit chloroplast protein translation was also confirmed using a labelled methionine incorporation assay, as **1** strongly inhibited protein translation at a concentration 1000 times lower than spectinomycin, but did not inhibit electron transfer at any tested rate of application, as confirmed by the DCPIP assay that measures electron transfer in thylakoid membranes.

The utility of **1** was also demonstrated in its broad-spectrum herbicidal activity against a wide range of plant species where in all cases it was effective. Despite its promise, we cannot consider **1** itself as a potential herbicide unless its antibacterial and herbicidal activities are divorced from each other. To that end we explored which moieties of **1** are required for herbicidal activity, taking advantage of a physicochemical database^9, 43^ to prepare analogues of **1** that retain suitable herbicidal properties. We demonstrate the critical elements that retain herbicidal activity but also show, using antimicrobial susceptibility assays, that separation of antibacterial and herbicidal activities was possible exemplified by **6, 24** and **34**. Overall these results demonstrate that inhibiting chloroplast translation is a viable herbicidal mode of action and the structure of **1** could be used as a scaffold to create herbicides that target this important plant process.

## Supporting information

Supporting Figures 1-2, General Experimental

Supporting Dataset - NMR spectra

## Author Contributions

K.V.S., J.S.M. and K.A.S. conceived the study and analysed data. K.V.S. performed experiments validating the target of **1** and initial herbicidal analysis. K.J.B. and K.A.S. performed chemical synthesis and K. J. B. conducted herbicidal analysis of prepared compounds. K.V.S. and A.W.D. performed antimicrobial assays. M.W.M. extracted organelles and performed translation assays. K.V.S., K.A.S. and J.S.M. wrote the manuscript with help from all authors.

## Acknowledgements

The authors thank Philippe Hervé and Bruce Lee from Nexgen Plants for fruitful discussions. The authors are grateful to Aneta Ivanova and Mabel Gill-Hille for assistance establishing translation assays and to Maxime Corral and Joel Haywood for help establish *A. thaliana* herbicide assays. K.V.S. was supported by a Research Training Program Stipend and a Research Training Program Fee Offset. This work was funded in part by Nexgen Plants and in part by an Australian Research Council (ARC) Discovery Project (DP190101048) to J.S.M. and

K.A.S. M.W.M. was supported by ARC Future Fellowship FT130100112.

## Experimental procedures

### Source of commercial oxazolidinones

Linezolid **1** was purchased from Fluorochem (United Kingdom). Sutezolid **2** and the piperazyl analogue **3** were purchased from Angene Chemical (United Kingdom). Compound 55 was purchased from Carbosynth (United Kingdom).

### Herbicidal activity assay

The herbicidal activity of compounds was determined based on the published method from.^16^ *A. thaliana* Col-0 seeds (∼30) were sown in 63 × 63 × 59 mm pots consisting of Irish peat that was pre-wet prior to sowing. Seeds were cold-treated for 3 days in the dark at 4°C to synchronise germination and then grown in a chamber at 22°C, with 60% relative humidity and in a 16 h light/8 h dark photoperiod. Compounds were initially dissolved in dimethyl sufloxide (DMSO) at 20 mg/mL and further diluted in water prior to treatments. The surfactant Brushwet (SST Australia) was added to a final concentration of 0.02%. The carrier DMSO was used as a negative control. Seeds or seedlings were treated with 500 µL of 0, 25, 50, 100, 200 or 400 mg/L solutions that contained 2% of DMSO, using a pipette. Pre-emergence, treatments were given at day zero as trays were moved into their first long day, whereas post-emergence treatments were done at three and six days after germination. Seedlings were grown for 16 days after treatment before photos were taken.

### Herbicidal activity assay in etiolated plants

Ethanol-sterilised *A. thaliana* Col-0 seeds were resuspended in 1 g/L agar and stratified for three days before 30-40 seeds were sown on 10 cm square Petri-dishes contained 10 g/L agar medium with 4 g/L of Murashige-Skoog salt and 10 µM of **1**. Ciprofloxacin at 10 µM and lincomycin at 500 µM were used as controls. For etiolated plants the medium additionally contained 10 g/L of glucose and plates were grown vertically. Control plants were treated in the same manner, but the medium did not contain **1**. After ten days of incubation pictures were taken and root length was measured by ImageJ software and compared to control samples using one-way ANOVA test (p = 0.05, n = 20).

### Isolation of chloroplasts

Chloroplasts were isolated from *P. sativum* according to published methods.^44^ Peas were sown in vermiculite, grown for 10-12 days under long day conditions (16 h light/8 h dark, 9660 lux) at 26°C and 60% relative humidity. Prior to isolation, plants underwent dark treatment for 14-16 h to prevent starch accumulation. All subsequent operations were carried out at 4°C. Approximately 100 g of leaves were ground in 300 mL of grinding buffer (0.33 M sorbitol, 50 mM HEPES buffer, pH 8.0, 0.45 mM ascorbic acid, 1 mM magnesium chloride, 1 mM manganese chloride and 2 mM EDTA disodium salt). The extract was filtered through two layers of miracloth that was pre-wet in grinding buffer, and the filtrate centrifuged at 2,000 *g* for 5 min to pellet both intact and broken plastids. The supernatant was decanted and the plastid pellet resuspended in the residual supernatant using a soft painting brush and loaded onto 50% percoll-based density gradient prepared with grinding buffer as previously described.^44^ The loaded gradient was centrifuged at 12,100 *g* for 10 min and the higher band of broken plastids was discarded by vacuum aspiration, whereas a lower dark-green band of intact chloroplasts was collected by a disposable Pasteur pipette, mixed with three volumes of 0.33 M sorbitol, 50 mM HEPES buffer (pH 8.0) and centrifuged at 2,000 *g* for 4 min to pellet intact plastids. The supernatant was decanted and plastids resuspended in the residual liquid. The plastid suspension was kept on ice and used immediately for *in vitro in organello* translation and Hill reaction assays.

### Thylakoid membrane electron transfer assay

The inhibitory effect of different compounds on the rate of electron transport in thylakoid membranes of intact chloroplasts was measured in an assay based on the reduction of dichlorophenolindophenol by the plastid electron transfer chain.^40, 41^ In a typical experiment 18 µL of intact chloroplast suspension was added to 1800 µL of a reaction mixture containing 40 µM dichlorophenolindophenol, 0.01-100 µM asulam, atrazine or linezolid and 0.05% ethanol in 0.33 M sorbitol, 50 mM HEPES buffer (pH 8.0), which was then mixed by gentle inversion of the reaction tube and shook by hand under light of a halogen bulb lamp (70W, 6110 lux) for 10 min at room temperature. After this incubation, the samples were centrifuged at 500 *g* for 1 min and the absorbance at 600 nm recorded. Photosynthetic activity was defined as a relative decrease of absorbance at 600 nm which corresponds to the oxidized state of dichlorophenolindophenol. All measurements were repeated three times and averaged. Control samples with no compound treatment were also prepared.

### *In vitro* in organelle protein translation assay

Chloroplast protein translation assays were carried out as previously described.^45^ In brief, 10 µL of isolated chloroplasts was added to 50 µL of reaction mixture containing 40 µM of each amino acid except for methionine, 10 mM DTT, 10 mM ATP and 10 mM magnesium chloride, 10 µCi of [^35^S]-methionine and 25-100 µM of **1** or 250-1000 µM of spectinomycin in buffer (0.33 M sorbitol, 50 mM HEPES buffer, pH 8.0). The reactions were then incubated for 10 min at 26°C with gentle shaking before the reaction was stopped by addition of 1 mL of 10 mM methionine in cold buffer. To assess the amount of synthesized protein, 45 µL aliquots of the reaction were taken and denatured by boiling for 5 min with 20 g/L sodium dodecyl sulfate, 100 g/L sucrose and 0.3 g/L bromophenol blue. The proteins were then separated by 16% polyacrylamide SDS-PAGE analysis, stained with Coomassie dye and dried under vacuum. Radioactive gels were exposed to a BAS phosphor-imaging plate (TR2040, Fuji) for 24 h and visualised using a Typhoon scanner (GE Healthcare, FLA-9500).

### Antibacterial assay

Antibacterial activity of compounds was tested against *Bacillus subtilis* (strain Marburg 168) using a paper-disk diffusion method.^46^ The starting culture of bacteria was obtained by inoculation of 2 mL of sterile Luria-Bertani (LB) broth with glycerol stock of *B. subtilis*, followed by overnight incubation in shaker incubator at 37°C/180 rpm. After this time, an aliquot (0.5 mL) was added to 4 mL of sterile LB and grown at 37°C/180 rpm for ∼ 3 h, until A_600_ ∼0.5 was reached. At this time the bacterial culture was diluted 5-fold with sterile LB, and the culture spread evenly over the surface of pre-warmed (37°C) LB-agar plates using a sterile cotton swab. The surface of the agar was allowed to dry for 10-15 min before sterile paper disks (6 mm diameter, Whatman) containing 1.25 µg of compound or 8 µg of kanamycin were placed onto the agar surface. The disks were gently flattened with forceps to ensure full contact with agar surface. The plates were then incubated upside down at 37°C for 24 h before the diameter of the zone of inhibition was measured. Determinations were performed in triplicate and data presented as the mean of the measurements with the error given as the standard error of the mean.

