## Supporting Figures 1-2, General Experimental for "Inhibition of chloroplast translation as a new target for herbicides"

### Inhibition of chloroplast translation as a new mode of action for herbicides

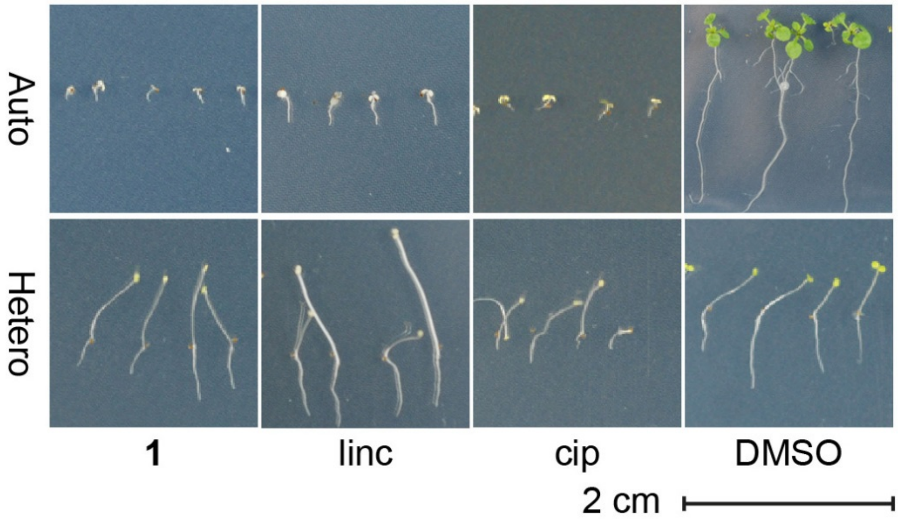


**Supporting Figure 1 |** Linezolid **1**, similarly to the known chloroplast translation inhibitor lincomycin (linc) causes bleaching and root length reduction, only in photosynthesising plants, unlike ciprofloxacin (cip) that affects plants regardless of photosynthesis.

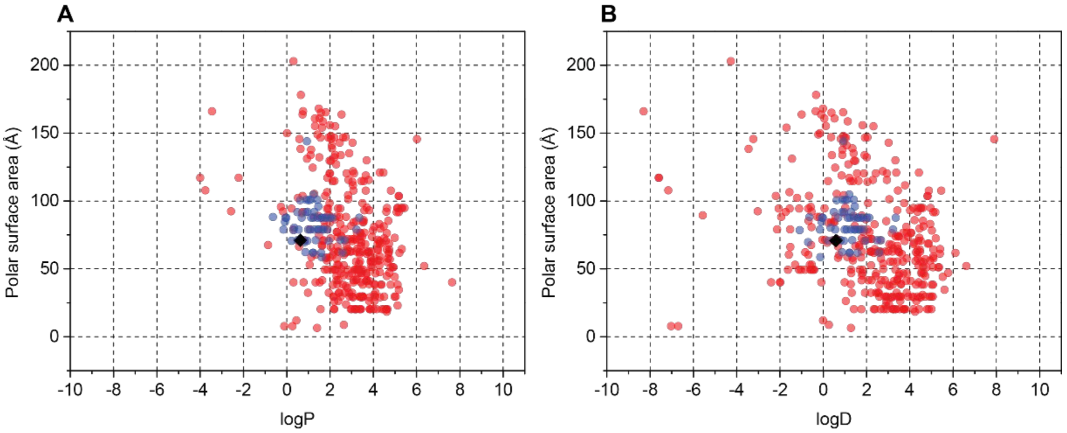


**Supporting Figure 2 |** Examples of cluster analysis of physicochemical properties of linezolid **1** and prepared analogues versus known herbicides. Charts were extracted from an interactive database containing the physicochemical properties of commercial herbicides. (A) polar surface area vs partition coefficient (logP); (B) polar surface area vs distribution coefficient (logD). Linezolid **1** is displayed as a black rhombus, blue dots represent the analogues prepared and the red dots represent the commercial herbicides.

**General Experimental**

All reagents and materials were purchased from commercial suppliers. Compounds **10** and **23** were purchased from SYNthesis Med Chem. Thin layer chromatography (TLC) was affected on Merck silica gel 60 F254 aluminium-backed plates and spots stained by heating with vanillin dip (6 g vanillin, 1 mL conc. $H_2SO_4$, 100 mL ethanol), unless stated otherwise. Flash column chromatography was performed on Merck silica gel using the specified solvents. NMR spectra were obtained on a Bruker Avance IIIHD 400, 500 or 600 spectrometers. The solvents used were $CDCl_3$ or DMSO-$d_6$ with $CHCl_3$ ($^1$H, δ 7.26 ppm), $CDCl_3$ ($^{13}$C, δ 77.16 ppm), $CD_3S(O)CD_2H$ ($^1$H, δ 2.50 ppm) or $(CD_3)_2SO$ ($^{13}$C, δ 39.52 ppm) used as an internal standard. Infrared spectra were obtained with neat samples on a PerkinElmer spectrum one FT-IR spectrometer fitted with a PerkinElmer Universal Attenuated Total Reflectance (ATR) sampling accessory. High resolution mass spectra (HR-MS) were obtained on a Waters LCT Premier XE TOF spectrometer, run in W-mode, using either the ESI or APCI equipped ion source, in positive or negative mode.

**Preparation of *N*-acyl analogues**

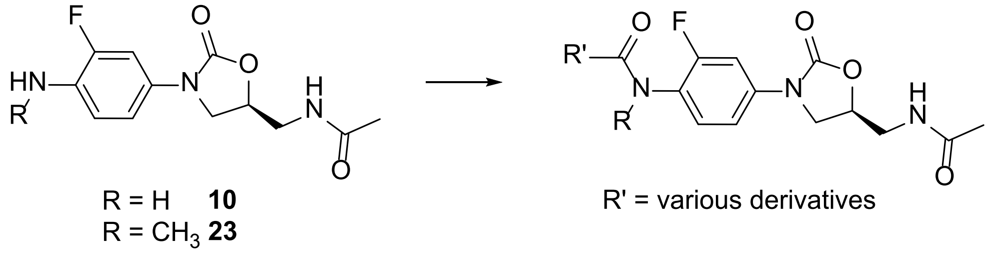


***General Procedure A***

To a stirred suspension of **10** or **23** (0.18 mmol, 1.0 equiv) in $CH_2Cl_2$ (1.5 mL) was added pyridine (0.36 mmol, 2.0 equiv) and the acid anhydride (0.36 mmol, 2.0 equiv). The mixture was stirred at r.t. for 18 h, then quenched with saturated aqueous $NaHCO_3$ (5 mL) and extracted with $CH_2Cl_2$ (3 x 5 mL), dried ($MgSO_4$), filtered and concentrated under reduced pressure. The residue was purified by silica gel chromatography (0-3% MeOH/$CH_2Cl_2$) to yield the compound of interest.

***General Procedure B***

To a stirred suspension of **10** or **23** (0.18 mmol, 1.0 equiv) in $CH_2Cl_2$ (1.5 mL) was added pyridine (0.36 mmol, 2.0 equiv) and the acid chloride (0.36 mmol, 2.0 equiv). The mixture was stirred at r.t. for 2 h, then quenched with saturated aqueous $NaHCO_3$ (5 mL) and extracted with $CH_2Cl_2$ (3 x 5 mL), dried ($MgSO_4$), filtered and concentrated under reduced pressure. The residue was purified by silica gel chromatography (0-3% $MeOH/CH_2Cl_2$) to yield the compound of interest.

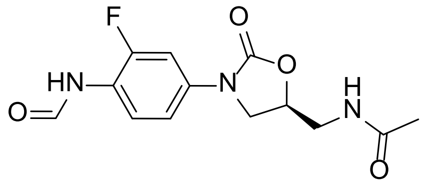


***N*-[[(5*S*)-3-[3-fluoro-4-(formylamino)phenyl]-2-oxo-5-oxazolidinyl]methyl]acetamide 11**

A stirred solution of formic acid (90%, 0.065 mL, 1.7 mmol) and acetic anhydride (0.095 mL, 1.0 mmol) was stirred under $N_2$ at 70 °C for 1 h, then allowed to cool to r.t. At this time **10** (53 mg, 0.20 mmol) was then added and the resultant mixture heated at 50 °C for 0.5 h, then $CH_2Cl_2$ (1 mL) was added and heating was continued for 0.5 h. The residue was purified by silica gel chromatography (0-10% $MeOH/CHCl_3$), to yield the title compound **11** as a white powder (32 mg, 54%). $R_f$ 0.30 (10% $MeOH/CHCl_3$); $^1H$ NMR (500 MHz, DMSO-$d_6$): major rotamer δ 10.08 (s, 1H), 8.28 (s, 1H), 8.24 (bt, 1H), 8.03 (dd, *J* = 8.9, 8.8 Hz), 7.60 (dd, *J* = 13.2, 2.0 Hz, 1H), 7.23 (dd, *J* = 8.8, 2.0 Hz, 1H), 4.75-4.70 (m, 1H), 4.10 (dd, *J* = 8.9, 8.8 Hz, 1H), 3.72 (dd, *J* = 8.8, 6.7 Hz. 1H), 3.42-3.40 (m, 2H), 1.83 (s, 3H); $^{13}C$ NMR (126 MHz, DMSO-$d_6$): major rotamer δ 170.1, 159.9, 154.1, 152.6 (d, *J* = 243 Hz), 135.4 (d, *J* = 10 Hz), 123.3 (d, *J* = 3 Hz), 120.8 (d, *J* = 12 Hz), 113.5 (d, *J* = 3 Hz), 105.5 (d, *J* = 25 Hz), 71.7, 42.3, 41.4, 22.5; FTIR (ATR): υ 3297, 1724, 1680, 1651, 1528 $cm^{-1}$; HR-MS (ESI) *m/z*: found 296.1047; calculated for $C_{13}H_{15}N_3O_4F$ $[M+H]^+$ 296.1047.

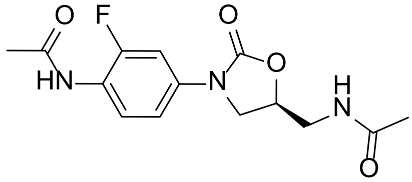


**(*S*)-*N*-(4-(5-(acetamidomethyl)-2-oxooxazolidin-3-yl)-2-fluorophenyl)acetamide 12**

The title compound was prepared according to General Procedure A to give **12** as an off white powder (51 mg, 92%). $R_f$ 0.25 (5% $MeOH/CHCl_3$); $^1H$ NMR (600 MHz, DMSO-$d_6$): δ 9.69 (s, 1H), 8.24 (t, *J* = 5.7 Hz, 1H), 7.80 (dd, *J* = 8.8, 8.9 Hz, 1H), 7.56 (dd, *J* = 13.2, 2.2 Hz, 1H), 7.21 (dd, *J* = 8.8, 2.2 Hz, 1H), 4.74-4.70 (m, 1H), 4.10 (dd, *J* = 8.9, 8.9 Hz, 1H), 3.72 (dd, *J* = 8.9, 6.5 Hz, 1H), 3.41 (app t, *J* = 5.5 Hz, 2H), 2.06 (s, 3H), 1.83 (s, 3H); $^{13}C$ NMR (151 MHz, DMSO-$d_6$): δ 170.0, 168.6, 154.0, 153.6 (d, *J* = 244 Hz), 135.5 (d, *J* = 10 Hz), 124.8, 121.5 (d, *J* = 12 Hz), 113.3 (d, *J* = 3 Hz), 105.4 (d, *J* = 25 Hz), 71.6, 47.2, 41.4,

23.3, 22.4; FTIR (ATR): υ 3334, 1749, 1661, 1539 $cm^{-1}$; HR-MS (ESI) *m/z*: found 310.1202; calculated for $C_{14}H_{17}FN_3O_4$ $[M+H]^+$ 310.1203.

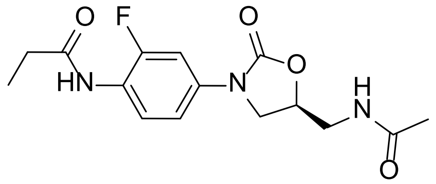

**(*S*)-*N*-(4-(5-(acetamidomethyl)-2-oxooxazolidin-3-yl)-2-fluorophenyl)propanamide 13**

The title compound was prepared according to General Procedure A to give **13** as a beige powder (54 mg, 94%). $R_f$ 0.28 (5% MeOH/$CHCl_3$); $^1H$ NMR (500 MHz, DMSO-$d_6$): δ 9.60 (s, 1H), 8.24 (t, *J* = 5.8 Hz, 1H), 7.80 (dd, *J* = 8.9, 8.9 Hz, 1H), 7.56 (dd, *J* = 13.2, 2.4 Hz, 1H), 7.22 (dd, *J* = 8.9, 2.4 Hz, 1H), 4.75-4.69 (m, 1H), 4.10 (dd, *J* = 9.0, 9.0 Hz, 1H), 3.72 (dd, *J* = 9.0, 6.5 Hz, 1H), 3.41 (app t, *J* = 5.5 Hz, 2H), 2.36 (q, *J* = 7.5 Hz, 2H), 1.83 (s, 3H), 1.07 (t, *J* = 7.5 Hz, 3H); $^{13}C$ NMR (126 MHz, DMSO-$d_6$): δ 172.3, 170.0, 154.0, 153.7 (d, *J* = 244 Hz), 135.5 (d, *J* = 10 Hz), 124.9, 121.5 (d, *J* = 12 Hz), 113.3 (d, *J* = 3 Hz), 105.4 (d, *J* = 25 Hz), 71.6, 47.2, 41.4, 28.8, 22.4, 9.6; FTIR (ATR): υ 3301, 1737, 1662, 1532 $cm^{-1}$; HR-MS (ESI) *m/z*: found 324.1358; calculated for $C_{15}H_{19}FN_3O_4$ $[M+H]^+$ 324.1360.

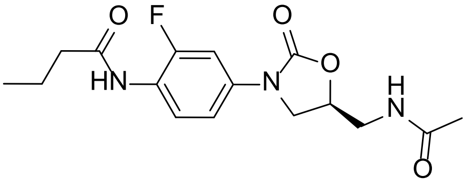

**(*S*)-*N*-(4-(5-(acetamidomethyl)-2-oxooxazolidin-3-yl)-2-fluorophenyl)butanamide 14**

The title compound was prepared according to General Procedure A to give **14** as an off-white powder (49 mg, 82%). $R_f$ 0.30 (5% MeOH/$CHCl_3$); $^1H$ NMR (500 MHz, DMSO-$d_6$): δ 9.61 (s, 1H), 8.24 (t, *J* = 5.8 Hz, 1H), 7.78 (dd, *J* = 8.9, 8.9 Hz, 1H), 7.56 (dd, *J* = 13.2, 2.4 Hz, 1H), 7.22 (dd, *J* = 8.9, 2.4 Hz, 1H), 4.75-4.70 (m, 1H), 4.10 (dd, *J* = 9.1, 9.0 Hz, 1H), 3.72 (dd, *J* = 9.1, 6.5 Hz, 1H), 3.41 (app t, *J* = 5.5 Hz, 2H), 2.32 (t, *J* = 7.3 Hz, 2H), 1.83 (s, 3H), 1.59 (tt, *J* = 7.4, 7.3 Hz, 2H), 0.91 (t, *J* = 7.4 Hz, 3H); $^{13}C$ NMR (126 MHz, DMSO-$d_6$): δ 171.4, 170.0, 154.0, 153.8 (d, *J* = 244 Hz), 135.6 (d, *J* = 10 Hz), 125.1, 121.5 (d, *J* = 12 Hz), 113.3 (d, *J* = 3 Hz), 105.4 (d, *J* = 25 Hz), 71.6, 47.2, 41.4, 37.5, 22.4, 18.6, 13.6; FTIR (ATR): υ 3268, 1727, 1660, 1532 $cm^{-1}$; HR-MS (ESI) *m/z*: found; calculated for $C_{16}H_{21}FN_3O_4$ $[M+H]^+$ 338.1516.

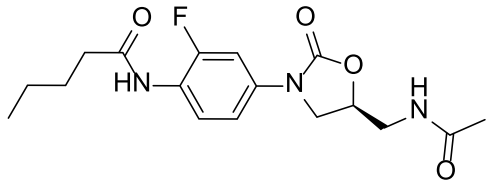

**(*S*)-*N*-(4-(5-(acetamidomethyl)-2-oxooxazolidin-3-yl)-2-fluorophenyl)pentanamide 15**

The title compound was prepared according to General Procedure A to give **15** as a white powder (38 mg, 60%). $R_f$ 0.13 (5% MeOH/ $CH_2Cl_2$); $^1$H NMR (600 MHz, DMSO-$d_6$): δ 9.61 (s, 1H), 8.24 (t, *J* = 5.8 Hz, 1H), 7.77 (dd, *J* = 9.0, 9.0 Hz, 1H), 7.55 (dd, *J* = 13.2, 2.4 Hz, 1H), 7.21 (dd, *J* = 9.0, 2.4 Hz, 1H), 4.74-4.70 (m, 1H), 4.10 (dd, *J* = 9.0, 9.0 Hz, 1H), 3.72 (dd, *J* = 9.0, 6.6 Hz, 1H), 3.41 (app t, 5.5 Hz, 2H), 2.34 (t, *J* = 7.5 Hz, 2H), 1.83 (s, 3H), 1.56 (tt, *J* = 7.5, 7.5 Hz, 2H), 1.32 (tq, *J* = 7.5, 7.5 Hz, 2H), 0.89 (t, *J* = 7.5 Hz, 3H); $^{13}$C NMR (151 MHz, DMSO-$d_6$): δ 171.6, 170.0, 154.0, 153.8 (d, *J* = 244 Hz), 135.6 (d, *J* = 11 Hz), 125.1, 121.5 (d, *J* = 12 Hz), 113.3 (d, *J* = 3 Hz), 105.5 (d, *J* = 26 Hz), 71.6, 47.2, 41.4, 35.4, 27.3, 22.4, 21.8, 13.8; FTIR (ATR): υ 3289, 1727, 1659, 1532 $cm^{-1}$; HR-MS (ESI) *m/z*: found 374.1478; calculated for $C_{17}H_{22}FN_3O_4Na$ $[M+Na]^+$ 374.1492.

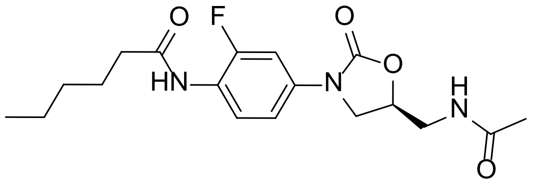

**(*S*)-*N*-(4-(5-(acetamidomethyl)-2-oxooxazolidin-3-yl)-2-fluorophenyl)hexanamide 16**

The title compound was prepared according to General Procedure A to give **16** as a white powder (35 mg, 51%). $R_f$ 0.13 (5% MeOH/ $CH_2Cl_2$); $^1$H NMR (600 MHz, DMSO-$d_6$): δ 9.61 (s, 1H), 8.25 (t, *J* = 5.9 Hz, 1H), 7.77 (dd, *J* = 8.9, 8.9 Hz, 1H), 7.55 (dd, *J* = 13.1, 2.5 Hz, 1H), 7.21 (dd, *J* = 8.9, 2.5 Hz, 1H), 4.74-4.70 (m, 1H), 4.10 (dd, *J* = 9.0, 9.0 Hz, 1H), 3.72 (dd, *J* = 9.0, 6.6 Hz, 1H), 3.41 (app t, 5.5 Hz, 2H), 2.33 (t, *J* = 7.4 Hz, 2H), 1.83 (s, 3H), 1.58 (tt, *J* = 7.4, 7.3 Hz, 2H), 1.37-1.25 (m, 4H), 0.87 (t, *J* = 7.0 Hz, 3H); $^{13}$C NMR (151 MHz, DMSO-$d_6$): δ 171.6, 170.0, 154.0, 153.8 (d, *J* = 243 Hz), 135.6 (d, *J* = 11 Hz), 125.0, 121.5 (d, *J* = 12 Hz), 113.3 (d, *J* = 3 Hz), 105.5 (d, *J* = 26 Hz), 71.6, 47.2, 41.4, 35.6, 30.9, 24.8, 22.4, 21.9, 13.9; FTIR (ATR): υ 3269, 1740, 1662, 1531 $cm^{-1}$; HR-MS (ESI) *m/z*: found 366.1819; calculated for $C_{18}H_{25}FN_3O_4$ $[M+H]^+$ 366.1829.

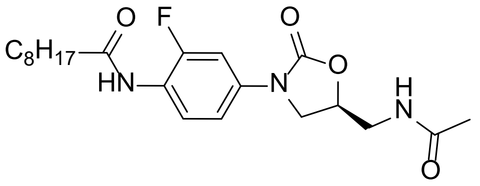

**(*S*)-*N*-(4-(5-(acetamidomethyl)-2-oxooxazolidin-3-yl)-2-fluorophenyl)nonanamide 17**

The title compound was prepared according to General Procedure B to give **17** as a white powder (49 mg, 67%). $R_f$ 0.10 (5% MeOH/ $CH_2Cl_2$); $^1$H NMR (400 MHz, DMSO-$d_6$): δ 9.60 (s, 1H), 8.24 (t, *J* = 5.8 Hz, 1H), 7.77 (dd, *J* = 8.9, 8.9 Hz, 1H), 7.55 (dd, *J* = 13.2, 2.4 Hz, 1H), 7.21 (dd, *J* = 8.9, 2.4 Hz, 1H), 4.75-4.69 (m, 1H), 4.10 (dd, *J* = 9.0, 9.0 Hz, 1H), 3.72 (dd, *J* = 9.0, 6.4 Hz, 1H), 3.41 (app t, *J* = 5.5, 2H), 2.33 (t, *J* = 7.4 Hz, 2H), 1.83 (s, 3H), 1.57 (tt, *J* = 7.4, 7.0 Hz, 2H), 1.32-1.22 (m, 10H), 0.88 (t, *J* = 6.8 Hz, 3H); $^{13}$C NMR (101 MHz, $CDCl_3$): δ 171.6, 170.0, 154.0, 153.8 (d, *J* = 243 Hz), 135.6 (d, *J* = 11 Hz), 125.0, 121.5 (d, *J* = 12 Hz), 113.3 (d, *J* = 3 Hz), 105.5 (d, *J* = 25 Hz), 71.6, 47.2, 41.4, 35.6, 31.3, 28.7, 28.6, 25.1, 22.4, 22.1, 14.0; FTIR (ATR): υ 3291, 1729, 1675, 1538 cm$^{-1}$; HR-MS (ESI) *m/z*: found 430.2117; calculated for $C_{21}H_{30}FN_3O_4Na$ $[M+Na]^+$ 430.2118.

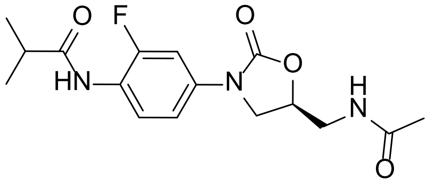

**(*S*)-*N*-(4-(5-(acetamidomethyl)-2-oxooxazolidin-3-yl)-2-fluorophenyl)isobutyramide 18**

The title compound was prepared according to General Procedure A to give **18** as a white powder (31 mg, 51%). $R_f$ 0.13 (5% MeOH/ $CH_2Cl_2$); $^1$H NMR (500 MHz, DMSO-$d_6$): δ 9.56 (s, 1H), 8.24 (t, *J* = 5.8 Hz, 1H), 7.75 (dd, *J* = 8.9, 8.9 Hz, 1H), 7.55 (dd, *J* = 13.2, 2.4 Hz, 1H), 7.21 (dd, *J* = 8.9, 2.4 Hz, 1H), 4.75-4.70 (m, 1H), 4.10 (dd, *J* = 9.0, 9.0 Hz, 1H), 3.73 (dd, *J* = 9.0, 6.5 Hz, 1H), 3.41 (app t, *J* = 5.5 Hz, 2H), 2.70 (sept, *J* = 7.0 Hz, 1H), 1.83 (s, 3H), 1.09 (d, *J* = 7.0 Hz, 6H); $^{13}$C NMR (126 MHz, DMSO-$d_6$): δ 175.5, 170.0, 154.0, 154.0 (d, *J* = 244 Hz), 135.7 (d, *J* = 10 Hz), 125.3, 121.5 (d, *J* = 13 Hz), 113.3 (d, *J* = 3 Hz), 105.4 (d, *J* = 25 Hz), 71.6, 47.2, 41.4, 34.2, 22.4, 19.5; FTIR (ATR): υ 3284, 1729, 1670, 1657, 1537 cm$^{-1}$; HR-MS (ESI) *m/z*: found 360.1342; calculated for $C_{16}H_{20}FN_3O_4Na$ $[M+Na]^+$ 360.1336.

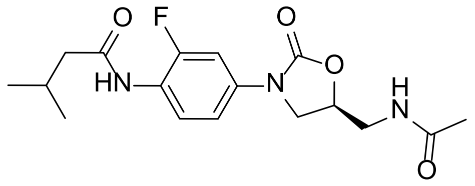

**(*S*)-*N*-(4-(5-(acetamidomethyl)-2-oxooxazolidin-3-yl)-2-fluorophenyl)-3-methylbutanamide 19**

The title compound was prepared according to General Procedure A to give **19** as a white powder (38 mg, 60%). $R_f$ 0.13 (5% MeOH/ $CH_2Cl_2$); $^1$H NMR (500 MHz, DMSO-$d_6$): δ 9.60 (s, 1H), 8.24 (t, *J* = 5.6 Hz, 1H), 7.74 (dd, *J* = 8.8, 8.8 Hz, 1H), 7.56 (dd, *J* = 13.1, 2.2 Hz, 1H), 7.22 (dd, *J* = 8.8, 2.2 Hz, 1H), 4.75-4.70 (m, 1H), 4.10 (dd, *J* = 9.0, 9.0 Hz, 1H), 3.72 (dd, *J* = 9.0, 6.5 Hz, 1H), 3.41 (app t, *J* = 5.5 Hz, 2H), 2.22 (d, *J* = 7.0 Hz, 2H), 2.05 (dsept, *J* = 7.0, 6.5 Hz, 1H), 1.83 (s, 3H), 0.93 (d, *J* = 6.5 Hz, 6H); $^{13}$C NMR (126 MHz, DMSO-$d_6$): δ 170.9, 170.0, 154.0, 154.0 (d, *J* = 244 Hz), 135.7 (d, *J* = 11 Hz), 125.3, 121.4 (d, *J* = 13 Hz), 113.3 (d, *J* = 3 Hz), 105.5 (d, *J* = 26 Hz), 71.6, 47.2, 44.8. 41.4, 25.7, 22.4, 22.3; FTIR (ATR): υ 3285, 1736, 1658, 1534 cm$^{-1}$; HR-MS (ESI) *m/z*: found 374.1502; calculated for $C_{17}H_{22}FN_3O_4Na$ $[M+Na]^+$ 374.1516.

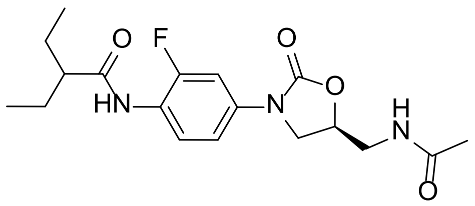

**(*S*)-*N*-(4-(5-(acetamidomethyl)-2-oxooxazolidin-3-yl)-2-fluorophenyl)-2-ethylbutanamide 20**

To a stirred suspension of **10** (48 mg, 0.18 mmol) in pyridine (1.5 mL), diethylacetic anhydride (0.21 mL, 0.90 mmol) was added. The mixture was stirred at r.t. for 18 h, then diluted with toluene (3 mL) and concentrated under reduced pressure. The residue was purified by silica gel chromatography (0-3% MeOH/$CH_2Cl_2$), and the resultant solid was triturated with $CH_2Cl_2$ (2 x 0.2 mL) to yield the title compound as a white powder (40 mg, 61%). $R_f$ 0.13 (5% MeOH/ $CH_2Cl_2$); $^1$H NMR (500 MHz, DMSO-$d_6$): δ 9.60 (s, 1H), 8.24 (t, *J* = 5.8, 1H), 7.68 (dd, *J* = 8.9, 8.9 Hz, 1H), 7.55 (dd, *J* = 13.0, 2.4 Hz, 1H), 7.23 (dd, *J* = 8.9, 2.4 Hz, 1H), 4.75-4.70 (m, 1H), 4.11 (dd, *J* = 9.0, 9.0 Hz, 1H), 3.73 (dd, *J* = 9.0, 7.7 Hz, 1H), 3.41 (app t, *J* = 5.5 Hz, 2H), 2.35-2.30 (m, 1H), 1.58-1.49 (m, 2H), 1.47-1.39 (m, 2H), 0.87

(t, $J$ = 7.4 Hz, 3H); $^{13}$C NMR (126 MHz, DMSO-d$_6$): δ 174.2, 170.0, 154.4 (d, $J$ = 244 Hz), 154.0, 136.0 (d, $J$ = 10 Hz), 125.9, 121.2 (d, $J$ = 13 Hz), 113.3 (d, $J$ = 3 Hz), 105.5 (d, $J$ = 26 Hz), 71.6, 49.0, 47.3, 41.4, 25.3, 22.4, 11.8; FTIR (ATR): υ 3276, 1739, 1658, 1525 cm$^{-1}$; HR-MS (ESI) *m/z*: found 388.1646; calculated for $C_{18}H_{24}FN_3O_4Na$ [M+Na]$^+$ 388.1649.

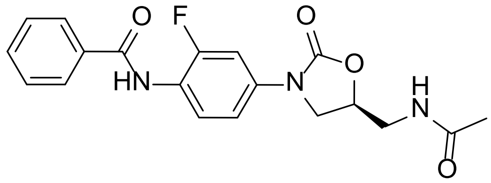

**(*S*)-*N*-(4-(5-(acetamidomethyl)-2-oxooxazolidin-3-yl)-2-fluorophenyl)benzamide 35**

The title compound was prepared according to General Procedure B followed by recrystallisation from MeOH to give **35** as a white powder (23 mg, 34%). R$_f$ 0.07 (5% MeOH/ $CH_2Cl_2$); $^1$H NMR (500 MHz, DMSO-d$_6$): δ 10.09 (s, 1H), 8.26 (t, $J$ = 5.8 Hz, 1H), 7.62-7.52 (m, 5H), 7.32 (dd, $J$ = 9.0, 2.5 Hz, 1H), 4.77-4.72 (m, 1H), 4.15 (dd, $J$ = 9.0, 9.0 Hz, 1H), 3.76 (dd, $J$ = 9.0, 6.5 Hz, 1H), 3.43 (app t, $J$ = 5.5 Hz, 2H), 1.84 (s, 3H); $^{13}$C NMR (126 MHz, DMSO-d$_6$): δ 170.1, 165.5, 155.8 (d, $J$ = 245 Hz), 154.1, 137.1 (d, $J$ = 11 Hz), 133.8, 131.8, 128.5, 127.7, 127.7 (d, $J$ = 3 Hz), 120.9 (d, $J$ = 13 Hz), 113.4 (d, $J$ = 3 Hz), 105.6 (d, $J$ = 25 Hz), 71.7, 47.3, 41.4, 22.5; FTIR (ATR): υ 3288, 1731, 1653, 1515 cm$^{-1}$; HR-MS (ESI) *m/z*: found 394.1188; calculated for $C_{19}H_{18}FN_3O_4Na$ [M+Na]$^+$ 394.1179.

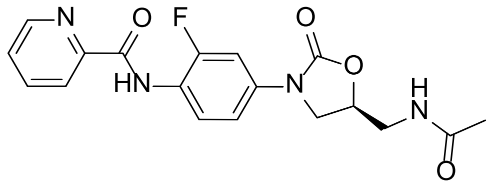

***N*-[4-[(5*S*)-5-[(acetylamino)methyl]-2-oxo-3-oxazolidinyl]-2-fluorophenyl]-2-pyridinecarboxamide 38**

The title compound was prepared according to General Procedure B but instead using pyridine-2-carbonyl chloride hydrochloride (64 mg, 0.36 mmol) and pyridine (0.04 mL, 0.5 mmol) to give **38** as a white powder (31 mg, 46%). R$_f$ 0.17 (5% MeOH/ $CH_2Cl_2$); $^1$H NMR (600 MHz, DMSO-d$_6$): δ 10.37 (s, 1H), 8.75 (d, $J$ = 4.8 Hz), 8.25 (t, $J$ = 5.8 Hz, 1H), 8.16 (d, $J$ = 7.8 Hz, 1H), 8.09 (ddd, $J$ = 7.8, 7.7, 1.6 Hz), 8.04 (dd, $J$ = 8.9, 8.9 Hz, 1H), 7.71 (ddd, $J$ = 7.7, 4.8, 1.1 Hz, 1H), 7.66 (dd, $J$ = 13.2, 1.1 Hz, 1H), 7.32 (dd, $J$ = 8.9, 1.1 Hz, 1H),

4.77-4.73 (m, 1H), 4.14 (dd, $J$ = 8.9, 8.9 Hz, 1H), 3.76 (dd, $J$ = 8.9, 6.5 Hz, 1H), 3.43 (app t, $J$ = 5.5 Hz, 2H), 1.84 (s, 3H); $^{13}$C NMR (151 MHz, DMSO-d$_6$): δ 170.0, 162.1, 154.0, 153.9 (d, $J$ = 244 Hz), 149.0, 148.6, 138.3, 136.1 (d, $J$ = 11 Hz), 127.3, 124.1, 122.2, 120.9 (d, $J$ = 11 Hz), 113.6 (d, $J$ = 2 Hz), 105.5 (d, $J$ = 25 Hz), 71.7, 47.3, 41.4, 22.4; FTIR (ATR): υ 3276, 1744, 1674, 1654, 1526 cm$^{-1}$; HR-MS (ESI) *m/z*: found 395.1138; calculated for $C_{18}H_{17}FN_4O_4Na$ [M+Na]$^+$ 395.1132.

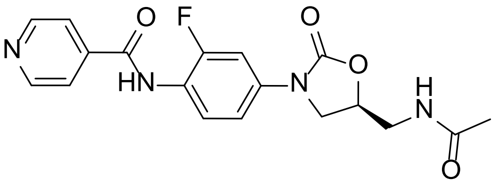

***N*-[4-[(5*S*)-5-[(acetylamino)methyl]-2-oxo-3-oxazolidinyl]-2-fluorophenyl]-4-pyridinecarboxamide 39**

The title compound was prepared according to General Procedure B but instead using pyridine-4-carbonyl chloride hydrochloride (64 mg, 0.36 mmol) and pyridine (0.04 mL, 0.5 mmol) to give **39** as a white powder (21 mg, 31%). $R_f$ 0.27 (10% MeOH/ $CH_2Cl_2$); $^1$H NMR (600 MHz, DMSO-d$_6$): δ 10.41 (s, 1H), 8.79 (d, $J$ = 5.4 Hz, 2H), 8.26 (t, $J$ = 5.7 Hz, 1H), 7.87 (d, $J$ = 5.4 Hz, 2H), 7.63-7.58 (m, 2H), 7.34 (dd, $J$ = 8.8, 1.9 Hz, 1H), 4.77-4.73 (m, 1H), 4.15 (dd, $J$ = 9.0, 9.0 Hz, 1H), 3.77 (dd, $J$ = 9.0, 6.5 Hz, 1H), 3.43 (app t, $J$ = 5.5 Hz, 2H), 1.84 (s, 3H); $^{13}$C NMR (151 MHz, DMSO-d$_6$): δ 170.1, 164.0, 155.7 (d, $J$ = 246 Hz), 154.1, 150.4, 140.9, 137.5 (d, $J$ = 10 Hz), 127.6, 121.6, 120.1 (d, $J$ = 13 Hz), 113.4 (d, $J$ = 2 Hz), 105.7 (d, $J$ = 26 Hz), 71.7, 47.3, 41.4, 22.5; FTIR (ATR): υ 3277, 1729, 1655, 1532 cm$^{-1}$; HR-MS (ESI) *m/z*: found 373.1320; calculated for $C_{18}H_{18}FN_4O_4$ [M+H]$^+$ 373.1312.

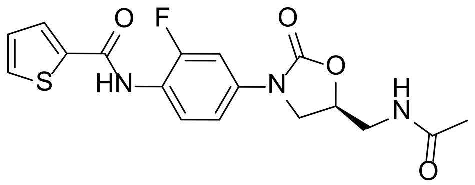

***N*-[4-[(5*S*)-5-[(acetylamino)methyl]-2-oxo-3-oxazolidinyl]-2-fluorophenyl]-2-thiophenecarboxamide 40**

The title compound was prepared according to General Procedure B followed by trituation with $CH_2Cl_2$ to give **40** as a white powder (18 mg, 26%). $R_f$ 0.11 (5% MeOH/ $CH_2Cl_2$); $^1$H

NMR (500 MHz, DMSO-$d_6$): δ 10.13 (bs, 1H), 8.25 (t, *J* = 5.8 Hz, 1H), 7.99 (dd, *J* = 3.7, 0.9 Hz, 1H), 7.86 (dd, *J* = 5.0, 0.9 Hz, 1H), 7.60 (dd, *J* = 12.8, 2.4 Hz, 1H), 7.55 (dd, *J* = 8.8, 8.8 Hz, 1H), 7.32 (dd, *J* = 8.8, 2.4 Hz, 1H), 7.22 (dd, *J* = 5.0, 3.7 Hz, 1H), 4.77-4.72 (m, 1H), 4.14 (dd, *J* = 9.0, 9.0 Hz, 1H), 3.76 (dd, *J* = 9.0, 6.5 Hz, 1H), 3.43 (app t, *J* = 5.5 Hz, 2H), 1.84 (s, 3H); $^{13}$C NMR (126 MHz, DMSO-$d_6$-$d_6$): δ 170.1, 160.1, 155.8 (d, *J* = 246 Hz), 154.1, 139.1, 137.3 (d, *J* = 10 Hz), 132.0, 129.5, 128.2, 127.8 (d, *J* = 3 Hz), 120.2 (d, *J* = 13 Hz), 113.4 (d, *J* = 3 Hz), 105.6 (d, *J* = 26 Hz), 71.7, 47.3, 41.4, 22.5; FTIR (ATR): υ 3335, 1756, 1672, 1525 cm$^{-1}$; HR-MS (ESI) *m/z*: found 400.0736; calculated for $C_{17}H_{16}FN_3O_4SNa$ [M+Na]$^+$ 400.0743.

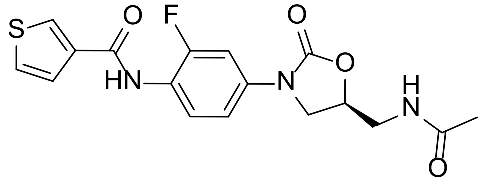


***N*-[4-[(5*S*)-5-[(acetylamino)methyl]-2-oxo-3-oxazolidinyl]-2-fluorophenyl]-3-thiophenecarboxamide 41**

The title compound was prepared according to General Procedure B to give **41** as a white powder (38 mg, 56%). $R_f$ 0.11 (5% MeOH/ $CH_2Cl_2$); $^1$H NMR (600 MHz, DMSO-$d_6$): δ 9.93 (s, 1H), 8.34 (dd, *J* = 2.9, 1.2 Hz, 1H), 8.26 (t, *J* = 5.9 Hz, 1H), 7.65 (dd, *J* = 5.0, 2.9 Hz, 1H), 7.61 (dd, *J* = 5.0, 1.2 Hz, 1H), 7.59 (dd, *J* = 12.8, 2.5 Hz, 1H), 7.55 (dd, *J* = 8.8, 8.8 Hz, 1H), 7.31 (dd, *J* = 8.8, 2.5 Hz, 1H), 4.76-4.72 (m, 1H), 4.14 (dd, *J* = 9.0, 9.0 Hz, 1H), 3.76 (dd, *J* = 9.0, 6.5 Hz, 1H), 3.43 (app t, *J* = 5.5 Hz, 2H), 1.84 (s, 3H); $^{13}$C NMR (151 MHz, DMSO-$d_6$): δ 170.1, 160.9, 155.8 (d, *J* = 245 Hz), 154.1, 137.1 (d, *J* = 10 Hz), 137.0, 130.1, 127.7 (d, *J* = 2 Hz), 127.1, 127.0, 120.6 (d, *J* = 12 Hz), 113.4 (d, *J* = 3 Hz), 105.6 (d, *J* = 26 Hz), 71.7, 47.3, 41.4, 22.5; FTIR (ATR): υ 3289, 1728, 1651, 1530 cm$^{-1}$; HR-MS (ESI) *m/z*: found 400.0746; calculated for $C_{17}H_{16}FN_3O_4SNa$ [M+Na]$^+$ 400.0743.

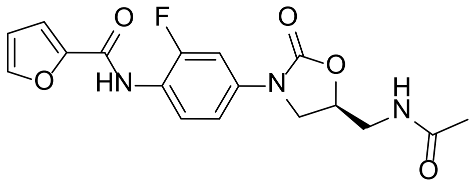

***N*-[4-[(5*S*)-5-[(acetylamino)methyl]-2-oxo-3-oxazolidinyl]-2-fluorophenyl]-2-furancarboxamide 42**

The title compound was prepared according to General Procedure B but instead when the reaction was complete the mixture was filtered, and the solid was washed with $CH_2Cl_2$ (2 x 1 mL) and MeOH (1 mL) to give **42** as a white powder (42 mg, 65%). $R_f$ 0.16 (5% MeOH/ $CH_2Cl_2$); $^1$H NMR (600 MHz, DMSO-$d_6$): δ 9.98 (s, 1H), 8.25 (t, *J* = 5.8 Hz, 1H), 7.94 (d, *J* = 1.7 Hz, 1H), 7.59 (dd, *J* = 12.8, 2.4 Hz, 1H), 7.55 (dd, *J* = 8.8, 8.8 Hz, 1H), 7.32 (d, *J* = 3.5 Hz), 7.30 (dd, *J* = 8.8, 2.2 Hz, 1H), 6.70 (dd, *J* = 3.5, 1.7 Hz, 1H), 4.76-4.72 (m, 1H), 4.14 (dd, *J* = 9.0, 8.9, 1H), 3.75 (dd, *J* = 9.0, 6.4 Hz, 1H), 3.42 (app t, *J* = 5.5 Hz, 2H), 1.84 (s, 3H); $^{13}$C NMR (151 MHz, DMSO-$d_6$): δ 170.0, 156.4, 155.7 (d, *J* = 247 Hz), 154.0, 147.1, 145.9, 137.2 (d, *J* = 11 Hz), 127.5 (d, *J* = 2 Hz), 120.0 (d, *J* = 12 Hz), 114.9, 113.4 (d, *J* = 3 Hz), 112.2, 105.6 (d, *J* = 26 Hz), 71.7, 47.3, 41.4, 22.4; FTIR (ATR): υ 3310, 1764, 1677, 1657, 1350 cm$^{-1}$; HR-MS (ESI) *m/z*: found 384.0978; calculated for $C_{17}H_{16}FN_3O_5Na$ $[M+Na]^+$ 384.0972.

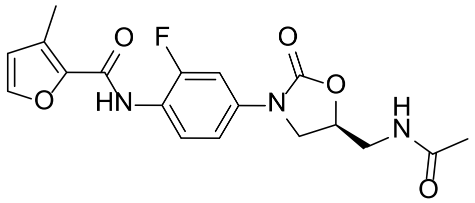

***N*-[4-[(5*S*)-5-[(acetylamino)methyl]-2-oxo-3-oxazolidinyl]-2-fluorophenyl]-3-methyl-2-furancarboxamide 43**

The title compound was prepared according to General Procedure B followed by trituation with $CH_2Cl_2$ to give **43** as a white powder (29 mg, 43%). $R_f$ 0.14 (5% MeOH/ $CH_2Cl_2$); $^1$H NMR (600 MHz, DMSO-$d_6$): δ 9.71 (s, 1H), 8.25 (t, *J* = 5.9 Hz, 1H), 7.79 (d, *J* = 1.5 Hz, 1H), 7.59-7.56 (m, 2H), 7.28 (dd, *J* = 8.7, 2.0 Hz, 1H), 6.58 (d, *J* = 1.5 Hz, 1H), 4.76-4.72 (m, 1H), 4.13 (dd, *J* = 9.0, 9.0, 1H), 3.75 (dd, *J* = 9.0, 6.4 Hz, 1H), 3.43 (app t, *J* = 5.5 Hz, 2H), 2.32 (s, 3H), 1.84 (s, 3H); $^{13}$C NMR (151 MHz, DMSO-$d_6$): δ 170.1, 157.4, 155.6 (d, *J* = 245 Hz), 154.1, 143.9, 141.6, 136.9 (d, *J* = 11 Hz), 127.9, 127.3, 120.3 (d, *J* = 13 Hz),

115.6, 113.4 (d, $J$ = 3 Hz), 105.6 (d, $J$ = 26 Hz), 71.7, 47.3, 41.4, 22.5, 11.0; FTIR (ATR): υ 3317, 1743, 1677, 1664, 1540 $cm^{-1}$; HR-MS (ESI) *m/z*: found 398.1130; calculated for $C_{18}H_{18}FN_3O_5Na$ $[M+Na]^+$ 398.1128.

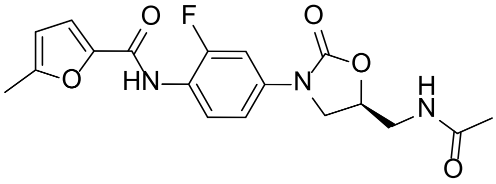

**$N$-[4-[(5$S$)-5-[(acetylamino)methyl]-2-oxo-3-oxazolidinyl]-2-fluorophenyl]-5-methyl-2-furancarboxamide 44**

The title compound was prepared according to General Procedure B to give **44** as an off-white powder (47 mg, 70%). $R_f$ 0.13 (5% MeOH/ $CH_2Cl_2$); $^1$H NMR (500 MHz, DMSO-$d_6$): δ 9.82 (s, 1H), 8.25 (t, $J$ = 5.9 Hz, 1H), 7.60-7.53 (m, 2H), 7.29 (dd, $J$ = 8.7, 1.9 Hz, 1H), 7.21 (d, $J$ = 3.3 Hz, 1H), 6.32 (d, $J$ = 3.3 Hz, 1H), 4.77-4.72 (m, 1H), 4.13 (dd, $J$ = 9.0, 9.0, 1H), 3.75 (dd, $J$ = 9.0, 6.5 Hz, 1H), 3.42 (app t, $J$ = 5.5 Hz, 2H), 2.37 (s, 3H), 1.84 (s, 3H); $^{13}$C NMR (126 MHz, DMSO-$d_6$): δ 170.1, 156.6, 155.6 (d, $J$ = 245 Hz), 155.3, 154.1, 145.6, 137.0 (d, $J$ = 10 Hz), 127.4 (d, $J$ = 3 Hz), 120.2 (d, $J$ = 13 Hz), 116.2, 113.4 (d, $J$ = 3 Hz), 108.6, 105.7 (d, $J$ = 26 Hz), 71.7, 47.3, 41.4, 22.5, 13.6; FTIR (ATR): υ 3303, 1766, 1678, 1655, 1529 $cm^{-1}$; HR-MS (ESI) *m/z*: found 398.1127; calculated for $C_{18}H_{18}FN_3O_5Na$ $[M+Na]^+$ 398.1128.

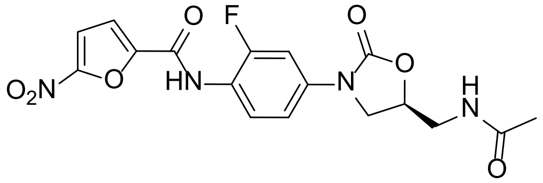

**$N$-[4-[(5$S$)-5-[(acetylamino)methyl]-2-oxo-3-oxazolidinyl]-2-fluorophenyl]-5-nitro-2-furancarboxamide 45**

The title compound was prepared according to General Procedure B but instead when the reaction was complete the mixture was filtered, and the solid was washed with $CH_2Cl_2$ (2 x 1 mL) and MeOH (1 mL) to give **45** as a white powder (32 mg, 44%). $R_f$ 0.16 (5% MeOH/ $CH_2Cl_2$); $^1$H NMR (600 MHz, DMSO-$d_6$): δ 10.55 (s, 1H), 8.25 (t, $J$ = 5.8 Hz, 1H), 7.81 (d, $J$ = 3.9 Hz, 1H), 7.64-7.61 (m, 2H), 7.56 (dd, $J$ = 8.8, 8.8 Hz, 1H), 7.34 (dd, $J$ = 8.8, 2.0 Hz,

1H), 4.77-4.73 (m, 1H), 4.15 (dd, *J* = 9.0, 9.0, 1H), 3.76 (dd, *J* = 9.0, 6.5 Hz, 1H), 3.43 (app t, *J* = 5.5 Hz, 2H), 1.84 (s, 3H); $^{13}$C NMR (151 MHz, DMSO-d$_6$): δ 170.0, 156.5 (d, *J* = 246 Hz), 155.0, 154.0, 151.8, 147.5, 137.8 (d, *J* = 11 Hz), 127.6, 119.1 (d, *J* = 13 Hz), 116.8, 113.5 (d, *J* = 2 Hz), 113.4, 105.7 (d, *J* = 25 Hz), 71.7, 47.2, 41.4, 22.5; FTIR (ATR): ʋ 3324, 1732, 1677, 1531 cm$^{-1}$; HR-MS (ESI) *m/z*: found 429.0828; calculated for $C_{17}H_{15}FN_4O_7Na$ $[M+Na]^+$ 429.0822.

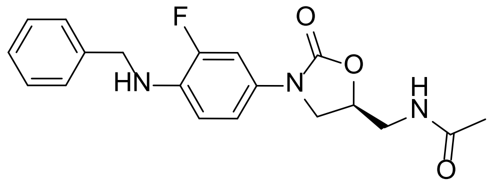


***N*-[[(5*S*)-3-[3-fluoro-4-(benzylamino)phenyl]-2-oxo-5-oxazolidinyl]methyl]acetamide 36**

To a stirred suspension of **10** (75 mg, 0.28 mmol) and benzaldehyde (0.035 mL, 0.34 mmol) in DCE (1.0 mL) was added STAB (0.27g, 1.3 mmol). The mixture was stirred at r.t. for 18 h, and then quenched with 1M NaOH (1 mL). After stirring for 5 min, water (5 mL) was added, then the mixture was extracted with $CH_2Cl_2$ (3 x 5 mL), dried ($MgSO_4$), filtered and concentrated under reduced pressure. The residue was purified by silica gel chromatography (0-3% MeOH/$CH_2Cl_2$) to yield the title compound **36** as a white powder (45 mg, 45%). $R_f$ 0.11 (5% MeOH/ $CH_2Cl_2$); $^1$H NMR (600 MHz, DMSO-d$_6$): δ 8.21 (bt, 1H), 7.39-7.28 (m, 5H), 7.21-7.19 (m, 1H), 6.90 (d, *J* = 8.4 Hz, 1H), 6.57 (dd, *J* = 9.2, 8.4 Hz, 1H), 6.12 (bt, 1H), 4.66-4.61 (m, 1H), 4.33 (d, *J* = 6.0 Hz, 2H), 3.99 (dd, *J* = 9.4, 9.4 Hz, 1H), 3.61 (dd, *J* = 6.5, 8.6 Hz, 1H), 3.36 (m, under $H_2O$ signal, 2H assumed), 1.82 (s, 3H); $^{13}$C NMR (151 MHz, DMSO-d$_6$): δ 170.0, 154.2, 150.2 (d, *J* = 238), 139.9, 133.0 (d, *J* = 12 Hz), 128.3, 127.3 (d, *J* = 9 Hz), 127.0, 126.6, 115.0, 112.2 (d, *J* = 5 Hz), 106.5 (d, *J* = 23 Hz), 71.3, 47.6, 46.0, 41.5, 22.4; FTIR (ATR): ʋ 3295, 1730, 1644, 1526 cm$^{-1}$; HR-MS (ESI) *m/z*: found 358.1569; calculated for $C_{19}H_{21}FN_3O_3$ $[M+H]^+$ 358.1567.

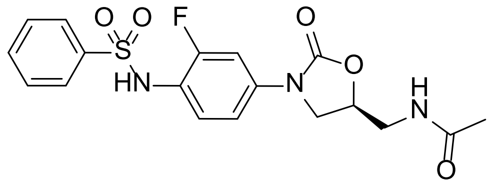

***N*-[[(5*S*)-3-[3-fluoro-4-[(phenylsulfonyl)amino]phenyl]-2-oxo-5-oxazolidinyl]methyl]acetamide 37**

To a stirred suspension of **10** (48 mg, 0.18 mmol) in $CH_2Cl_2$ (1.0 mL) was added pyridine (0.029 mL, 0.22 mmol) and phenylsulfonyl chloride (0.028 mL, 0.22 mmol). The mixture was stirred at r.t. for 18 h, and then quenched with saturated aqueous $NaHCO_3$ (5 mL) and extracted with $CH_2Cl_2$ (3 x 5 mL), dried ($MgSO_4$), filtered and concentrated under reduced pressure. The residue was purified by silica gel chromatography (0-3% MeOH/$CH_2Cl_2$) to yield the title compound **37** as a white powder (36 mg, 49%). $R_f$ 0.06 (5% MeOH/ $CH_2Cl_2$); $^1$H NMR (600 MHz, DMSO-$d_6$): δ 10.06 (bs, 1H), 8.21 (t, *J* = 5.8 Hz, 1H), 7.70-7.68 (m, 2H), 7.65-7.62 (m, 1H), 7.56-7.53 (m, 2H), 7.45-7.42 (m, 1H), 7.22-7.19 (m, 2H), 4.71-4.67 (m, 1H), 4.05 (dd, *J* = 9.0, 9.0 Hz, 1H), 3.67 (dd, *J* = 9.0, 6.5 Hz, 1H), 3.39-3.37 (m, 2H), 1.81 (s, 3H); $^{13}$C NMR (151 MHz, DMSO-$d_6$): δ 170.0, 156.1 (d, *J* = 246 Hz), 153.9, 140.0, 137.6 (d, *J* = 10 Hz), 132.9, 129.2, 127.9, 126.6, 119.1 (d, *J* = 13 Hz), 113.6 (d, *J* = 3 Hz), 105.5 (d, *J* = 26 Hz), 71.7, 47.1, 41.3, 22.4; FTIR (ATR): ʋ 3403, 1750, 1659, 1513 $cm^{-1}$; HR-MS (ESI) *m/z*: found 408.1027; calculated for $C_{18}H_{19}FN_3O_5S$ $[M+H]^+$ 408.1029.

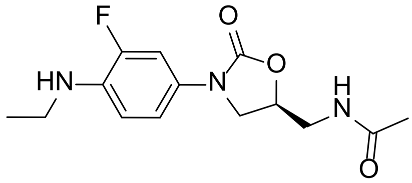

***N*-[[(5*S*)-3-[3-fluoro-4-(ethylamino)phenyl]-2-oxo-5-oxazolidinyl]methyl]acetamide 21**

To a stirred suspension of **10** (50 mg, 0.19 mmol) and $K_2CO_3$ (49 mg, 0.36 mmol) in DMF (1 mL) was added ethyl bromide (0.2 mL, 3 mmol). The mixture was heated at 60 °C in a sealed tube for 18 h, then diluted with EtOAc (5 mL), washed with water (3 x 5 mL), dried ($MgSO_4$), filtered and concentrated under reduced pressure. The residue was purified by silica gel chromatography (0-3% MeOH/$CH_2Cl_2$), followed by C18 silica chromatography (5-50% MeCN/$H_2O$) to yield the title compound **21** as an off-white powder (20 mg, 36%). $R_f$ 0.25 (5% MeOH/ $CH_2Cl_2$); $^1$H NMR (500 MHz, $CDCl_3$): δ 7.31 (dd, *J* = 13.3, 2.5 Hz, 1H),

6.95 (ddd, *J* = 8.7, 2.5, 1.1 Hz, 1H), 6.62 (dd, *J* = 9.1, 8.7 Hz, 1H), 6.61 (bs, 1H), 4.76-4.71 (m, 1H), 3.96 (dd, *J* = 9.1, 9.0 Hz, 1H), 3.73 (bs, 1H), 3.71 (dd, *J* = 9.1, 6.7 Hz, 1H), 3.65 (ddd, *J* = 14.7, 6.0, 3.3 Hz, 1H), 3.58 (ddd, *J* = 14.7, 6.0, 6.0 Hz, 1H), 3.15 (q, *J* = 7.1 Hz, 1H), 2.00 (s, 3H), 1.26 (t, *J* = 7.1 H, 1H); $^{13}$C NMR (126 MHz, $CDCl_3$): δ 171.4, 154.9, 151.0 (d, *J* = 239 Hz), 134.2 (d, *J* = 12 Hz), 127.4 (d, *J* = 10 Hz), 115.3 (d, *J* = 3 Hz), 111.9 (d, *J* = 5 Hz), 107.2 (d, *J* = 24 Hz), 72.0, 48.2, 42.1, 38.4, 23.1, 14.8; FTIR (ATR): ʋ 3330, 1734, 1667, 1525 cm$^{-1}$; HR-MS (ESI) *m/z*: found 296.1415; calculated for $C_{14}H_{19}N_3O_3F$ $[M+H]^+$ 296.1410.

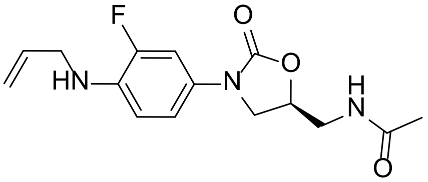


***N*-[[(5*S*)-3-[3-fluoro-4-(allylamino)phenyl]-2-oxo-5-oxazolidinyl]methyl]acetamide 22**

To a stirred suspension of **10** (100 mg, 0.37 mmol) and $K_2CO_3$ (103 mg, 0.75 mmol) in DMF (1.0 mL) was added allyl bromide (45 mg, 0.37 mmol) in DMF (1.0 mL). The mixture was stirred at r.t. for 18 h, then quenched with water (5 mL) and extracted with $CH_2Cl_2$ (2 x 5 mL), washed with water (2 x 5 mL), dried ($MgSO_4$), filtered and concentrated under reduced pressure. The residue was purified by silica gel chromatography (0-3% MeOH/$CH_2Cl_2$) to yield the title compound **22** as a yellow resin (24 mg, 28%). $R_f$ 0.19 (5% MeOH/ $CH_2Cl_2$); $^1$H NMR (600 MHz, $CDCl_3$): δ 7.35 (dd, *J* = 13.2, 2.5 Hz, 1H), 6.95 (ddd, *J* = 8.8, 2.5, 1.2 Hz, 1H), 6.65 (dd, *J* = 9.1, 8.8 Hz, 1H), 6.22 (t, *J* = 5.9 Hz, 1H), 5.96-5.90 (m, 1H), 5.29-5.26 (m, 1H), 5.19-5.17 (m, 1H), 4.76-4.72 (m, 1H), 4.01 (bs, 1H), 3.98 (dd, *J* = 9.1, 8.9 Hz, 1H), 3.80 (d, *J* = 5.3 Hz, 2H), 3.71 (dd, *J* = 9.1, 6.7 Hz, 1H), 3.69 (ddd, 14.7, 6.1, 3.2 Hz, 1H), 3.58 (ddd, 14.7, 6.1, 6.1 Hz, 1H), 2.02 (s, 3H); $^{13}$C NMR (151 MHz, $CDCl_3$): δ 171.2, 154.87, 151.2 (d, *J* = 239 Hz), 134.9, 133.8 (d, *J* = 12 Hz), 127.8 (d, *J* = 10 Hz), 116.7, 115.1 (d, *J* = 3 Hz), 112.3 (d, *J* = 4 Hz), 107.2 (d, *J* = 24 Hz), 71.9, 48.2, 46.3, 42.2, 23.3; FTIR (ATR): ʋ 3288, 1730, 1645, 1526 cm$^{-1}$; HR-MS (ESI) *m/z*: found 308.14156; calculated for $C_{15}H_{19}FN_3O_3$ $[M+H]^+$ 308.1410.

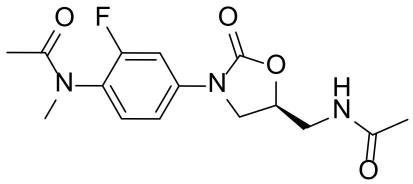

**(*S*)-*N*-(4-(5-(acetamidomethyl)-2-oxooxazolidin-3-yl)-2-fluorophenyl)-*N*-methylacetamide 24**

The title compound was prepared according to General Procedure A to give **24** as a colourless resin (46 mg, 80%). $R_f$ 0.20 (5% MeOH/$CHCl_3$); $^1$H NMR (500 MHz, $CDCl_3$): δ 7.57 (d, *J* = 11.7 Hz, 1H), 7.24-7.19 (m, 2H), 6.86 (t, *J* = 6.1 Hz, 1H), 4.82-4.77 (m, 1H), 4.06 (dd, *J* = 9.1, 8.9 Hz, 1H), 3.82 (dd, *J* = 9.1, 6.7 Hz, 1H), 3.65-3.63 (m, 2H), 3.16 (s, 3H), 2.00 (s, 3H). $^{13}$C NMR (126 MHz, $CDCl_3$): δ 171.6, 171.1, 158.0 (d, *J* = 249 Hz), 154.3, 139.1 (d, *J* = 10 Hz), 129.7 (d, *J* = 2 Hz), 127.6 (d, *J* = 14 Hz), 114.1 (d, *J* = 3 Hz), 107.1 (d, *J* = 26 Hz), 72.3, 47.6, 41.9, 36.5, 23.0, 21.8. FTIR (ATR): υ 3298, 1748, 1646, 1518 cm$^{-1}$; HR-MS (ESI) *m/z*: found 324.1363; calculated for $C_{15}H_{19}FN_3O_4$ $[M+H]^+$ 324.1360.

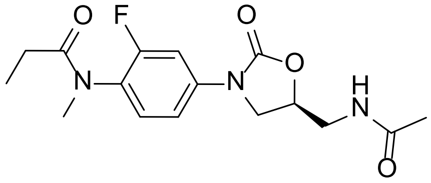

**(*S*)-*N*-(4-(5-(acetamidomethyl)-2-oxooxazolidin-3-yl)-2-fluorophenyl)-*N*-methylpropanamide 25**

The title compound was prepared according to General Procedure A to give **25** as a colourless resin (50 mg, 83%). $R_f$ 0.26 (5% MeOH/$CHCl_3$); $^1$H NMR (500 MHz, $CDCl_3$): δ 7.57 (d, *J* = 11.6 Hz, 1H), 7.23-7.19 (m, 2H), 6.80 (t, *J* = 5.8 Hz, 1H), 4.82-4.77 (m, 1H), 4.06 (dd, *J* = 9.1, 8.9 Hz, 1H), 3.82 (dd, *J* = 9.1, 6.7 Hz, 1H), 3.66-3.63 (m, 2H), 3.16 (s, 3H), 2.05-2.00 (m, 5H) 1.00 (t, *J* = 7.5 Hz, 3H); $^{13}$C NMR (126 MHz, $CDCl_3$): δ 174.4, 171.6, 158.1 (d, *J* = 249 Hz), 154.3, 139.1 (d, *J* = 10 Hz), 130.0 (d, *J* = 2 Hz), 127.2 (d, *J* = 14 Hz), 114.1 (d, *J* = 3 Hz), 107.1 (d, *J* = 26 Hz), 72.2, 47.6, 41.9, 36.6, 27.0, 23.1, 9.5; FTIR (ATR): υ 3298, 1748, 1646, 1518 cm$^{-1}$; HR-MS (ESI) *m/z*: found 338.1514; calculated for $C_{16}H_{21}FN_3O_4$ $[M+H]^+$ 338.1516.

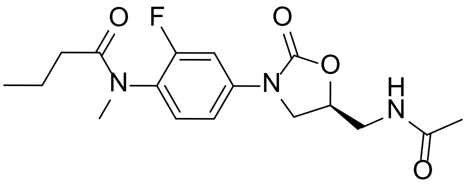

**(*S*)-*N*-(4-(5-(acetamidomethyl)-2-oxooxazolidin-3-yl)-2-fluorophenyl)-*N*-methylbutanamide 26**

The title compound was prepared according to General Procedure A to give **26** as a colourless resin (52 mg, 82%). $R_f$ 0.29 (5% MeOH/$CHCl_3$); $^1$H NMR (600 MHz, $CDCl_3$): δ 7.59 (d, *J* = 11.8 Hz, 1H), 7.24-7.20 (m, 2H), 6.16 (bs), 4.82-4.79 (m, 1H), 4.07 (dd, *J* = 9.1, 8.9 Hz, 1H), 3.82 (dd, *J* = 9.1, 6.8 Hz, 1H), 3.72 (ddd, *J* = 14.7, 6.1, 3.5 Hz, 1H), 3.64 (ddd, *J* = 14.7, 6.1, 6.1 Hz, 1H), 3.20 (s, 3H), 2.03 (s, 3H), 2.00 (t, *J* = 7.5 Hz, 2H), 1.61-1.55 (m, 2H), 0.82 (t, *J* = 7.4 Hz, 3H); $^{13}$C NMR (151 MHz, $CDCl_3$): δ 173.5, 171.3, 158.3 (d, *J* = 250 Hz), 154.2, 139.0 (d, *J* = 10 Hz), 130.2 (d, *J* = 2 Hz), 127.5 (d, *J* = 13 Hz), 114.1 (d, *J* = 3 Hz), 107.2 (d, *J* = 26 Hz), 72.2, 47.6, 42.0, 36.6, 35.7, 23.3, 18.7, 13.9; FTIR (ATR): υ 3300, 1748, 1646, 1518 $cm^{-1}$; HR-MS (ESI) *m/z*: found 352.1674; calculated for $C_{17}H_{23}FN_3O_4$ $[M+H]^+$ 352.1673.

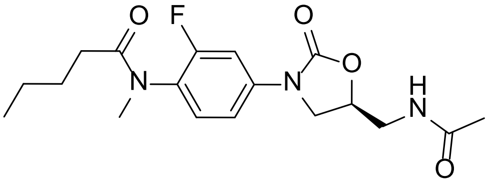

**(*S*)-*N*-(4-(5-(acetamidomethyl)-2-oxooxazolidin-3-yl)-2-fluorophenyl)-*N*-methylpentanamide 27**

The title compound was prepared according to General Procedure A to give **27** as a colourless resin (52 mg, 80%). $R_f$ 0.32 (5% MeOH/$CHCl_3$); $^1$H NMR (600 MHz, $CDCl_3$): δ 7.59 (d, *J* = 11.6 Hz, 1H), 7.24-7.20 (m, 2H), 6.06 (bs), 4.82-4.79 (m, 1H), 4.07 (dd, *J* = 9.0, 8.9 Hz, 1H), 3.82 (dd, *J* = 9.0, 6.8 Hz, 1H), 3.73 (ddd, *J* = 14.7, 5.9, 3.5 Hz, 1H), 3.64 (ddd, *J* = 14.7, 6.1, 6.1 Hz, 1H), 3.20 (s, 3H), 2.04-2.02 (m, 5H), 1.57-1.51 (m, 2H), 1.21 (tt, *J* = 7.5 Hz, 7.4 Hz, 2H), 0.81 (t, *J* = 7.4 Hz, 3H); $^{13}$C NMR (151 MHz, $CDCl_3$): δ 173.7, 171.2, 158.3 (d, *J* = 250 Hz), 154.1, 139.0 (d, *J* = 10 Hz), 130.1 127.6 (d, *J* = 13 Hz), 114.0 (d, *J* = 3 Hz), 107.2 (d, *J* = 26 Hz), 72.1, 47.6, 42.1, 36.6, 33.5, 27.5, 23.3, 22.5, 13.9; FTIR (ATR): υ 3299,

1748, 1646, 1518 cm$^{-1}$; HR-MS (ESI) *m/z*: found 366.1825; calculated for $C_{18}H_{25}FN_3O_4$ $[M+H]^+$ 366.1829.

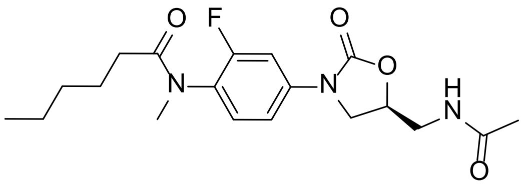

**(*S*)-*N*-(4-(5-(acetamidomethyl)-2-oxooxazolidin-3-yl)-2-fluorophenyl)-*N*-methylhexanamide 28**

The title compound was prepared according to General Procedure A to give **28** as a colourless resin (59 mg, 87%). $R_f$ 0.32 (5% MeOH/$CHCl_3$); $^1$H NMR (600 MHz, $CDCl_3$): δ 7.59 (d, *J* = 11.3 Hz, 1H), 7.24-7.20 (m, 2H), 6.06 (bs), 4.83-4.79 (m, 1H), 4.07 (dd, *J* = 9.0, 8.9 Hz, 1H), 3.82 (dd, *J* = 9.0, 6.8 Hz, 1H), 3.73 (ddd, *J* = 14.7, 5.8, 3.4 Hz, 1H), 3.64 (ddd, *J* = 14.7, 6.1, 6.1 Hz, 1H), 3.20 (s, 3H), 2.04-2.01 (m, 5H), 1.58-1.53 (m, 2H), 1.25-1.13 (m, 4H), 0.83 (t, *J* = 7.2 Hz, 3H); $^{13}$C NMR (151 MHz, $CDCl_3$): δ 173.7, 171.2, 158.3 (d, *J* = 250 Hz), 154.1, 139.0 (d, *J* = 10 Hz), 130.1 127.6 (d, *J* = 13 Hz), 114.0 (d, *J* = 3 Hz), 107.2 (d, *J* = 26 Hz), 72.1, 47.6, 42.1, 36.6, 33.7, 31.5, 25.0, 23.3, 22.5, 14.0; FTIR (ATR): υ 3302, 1749, 1646, 1518 cm$^{-1}$; HR-MS (ESI) *m/z*: found 380.1986; calculated for $C_{19}H_{27}FN_3O_4$ $[M+H]^+$ 380.1986.

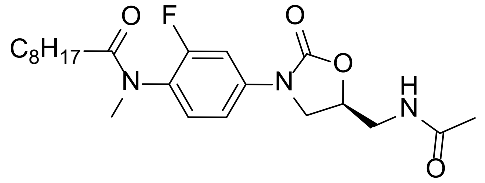

**(*S*)-*N*-(4-(5-(acetamidomethyl)-2-oxooxazolidin-3-yl)-2-fluorophenyl)-*N*-methylnonanamide 29**

The title compound was prepared according to General Procedure B to give **29** as a colourless resin (45 mg, 60%). $R_f$ 0.13 (5% MeOH/$CH_2Cl_2$); $^1$H NMR (500 MHz, $CDCl_3$): δ 7.60-7.57 (m, 1H), 7.22-7.19 (m, 2H), 6.12 (t, *J* = 5.6 Hz, 1H), 4.82-8.79 (m, 1H), 4.07 (dd, *J* = 9.0, 9.0 Hz, 1H), 3.82 (dd, *J* = 9.0, 6.8 Hz, 1H), 3.72 (ddd, *J* = 14.7, 6.1, 3.4 Hz, 1H), 3.63 (ddd, *J* = 14.7, 6.1, 6.1 Hz, 1H), 3.19 (s, 3H), 2.03 (s, 3H), 2.02 (t, *J* = 7.6 Hz, 2H), 1.57-1.51 (m, 2H), 1.27-1.15 (m, 10H), 0.85 (t, *J* = 7.0, 3H); $^{13}$C NMR (126 MHz, $CDCl_3$): δ 173.7, 171.3,

154.0, 158.3 (d, $J$ = 250 Hz), 154.2, 139.0 (d, $J$ = 10 Hz), 130.1, 127.6 (d, $J$ = 14 Hz), 114.0 (d, $J$ = 3 Hz), 107.2 (d, $J$ = 26 Hz), 72.2, 47.6, 42.1, 36.6, 33.8, 31.9, 29.4, 29.4, 29.2, 25.4, 23.3, 22.8, 14.2; FTIR (ATR): ʋ 3302, 1751, 1648, 1518 cm$^{-1}$; HR-MS (ESI) *m/z*: found 444.2265; calculated for $C_{22}H_{32}FN_3O_4Na$ $[M+Na]^+$ 444.2275.

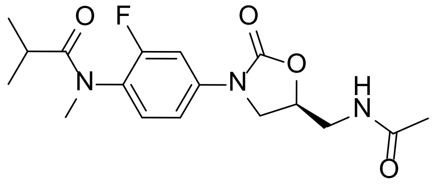

**(*S*)-*N*-(4-(5-(acetamidomethyl)-2-oxooxazolidin-3-yl)-2-fluorophenyl)-*N*-methylisobutyramide 30**

The title compound was prepared according to General Procedure A to give **30** as a colourless resin (50 mg, 80%). $R_f$ 0.30 (5% MeOH/$CHCl_3$); $^1$H NMR (600 MHz, $CDCl_3$): δ 7.60-7.58 (m, 1H), 7.24-7.21 (m, 2H), 6.10 (t, $J$ = 5.8 Hz, 1H), 4.83-4.78 (m, 1H), 4.07 (dd, $J$ = 9.0, 8.9 Hz, 1H), 3.82 (dd, $J$ = 9.0, 6.8 Hz, 1H), 3.72 (ddd, $J$ = 14.7, 5.8, 3.8 Hz, 1H), 3.66-3.62 (m, 1H), 3.19 (s, 3H), 2.40 (sept, $J$ = 6.7 Hz, 1H), 2.03 (s, 3H), 1.02 (d, $J$ = 6.7 Hz, 3H); $^{13}$C NMR (151 MHz, $CDCl_3$): δ 177.9, 171.2, 158.3 (d, $J$ = 249 Hz), 154.2, 139.0 (d, $J$ = 10 Hz), 130.1, 127.5 (d, $J$ = 14 Hz), 114.1, 107.2 (d, $J$ = 26 Hz), 72.2, 47.6, 42.0, 36.8, 31.3, 23.3, 19.9, 19.5; FTIR (ATR): ʋ 3298, 1749, 1646, 1518 cm$^{-1}$; HR-MS (ESI) *m/z*: found 352.1674; calculated for $C_{17}H_{23}FN_3O_4$ $[M+H]^+$ 352.1673.

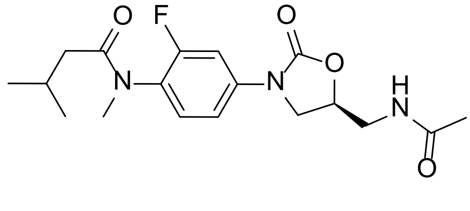

**(*S*)-*N*-(4-(5-(acetamidomethyl)-2-oxooxazolidin-3-yl)-2-fluorophenyl)-*N*,3-dimethylbutanamide 31**

The title compound was prepared according to General Procedure A to give **31** as a colourless resin (39 mg, 60%). $R_f$ 0.15 (5% MeOH/$CH_2Cl_2$); $^1$H NMR (600 MHz, $CDCl_3$): δ 7.59 (d, $J$ = 11.7 Hz, 1H), 7.24-7.18 (m, 2H), 6.04 (t, $J$ = 6.0 Hz, 1H), 4.82-4.79 (m, 1H), 4.08 (dd, $J$ = 8.9, 8.9 Hz, 1H), 3.82 (dd, $J$ = 8.9, 6.8 Hz, 1H), 3.73 (ddd, $J$ = 14.7, 6.0, 3.3 Hz, 1H), 3.63 (ddd, $J$ = 14.7, 6.0, 6.0 Hz, 1H), 3.20 (s, 3H), 2.12 (dsept, 7.0, 6.7 Hz, 1H), 2.03 (s,

3H), 1.92 (d, *J* = 7.0 Hz, 2H), 0.83 (d, *J* = 6.7 Hz, 6H); $^{13}$C NMR (151 MHz, $CDCl_3$): δ 173.0, 171.2, 158.2 (d, *J* = 250 Hz), 154.1, 139.0 (d, *J* = 11 Hz), 130.2, 127.6 (d, *J* = 14 Hz), 114.0 (d, *J* = 3 Hz), 107.1 (d, *J* = 26 Hz), 72.1, 47.6, 42.5, 42.1, 36.6, 25.8, 23.3, 22.7, 22.6; FTIR (ATR): υ 3309, 1749, 1646, 1517 cm$^{-1}$; HR-MS (ESI) *m/z*: found 388.1650; calculated for $C_{18}H_{24}FN_3O_4Na$ $[M+Na]^+$ 388.1649.

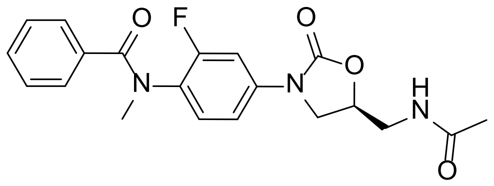

**(*S*)-*N*-(4-(5-(acetamidomethyl)-2-oxooxazolidin-3-yl)-2-fluorophenyl)-*N*-methylbenzamide 46**

The title compound was prepared according to General Procedure B to give **46** as a colourless resin (35 mg, 51%). $R_f$ 0.15 (5% MeOH/$CH_2Cl_2$); $^1$H NMR (600 MHz, $CDCl_3$): δ 7.44 (d, *J* = 11.6 Hz, 1H), 7.31 (bs, 1H), 7.25 (bs, 1H), 7.18 (bs, 2H), 7.02 (bs, 2H), 6.11 (bt, 1H), 4.75-4.73 (m, 1H), 3.96 (dd, *J* = 9.0, 9.0 Hz, 1H), 3.70 (dd, 9.0, 7.2 Hz, 1H), 3.66 (ddd, *J* = 14.7, 6.1, 3.4 Hz, 1H), 3.58 (ddd, *J* = 14.7, 6.1, 6.0 Hz, 1H), 3.39 (s, 3H), 1.99 (s, 3H); $^{13}$C NMR (151 MHz, $CDCl_3$): δ 171.4, 171.2, 157.5 (d, *J* = 250 Hz), 154.1, 138.4 (d, *J* = 9 Hz), 135.7, 130.0, 130.0, 128.5 (d, *J* = 10 Hz), 128.1, 113.5, 106.7 (d, *J* = 26 Hz), 72.1, 47.5, 42.0, 37.6, 23.2; FTIR (ATR): υ 3301, 1748, 1639, 1518 cm$^{-1}$; HR-MS (ESI) *m/z*: found 408.1356; calculated for $C_{20}H_{20}FN_3O_4Na$ $[M+Na]^+$ 408.1336.

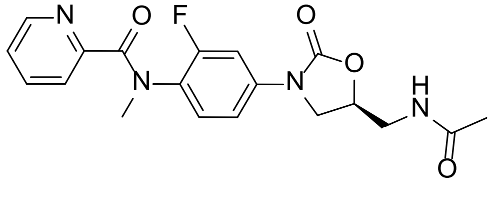

***N*-[4-[(5*S*)-5-[(acetylamino)methyl]-2-oxo-3-oxazolidinyl]-2-fluorophenyl]-*N*-methyl-2-pyridinecarboxamide 49**

The title compound was prepared according to General Procedure B but instead using pyridine-2-carbonyl chloride hydrochloride (64 mg, 0.36 mmol) and pyridine (0.04 mL, 0.5 mmol) to give **49** as a light yellow resin (36 mg, 52%). $R_f$ 0.40 (10% MeOH/ $CH_2Cl_2$); $^1$H NMR (600 MHz, $CDCl_3$): δ 8.25 (m, 1H), 7.64-7.62 (m, 1H), 7.59-7.58 (m, 1H), 7.42-

7.40 (m, 1H), 7.14-7.12 (m, 1H), 7.11-7.08 (m, 1H), 6.99-9.98 (m, 1H), 6.25 (bt, 1H), 4.75-4.71 (m, 1H), 3.97 (dd, *J* = 8.9, 8.9 Hz), 3.73-3.70 (m, 1H), 3.66 (ddd, *J* = 14.6, 6.1, 3.5 Hz, 1H), 3.59-3.55 (m, 1H), 3.42 (s, 3H), 1.99 (s, 3H); $^{13}$C NMR (151 MHz, $CDCl_3$): δ 171.3, 169.0, 157.5 (d, *J* = 249 Hz), 154.1, 153.6, 148.3, 138.2 (d, *J* = 10 Hz), 136.6, 129.7, 128.3 (d, *J* = 12 Hz), 124.5, 123.6, 113.3, 106.4 (d, *J* = 26 Hz), 72.1, 47.5, 42.1, 37.6, 23.2; FTIR (ATR): υ 3300, 1747, 1645, 1518 cm$^{-1}$; HR-MS (ESI) *m/z*: found 409.1288; calculated for $C_{19}H_{19}FN_4O_4Na$ $[M+Na]^+$ 409.1288.

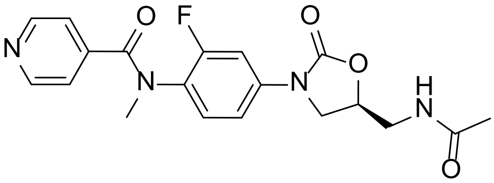


***N*-[4-[(5*S*)-5-[(acetylamino)methyl]-2-oxo-3-oxazolidinyl]-2-fluorophenyl]-*N*-methyl-4-pyridinecarboxamide 50**

The title compound was prepared according to General Procedure B but instead using pyridine-4-carbonyl chloride hydrochloride (64 mg, 0.36 mmol) and pyridine (0.04 mL, 0.5 mmol) to give **50** as a white powder (28 mg, 41%). $R_f$ 0.42 (10% MeOH/ $CH_2Cl_2$); $^1$H NMR (600 MHz, $CDCl_3$): δ 8.47-8.46 (m, 2H), 7.44 (d, *J* = 11.6 Hz, 1H), 7.15-7.14 (m, 2H), 7.07-7.04 (m, 2H), 6.29 (bs, 1H), 4.77-4.73 (m, 1H), 3.96 (dd, *J* = 8.9, 8.9 Hz, 1H), 3.71 (dd, *J* = 8.9, 6.8 Hz, 1H), 3.65 (ddd, 14.6, 6.0, 3.4 Hz, 1H), 3.59 (ddd, 14.6, 5.8, 5.8 Hz, 1H), 3.39 (s, 3H), 1.98 (s, 3H); $^{13}$C NMR (151 MHz, $CDCl_3$): δ 171.3, 169.0, 157.5 (d, *J* = 249 Hz), 154.0, 149.9, 143.4, 139.2 (d, *J* = 10 Hz), 129.8, 127.0 (d, *J* = 13 Hz), 121.9, 113.7, 106.7 (d, *J* = 26 Hz), 72.2, 47.4, 42.0, 37.4, 23.2; FTIR (ATR): υ 3301, 1748, 1650, 1518 cm$^{-1}$; HR-MS (ESI) *m/z*: found 387.1469; calculated for $C_{19}H_{20}FN_4O_4$ $[M+H]^+$ 387.1469.

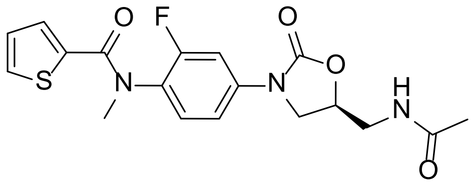

***N*-[4-[(5*S*)-5-[(acetylamino)methyl]-2-oxo-3-oxazolidinyl]-2-fluorophenyl]-*N*-methyl-2-thiophenecarboxamide 51**

The title compound was prepared according to General Procedure B to give **51** as a colourless resin (56 mg, 80%). $R_f$ 0.09 (5% MeOH/$CH_2Cl_2$); $^1$H NMR (500 MHz, $CDCl_3$): δ 7.59 (bd, 1H), 7.31 (dd, *J* = 5.0, 1.0 Hz, 1H), 7.26 (dd, *J* = 8.5, 8.5 Hz, 1H), 7.22-7.18 (bm, 1H), 6.99 (bd, 1H), 6.83 (dd, *J* = 5.0, 4.0 Hz, 1H), 6.20 (t, *J* = 6.0, 1H), 4.83-4.78 (m, 1H), 4.07 (dd, *J* = 9.0, 9.0, 1H), 3.81 (dd, *J* = 9.0, 6.7 Hz, 1H), 3.72 (ddd, *J* = 14.7, 6.0, 3.5 Hz, 1H), 3.63 (ddd, *J* = 14.7, 6.0, 6.0 Hz, 1H), 3.38 (s, 3H), 2.03 (s, 3H); $^{13}$C NMR (126 MHz, $CDCl_3$): δ 171.3, 163.3, 158.5 (d, *J* = 250 Hz), 154.1, 139.5 (d, 10 Hz), 137.1, 132.1, 130.7, 127.6 (d, *J* = 13 Hz), 126.9, 113.9 (d, *J* = 3 Hz), 107.1 (d, *J* = 26 Hz), 72.2, 47.6, 42.1, 38.3, 23.3; FTIR (ATR): υ 3299, 1748, 1627, 1516 $cm^{-1}$; HR-MS (ESI) *m/z*: found 414.0904; calculated for $C_{18}H_{18}FN_3O_4SNa$ $[M+Na]^+$ 414.0900.

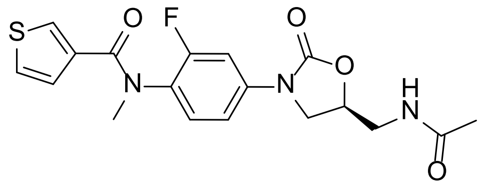

***N*-[4-[(5*S*)-5-[(acetylamino)methyl]-2-oxo-3-oxazolidinyl]-2-fluorophenyl]-*N*-methyl-3-thiophenecarboxamide 52**

The title compound was prepared according to General Procedure B to give **52** as a colourless resin (60 mg, 86%). $R_f$ 0.08 (5% MeOH/$CH_2Cl_2$); $^1$H NMR (500 MHz, $CDCl_3$): δ 7.52-7.50 (m, 1H), 7.26 (bs, 1H), 7.14-7.07 (m, 3H), 6.97 (bs, 1H), 6.29 (t, *J* = 6.0 Hz, 1H), 4.80-4.75 (m, 1H), 4.02 (dd, *J* = 9.0, 9.0, 1H), 3.77 (dd, *J* = 9.0, 6.8 Hz, 1H), 3.68 (ddd, *J* = 14.7, 6.0, 3.5 Hz, 1H), 3.61 (ddd, *J* = 14.7, 6.0, 6.0 Hz, 1H), 3.36 (s, 3H), 2.01 (s, 3H); $^{13}$C NMR (126 MHz, $CDCl_3$): δ 171.3, 165.8, 157.8 (d, *J* = 250 Hz), 154.1, 138.7 (d, 10 Hz), 136.4, 129.9, 129.1, 128.2 (d, *J* = 13 Hz), 128.0, 125.0, 113.7 (d, *J* = 3 Hz), 106.9 (d, *J* = 26 Hz), 72.1, 47.5, 42.0, 37.7, 23.2; FTIR (ATR): υ 3300, 1748, 1634, 1517 $cm^{-1}$; HR-MS (ESI) *m/z*: found 414.0896; calculated for $C_{18}H_{18}FN_3O_4SNa$ $[M+Na]^+$ 414.0900.

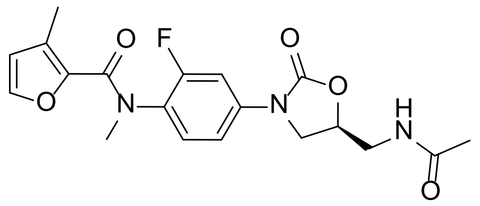

***N*-[4-[(5*S*)-5-[(acetylamino)methyl]-2-oxo-3-oxazolidinyl]-2-fluorophenyl]-*N*-methyl-3-methyl-2-furan carboxamide 53**

The title compound was prepared according to General Procedure B to give **53** as a colourless resin (44 mg, 64%). $R_f$ 0.18 (5% MeOH/$CH_2Cl_2$); $^{1}$H NMR (600 MHz, $CDCl_3$): δ 7.52 (d, $J$ = 11.9 Hz, 1H), 7.13-7.09 (m, 2H), 6.96 (bs, 1H), 6.27 (t, $J$ = 5.8 Hz, 1H), 6.18 (bs, 1H), 4.80-4.76 (m, 1H), 4.05 (dd, $J$ = 9.0, 8.9 Hz, 1H), 3.78 (dd, $J$ = 9.0, 6.7 Hz, 1H), 3.70 (ddd, 14.7, 6.1, 3.4 Hz, 1H), 3.62 (ddd, 14.7, 6.1, 6.1 Hz, 1H), 3.35 (s, 3H), 2.28 (s, 3H), 2.02 (s, 3H); $^{13}$C NMR (151 MHz, $CDCl_3$): δ 171.3, 161.2, 158.0 (d, $J$ = 249 Hz), 154.2, 142.7, 142.6, 138.1 (d, $J$ = 10 Hz), 129.4 (d, $J$ = 2 Hz), 129.2, 128.2 (d, $J$ = 13 Hz), 114.6, 113.4, 106.6 (d, $J$ = 26 Hz), 72.1, 47.6, 42.1, 37.6, 23.2, 11.6; FTIR (ATR): υ 3301, 1748, 1635, 1518 cm$^{-1}$; HR-MS (ESI) *m/z*: found 412.1287; calculated for $C_{19}H_{20}FN_3O_5Na$ $[M+Na]^+$ 412.1285.

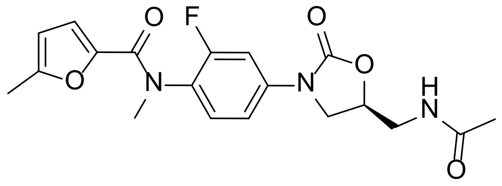

***N*-[4-[(5*S*)-5-[(acetylamino)methyl]-2-oxo-3-oxazolidinyl]-2-fluorophenyl]-*N*,5-methyl-2-furan carboxamide 54**

The title compound was prepared according to General Procedure B to give **54** as a colourless resin (60 mg, 87%). $R_f$ 0.18 (5% MeOH/$CH_2Cl_2$); $^{1}$H NMR (600 MHz, $CDCl_3$): δ 7.57 (bd, 1H), 7.24-7.18 (m, 2H), 6.44 (bs, 1H), 5.94 (bs, 1H), 5.84 (bs, 1H), 4.83-4.78 (m, 1H), 4.07 (dd, $J$ = 9.0, 9.0 Hz, 1H), 3.83 (dd, $J$ = 9.0, 6.7 Hz, 1H), 3.70 (ddd, 14.7, 6.1, 3.7 Hz, 1H), 3.64 (ddd, 14.7, 6.1, 6.0 Hz, 1H), 3.34 (s, 3H), 2.18 (s, 3H), 2.02 (s, 3H); $^{13}$C NMR (151 MHz, $CDCl_3$): δ 171.4, 159.8, 158.3 (d, $J$ = 250 Hz), 155.3, 154.2, 145.2, 139.0 (d, $J$ = 10 Hz), 130.2, 127.7 (d, $J$ = 13 Hz), 117.7, 113.7 (d, $J$ = 3 Hz), 107.8, 106.9 (d, $J$ = 26 Hz), 72.1, 47.7, 42.1, 37.7, 23.2, 13.8; FTIR (ATR): υ 3300, 1749, 1634, 1514 cm$^{-1}$; HR-MS (ESI) *m/z*: found 412.1286; calculated for $C_{19}H_{20}FN_3O_5Na$ $[M+Na]^+$ 412.1285.

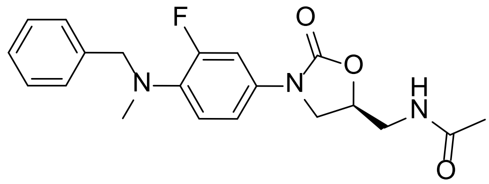

***N*-[[(5*S*)-3-[3-fluoro-4-(benzylmethylamino)phenyl]-2-oxo-5-oxazolidinyl]methyl]acetamide 47**

To a stirred suspension of **23** (50 mg, 0.18 mmol) and $K_2CO_3$ (49 mg, 0.36 mmol) in DMF (1.0 mL) was added benzyl bromide (0.025 mL, 0.21 mmol). The mixture was then stirred at r.t. for 18 h, and then quenched with water (10 mL) and extracted with $CH_2Cl_2$ (4 x 5 mL), washed with water (3 x 5 mL), dried ($MgSO_4$), filtered and concentrated under reduced pressure. The residue was purified by silica gel chromatography (0-3% MeOH/$CH_2Cl_2$) and co-evaporated with 1:1 MeOH/$H_2O$ (2 mL) to yield the title compound **47** as a white powder (39 mg, 60%). $R_f$ 0.19 (5% MeOH/$CH_2Cl_2$); $^1$H NMR (600 MHz, DMSO-$d_6$): δ 8.23 (t, *J* = 5.8 Hz, 1H), 7.48 (dd, *J* = 15.2, 2.6 Hz, 1H), 7.32-7.30 (m, 2H), 7.26-7.23 (m, 3H), 7.11 (ddd, *J* = 8.8, 2.6, 0.6 Hz, 1H), 6.97 (dd, *J* = 9.9, 8.8 Hz, 1H), 4.69 (m, 1H), 4.23 (s, 2H), 4.06 (dd, *J* = 9.1, 8.9, 1H), 3.69 (dd, *J* = 9.1, 6.4 Hz, 1H), 3.39 (dd, *J* = 5.6, 5.6 Hz, 2H), 2.67 (s, 3H), 1.83 (s, 2H); $^{13}$C NMR (151 MHz, DMSO-$d_6$): δ 170.0, 154.1 (d, *J* = 243 Hz), 154.1, 138.0, 135.4 (d, *J* = 8 Hz), 132.3 (d, *J* = 11 Hz), 128.3, 128.1, 127.1, 119.8 (d, *J* = 4 Hz), 114.0 (d, *J* = 2 Hz), 106.8 (d, *J* = 26 Hz), 71.5, 58.4 (d, *J* = 5 Hz), 47.3, 41.4, 39.4, 22.5; FTIR (ATR): υ 3293, 1739, 1660, 1516 cm$^{-1}$; HR-MS (ESI) *m/z*: found 372.1722; calculated for $C_{20}H_{23}FN_3O_3$ $[M+H]^+$ 372.1723.

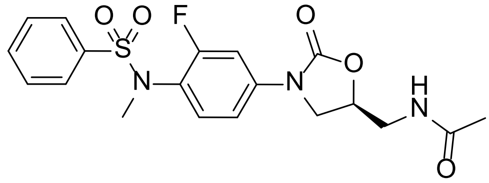

***N*-[[(5*S*)-3-[3-fluoro-4-[methyl(phenylsulfonyl)amino]phenyl]-2-oxo-5-oxazolidinyl]methyl]acetamide 48**

To a stirred suspension of **23** (50 mg, 0.18 mmol) in $CH_2Cl_2$ (1.0 mL) was added pyridine (0.029 mL, 0.22 mmol) and phenylsulfonyl chloride (0.028 mL, 0.22 mmol). The mixture was then stirred at r.t. for 18 h, and then quenched with saturated aqueous $NaHCO_3$ (5 mL) and extracted with $CH_2Cl_2$ (3 x 5 mL), dried ($MgSO_4$), filtered and concentrated under reduced pressure. The residue was purified by silica gel chromatography (0-3% MeOH/$CH_2Cl_2$) to yield the title compound **48** as a colourless resin (62 mg, 83%). $R_f$ 0.09 (5% MeOH/$CH_2Cl_2$); $^1$H NMR (600 MHz, $CDCl_3$): δ 7.68-7.67 (m, 2H), 7.62-7.59 (m, 1H), 7.50-7.47 (m, 3H), 7.29 (dd, *J* = 8.6, 8.6, 1H), 7.11 (dd, *J* = 8.6, 2.0 Hz, 1H), 6.21 (t, *J* = 6.0 Hz, 1H), 4.81-4.76 (m, 1H), 4.04 (dd, *J* = 9.0, 9.0 Hz, 1H), 3.78 (dd, *J* = 9.0, 6.8 Hz, 1H), 3.69 (ddd, *J* = 14.7, 6.1, 3.5 Hz, 1H), 3.62 (ddd, *J* = 14.7, 6.1, 6.1 Hz, 1H), 3.19 (d, *J* = 0.9 Hz, 3H), 2.02 (s, 3H); $^{13}$C NMR (151 MHz, $CDCl_3$): δ 171.3, 159.5 (d, *J* = 253 Hz), 154.1, 139.2 (d, *J* = 10 Hz), 138.0, 133.1, 131.9 (d, *J* = 2 Hz), 129.1, 127.6, 124.1 (d, *J* = 12 Hz), 113.4 (d, *J* = 3 Hz), 106.9 (d, *J* = 26 Hz), 72.1, 47.6, 42.0, 38.2 (d, *J* = 4 Hz), 23.2; FTIR (ATR): υ 3299, 1748, 1656, 1514 cm$^{-1}$; HR-MS (ESI) *m/z*: found 444.1022; calculated for $C_{19}H_{20}FN_3O_5SNa$ $[M+Na]^+$ 444.1005.

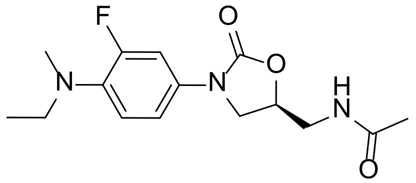


***N*-[[(5*S*)-3-[3-fluoro-4-(ethylmethylamino)phenyl]-2-oxo-5-oxazolidinyl]methyl]acetamide 32**

To a stirred suspension of **23** (50 mg, 0.18 mmol) and $K_2CO_3$ (49 mg, 0.36 mmol) in dried DMF (1 mL) under $N_2$ was added ethyl bromide (0.5 mL, 7 mmol). The mixture was then heated at 60 °C for 18 h, then diluted with EtOAc (5 mL), washed with water (3 x 5 mL), and concentrated under reduced pressure. The residue was purified by silica gel chromatography (0-3% MeOH/$CH_2Cl_2$), followed by C18 silica chromatography (5-50% MeCN/$H_2O$) to yield the title compound **32** as an off-white powder (25 mg, 45%). $R_f$ 0.21 (5% MeOH/$CH_2Cl_2$); $^1$H NMR (500 MHz, $CDCl_3$): δ 7.33 (dd, *J* = 14.6, 2.6 Hz, 1H), 7.01 (ddd, *J* = 8.8, 2.6, 0.9 Hz, 1H), 6.85 (dd, *J* = 9.2, 8.8, 1H), 6.71 (t, *J* = 6.0, 1H), 4.77-4.72 (m, 1H), 3.99 (dd, *J* = 9.1, 9.0 Hz, 1H), 3.73 (dd, *J* = 9.1, 6.7 Hz, 1H), 3.67-3.57 (m, 2H), 3.14 (q, *J* = 7.1 Hz, 2H), 2.76 (s, 3H), 2.00 (s, 3H), 1.07 (t, *J* = 7.1 Hz, 3H); $^{13}$C NMR (126 MHz, $CDCl_3$): δ 171.5, 155.0 (d, *J* = 245 Hz), 154.7, 136.7 (d, *J* = 9 Hz), 131.4 (d, *J* = 10 Hz), 119.4 (d, *J* = 5 Hz), 114.2 (d, *J* = 3 Hz), 107.8 (d, *J* = 27 Hz), 72.1, 49.5 (d, *J* = 5 Hz), 47.8, 42.0, 39.3 (*J* = 2 Hz), 23.1, 12.0; FTIR (ATR): υ 3335, 1733, 1658, 1517 cm$^{-1}$; HR-MS (ESI) *m/z*: found 310.1574; calculated for $C_{15}H_{21}N_3O_3F$ $[M+H]^+$ 310.1567.

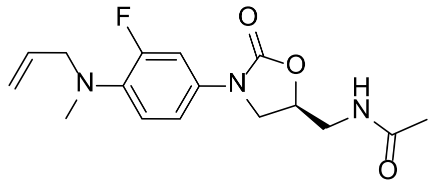

***N*-[[(5*S*)-3-[3-fluoro-4-[allylmethylamino]phenyl]-2-oxo-5-oxazolidinyl]methyl]acetamide 33**

To a stirred suspension of **23** (50 mg, 0.18 mmol) and $K_2CO_3$ (49 mg, 0.36 mmol) in DMF (1.0 mL) was added allyl bromide (0.02 mL, 0.2 mmol). The mixture was stirred at r.t. for 18 h, then diluted with $CH_2Cl_2$ (2 mL), filtered and concentrated under reduced pressure. The residue was purified by silica gel chromatography (0-3% MeOH/$CH_2Cl_2$) to yield the title compound **33** as a peach resin (21 mg, 37%). $R_f$ 0.21 (5% MeOH/$CH_2Cl_2$); $^1$H NMR (600 MHz, $CDCl_3$): δ 7.37 (dd, *J* = 14.5, 2.6 Hz, 1H), 7.03 (ddd, *J* = 8.8, 2.6, 0.9 Hz, 1H), 6.87 (dd, *J* = 9.2, 8.8 Hz, 1H), 6.20 (t, *J* = 6.1 Hz, 1H), 5.88-5.81 (m, 1H), 5.22-5.18 (m, 1H), 5.17-5.14 (m, 1H), 4.77-4.73 (m, 1H), 4.01 (dd, *J* = 9.0, 8.9 Hz, 1H), 3.73 (dd, *J* = 9.0, 6.7 Hz, 1H), 3.70 (d, *J* = 6.1 Hz, 2H), 3.70 (ddd, 14.7, 6.1, 3.2 Hz, 1H), 3.60 (ddd, 14.7, 6.1, 6.1 Hz, 1H), 2.78 (s, 3H), 2.02 (s, 3H); $^{13}$C NMR (151 MHz, $CDCl_3$): δ 171.2, 154.9 (d, *J* = 245 Hz), 154.5, 136.7 (d, *J* = 9 Hz), 134.4, 131.5 (d, *J* = 10 Hz), 119.4 (d, *J* = 4 Hz), 117.7, 114.1 (d, *J* = 3 Hz), 107.8 (d, *J* = 27 Hz), 72.0, 58.1 (d, *J* = 5 Hz), 47.9, 42.1, 39.4, 23.3; FTIR (ATR): υ 3285, 1731, 1648, 1519 cm$^{-1}$; HR-MS (ESI) *m/z*: found 322.1571; calculated for $C_{16}H_{21}FN_3O_3$ $[M+H]^+$ 322.1567.

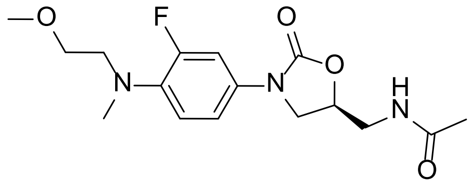

***N*-[[(5*S*)-3-[3-fluoro-4-((2-methoxyethyl)methylamino)phenyl]-2-oxo-5-oxazolidinyl]methyl]acetamide 34**

To a stirred suspension of **23** (50 mg, 0.18 mmol) and $K_2CO_3$ (50 mg, 0.36 mmol) in DMF (1.0 mL) was added 2-bromoethyl methyl ether (0.15 mL, 1.6 mmol). The mixture was stirred at 80 °C for 3 days, then diluted with $CH_2Cl_2$ (2 mL), purified by silica gel chromatography (0-3% MeOH/$CHCl_3$), followed by C18 silica chromatography (5-50% MeCN/$H_2O$) to yield the title compound **34** as a colourless resin (32 mg, 53%). $R_f$ 0.38 (10%

MeOH/$CHCl_3$); $^1$H NMR (600 MHz, $CDCl_3$): δ 7.35 (dd, *J* = 14.7, 2.6 Hz, 1H), 7.03 (ddd, *J* = 8.8, 2.6, 0.8 Hz, 1H), 6.89 (dd, *J* = 9.2, 8.8, 1H), 6.34 (t, *J* = 6.1, 1H), 4.76-4.73 (m, 1H), 4.00 (dd, *J* = 9.1, 9.0 Hz, 1H), 3.73 (dd, *J* = 9.1, 6.7 Hz, 1H), 3.68 (ddd, *J* = 14.7, 6.1, 3.2 Hz, 1H), 3.60 (ddd, *J* = 14.7, 6.1, 6.1 Hz, 1H), 3.54 (t, *J* = 5.8 Hz, 2H), 3.34 (s, 3H), 3.31 (t, *J* = 5.8 Hz, 2H), 2.88 (s, 3H), 2.01 (s, 3H); $^{13}$C NMR (151 MHz, $CDCl_3$): δ 171.3, 154.8 (d, *J* = 245 Hz), 154.6, 136.7 (d, *J* = 9 Hz), 131.3 (d, *J* = 10 Hz), 119.2 (d, *J* = 5 Hz), 114.2 (d, *J* = 3 Hz), 107.8 (d, *J* = 27 Hz), 72.0, 71.1, 59.0, 54.8 (d, *J* = 5 Hz), 47.9, 42.1, 40.7, 23.2; FTIR (ATR): υ 3287, 1731, 1650, 1519 cm$^{-1}$; HR-MS (ESI) *m/z*: found 340.1668; calculated for $C_{16}H_{23}N_3O_4F$ $[M+H]^+$ 340.1673.

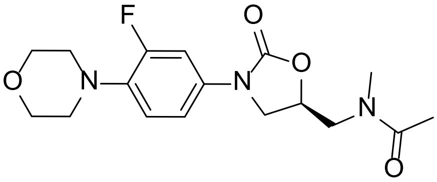

**N-[[(5S)-3-[3-fluoro-4-(4-morpholinyl)phenyl]-2-oxo-5-oxazolidinyl]methyl]-N-methyl-acetamide 9**

To a stirred solution of **1** (30 mg, 0.089 mmol) in DMF (1 mL) under $N_2$ at 0°C was added NaH (60% in mineral oil, 5 mg, 0.1 mmol), and stirred for 10 min before the addition of methyl iodide (15 mg, 0.11 mmol) in DMF (0.2 mL). The solution was stirred at 0°C for 45 min, then quenched with 1M HCl (0.1 mL), diluted with EtOAc (5 mL), and washed with water (3 x 5 mL). The combined aqueous phases were extracted with EtOAc (2 x 5 mL). The combined organic phases were washed with water (2 x 5 mL), dried ($MgSO_4$), filtered and concentrated under reduced pressure. The residue was purified by silica gel chromatography (0-3% MeOH/$CH_2Cl_2$), to yield the title compound **9** as a white powder (15 mg, 48%). $R_f$ 0.32 (5% MeOH/$CH_2Cl_2$); $^1$H NMR (600 MHz, $CDCl_3$): δ 7.44 (dd, *J* = 14.3, 2.6 Hz, 1H), 7.08 (ddd, *J* = 8.8, 2.5, 1.0 Hz, 1H), 6.91 (dd, *J* = 9.1, 8.8 Hz, 1H), 4.86-4.82 (m, 1H), 4.01 (dd, *J* = 9.1, 8.9 Hz, 1H), 3.91 (dd, *J* = 14.5, 3.2 Hz, 1H), 3.86-3.85 (m, 4H), 3.74 (dd, *J* = 9.1, 7.0 Hz, 1H), 3.51 (dd, *J* = 14.5, 6.4 Hz, 1H), 3.17 (s, 3H), 3.05-3.03 (m, 4H), 2.11 (s, 3H); $^{13}$C NMR (151 MHz, $CDCl_3$): δ 172.5, 155.6 (d, *J* = 246 Hz), 154.4, 136.6 (d, *J* = 9 Hz), 133.2 (d, *J* = 10 Hz), 118.9 (d, *J* = 4 Hz), 114.0 (d, *J* = 3 Hz), 107.6 (d, *J* = 26 Hz), 72.7, 67.1, 51.1 (d, *J* = 3 Hz), 51.0, 48.2, 38.7, 21.9; FTIR (ATR): υ 1722, 1665, 1518 cm$^{-1}$; HR-MS (ESI) *m/z*: found 352.1667; calculated for $C_{17}H_{22}N_3O_4F$ $[M+H]^+$ 352.1673.

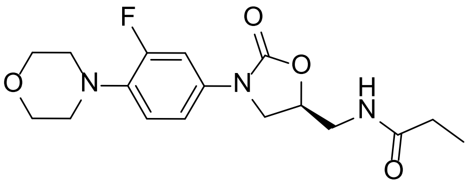

***N*-[(5*S*)-3-[3-fluoro-4-(4-morpholinyl)phenyl)-2-oxo-5-oxazolidiny]methyl}propanamide 5**

The title compound **5** was prepared as by Reddy *et al.*[1] $^{1}$H NMR (500 MHz, $CDCl_3$): δ 7.42 (dd, $J$ = 14.4, 2.6 Hz, 1H), 7.06 (ddd, $J$ = 8.9, 2.6, 1.0 Hz, 1H), 6.90 (dd, $J$ = 9.1, 8.9 Hz, 1H), 6.24 (t, $J$ = 6.0 Hz, 1H), 4.78-4.74 (m, 1H), 4.01 (dd, $J$ = 9.1, 9.0 Hz, 1H), 3.86-3.85 (m, 4H), 3.76 (dd, $J$ = 9.1, 6.5 Hz, 1H), 3.70-3.61 (m, 2H), 3.05-3.03 (m, 4H), 2.23 (q, $J$ = 7.6 Hz, 2H), 1.11 (t, $J$ = 7.6 Hz, 3H).

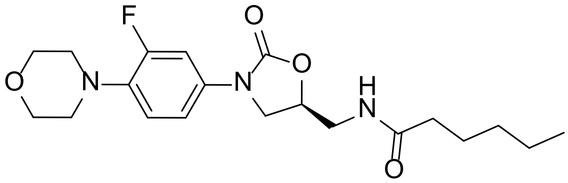

**(*S*)-*N*-(3-[3-fluoro-4-morpholinophenyl]-2-oxo-5-oxazolidinyl]methyl]hexanamide 6**

The title compound **6** was prepared as by Reddy *et al.*[1] $^{1}$H NMR (500 MHz, $CDCl_3$): δ 7.43 (dd, $J$ = 14.3, 2.6 Hz, 1H), 7.06 (ddd, $J$ = 8.9, 2.6, 1.0 Hz, 1H), 6.90 (dd, $J$ = 9.1, 8.9 Hz, 1H), 6.05 (t, $J$ = 6.1 Hz, 1H), 4.78-4.74 (m, 1H), 4.01 (dd, $J$ = 9.1, 9.0 Hz, 1H), 3.87-3.85 (m, 4H), 3.76 (dd, $J$ = 9.1, 6.5 Hz, 1H), 3.71-3.62 (m, 2H), 3.05-3.03 (m, 4H), 2.24-2.14 (m, 2H), 1.57 (tt, $J$ = 7.5, 7.5 Hz, 2H), 1.27-1.21 (m, 4H), 0.82 (t, $J$ = 7.0 Hz, 3H).

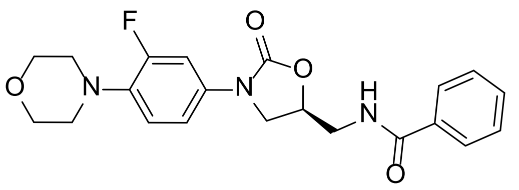

**(*S*)-*N*-[[3-[3-Fluoro-4-morpholinphenyl]-2-oxooxazolidin-5-yl)methyl]benzamide 7**

The title compound **7** was prepared as by Reddy *et al.*[2] $^{1}$H NMR (500 MHz, DMSO-$d_6$): δ 8.82 (t, $J$ = 5.7 Hz, 1H), 7.85-7.83 (m, 2H), 7.55-7.52 (m, 1H), 7.50-7.45 (m, 3H), 7.19 (dd, $J$ = 8.9, 2.1 Hz, 1H), 7.05 (dd, $J$ = 9.3, 8.9 Hz, 1H), 4.88-4.83 (m, 1H), 4.15 (dd, $J$ = 9.1, 9.0

Hz, 1H), 3.85 (dd, $J$ = 9.1, 5.9 Hz, 1H), 3.73-3.72 (m, 4H), 3.67-3.57 (m, 2H), 2.96-2.94 (m, 4H).

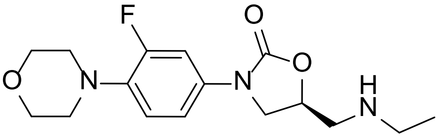


**(5*S*)-5-[(ethylamino)methyl]-3-[3-fluoro-4-(4-morpholinyl)phenyl]-2-oxazolidinone 8**

To a stirred suspension of **4** (75 mg, 0.25 mmol) and $K_2CO_3$ (27 mg, 0.20 mmol) in acetonitrile (1 mL) was added ethyl bromide (60 mg, 0.55 mmol). The mixture was heated at 60 °C in a sealed tube for 18 h, then concentrated under reduced pressure. The residue was taken up in water (5 mL) and extracted with $CH_2Cl_2$ (4 x 5 mL), dried ($MgSO_4$), filtered and concentrated under reduced pressure. The residue was purified by silica gel chromatography (0-10% MeOH/$CHCl_3$) to yield the title compound **8** as an off-white powder (10 mg, 38%). $R_f$ 0.25 (10% MeOH/$CH_2Cl_2$); $^1$H NMR (400 MHz, $CDCl_3$): δ 7.44 (dd, $J$ = 14.4, 2.6 Hz, 1H), 7.13 (ddd, $J$ = 8.8, 2.6, 1.1 Hz, 1H), 6.92 (dd, $J$ = 9.1, 8.8 Hz, 1H), 4.79-4.72 (m, 1H), 4.00 (dd, $J$ = 8.6, 8.5 Hz, 1H), 3.87-3.85 (m, 4H), 3.82 (dd, $J$ = 8.5, 6.8 Hz, 1H), 3.05-3.03 (m, 4H), 2.97 (dd, $J$ = 12.9, 4.4 Hz, 1H), 2.91 (dd, $J$ = 12.9, 6.1 Hz, 1H), 2.75-2.67 (m, 2H), 1.11 (t, $J$ = 7.1 Hz, 3H); FTIR (ATR): υ 1733, 1516 cm$^{-1}$; HR-MS (ESI) *m/z*: found 324.1723; calculated for $C_{16}H_{23}N_3O_3F$ $[M+H]^+$ 324.1723.

***Preparation of 56***

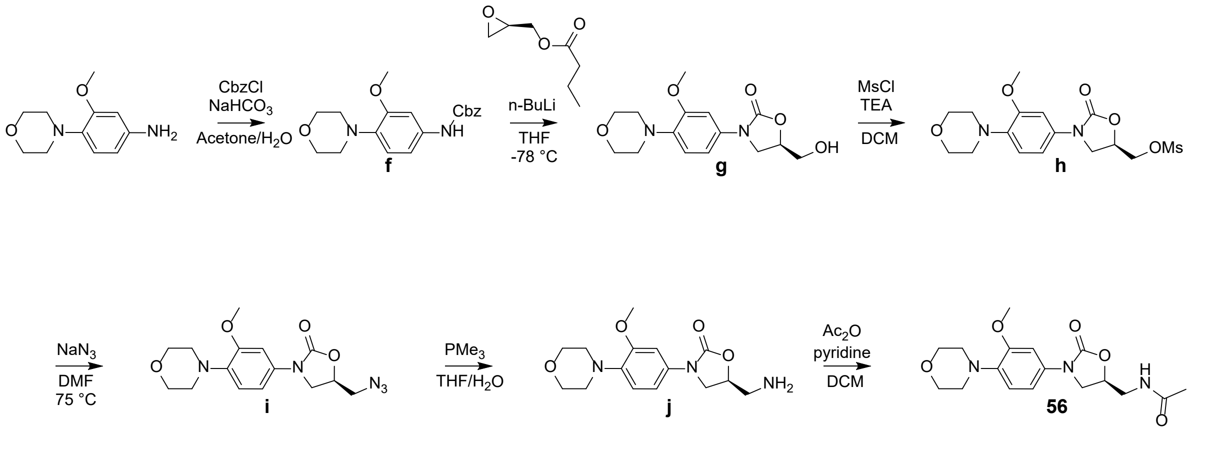

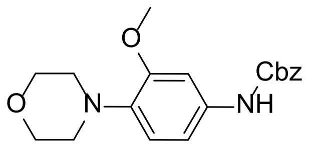


**[3-methoxy-4-(4-morpholinyl)phenyl]carbamic acid benzyl ester f**

To a stirred solution of 3-methoxy-4-(4-morpholinyl)aniline (1.00 g, 4.82 mmol) in acetone (18 mL) and water (9 mL) was added sodium bicarbonate (0.810 g, 9.64 mmol), and then cooled to 0 °C. Benzyl chloroformate (0.73 mL, 5.1 mmol) was added dropwise, and the mixture was stirred at 0 °C for 1 h, then at r.t. for 1 h, poured into water (30 mL) and stirred for 1 h. The resultant solid was filtered, washed with water (3 x 5 mL) and air dried to yield the title compound **f** as a dark brown powder (1.40 g, 85%). $^{1}$H NMR (400 MHz, $CDCl_3$): δ 7.42-7.32 (m, 5H), 7.20 (bs, 1H), 6.84 (d, *J* = 8.5 Hz, 1H), 6.74 (dd, *J* = 8.5, 2.3 Hz, 1H), 6.59 (bs, 1H), 5.19 (s, 2H), 3.89-3.86 (m, 7H), 3.02-3.00 (m, 4H).

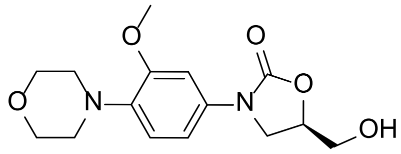


**(5*R*)-5-(hydroxymethyl)-3-[3-methoxy-4-(4-morpholinyl)phenyl]-2-oxazolidinone g**

To a stirred solution of **f** (1.40 g, 4.09 mmol) in THF (25 mL) under $N_2$, cooled to -78 °C, was added dropwise *n*-BuLi (1.6M in hexanes, 3.1 mL, 5.0 mmol). After an additional 30 min, (*R*)-glycidal butyrate (0.69 mL, 4.9 mmol) was added dropwise, and the mixture was allowed to warm to r.t. After 18 h, the mixture was quenched with saturated aqueous $NH_4Cl$ (10 mL), and concentrated under reduced pressure to remove THF, extracted with $CH_2Cl_2$ (3 x 20 mL), dried ($MgSO_4$), filtered and concentrated under reduced pressure. The residue was purified by silica gel chromatography (0-10% MeOH/$CHCl_3$) then triturated with EtOAc (2 mL) to yield the title compound **g** as an beige powder (592 mg, 47%). $R_f$ 0.38 (10% MeOH/$CHCl_3$); $^{1}$H NMR (600 MHz, $CDCl_3$): δ 7.47 (d, *J* = 2.4 Hz, 1H), 6.87 (d, *J* = 8.6 Hz, 1H), 6.75 (dd, *J* = 8.6, 2.4 Hz, 1H), 4.74-4.70 (m, 1H), 4.01 (dd, *J* = 8.8, 8.8 Hz, 1H), 3.98-3.95 (m, 2H), 3.89-3.87 (m, 4H), 3.87 (s, 3H), 3.76-3.73 (m, 1H), 3.03-3.01 (m, 4H), 2.58 (bs, 1H); $^{13}$C NMR (151 MHz, $CDCl_3$): δ 155.0, 152.7, 137.9, 134.0, 118.0, 110.3, 103.3, 72.9, 67.3, 63.0, 55.7, 51.3, 46.8; FTIR (ATR): υ 3435, 1727, 1515 $cm^{-1}$; HR-MS (ESI) *m/z*: found 309.1451 ; calculated for $C_{15}H_{21}N_2O_5$ $[M+H]^+$ 309.1450.

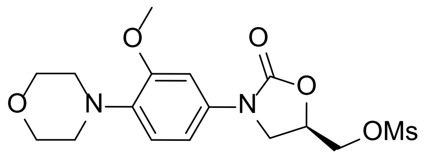


**(5*R*)-5-[[(methylsulfonyl)oxy]methyl]-3-[-3-methoxy-4-(4-morpholinyl)phenyl]-2-oxazolidinone h**

To a stirred suspension of **g** (500 mg, 1.55 mmol) in $CH_2Cl_2$ (25 mL) at 0 °C was added triethylamine (0.43 mL, 3.1 mmol), followed by methanesulfonyl chloride (0.15 mL, 1.9 mmol), and the solution was allowed to warm to r.t. After 1 h, the solution was quenched with water (20 mL), and the organic phase was separated, washed with water (20 mL) and brine (10 mL), dried ($MgSO_4$), filtered and concentrated under reduced pressure to yield presumably the title compound **h** as a brown resin (618 mg) and was used in the next step without further purification.

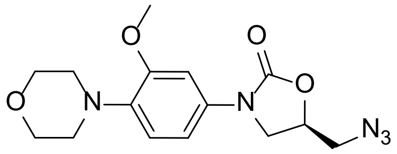


**(5*R*)-5-(azidomethyl)-3-[3-methoxy-4-(4-morpholinyl)phenyl]-2-oxazolidinone i**

To a stirred solution of **h** (618 mg) in anhydrous DMF (7 mL) under $N_2$ was added sodium azide (303 mg, 4.66 mmol) and the mixture was heated at 75 °C. After 3 h, the mixture was cooled to r.t., concentrated under reduced pressure to dryness and taken up in water (20 mL) and EtOAc (10 mL). The aqueous phase was separated and extracted with EtOAc (3 x 5 mL) and the combined organic phases were dried ($MgSO_4$), filtered and concentrated under reduced pressure, then purified by silica gel chromatography (50-100% EtOAc/hexanes) to yield the title compound **i** as a pale yellow crystalline solid (454 mg, 88% over 2 steps). $R_f$ 0.50 (EtOAc); $^1$H NMR (500 MHz, $CDCl_3$): δ 7.50 (d, $J$ = 2.5 Hz, 1H), 6.89 (d, $J$ = 8.6 Hz, 1H), 6.72 (dd, $J$ = 8.6, 2.5 Hz, 1H), 4.79-4.74 (m, 1H), 4.07 (dd, $J$ = 9.0, 8.9 Hz, 1H), 3.89 (s, 3H), 3.89-3.87 (m, 4H), 3.84 (dd, $J$ = 9.0, 6.2 Hz, 1H), 3.69 (dd, $J$ = 13.2, 4.7 Hz, 1H), 3.58 (dd, $J$ = 13.2, 4.5 Hz, 1H), 3.04-3.02 (m, 4H); $^{13}$C NMR (126 MHz, $CDCl_3$): δ 154.2, 152.8, 138.1, 133.7, 118.0, 110.1, 103.3, 70.6, 67.3, 55.8, 53.2, 51.3, 47.9; FTIR (ATR): υ 2107, 1733 $cm^{-1}$; HR-MS (ESI) *m/z*: found 334.1515; calculated for $C_{15}H_{20}N_5O_4$ $[M+H]^+$ 334.1515.

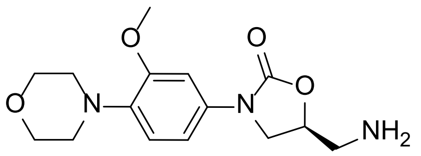

**(5*S*)-5-(aminomethyl)-3-[3-methoxy-4-(4-morpholinyl)phenyl]-2-oxazolidinone j**

To a stirred mixture of **i** (400 mg, 1.2 mmol) in THF (5 mL) and water (0.2 mL) at 0 °C was added trimethylphosphine (1.0 M in toluene, 1.3 mL, 1.3 mmol), then allowed to warm to r.t. After 18 h at r.t., the mixture was concentrated under reduced pressure to remove THF, then diluted with $CH_2Cl_2$ (5 mL) and water (5 mL), and the organic phase was separated. The aqueous phase was extracted with $CH_2Cl_2$ (3 x 5 mL), and the organic phases were combined, dried ($MgSO_4$), filtered and concentrated under reduced pressure, then purified by silica gel chromatography (0-10% MeOH/$CH_2Cl_2$) to yield the title compound **j** as a colourless resin (226 mg, 61%). $R_f$ 0.15 (10% MeOH/$CH_2Cl_2$); $^1$H NMR (600 MHz, $CDCl_3$): δ 7.52 (d, *J* = 2.5 Hz, 1H), 6.88 (d, *J* = 8.6 Hz, 1H), 6.74 (dd, *J* = 8.6, 2.5 Hz, 1H), 4.67-4.63 (m, 1H), 4.03 (dd, *J* = 8.7, 8.6 Hz, 1H), 3.89 (s, 3H), 3.89-3.87 (m, 4H), 3.83 (dd, *J* = 8.6, 6.7 Hz, 1H), 3.09 (dd, *J* = 13.6, 4.1 Hz, 1H), 3.03-3.02 (m, 4H), 2.98 (dd, *J* = 13.6, 5.9 Hz, 1H); $^{13}$C NMR (151 MHz, $CDCl_3$): δ 154.9, 152.8, 137.8, 134.2, 118.0, 110.0, 103.2, 73.9, 67.3, 55.7, 51.4, 48.1, 45.2; FTIR (ATR): υ 3382, 1738, 1511 cm$^{-1}$; HR-MS (ESI) *m/z*: found 308.1612; calculated for $C_{15}H_{22}N_3O_4$ $[M+H]^+$ 308.1610.

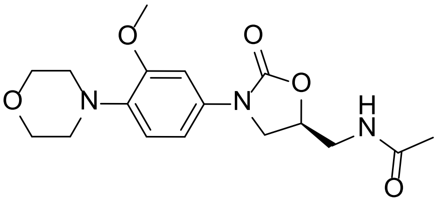

***N*-[[(5*S*)-3-[3-methoxy-4-(4-morpholinyl)phenyl]-2-oxo-5-oxazolidinyl]methyl]acetamide 56**

To a stirred solution of **j** (67 mg, 0.24 mmol) in $CH_2Cl_2$ (4 mL) was added pyridine (0.02 mL, 0.3 mmol) and acetic anhydride (0.03 mL, 0.3 mmol). After 15 min, the solution was concentrated under reduced pressure and purified by silica gel chromatography (5% MeOH/$CHCl_3$) to yield the title compound **56**[3] as a white powder (57 mg, 75%). $R_f$ 0.16 (5% MeOH/$CHCl_3$); $^1$H NMR (600 MHz, $CDCl_3$): δ 7.45 (d, *J* = 2.5 Hz, 1H), 6.89 (d, *J* = 8.6 Hz, 1H), 6.74 (dd, *J* = 8.6, 2.5 Hz, H), 5.97 (bt, 1H), 4.77-4.74 (m, 1H), 4.05 (dd, *J* = 9.1, 9.0 Hz,

1H), 3.89-3.88 (m, 7H), 3.76 (dd, $J$ = 9.1, 6.8 Hz, 1H), 3.72 (ddd, $J$ = 14.7, 6.2, 3.2 Hz, 1H), 3.59 (ddd, $J$ = 14.7, 6.2, 6.2 Hz, 1H), 3.04-3.03 (m, 4H), 2.02 (s, 3H); $^{13}$C NMR (151 MHz, $CDCl_3$): δ 171.2, 154.7, 152.7, 138.1, 133.7, 118.0, 110.4, 103.3, 72.0, 67.2, 55.7, 51.3, 48.0, 42.1, 23.2; FTIR (ATR): υ 3286, 1721, 1647, 1519 cm$^{-1}$; HR-MS (ESI) *m/z*: found 350.1721; calculated for $C_{17}H_{24}N_3O_5$ $[M+H]^+$ 350.1716.

***Preparation of 57***

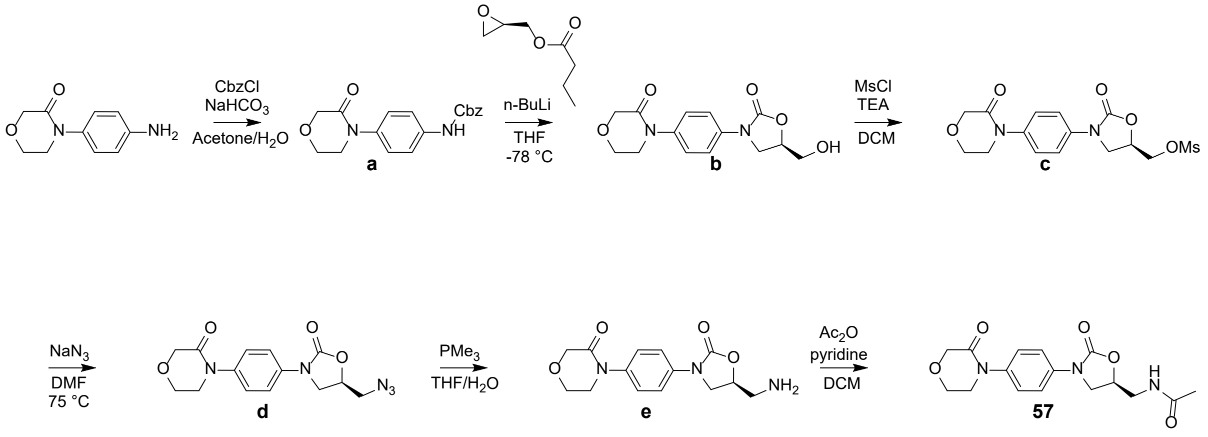


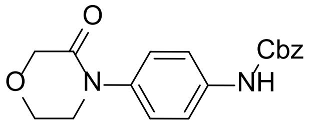

**[4-(3-Oxo-morpholin-4-yl)-phenyl]carbamic acid benzyl ester a**

The title compound **a** was prepared as by Sturm *et al.*[4] $^1$H NMR (400 MHz, $CDCl_3$): δ 7.43-7.32 (m, 7H), 7.26-7.23 (m, 2H), 5.20 (s, 2H), 4.32 (s, 2H), 4.02-4.00 (m, 2H), 3.73-3.71 (m, 2H).

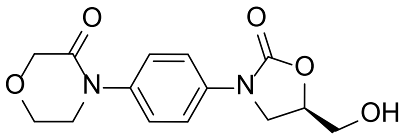


**4-[4-[(5*R*)-5-(Hydroxymethyl)-2-oxo-3-oxazolidinyl]phenyl]-3-morpholinone b**

The title compound **b** was prepared as by Maroju *et al.*[5] $^1$H NMR (400 MHz, $CDCl_3$): δ 7.58-7.56 (m, 2H), 7.33-7.31 (m, 2H), 4.72-4.65 (m, 1H), 4.33 (s, 2H), 4.04-3.94 (m, 4H), 3.91-3.88 (m, 1H), 3.75-3.66 (m, 3H), 2.86 (bt, 1H).

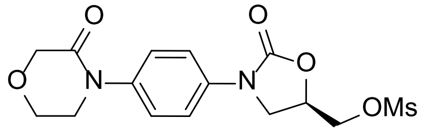


**4-[4-[(5*R*)-5-[[(methylsulfonyl)oxy]methyl]-2-oxo-3-oxazolidinyl]phenyl]-3-morpholinone c**

The title compound **c** was prepared as by Maroju *et al.*[5] $^1$H NMR (400 MHz, $CDCl_3$): δ 7.61-7.57 (m, 2H), 7.39-7.35 (m, 2H), 4.96-4.91 (m, 1H), 4.51 (dd, *J* = 11.7, 3.7 Hz, 1H), 4.43 (dd, *J* = 11.7, 4.1 Hz, 1H), 4.34 (s, 2H), 4.17 (dd, *J* = 9.2, 9.1 Hz, 1H), 4.05-4.03 (m, 2H), 3.98 (dd, *J* = 9.2, 6.1 Hz, 1H), 3.77-3.75 (m, 2H), 3.11 (s, 3H).

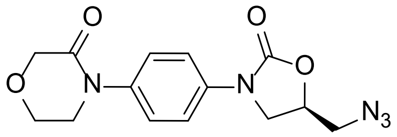


**4-[4-[(5*R*)-5-(Azidomethyl)-2-oxo-3-oxazolidinyl]phenyl]-3-morpholinone d**

The title compound **d** was prepared as by Maroju *et al.*[5] After 3 h of reaction though, the mixture was cooled to r.t., concentrated under reduced pressure to dryness, suspended in water (10 mL) then filtered, washed with water (3 x 5 mL) and air dried to yield the title compound as an off-white powder (787 mg, 71%). $^1$H NMR (400 MHz, $CDCl_3$): δ 7.60-7.58 (m, 2H), 7.37-7.35 (m, 2H), 4.82-4.76 (m, 1H), 4.34 (s, 2H), 4.09 (dd, *J* = 8.9, 8.9 Hz, 1H), 4.05-4.03 (m, 2H), 3.87 (dd, *J* = 8.9, 6.4 Hz, 1H), 3.77-3.74 (m, 2H), 3.71 (dd, *J* = 13.2, 4.5 Hz, 1H), 3.59 (dd, *J* = 13.2, 4.2 Hz, 1H).

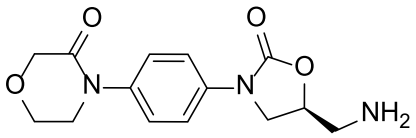

**4-[4-[(5S)-5-(Aminomethyl)-2-oxo-1,3-oxazolidin-3-yl]phenyl]morpholin-3-one e**

To a stirred mixture of **d** (765 mg, 2.41 mmol) in THF (10 mL) and water (0.5 mL) at 0 °C was added trimethylphosphine (1.0 M in toluene, 2.8 mL, 2.8 mmol), then allowed to warm to r.t. After 5 h at r.t., the mixture was concentrated under reduced pressure to remove THF, then diluted with $CH_2Cl_2$ (20 mL) and water (10 mL), and the organic phase was separated. The aqueous phase was extracted with $CH_2Cl_2$ (2 x 20 mL), and the organic phases were combined, dried ($MgSO_4$), filtered and concentrated under reduced pressure to yield the title compound **e** as an off-white solid (570 mg, 81%). The NMR data is consistent as that found in the literature.[6] $^{1}$H NMR (400 MHz, $CDCl_3$): δ 7.63-7.59 (m, 2H), 7.37-7.33 (m, 2H), 4.72-4.65 (m, 1H), 4.34 (s, 2H), 4.08-4.02 (m, 3H), 3.87 (dd, *J* = 8.6, 6.7 Hz, 1H), 3.77-3.74 (m, 2H), 3.12 (dd, *J* = 13.7, 4.1 Hz, 1H), 2.98 (dd, *J* = 13.7, 5.7 Hz, 1H).

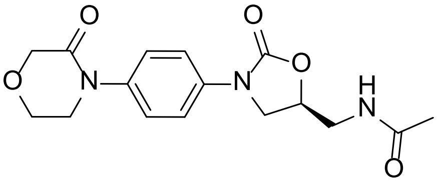

***N*-[[(5*S*)-3-[3-fluoro-4-(3-oxo-4-morpholinyl)phenyl]-2-oxo-5-oxazolidinyl]methyl]acetamide 57**

To a stirred solution of **e** (75 mg, 0.26 mmol) in $CH_2Cl_2$ (5 mL) was added pyridine (0.03 mL, 0.4 mmol) and acetic anhydride (0.03 mL, 0.3 mmol). After 15 min, the solution was concentrated under reduced pressure and purified by silica gel chromatography (0-3% MeOH/$CHCl_3$) to yield the title compound **57** as a white powder (70 mg, 81%). $R_f$ 0.31 (10% MeOH/$CHCl_3$); $^{1}$H NMR (500 MHz, $CDCl_3$): δ 7.56-7.54 (m, 2H), 7.34-7.32 (m, 2H), 6.43 (bt, 1H), 4.76-4.71 (m, 1H), 4.33 (s, 2H), 4.04-4.00 (m, 3H), 3.78 (dd, *J* = 8.9, 7.0 Hz, 1H), 3.75-3.73 (m, 2H), 3.66-3.62 (m, 1H), 3.60-3.56 (m, 1H), 1.99 (s, 3H). $^{13}$C NMR (126 MHz, $CDCl_3$): δ 171.3, 167.0, 154.6, 137.4, 136.8, 126.4, 119.1, 72.0, 68.7, 64.2, 49.8, 47.6, 42.0, 23.2. FTIR (ATR): υ 3290, 1745, 1724, 1647, 1519 $cm^{-1}$. HR-MS (ESI) *m/z*: found 356.1223; calculated for $C_{16}H_{19}N_3O_5Na$ $[M+Na]^+$ 356.1222.
