## Supporting Dataset - NMR spectra for "Inhibition of chloroplast translation as a new target for herbicides"

$^{1}$H NMR spectrum of **8**

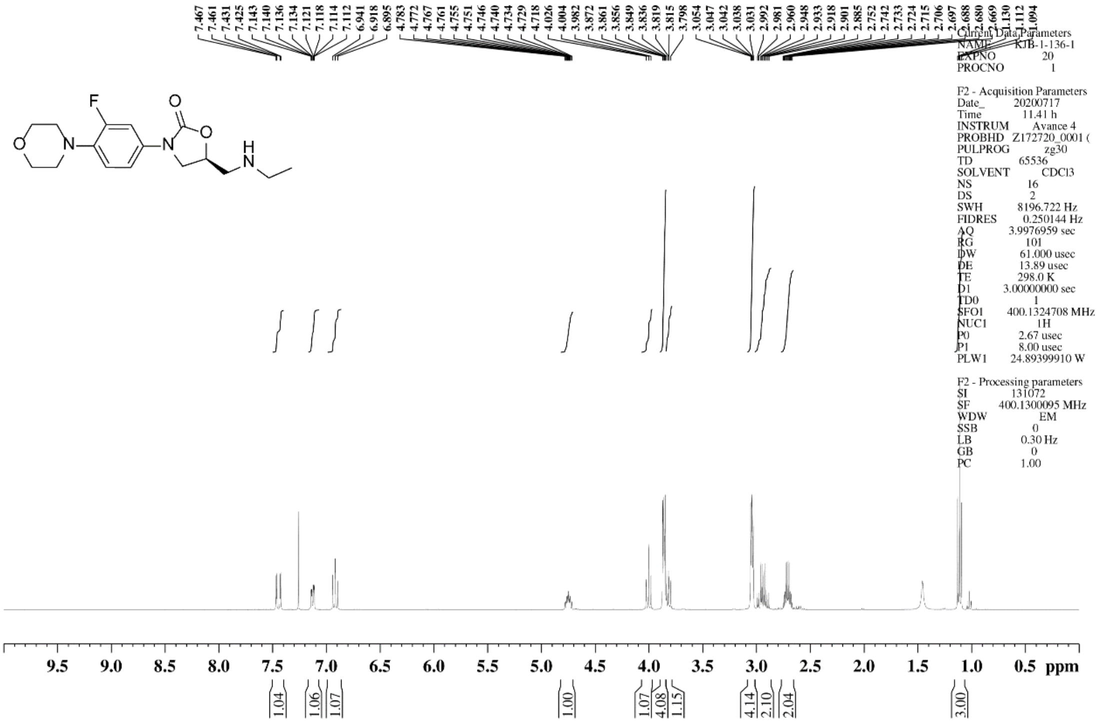

$^{1}H$ NMR spectrum of **9**

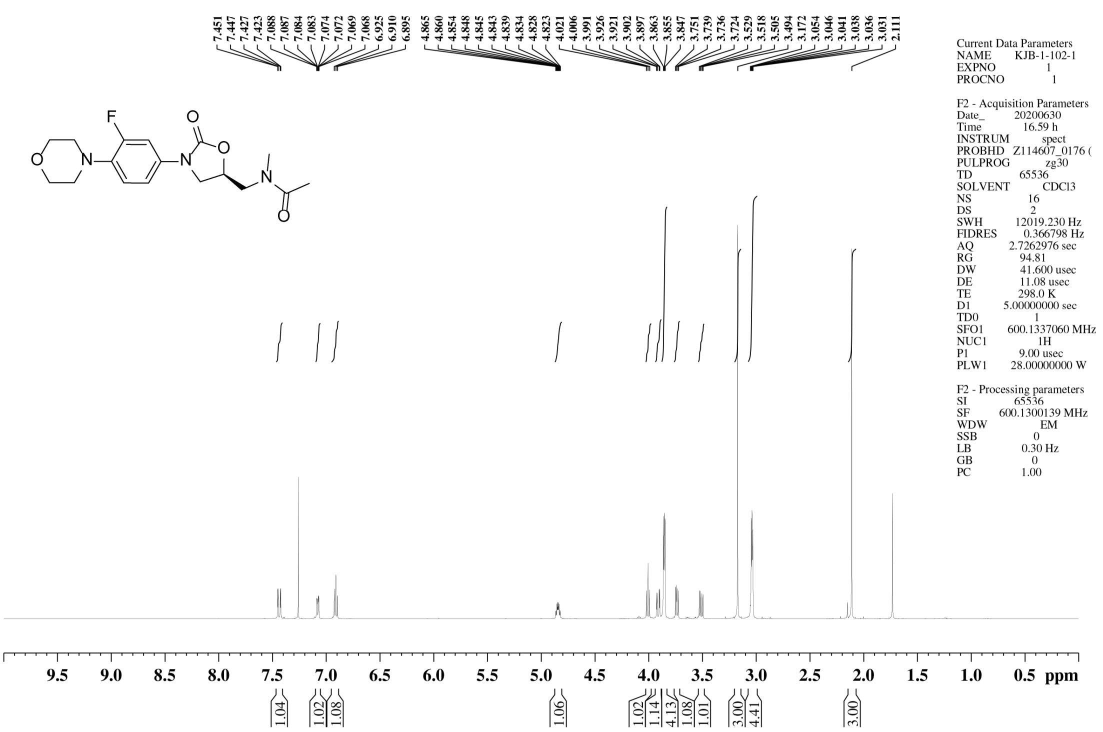


$^{13}C$ NMR spectrum of **9**

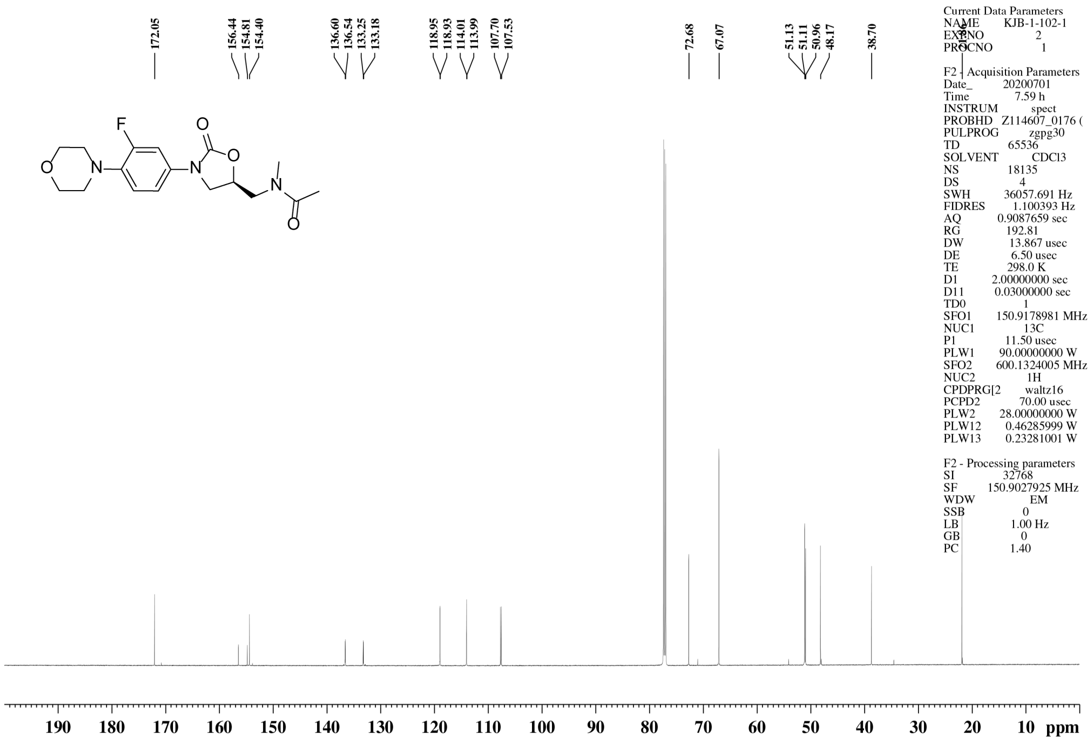

$^{1}$H NMR spectrum of **11**

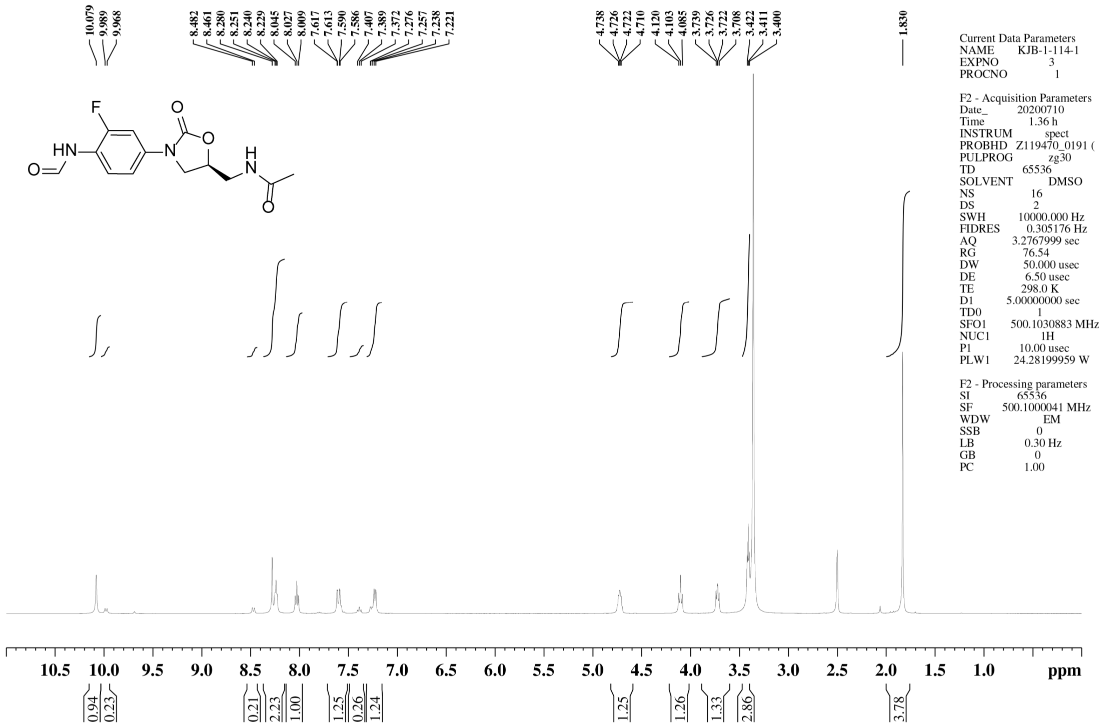


$^{13}$C NMR spectrum of **11**

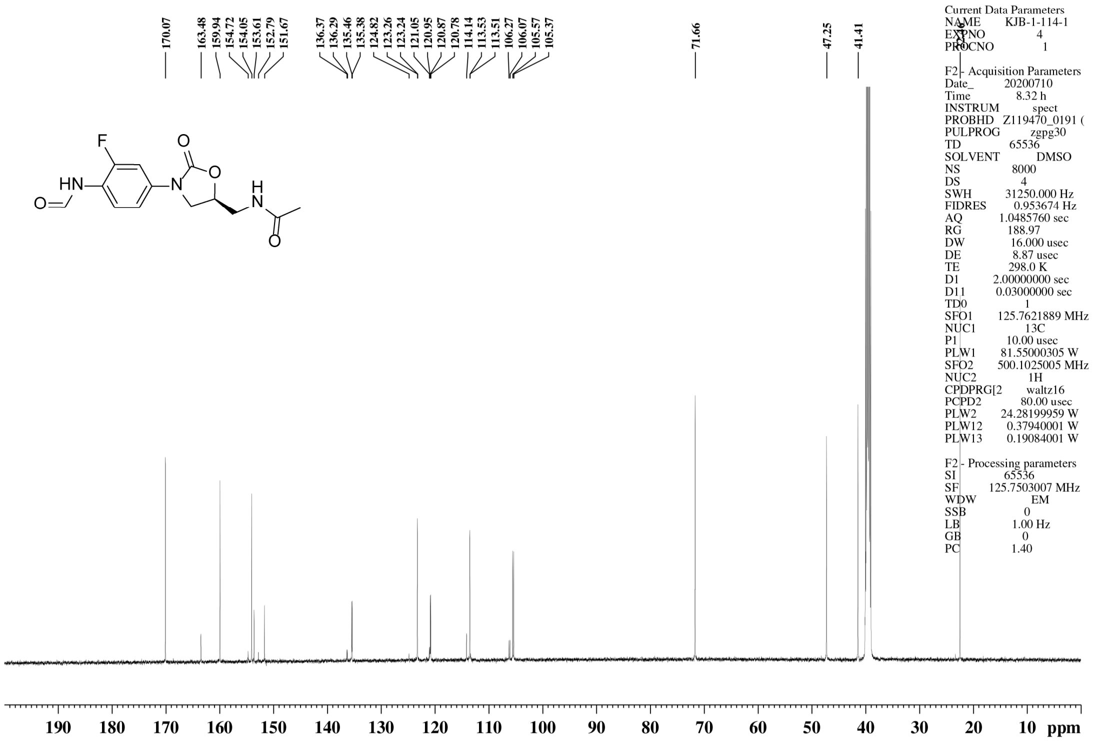

$^{1}$H NMR spectrum of **12**

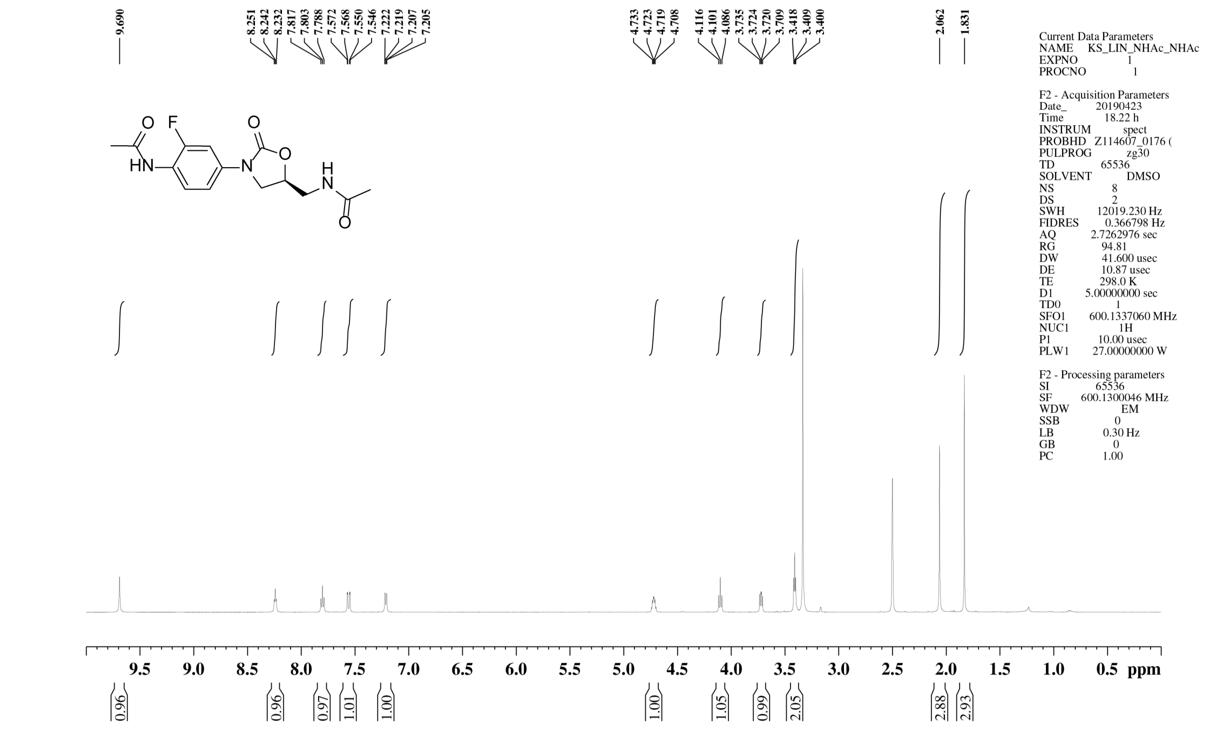


$^{13}$C NMR spectrum of **12**

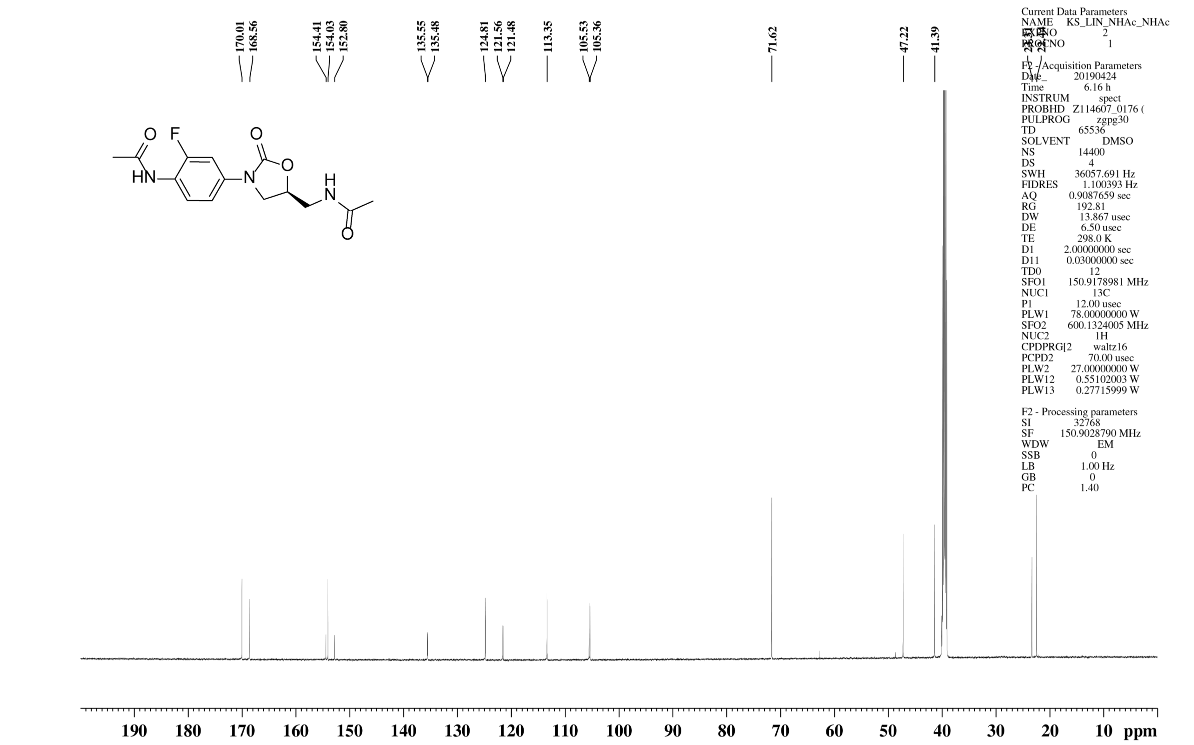

#### $^{1}$H NMR spectrum of **13**

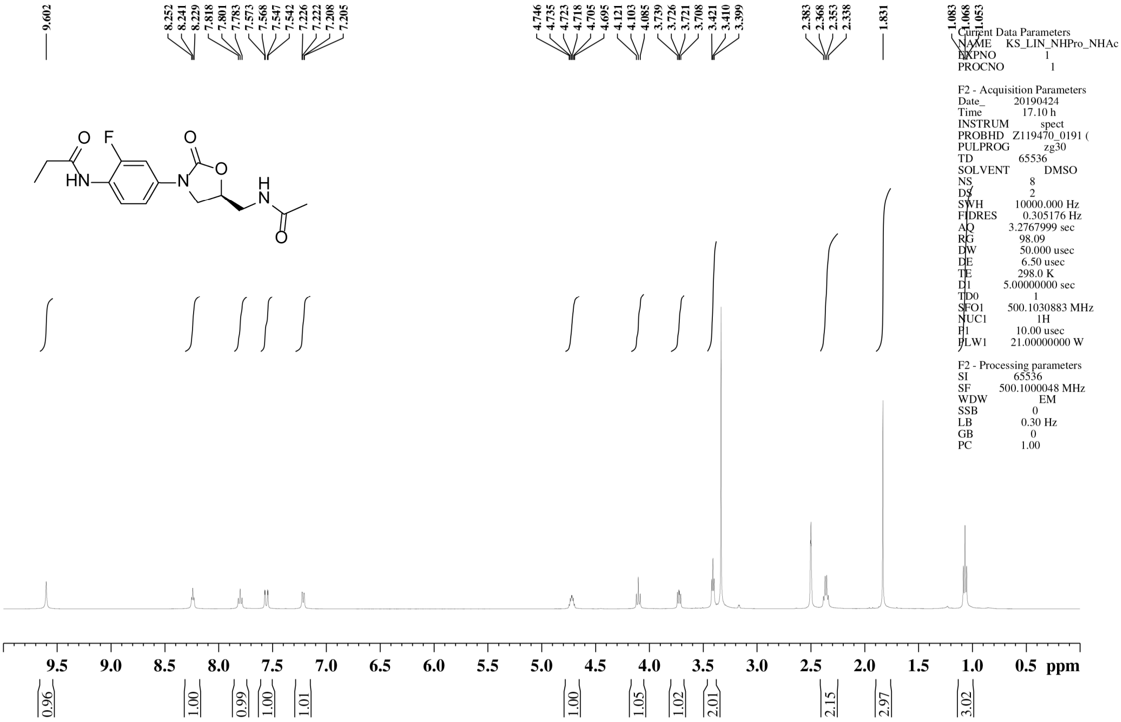


#### $^{13}$C NMR spectrum of **13**

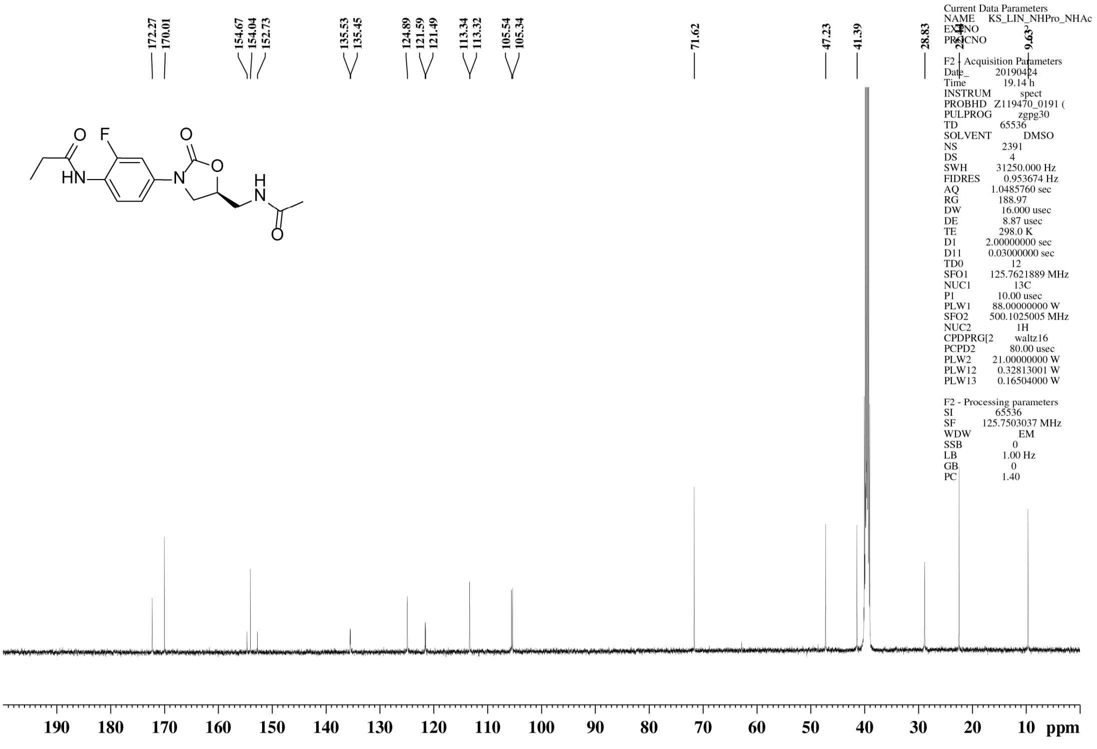

### $^{1}$H NMR spectrum of **14**

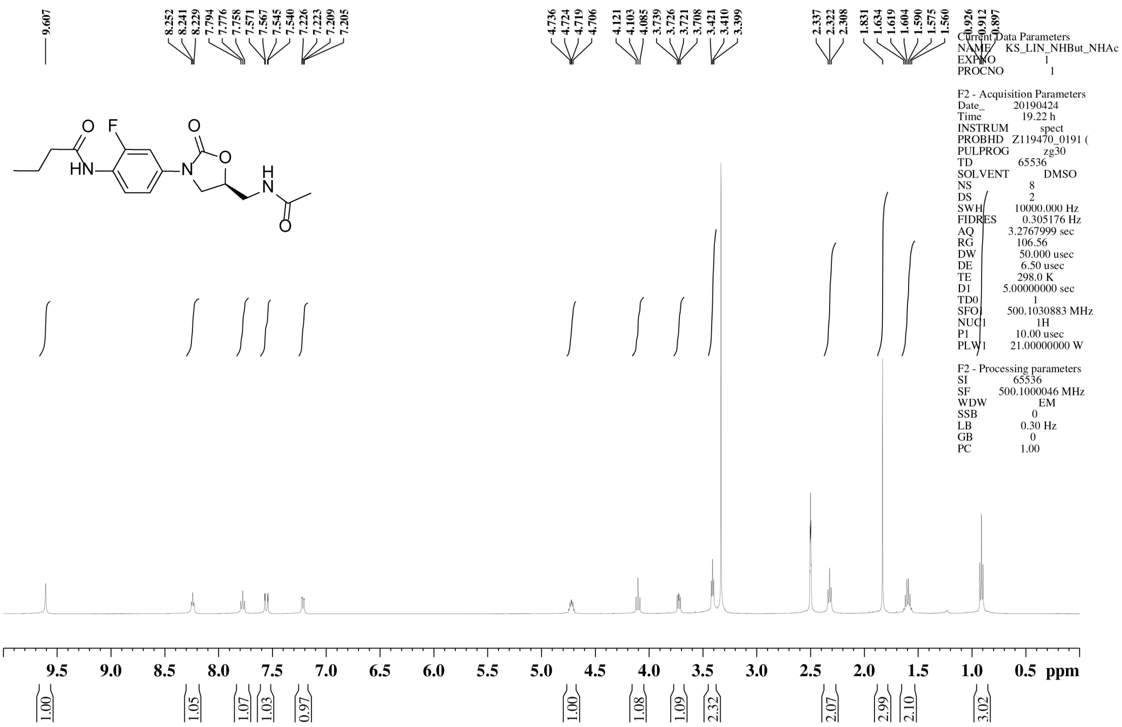


### $^{13}$C NMR spectrum of **14**

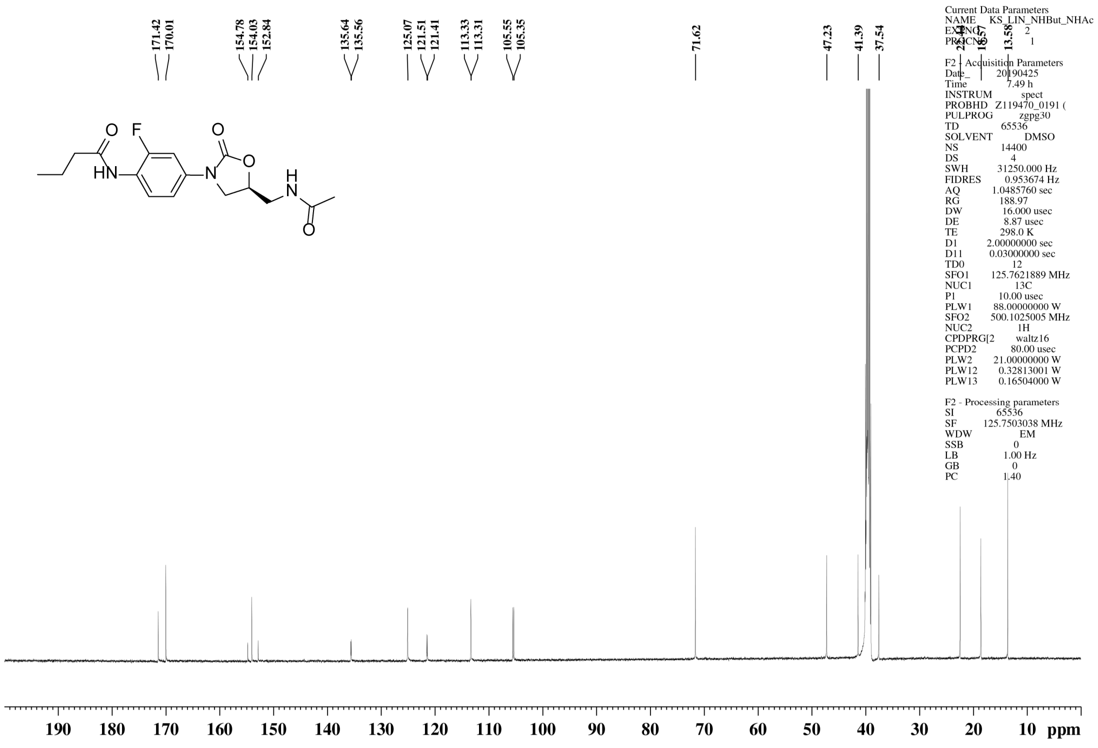

### $^{1}$H NMR spectrum of **15**

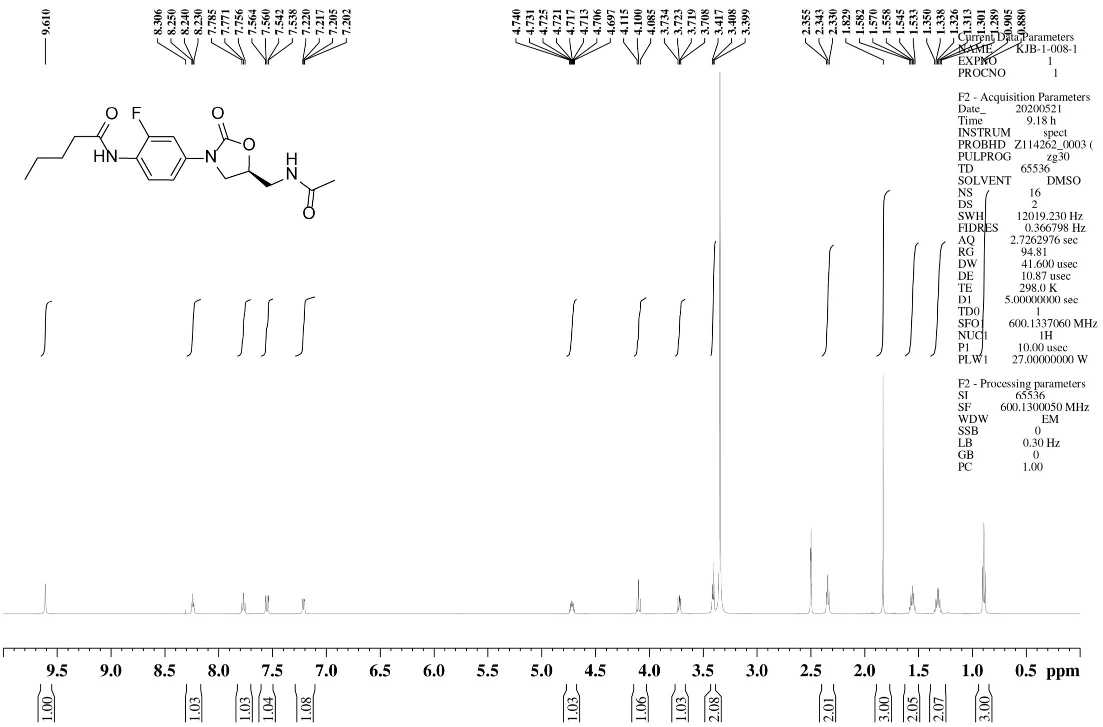


### $^{13}$C NMR spectrum of **15**

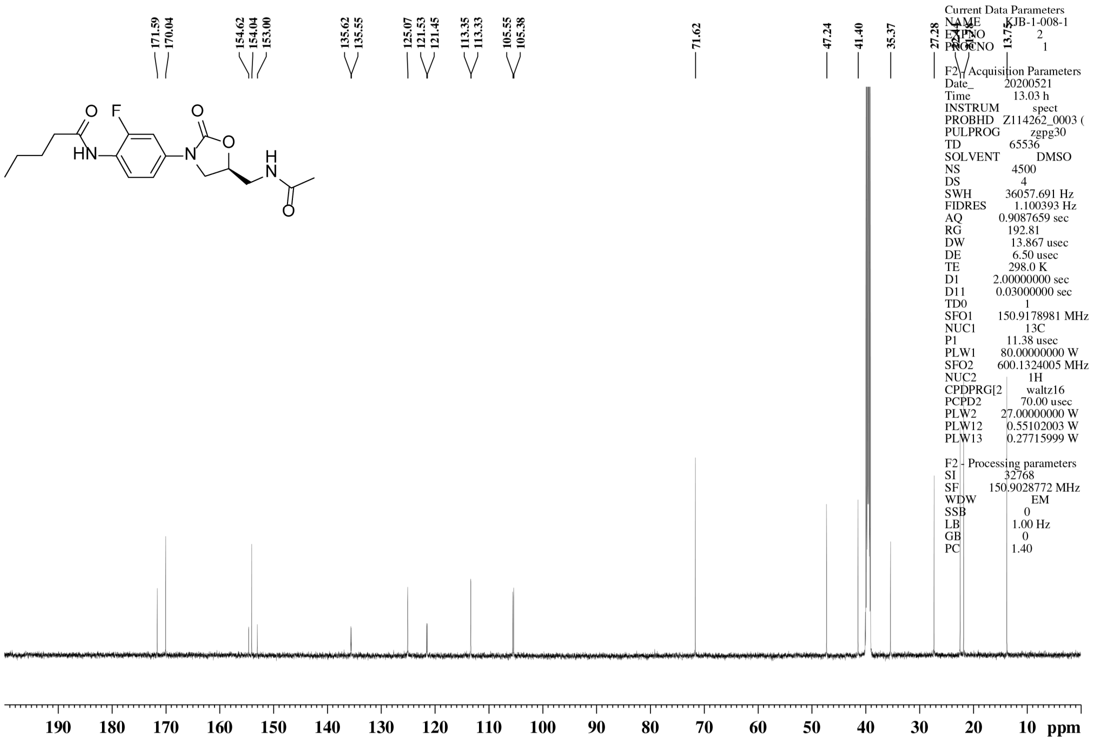

$^{1}$H NMR spectrum of **16**

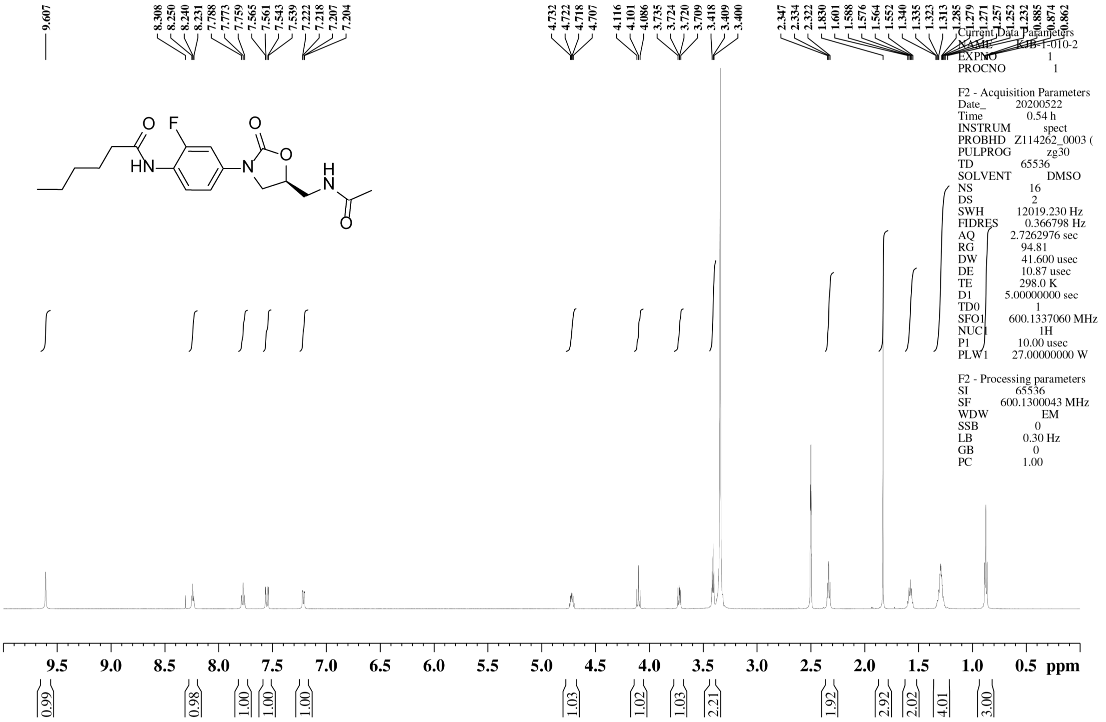


$^{13}$C NMR spectrum of **16**

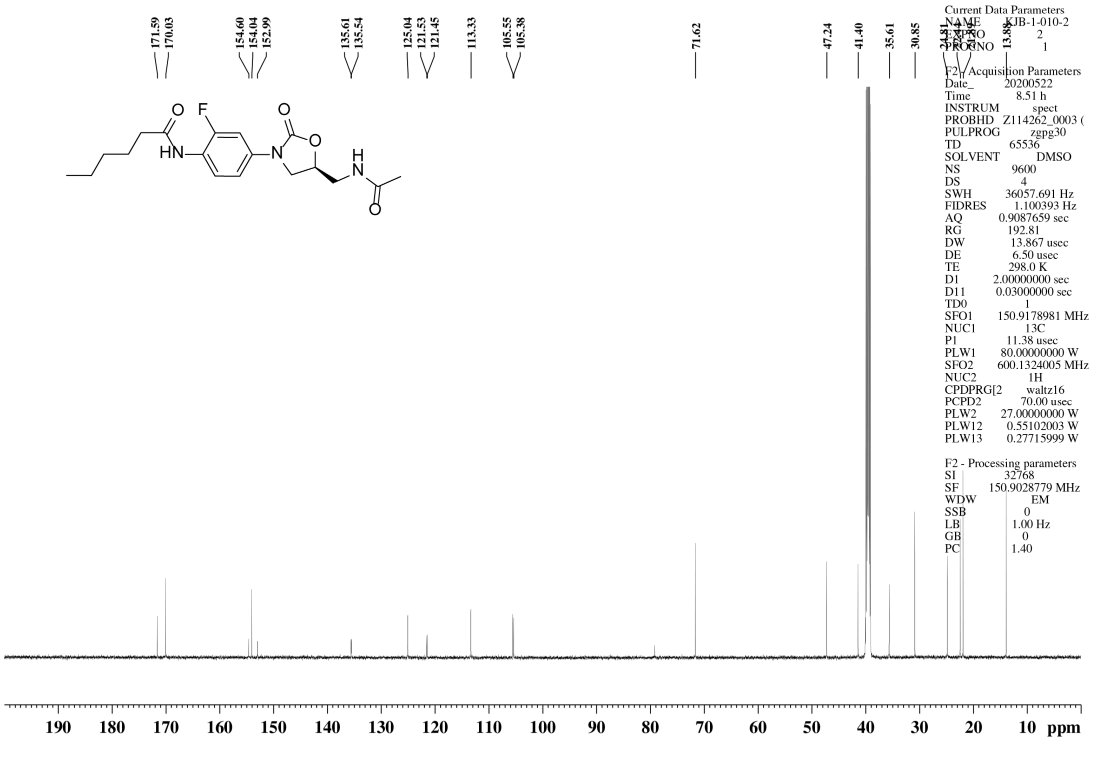

$^{1}$H NMR spectrum of **17**

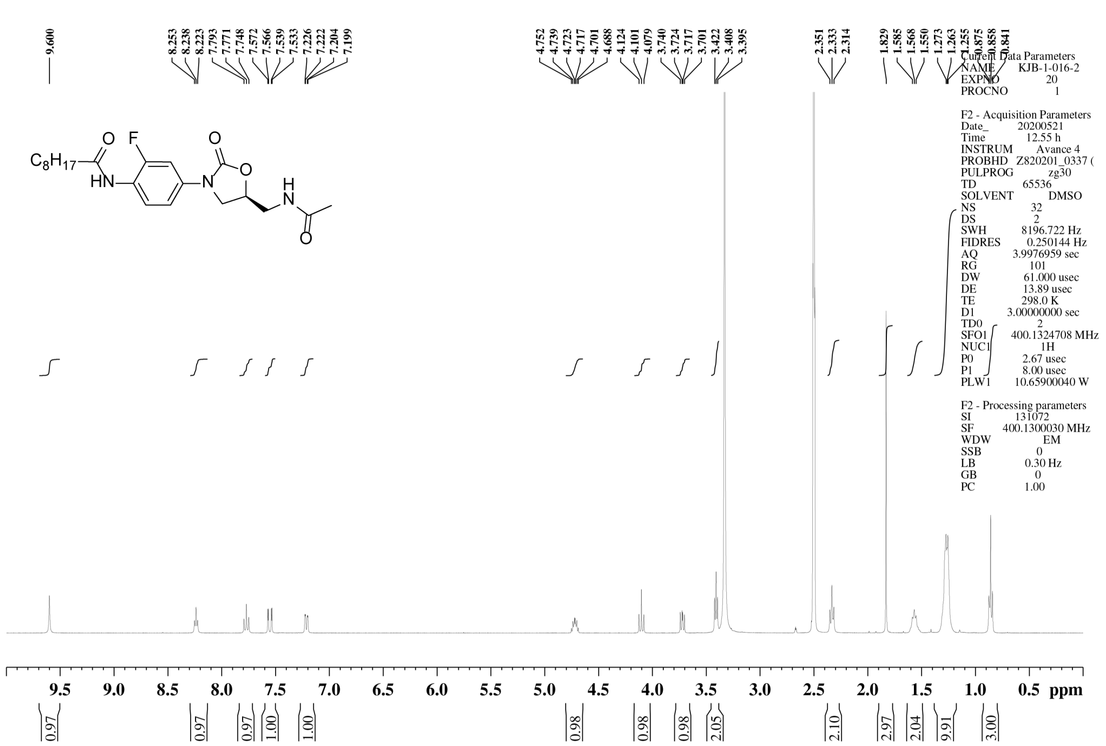


$^{13}$C NMR spectrum of **17**

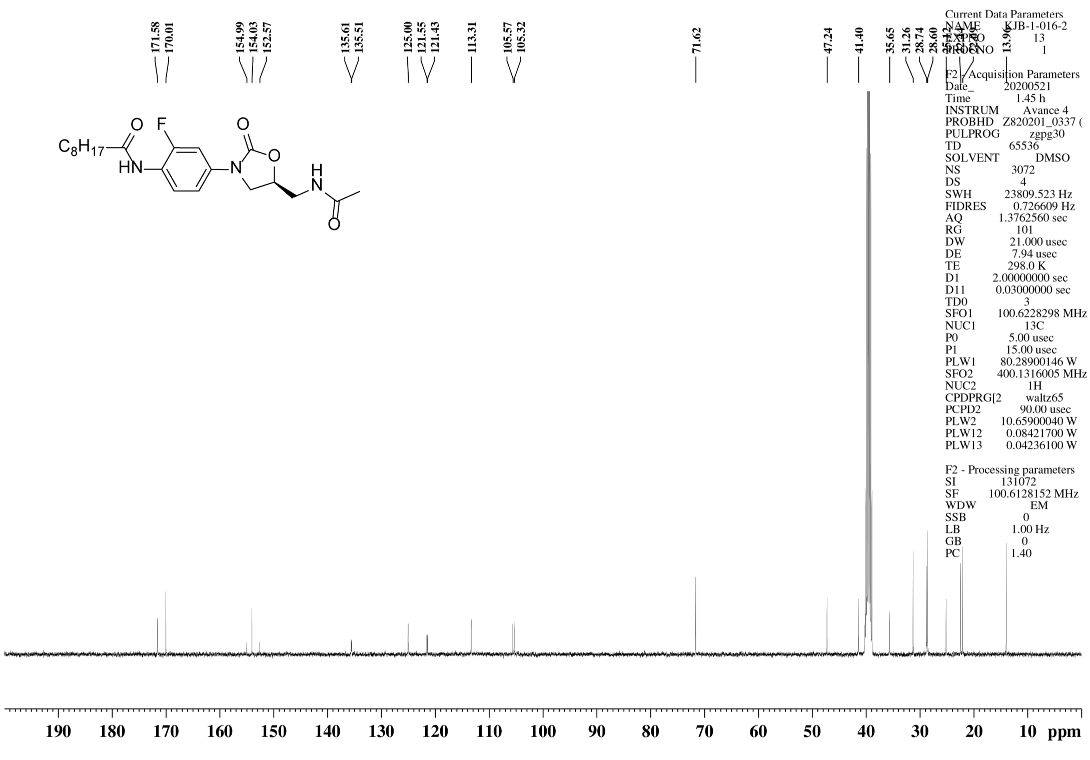

$^{1}$H NMR spectrum of **18**

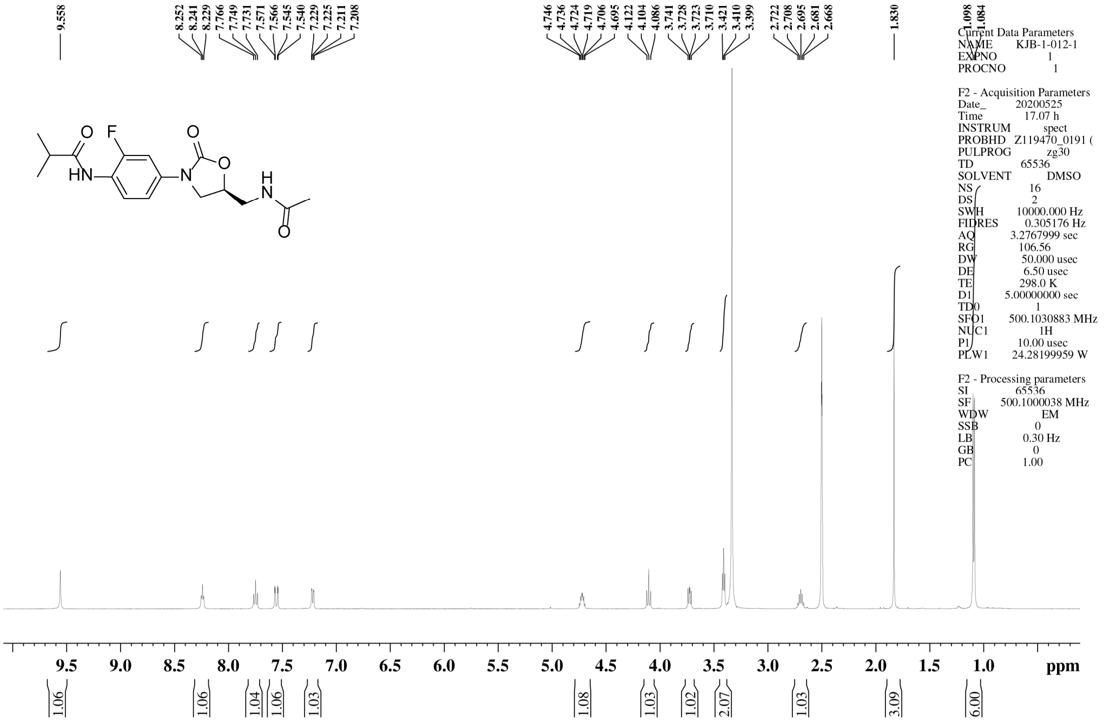


$^{13}$C NMR spectrum of **18**

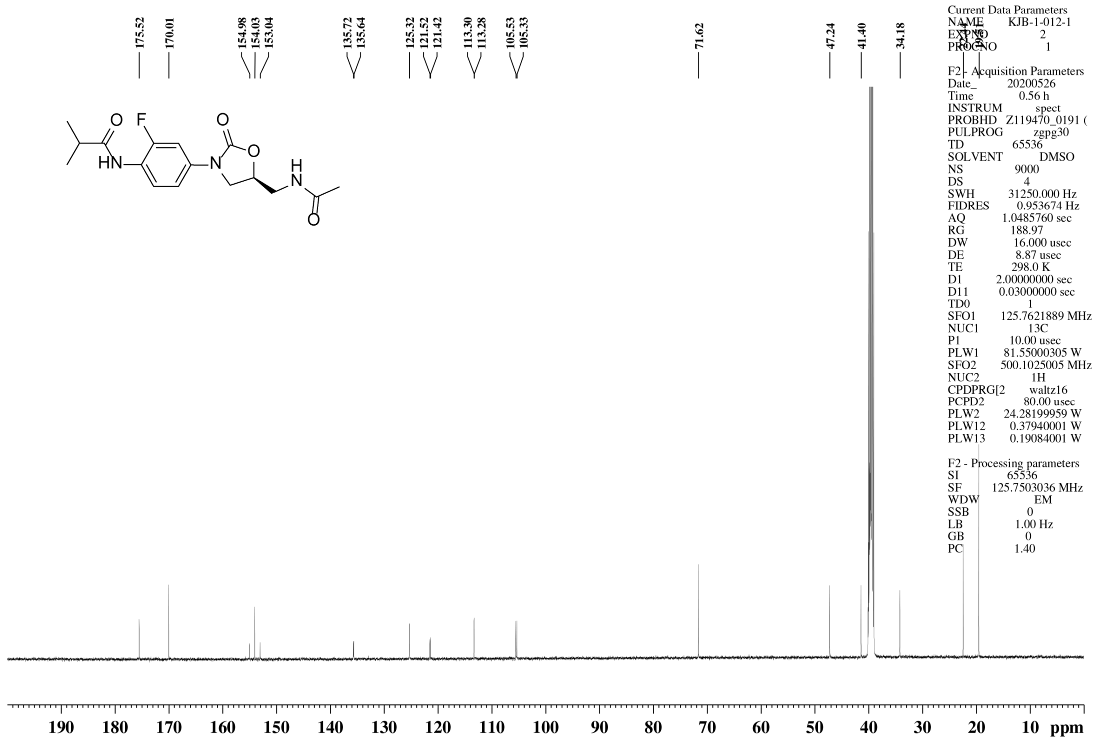

$^{1}$H NMR spectrum of **19**

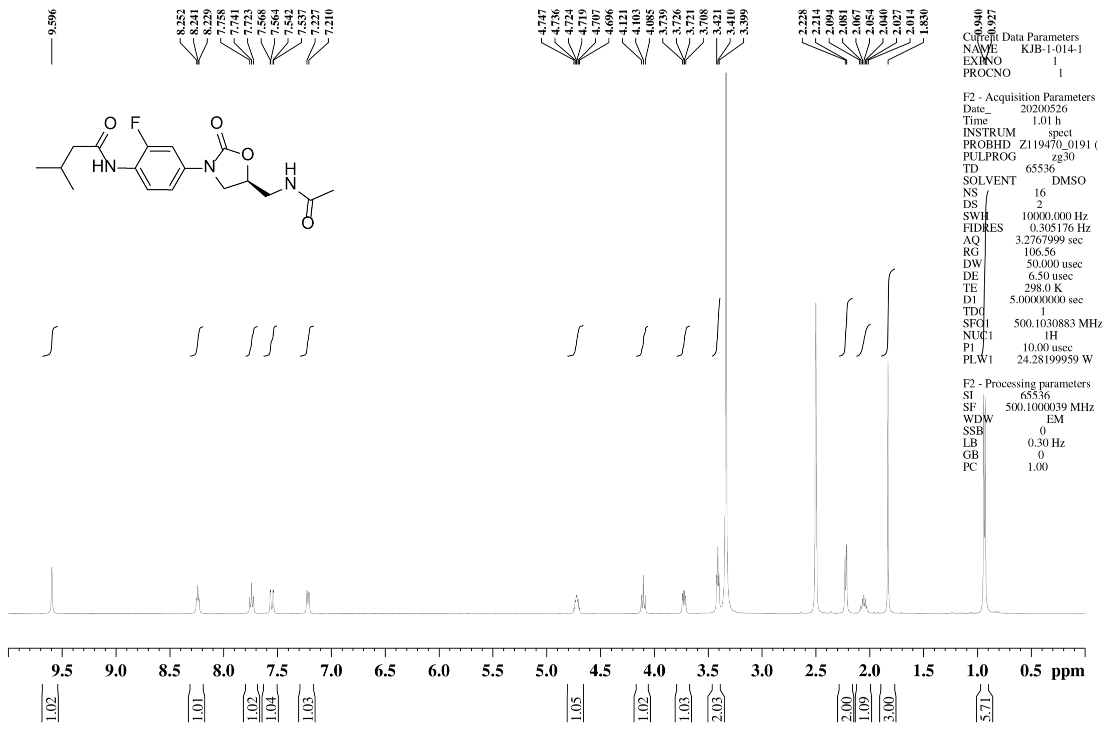

$^{13}$C NMR spectrum of **19**

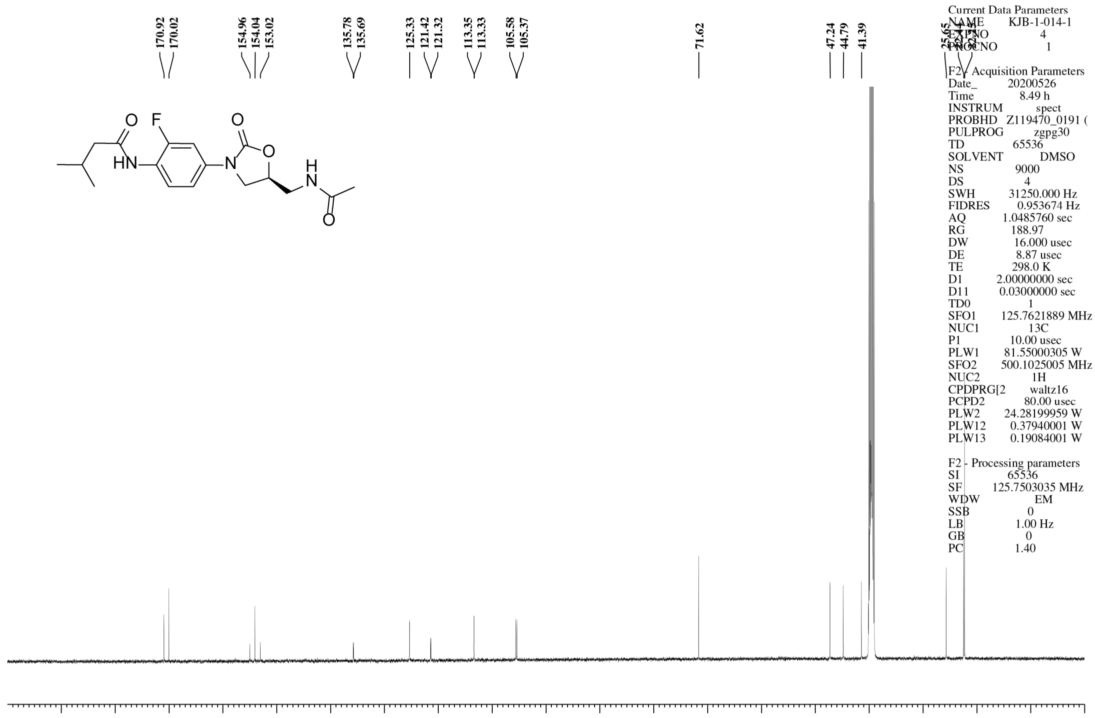

$^{1}$H NMR spectrum of **20**

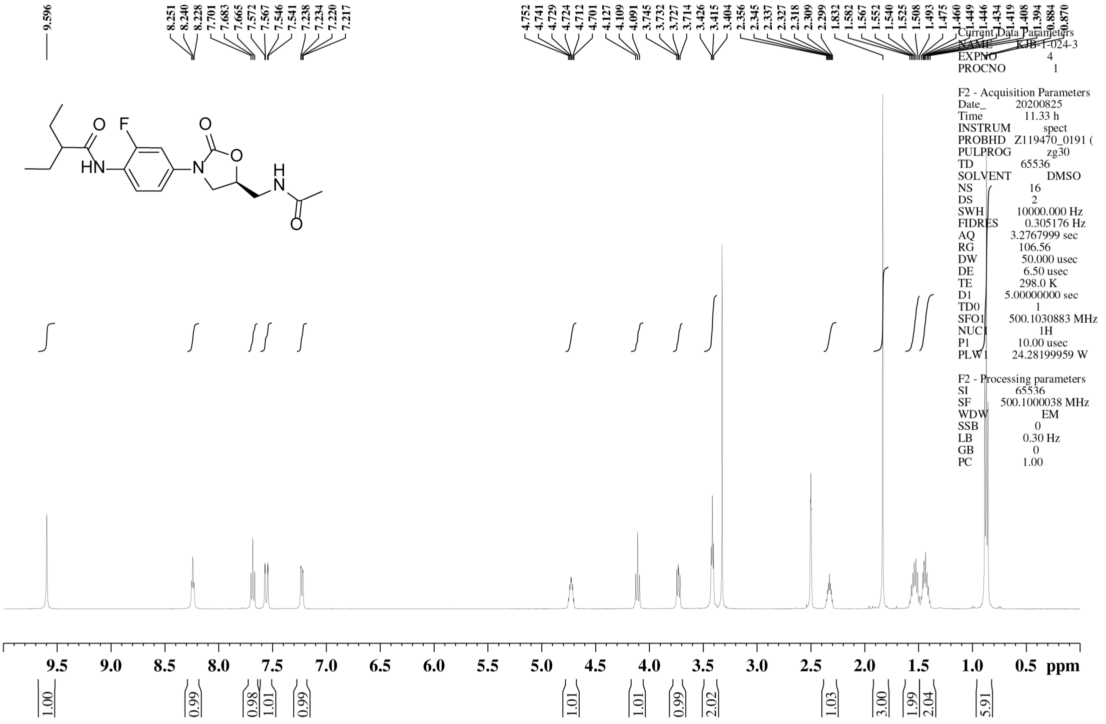

$^{13}$C NMR spectrum of **20**

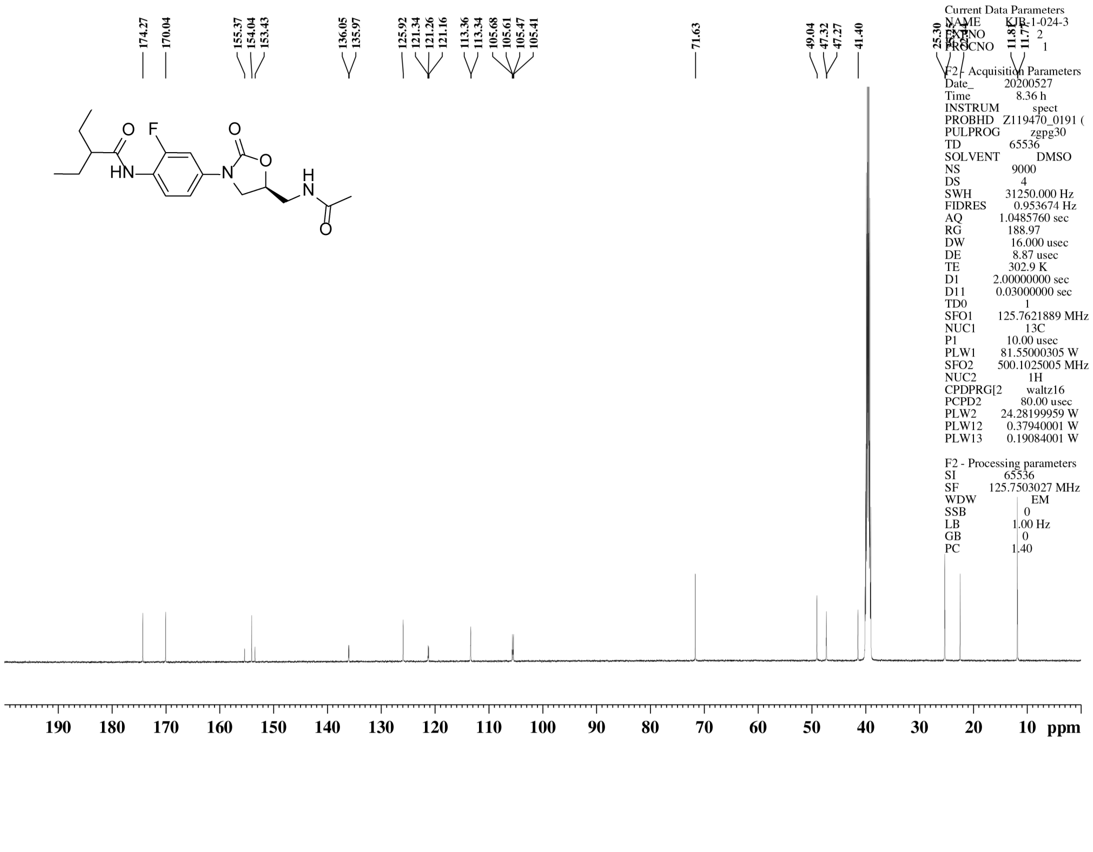

[1]H NMR spectrum of **21**

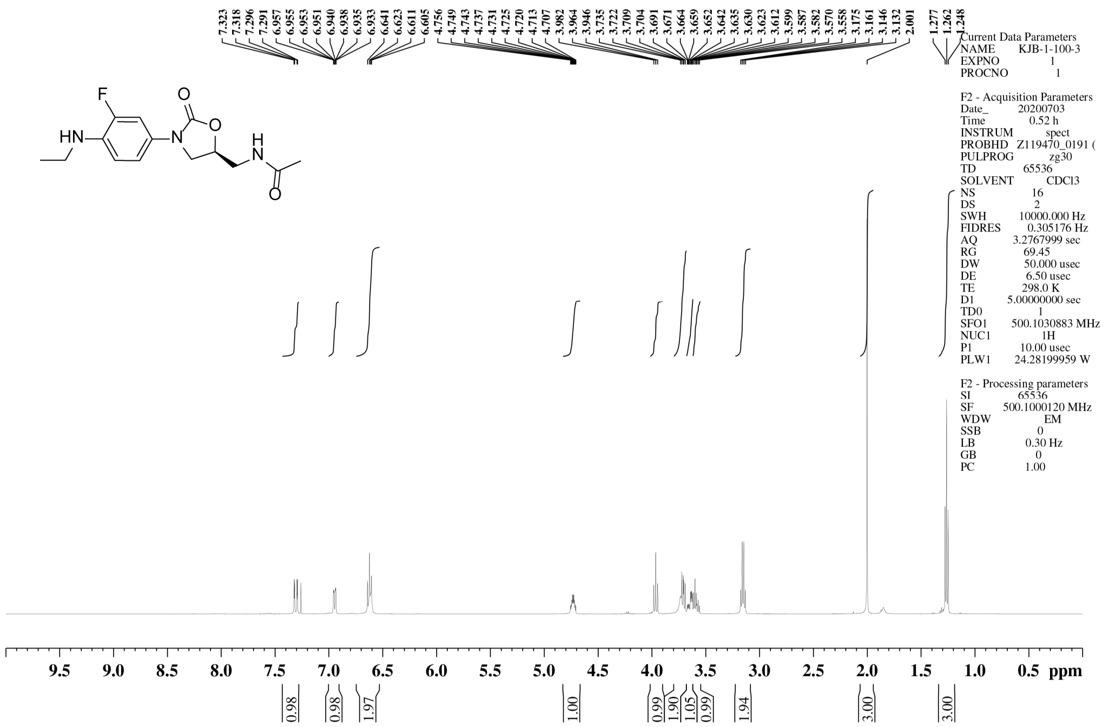

[13]C NMR spectrum of **21**

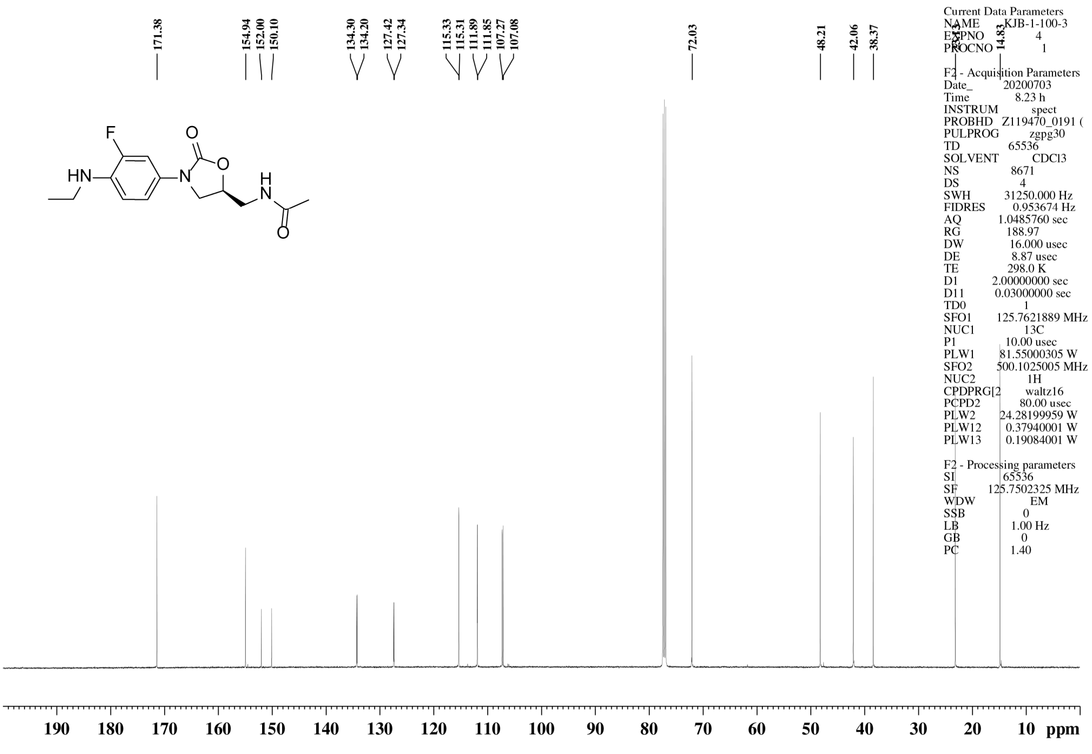

[1]H NMR spectrum of **22**

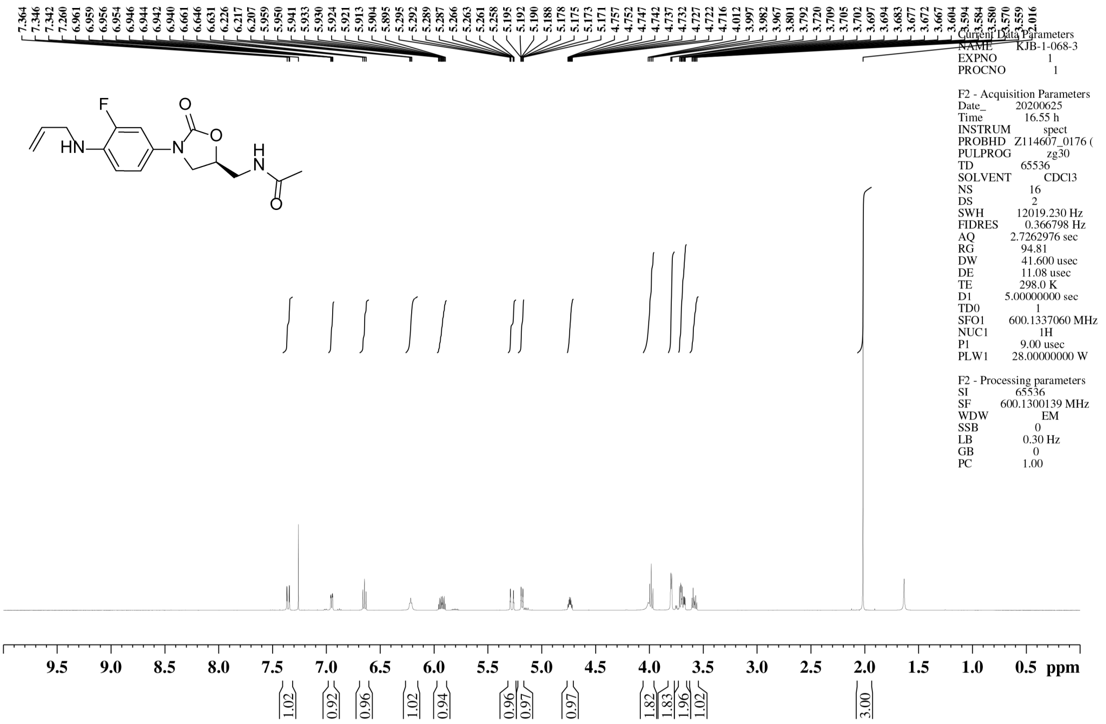


[13]C NMR spectrum of **22**

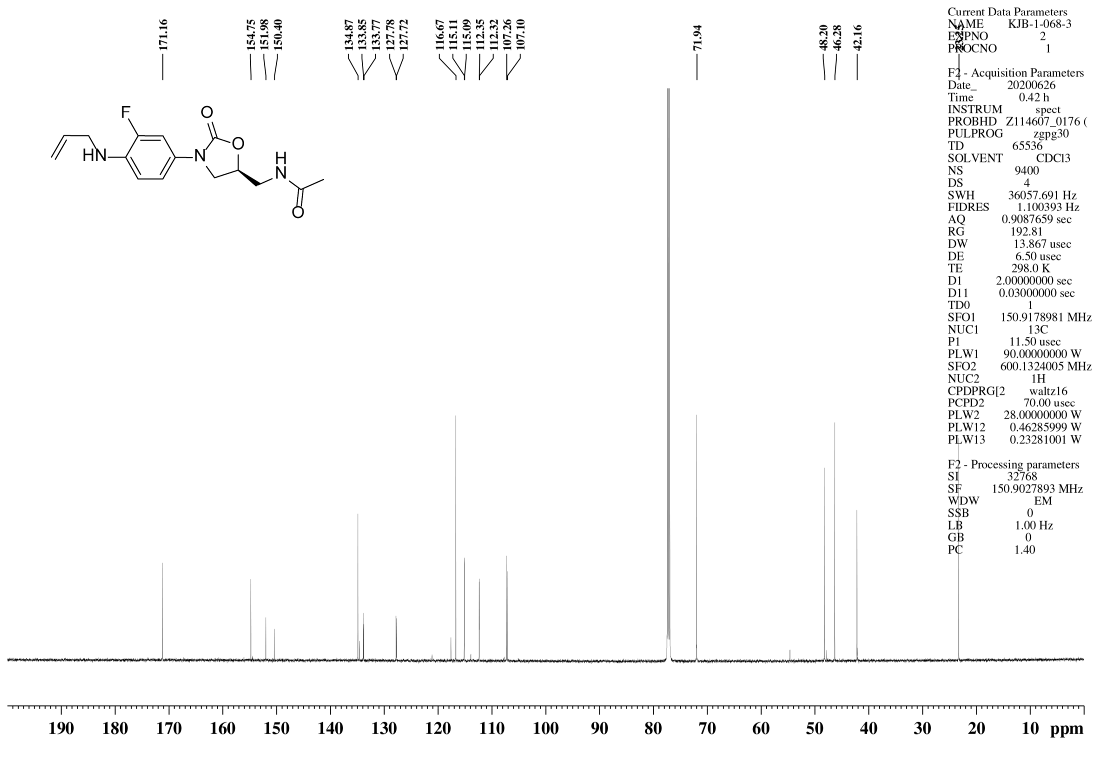

[1]H NMR spectrum of **24**

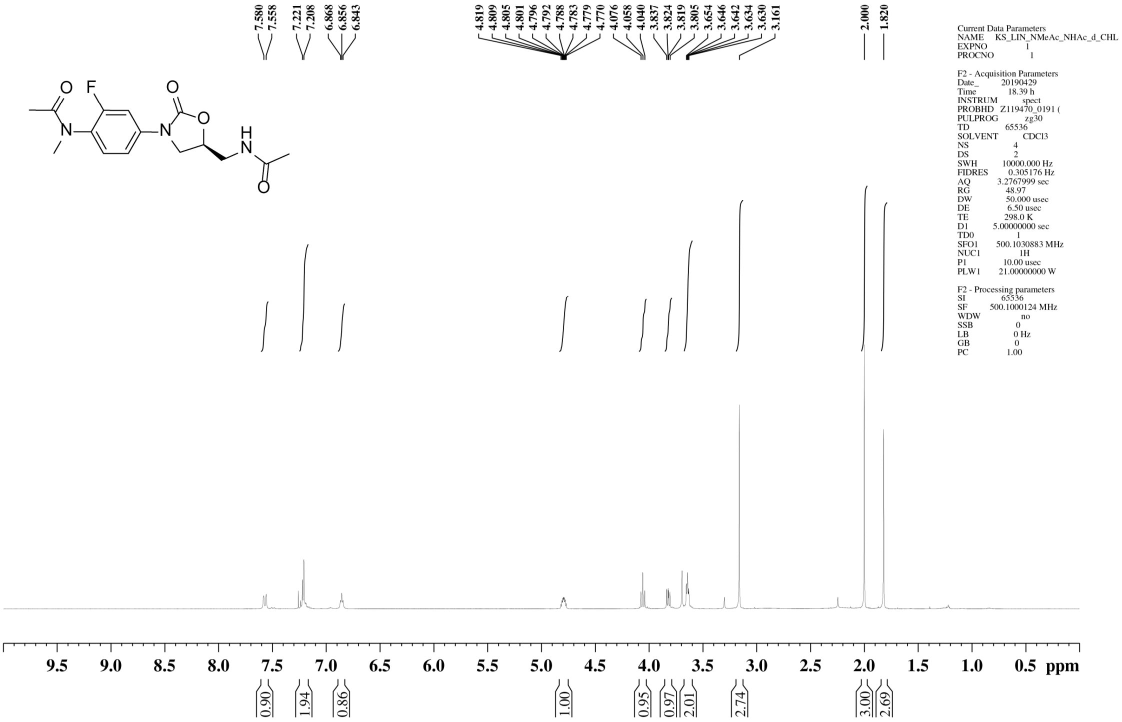


[13]C NMR spectrum of **24**

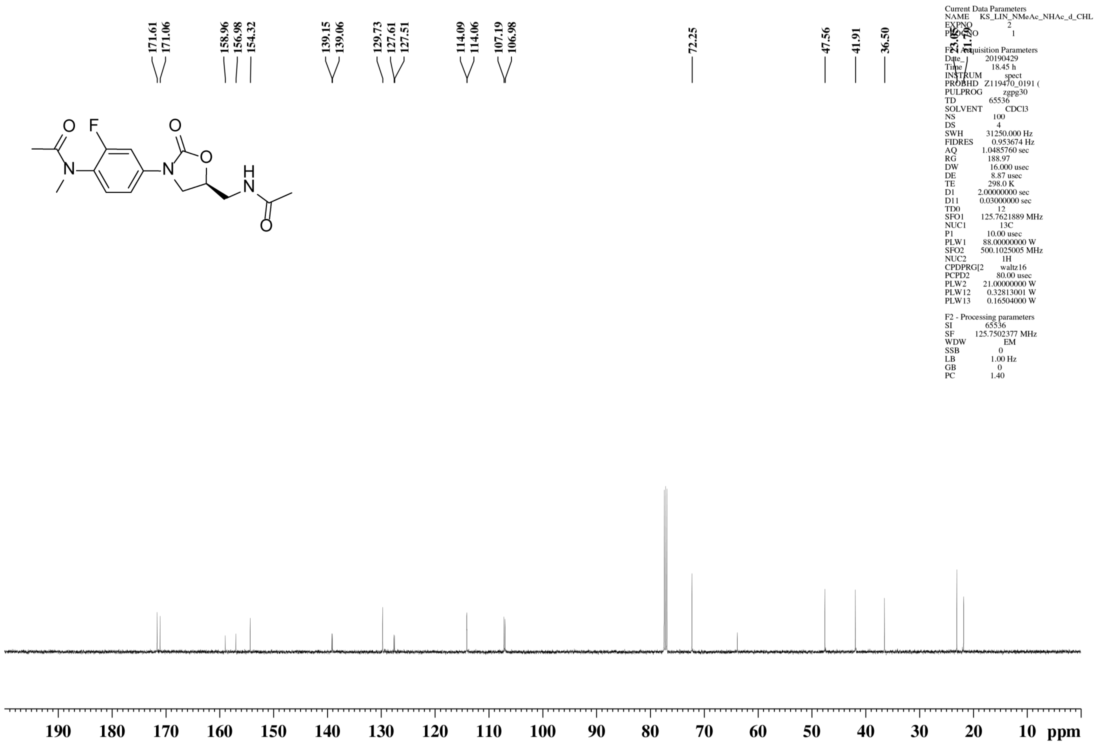

### [1]H NMR spectrum of **25**

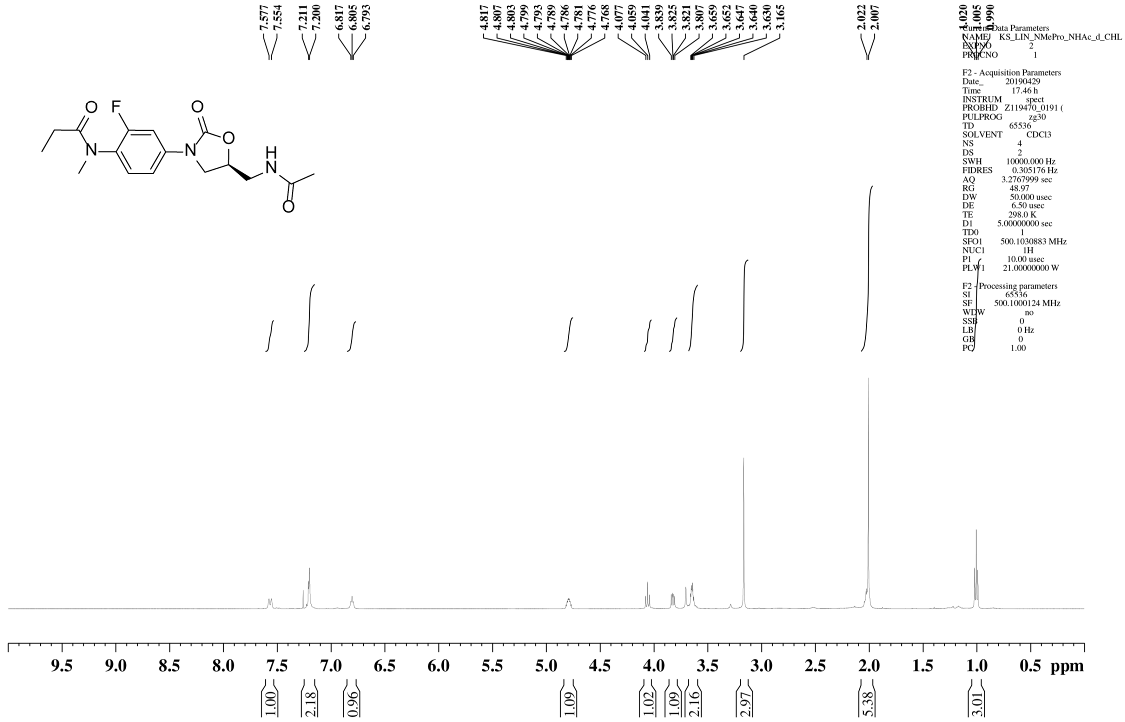


### [13]C NMR spectrum of **25**

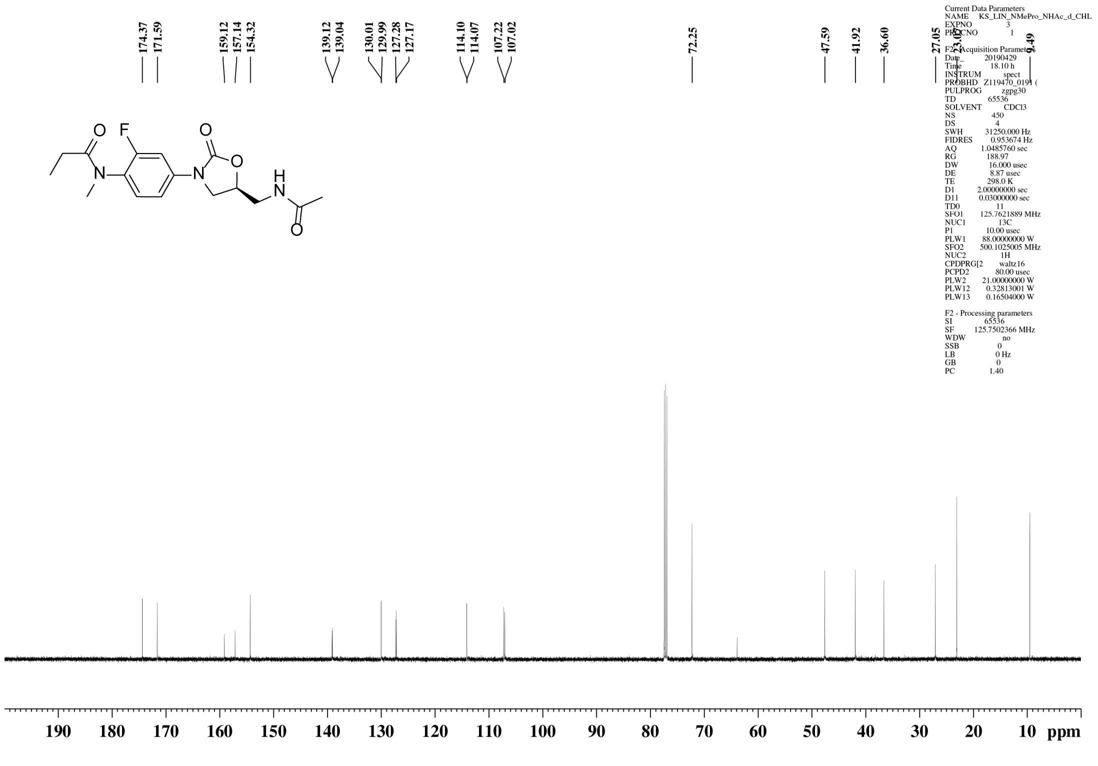

[1]H NMR spectrum of **26**

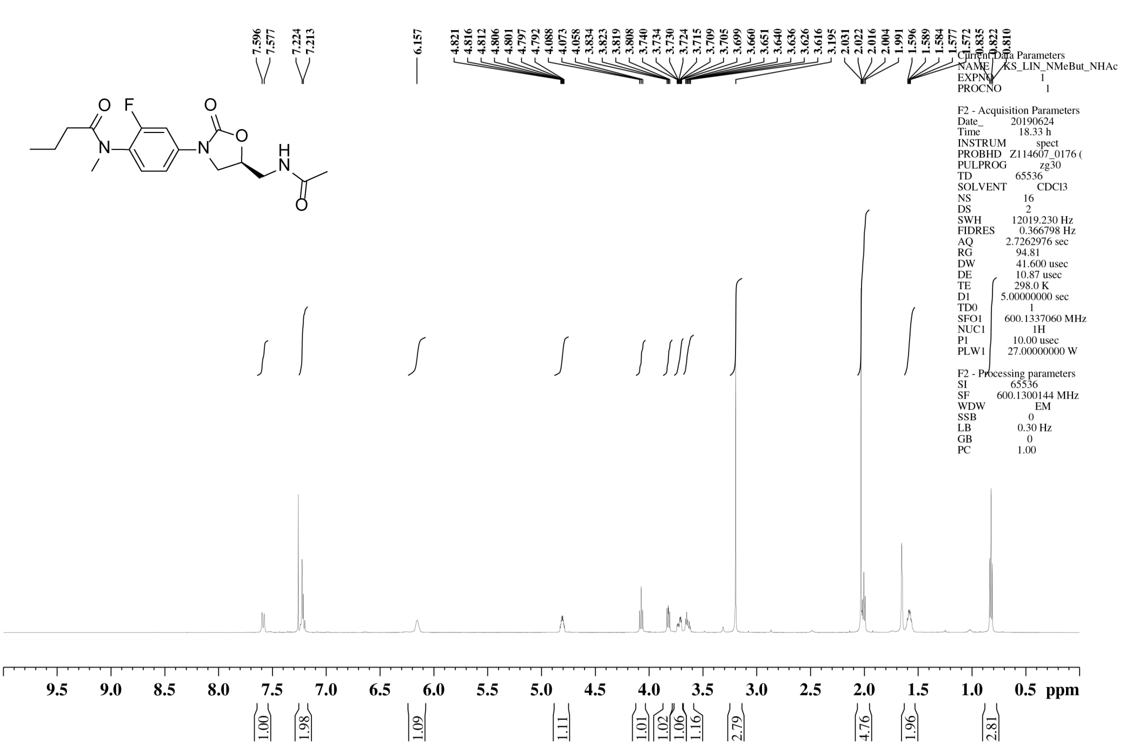


[13]C NMR spectrum of **26**

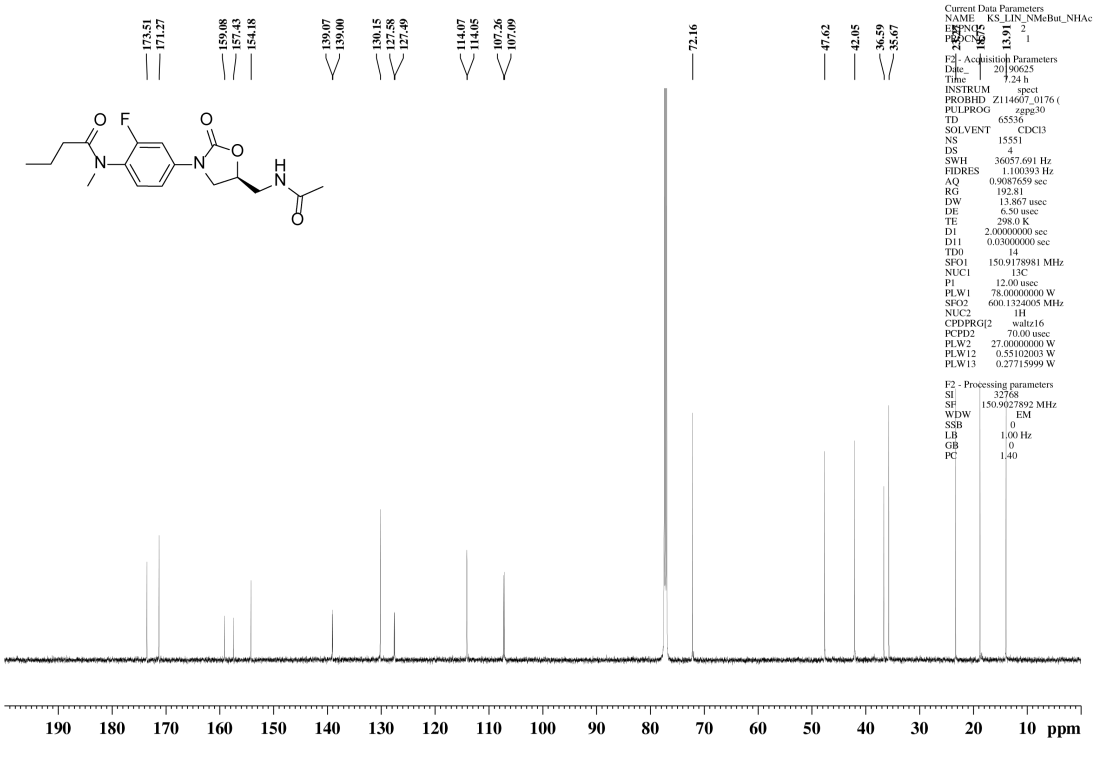

$^{1}$H NMR spectrum of **27**

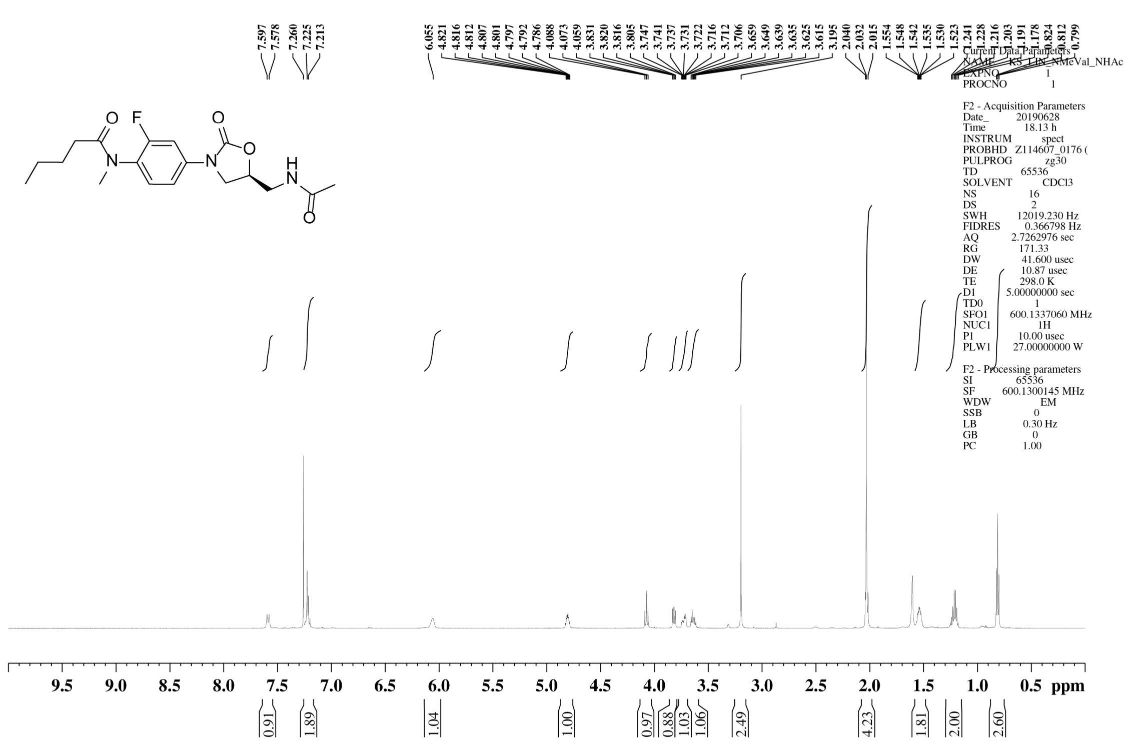


$^{13}$C NMR spectrum of **27**

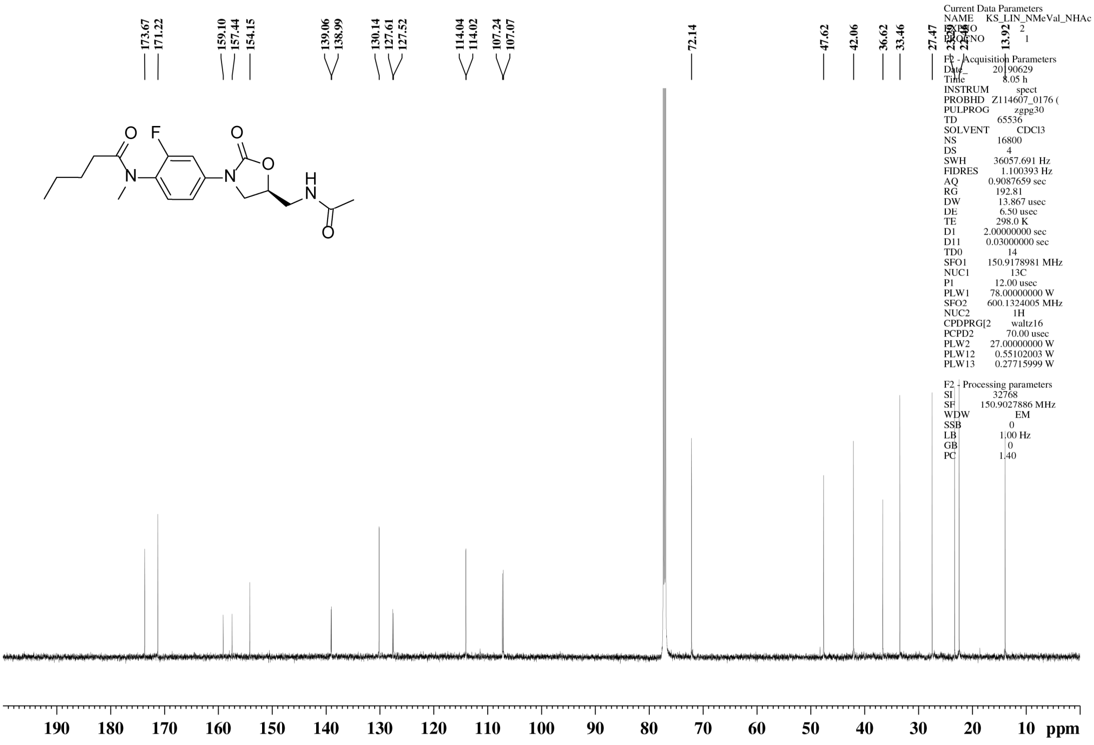

$^{1}$H NMR spectrum of **28**

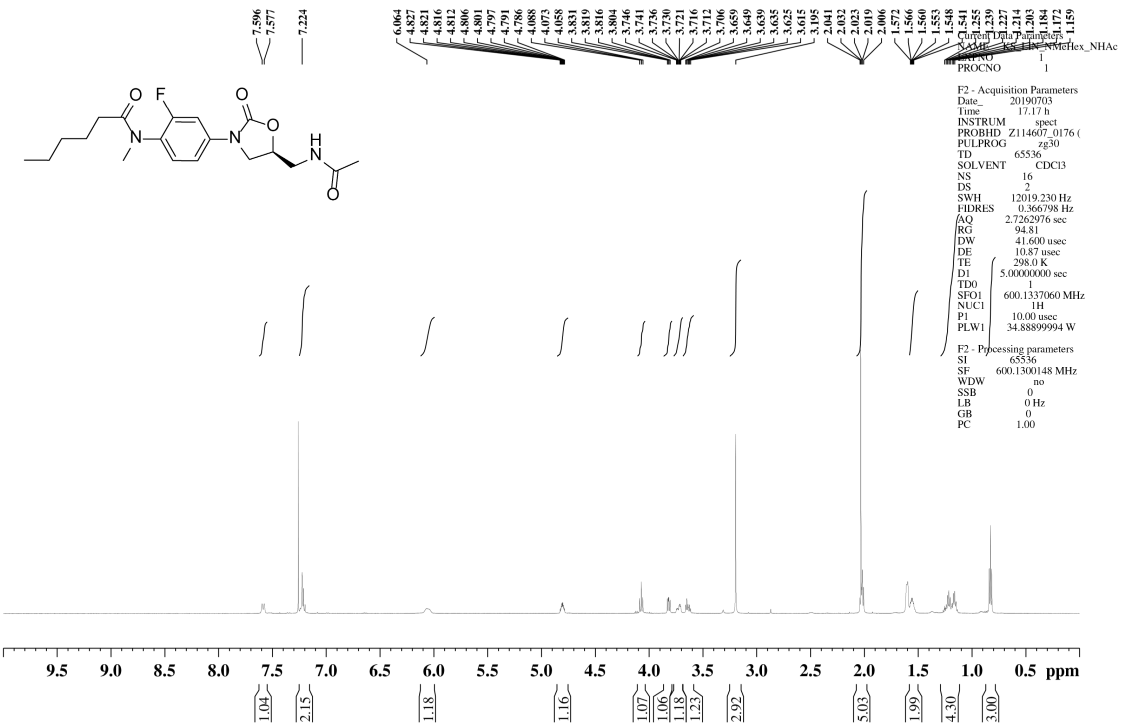

$^{13}$C NMR spectrum of **28**

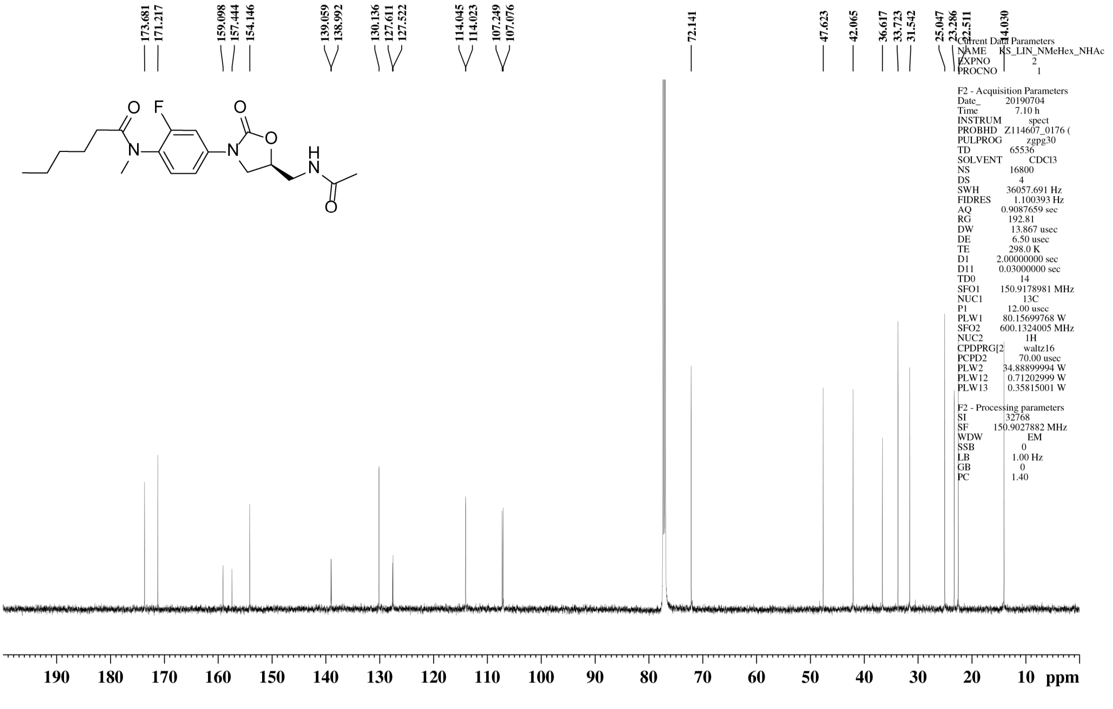

$^1$H NMR spectrum of **29**

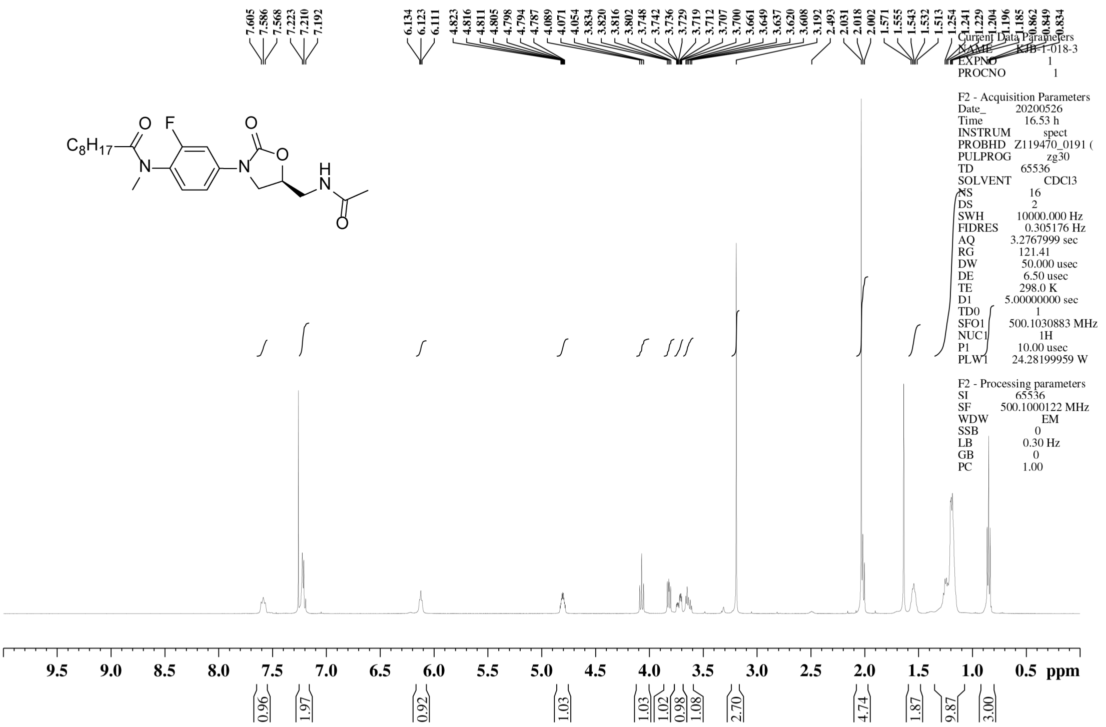

$^{13}$C NMR spectrum of **29**

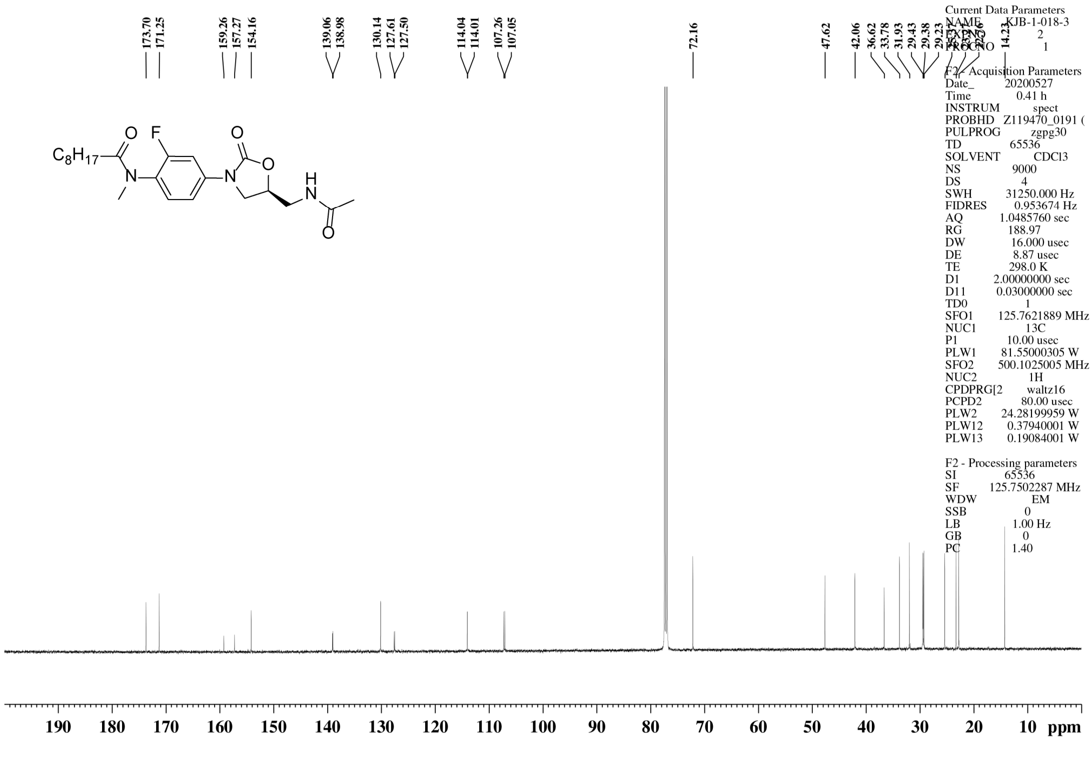

$^{1}$H NMR spectrum of **30**

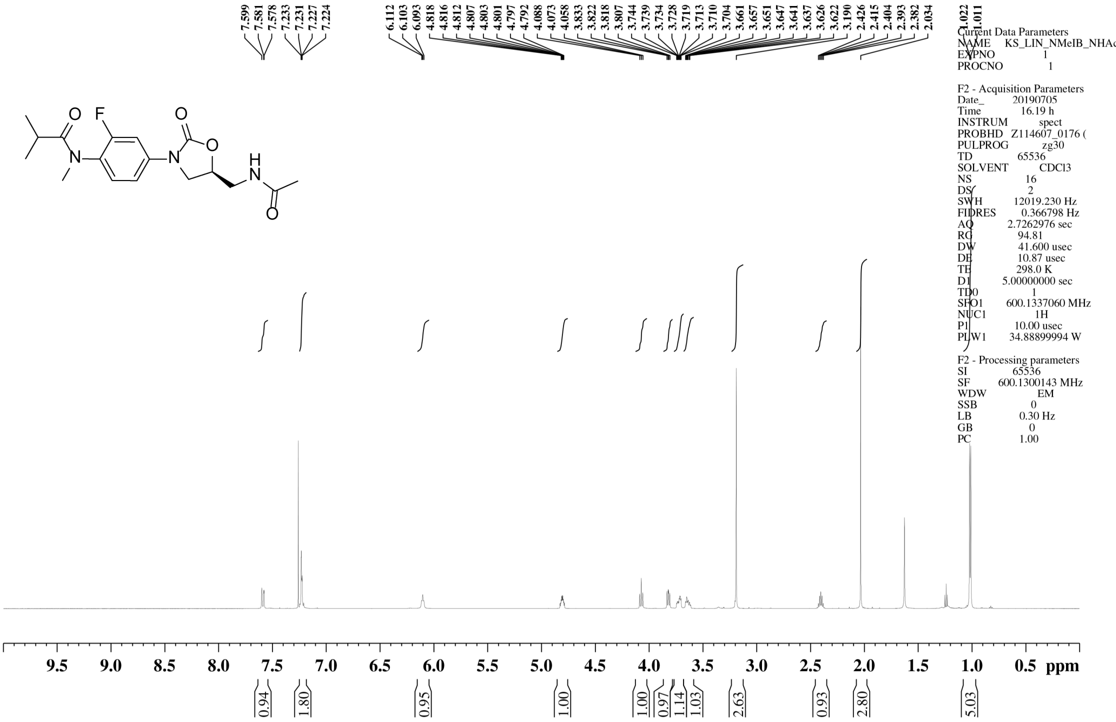


$^{13}$C NMR spectrum of **30**

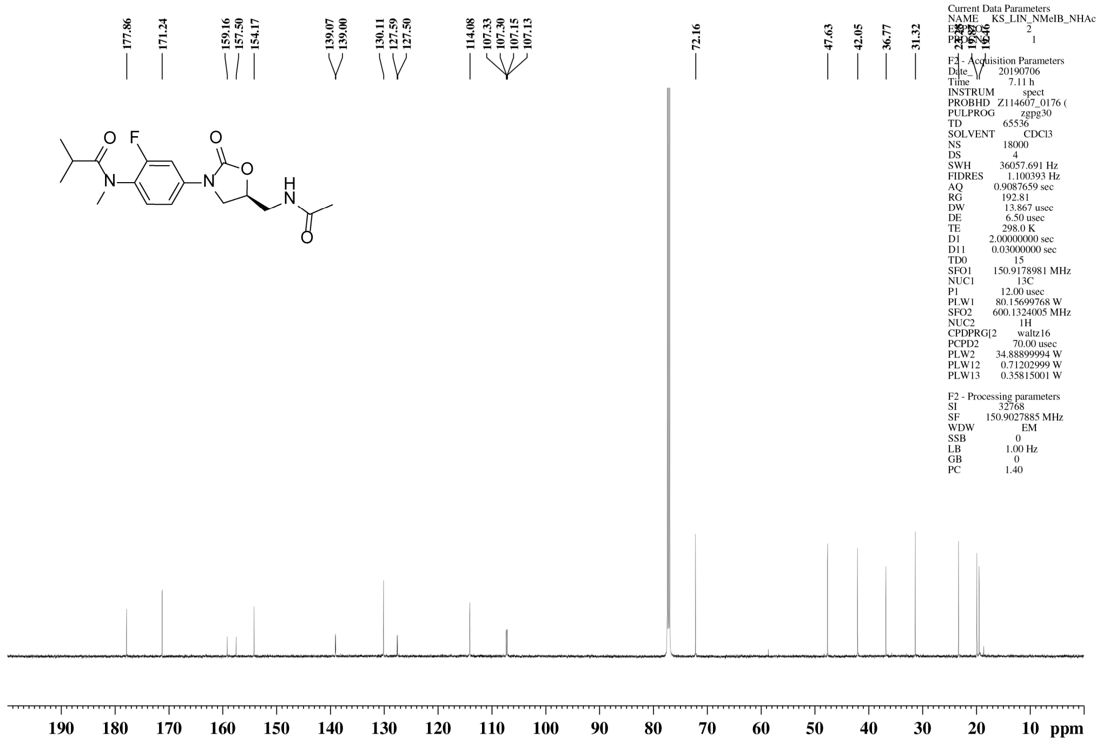

$^{1}$H NMR spectrum of **31**

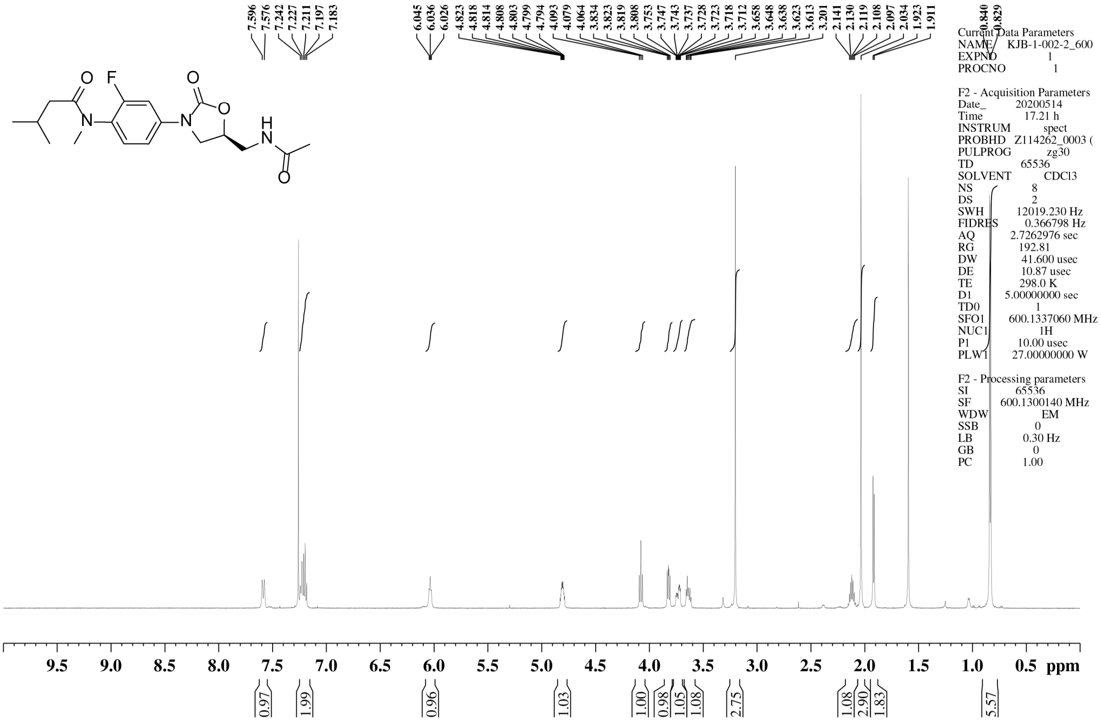


$^{13}$C NMR spectrum of **31**

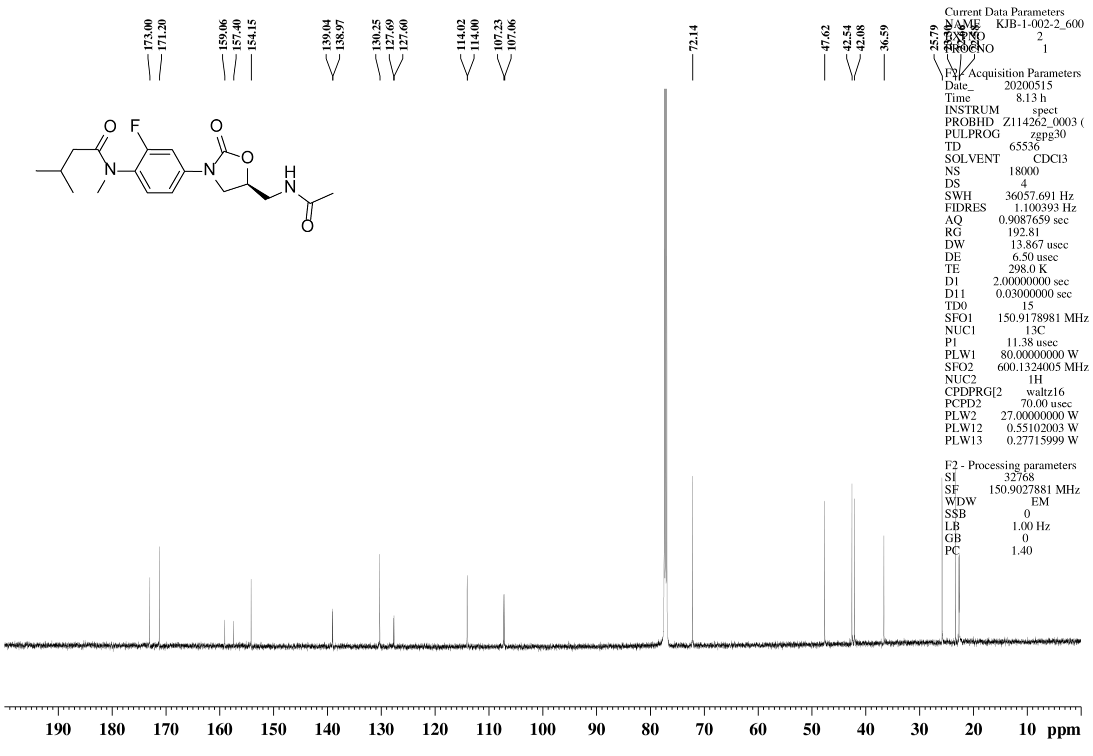

[1]H NMR spectrum of **32**

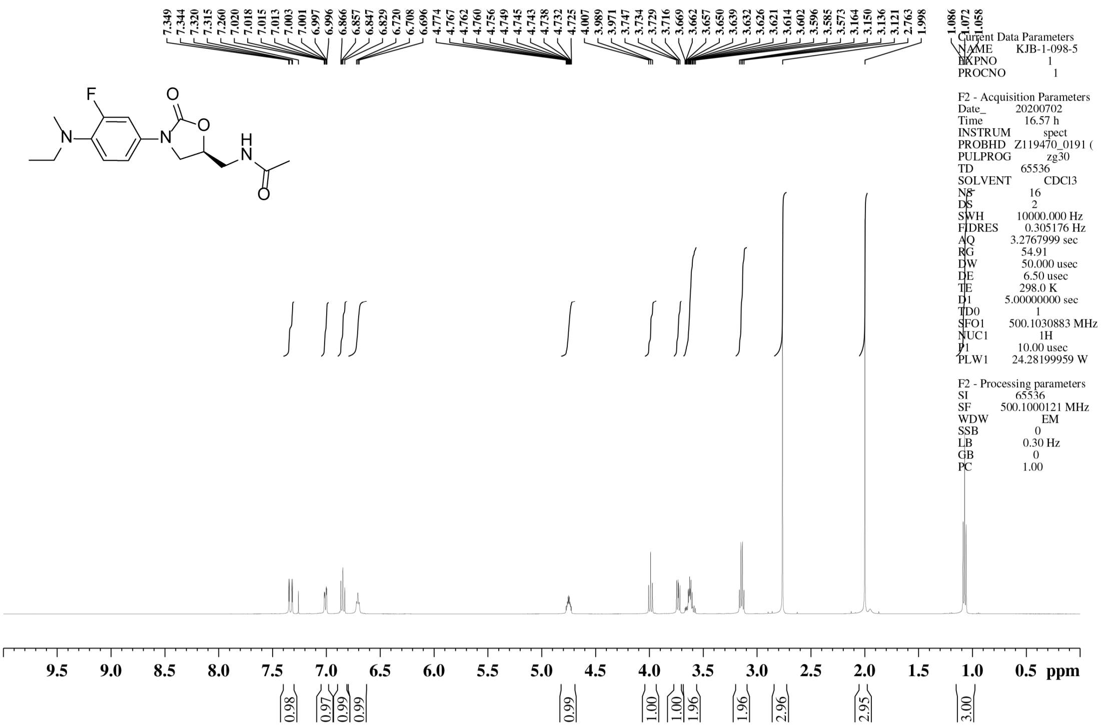


[13]C NMR spectrum of **32**

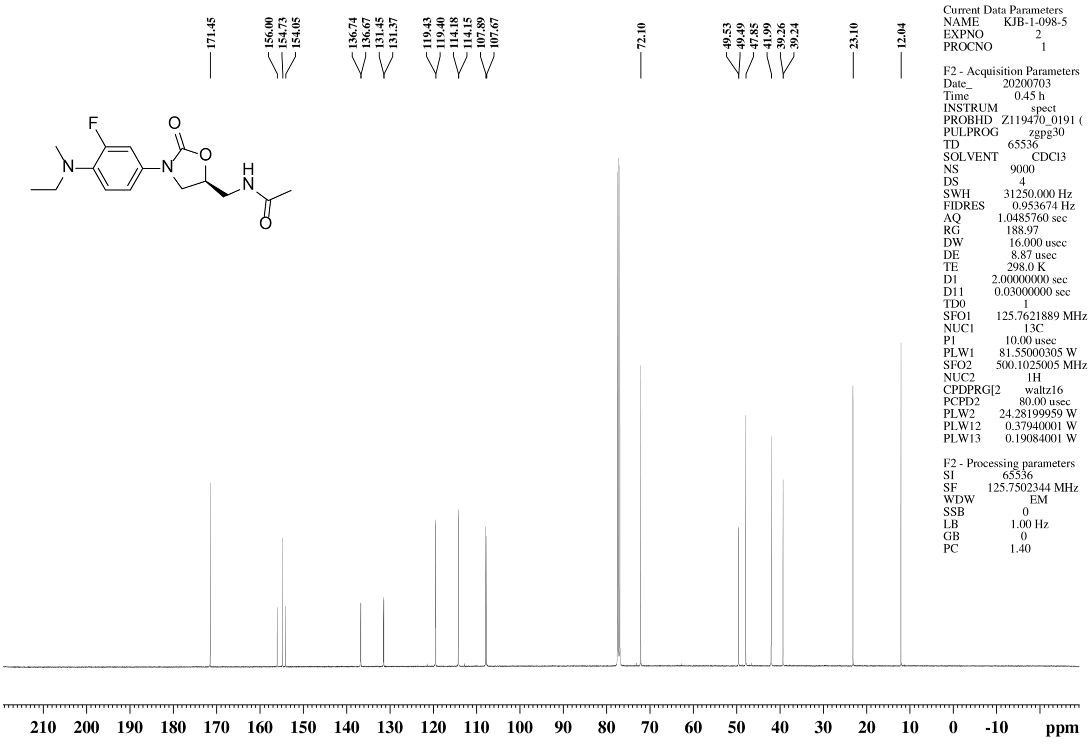

$^{1}$H NMR spectrum of **33**

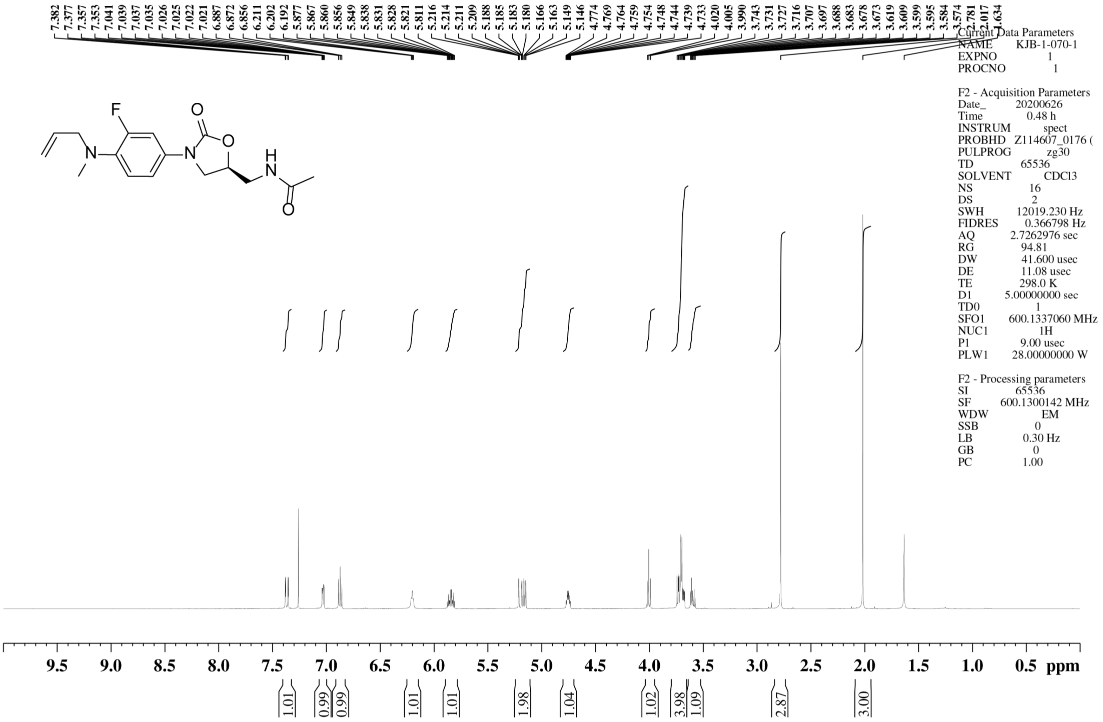


$^{13}$C NMR spectrum of **33**

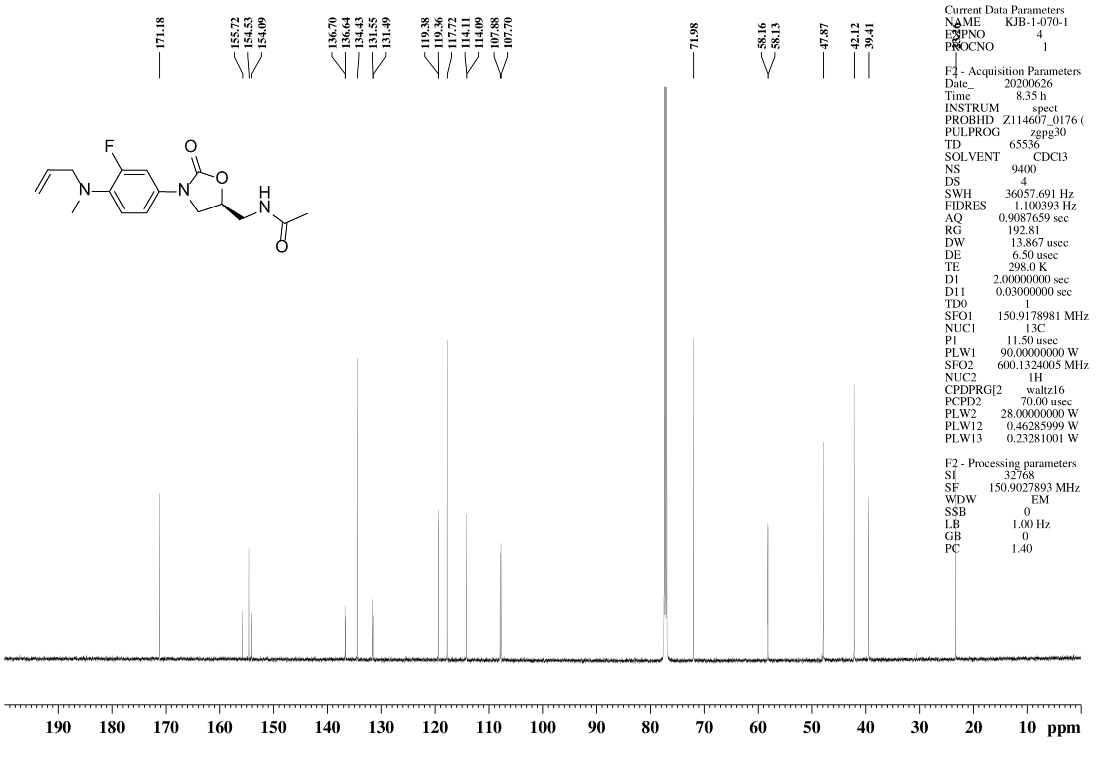

### [1]H NMR spectrum of **34**

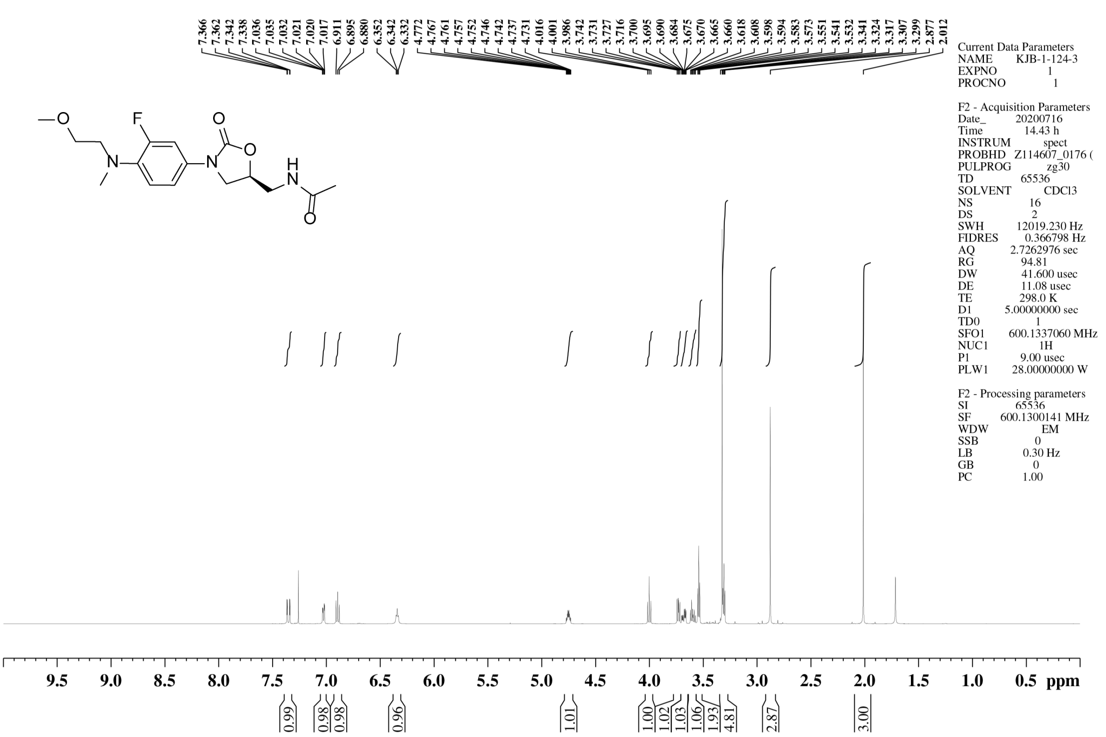


### [13]C NMR spectrum of **34**

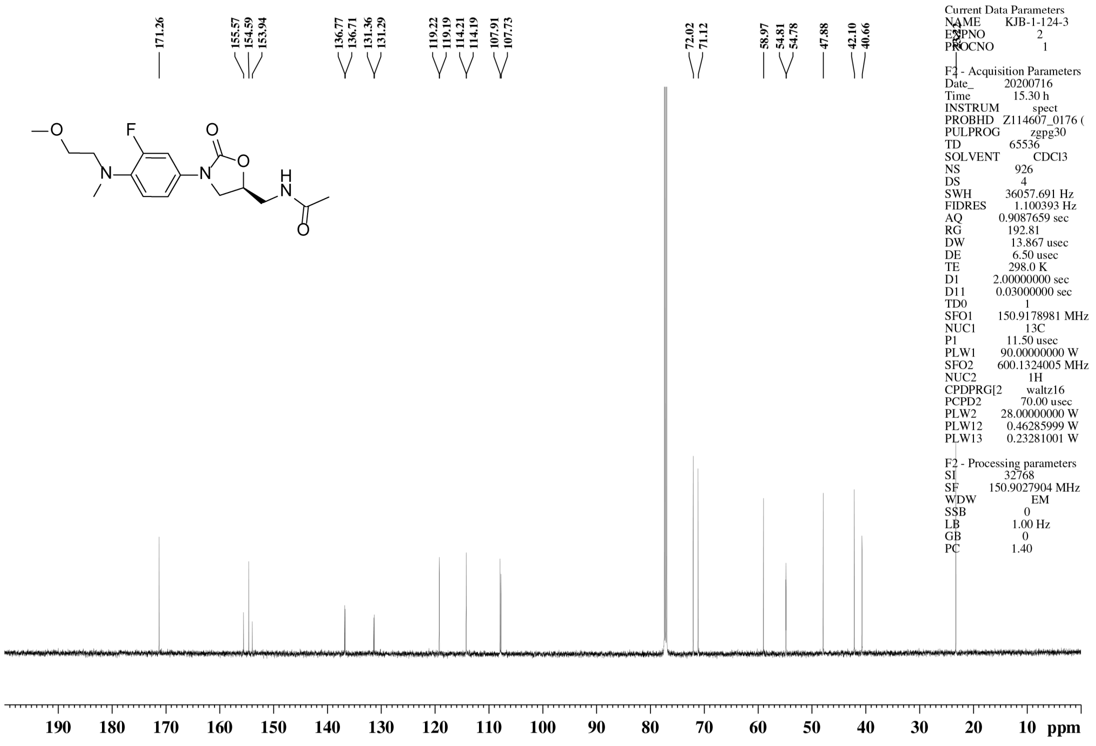

### $^{1}$H NMR spectrum of **35**

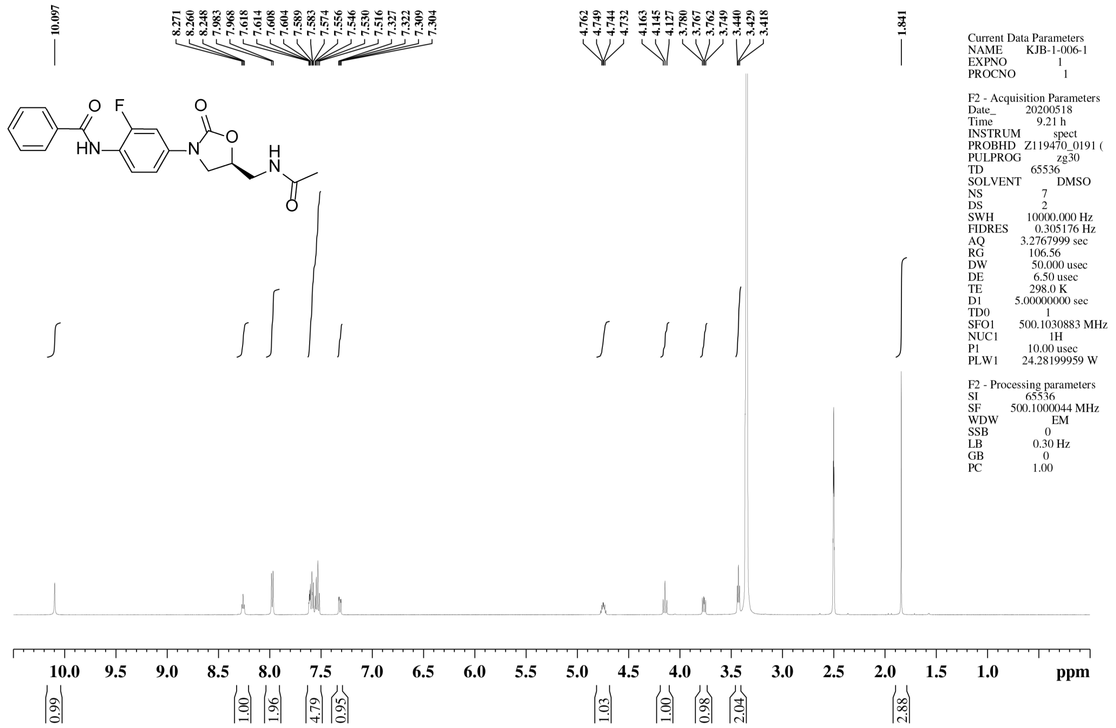


### $^{13}$C NMR spectrum of **35**

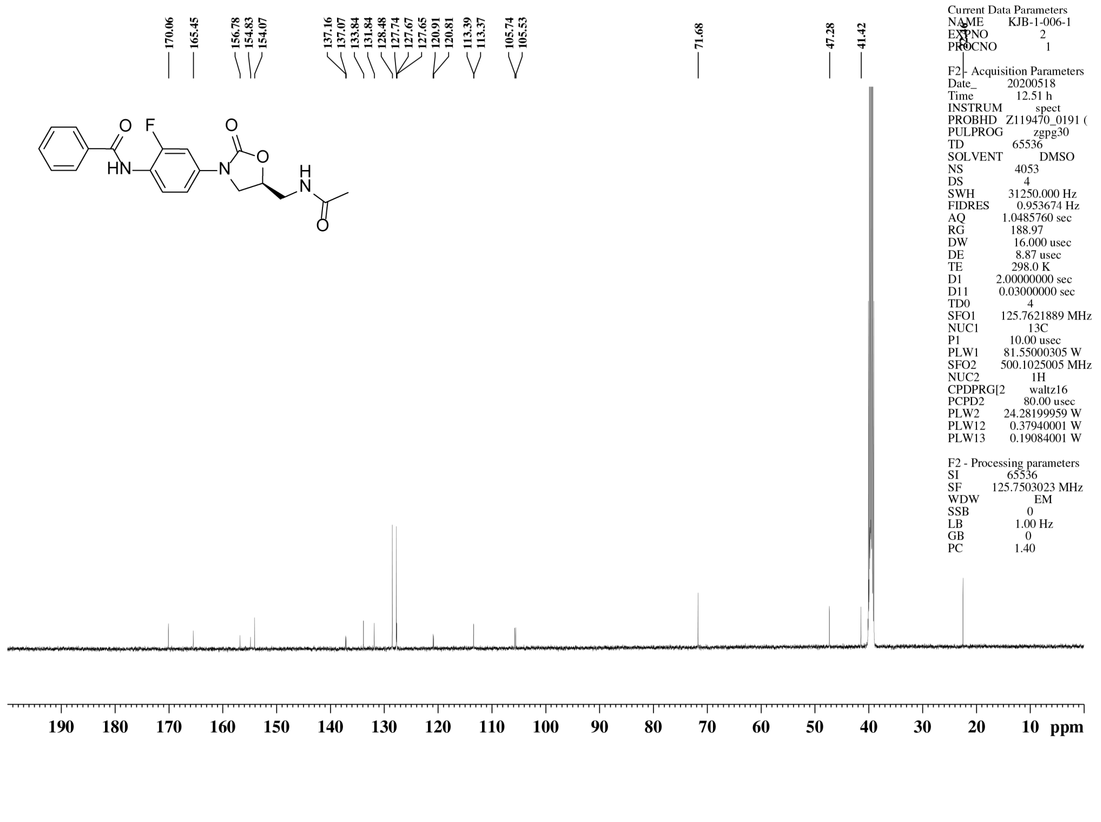

$^{1}$H NMR spectrum of **36**

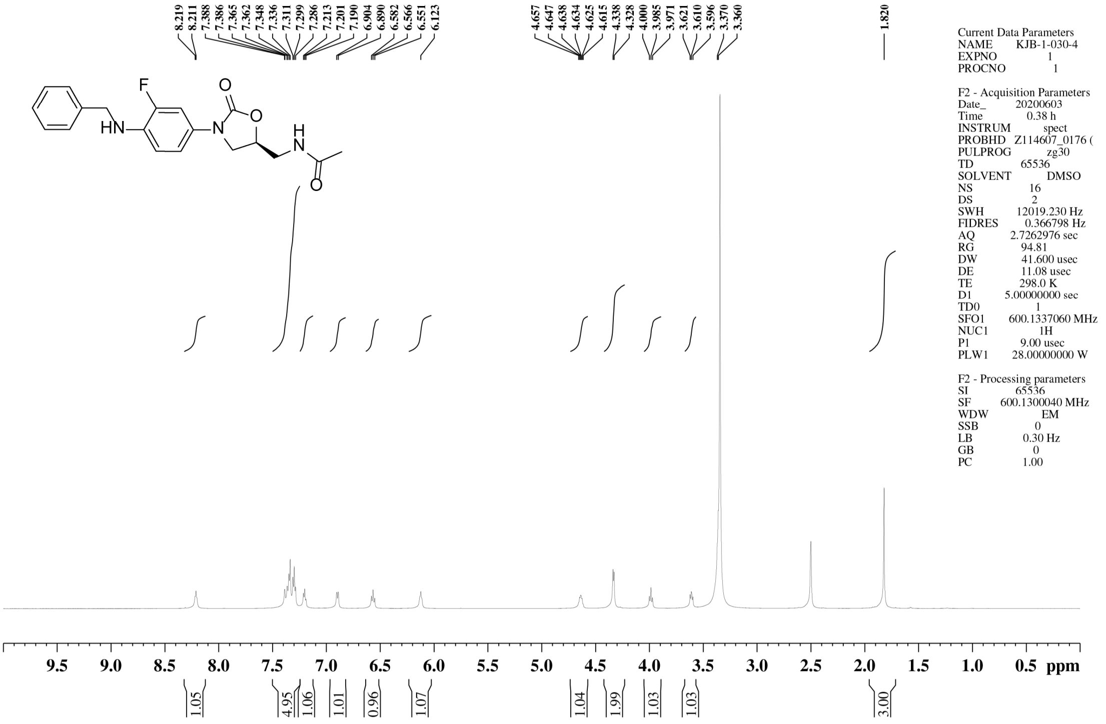

$^{13}$C NMR spectrum of **36**

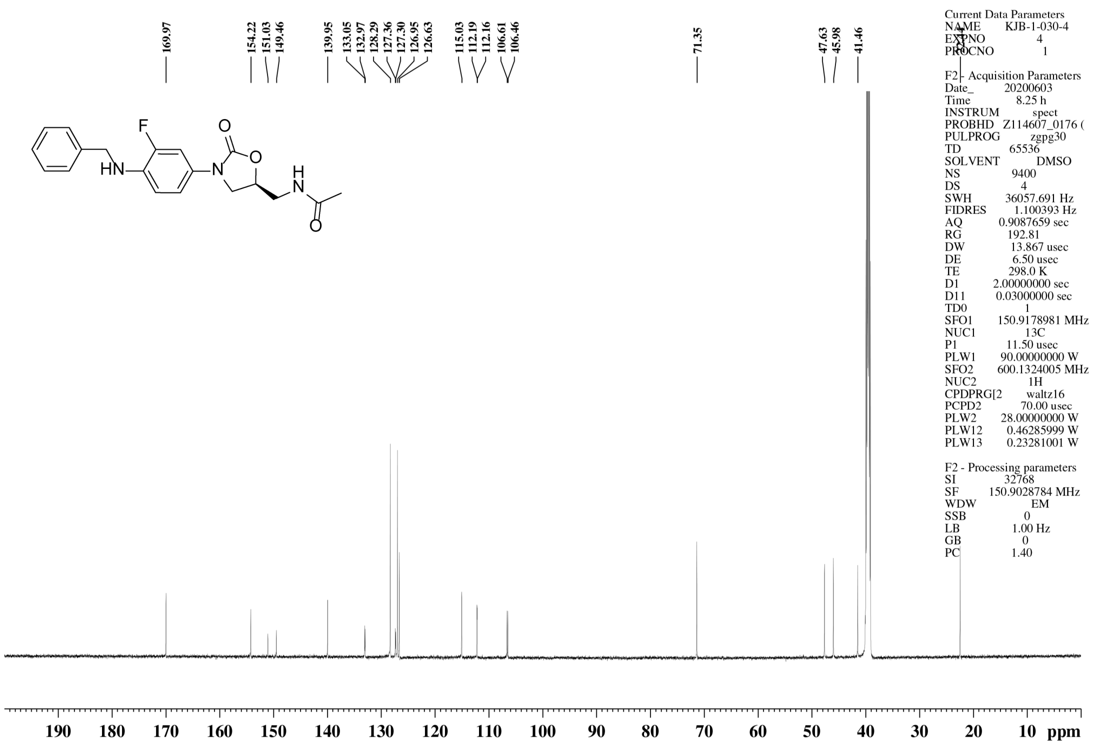

$^1$H NMR spectrum of **37**

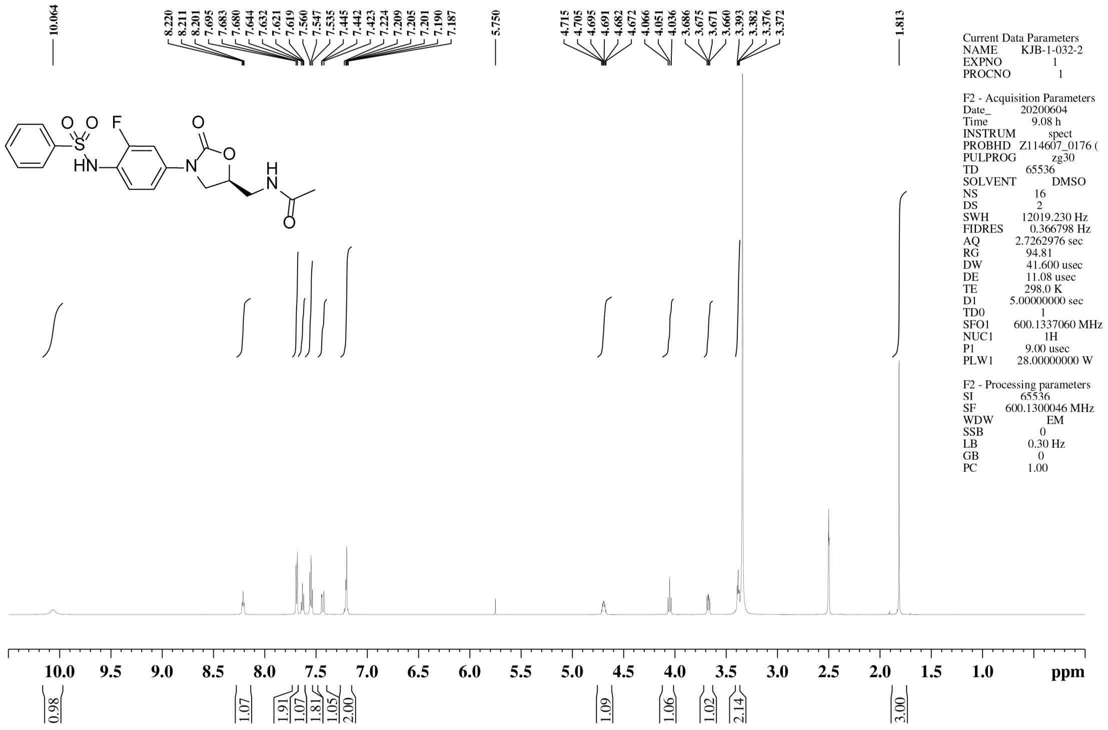


$^{13}$C NMR spectrum of **37**

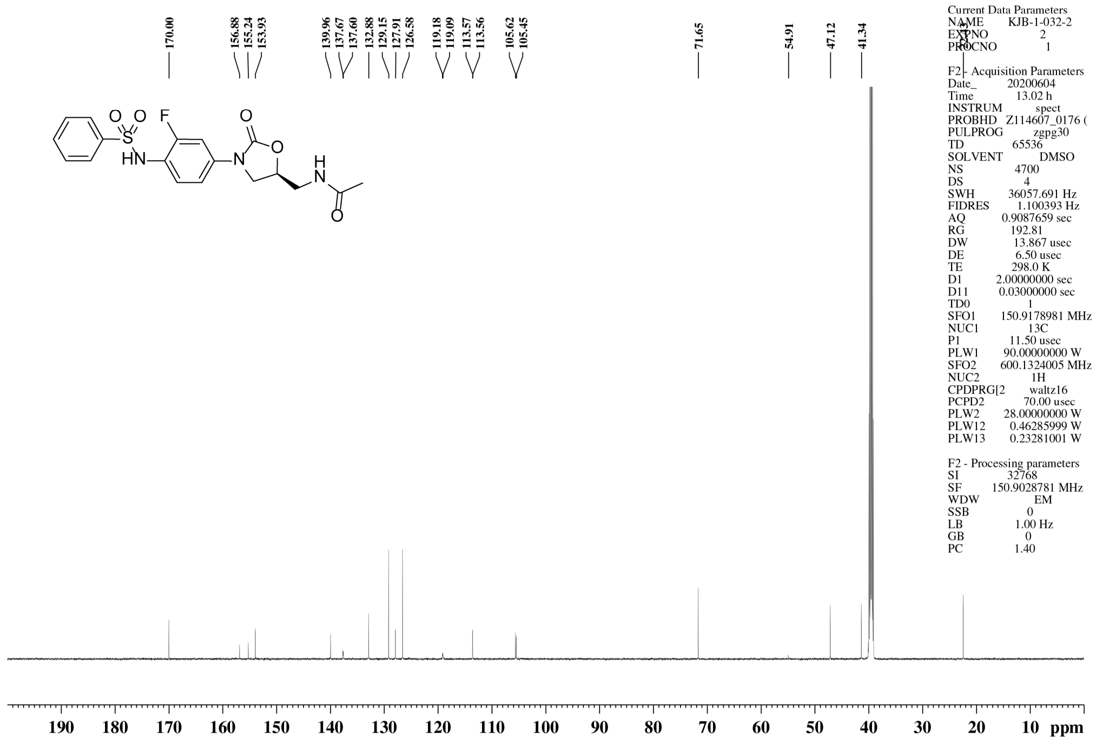

$^{1}$H NMR spectrum of **38**

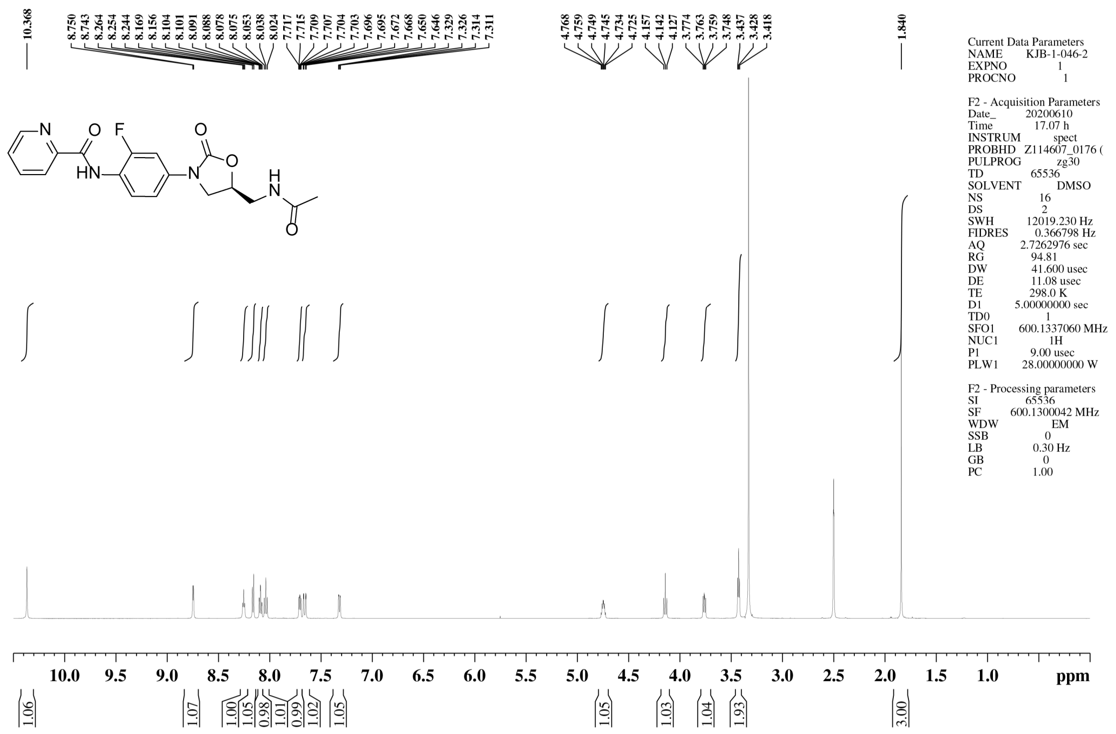


$^{13}$C NMR spectrum of **38**

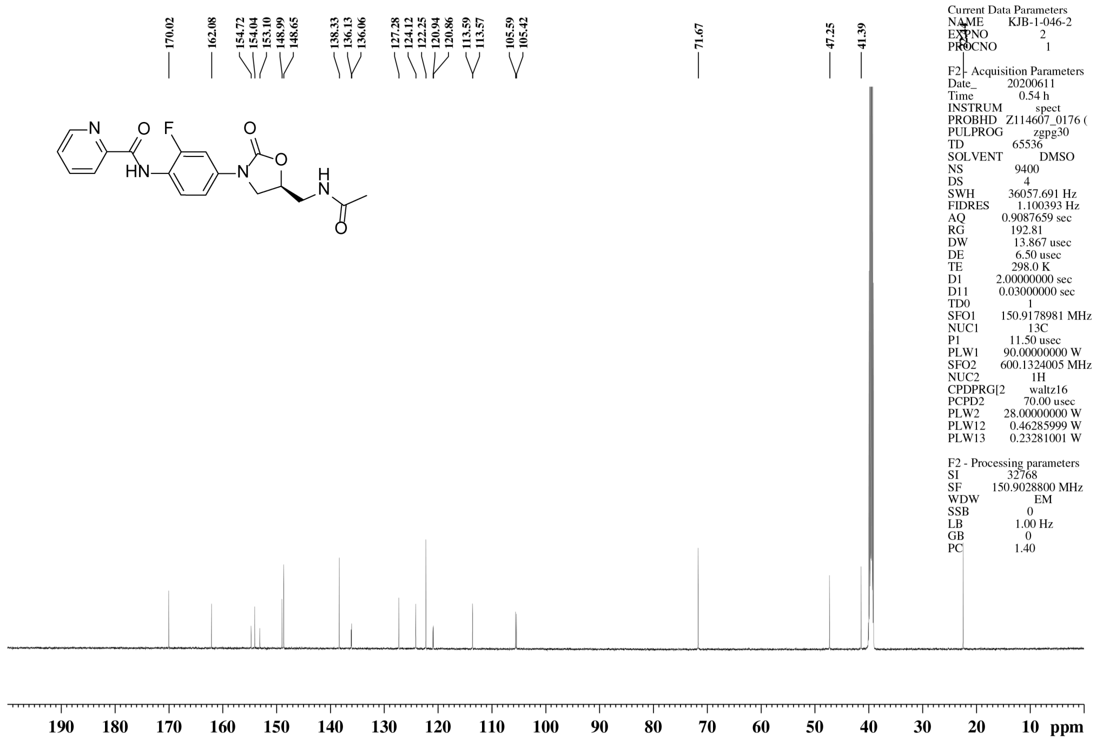

$^{1}$H NMR spectrum of **39**

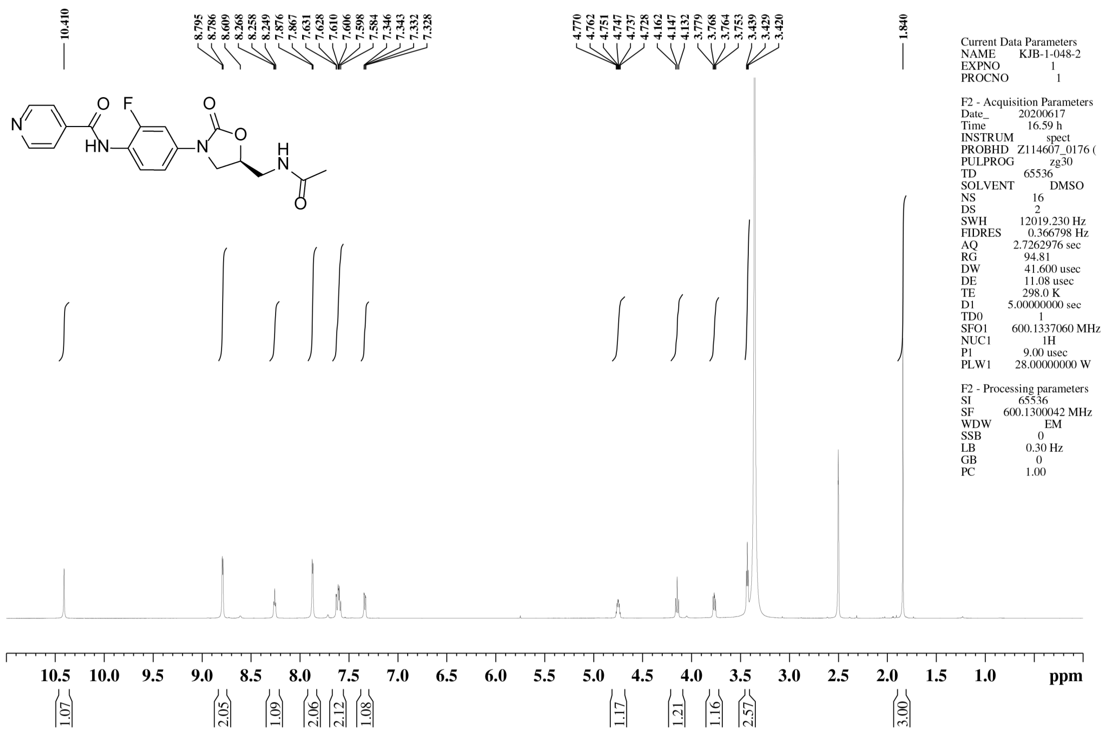


$^{13}$C NMR spectrum of **39**

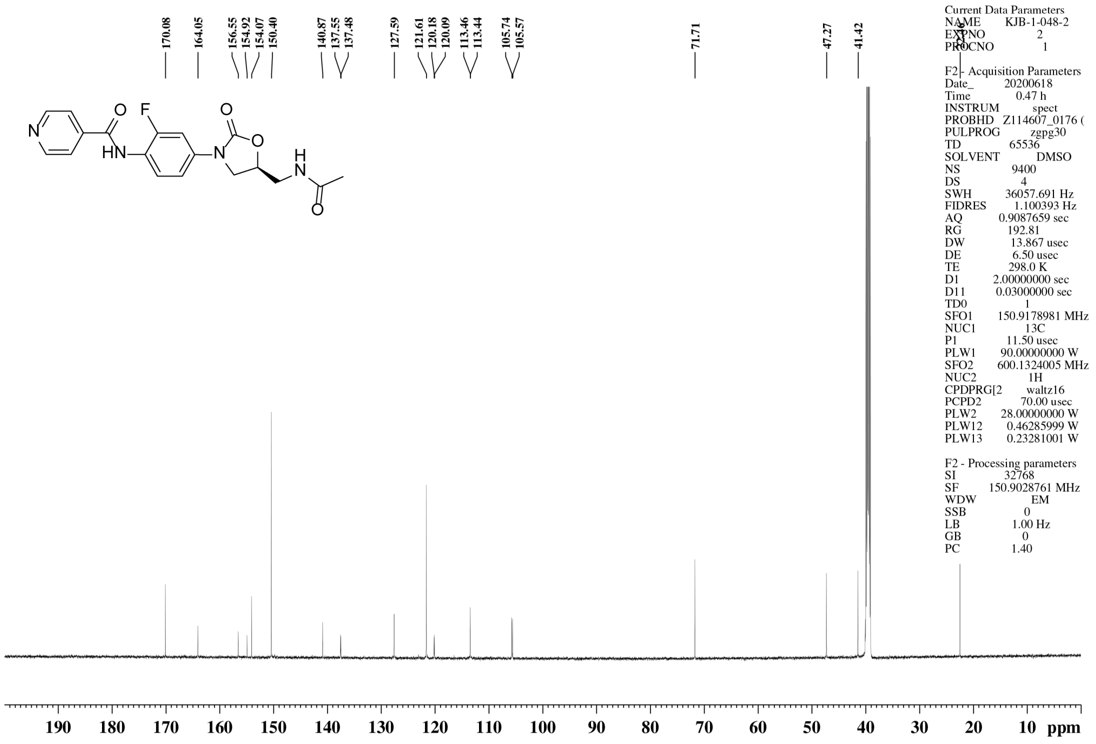

$^{1}$H NMR spectrum of **40**

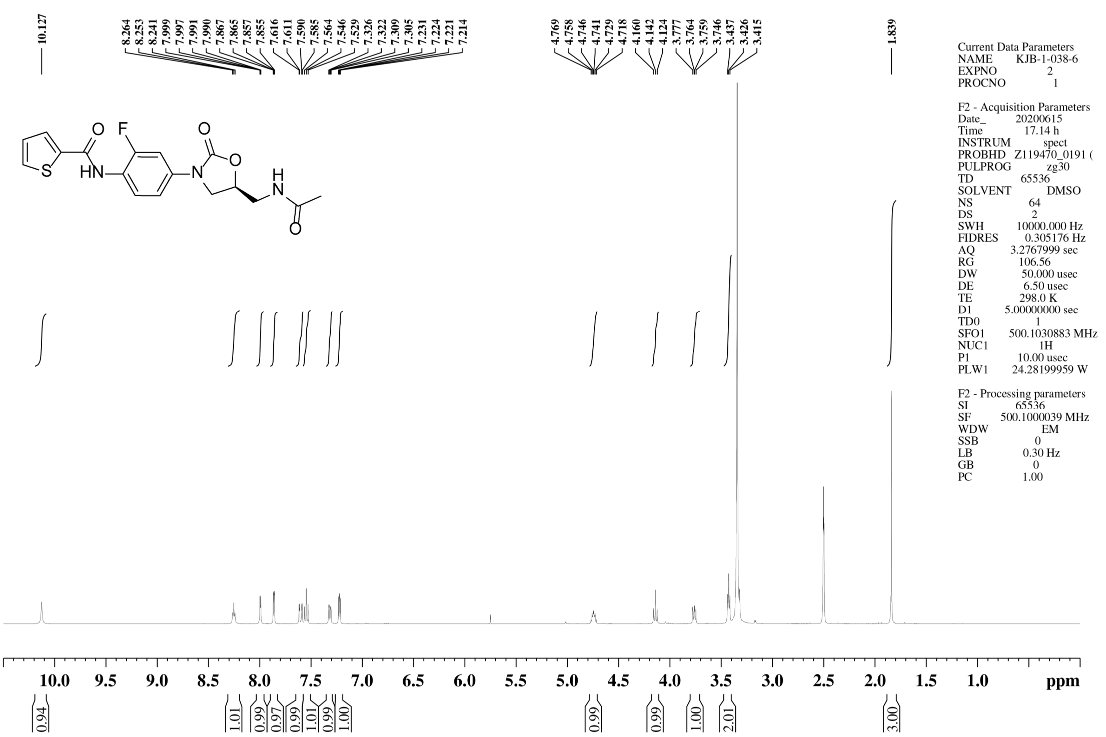


$^{13}$C NMR spectrum of **40**

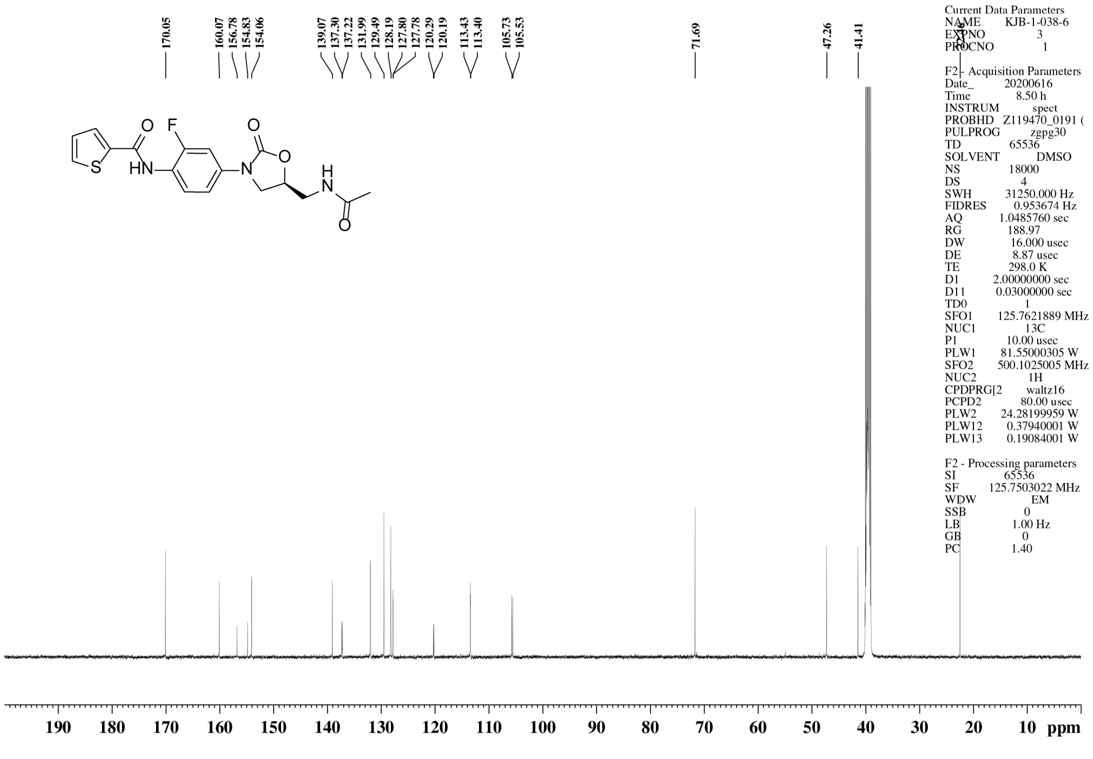

$^{1}$H NMR spectrum of **41**

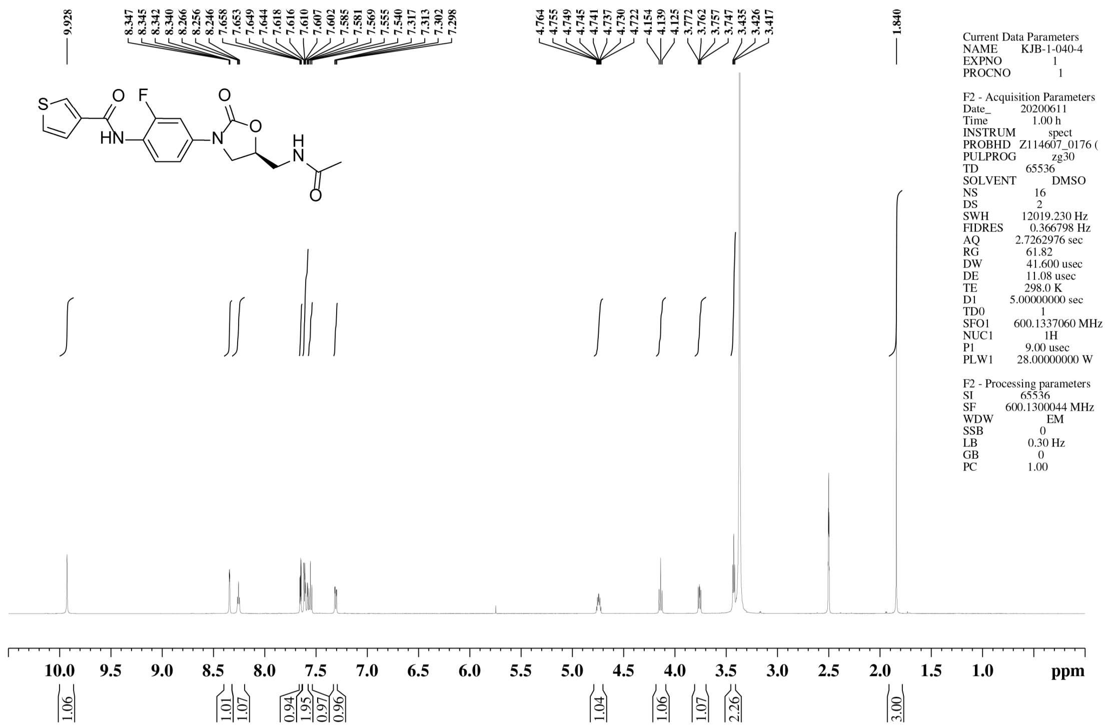


$^{13}$C NMR spectrum of **41**

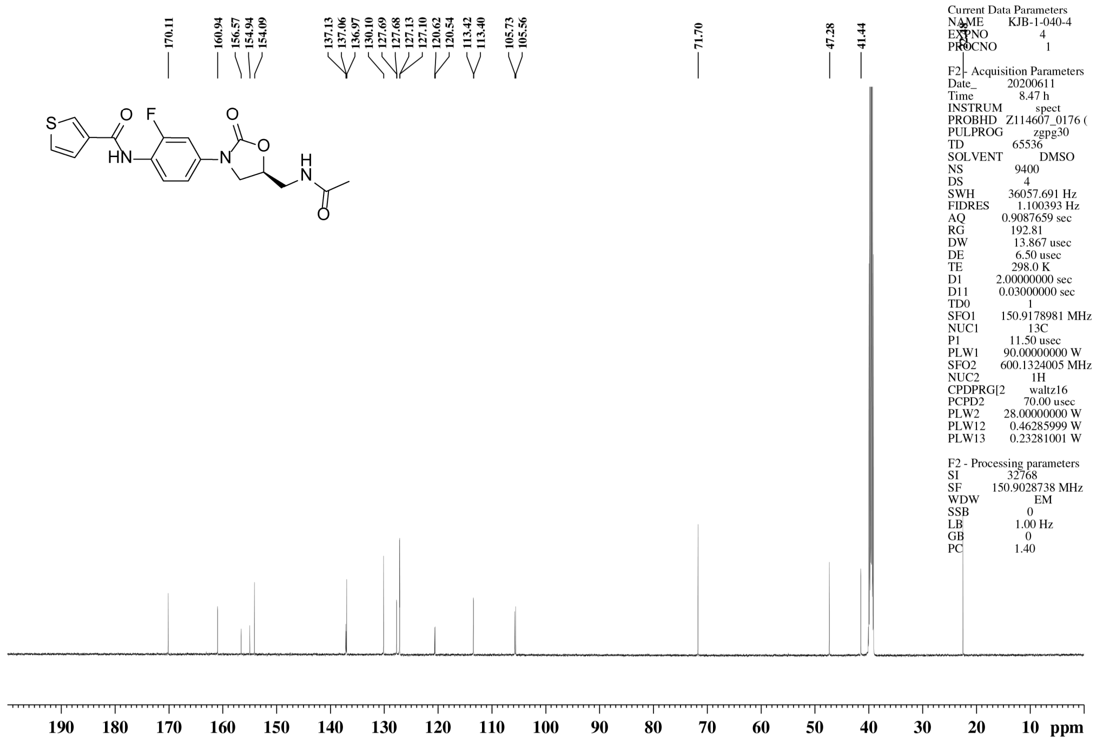

#### $^{1}$H NMR spectrum of **42**

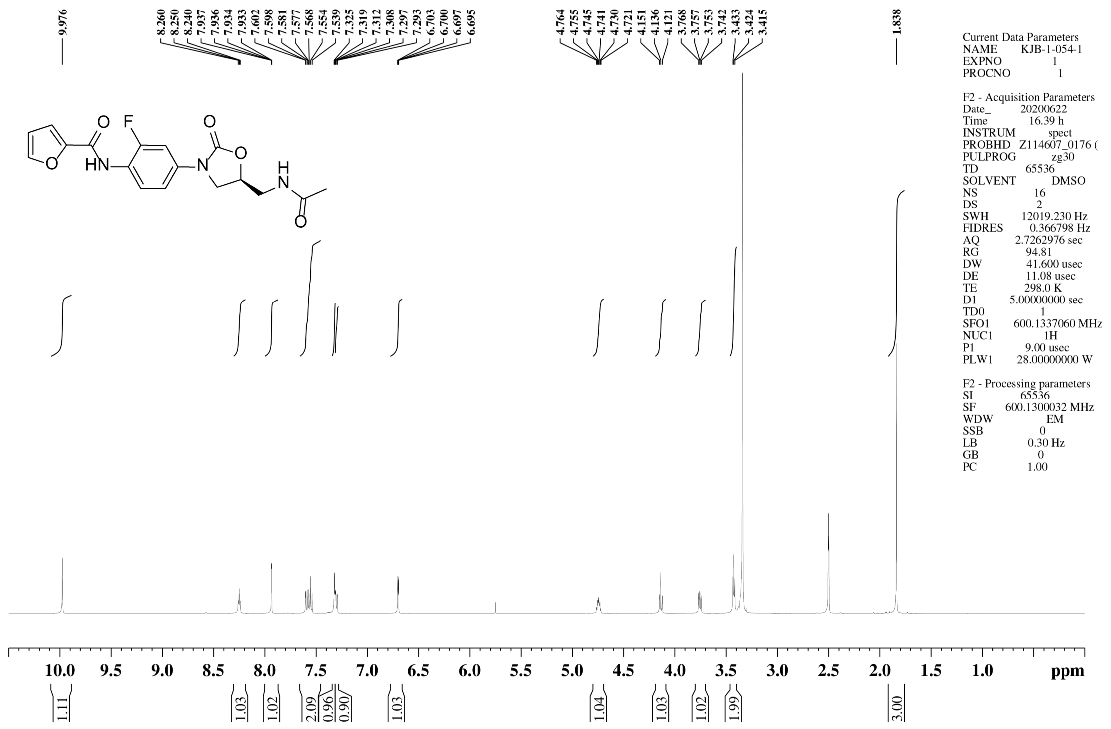


#### $^{13}$C NMR spectrum of **42**

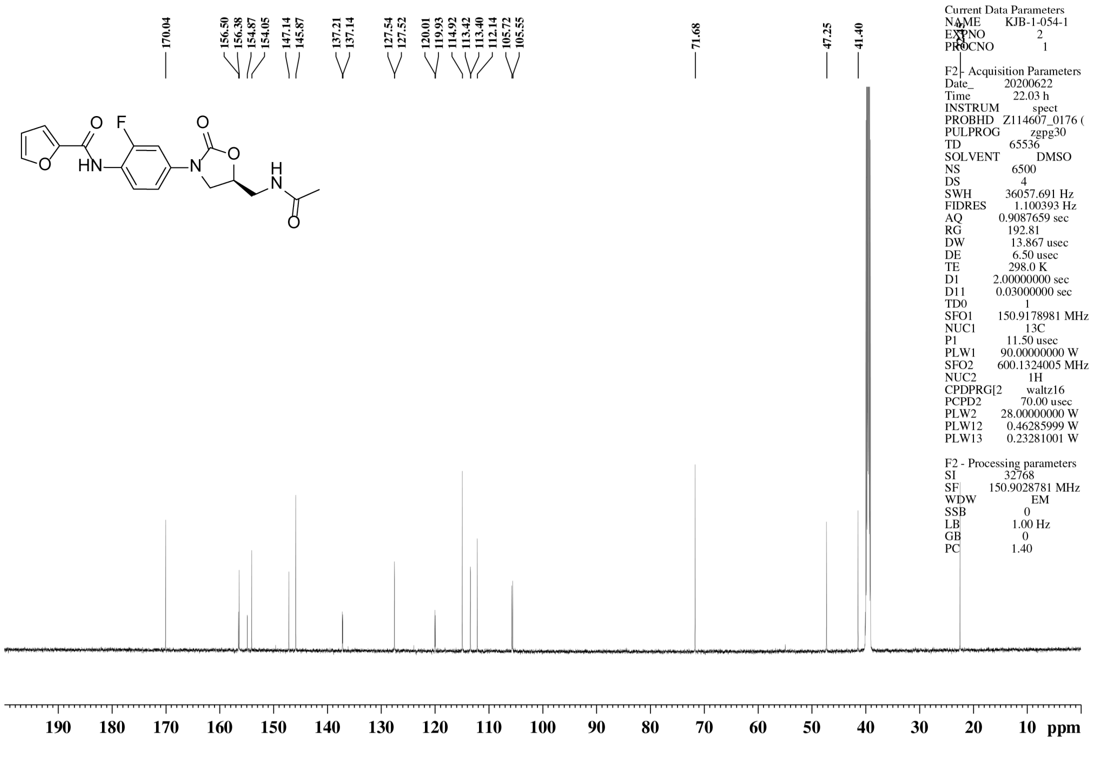

$^{1}$H NMR spectrum of **43**

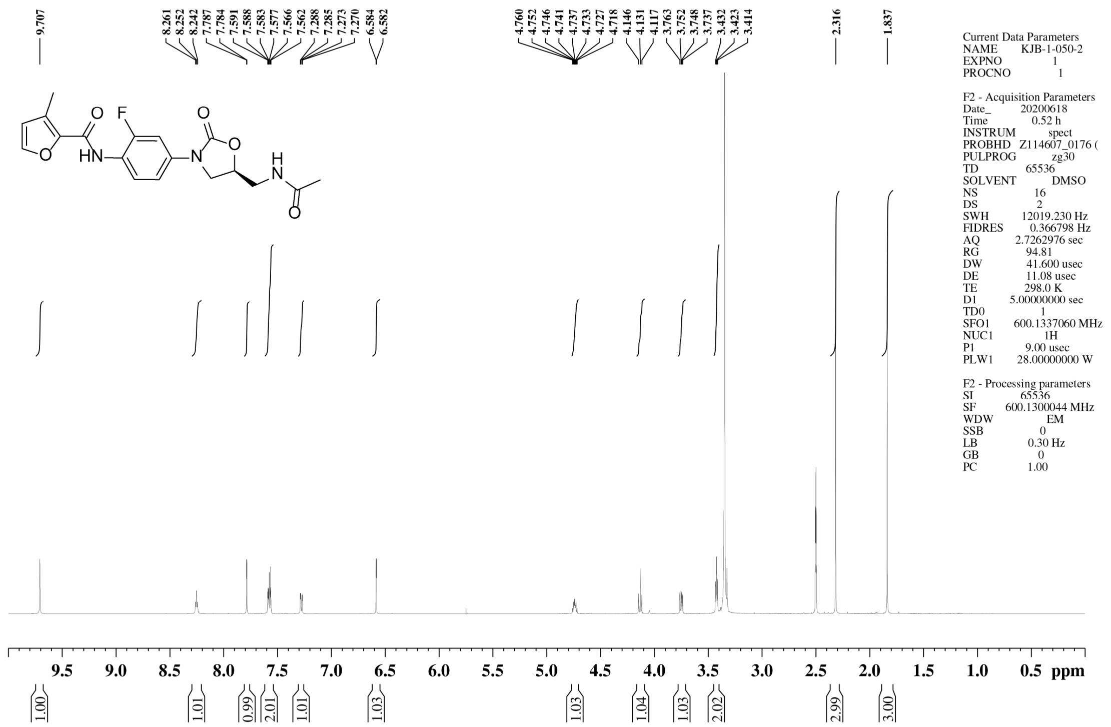


$^{13}$C NMR spectrum of **43**

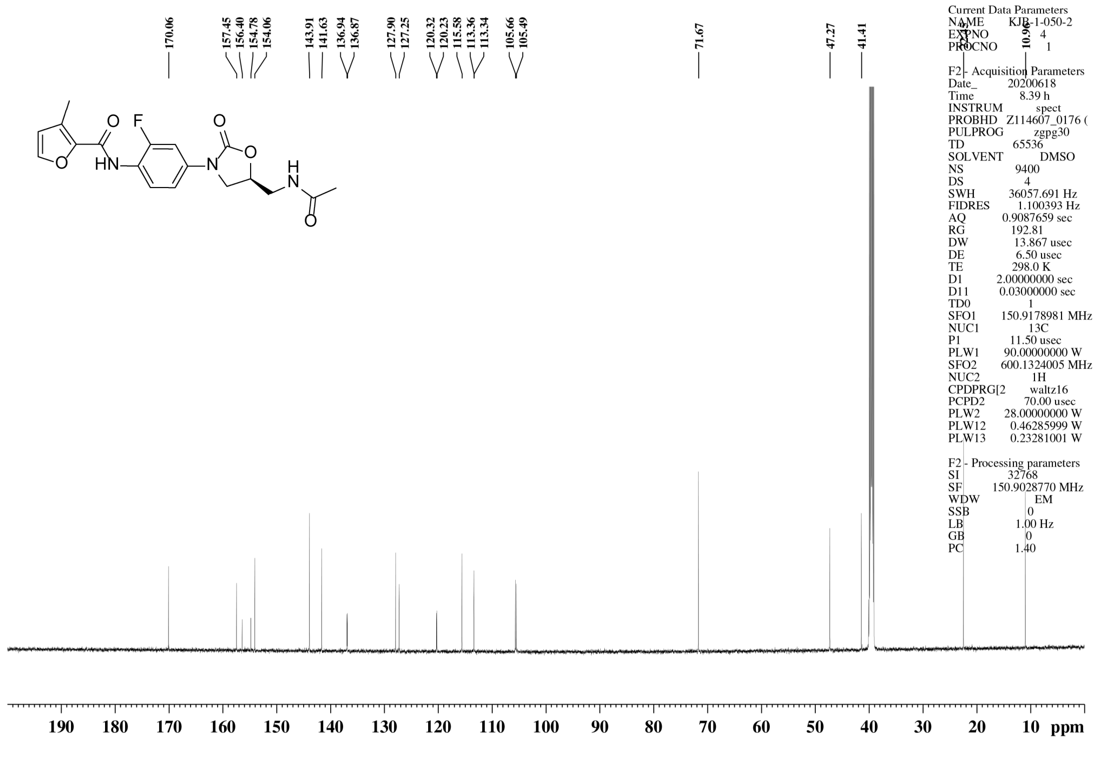

$^{1}$H NMR spectrum of **44**

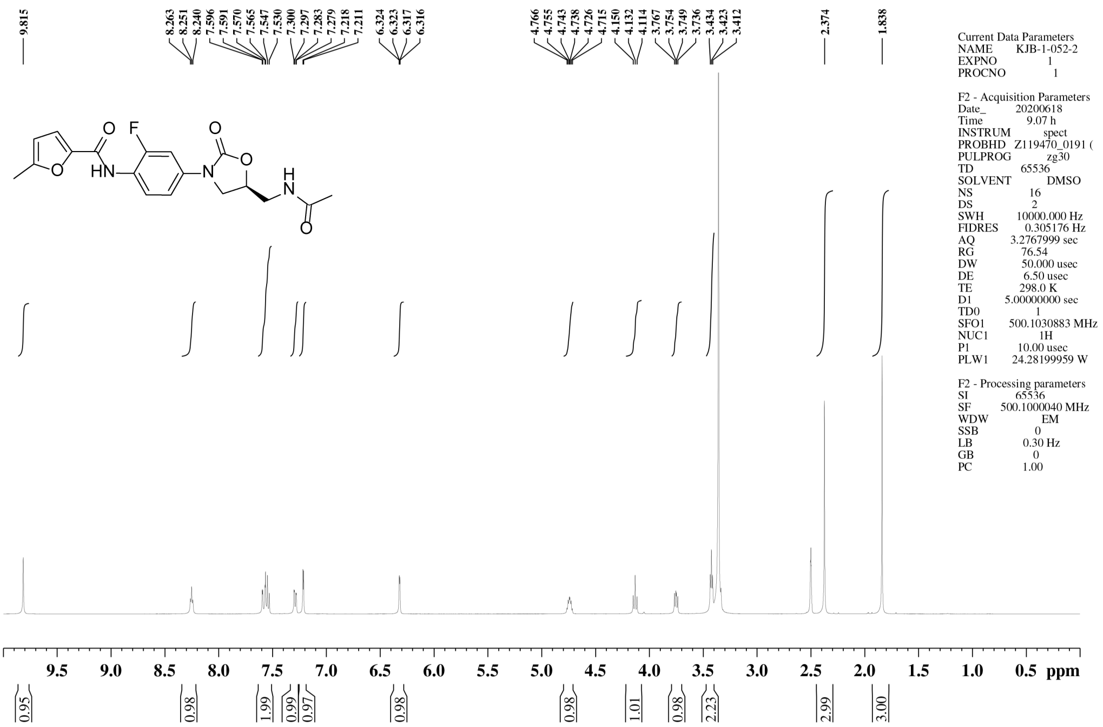


$^{13}$C NMR spectrum of **44**

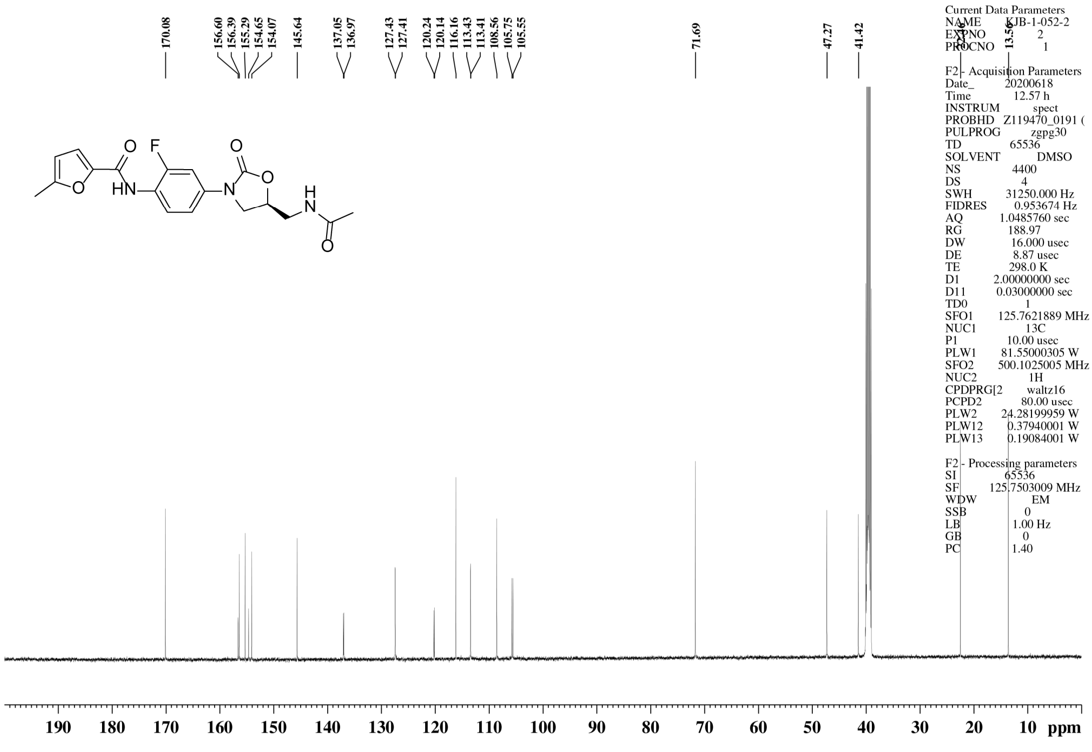

[1]H NMR spectrum of **45**

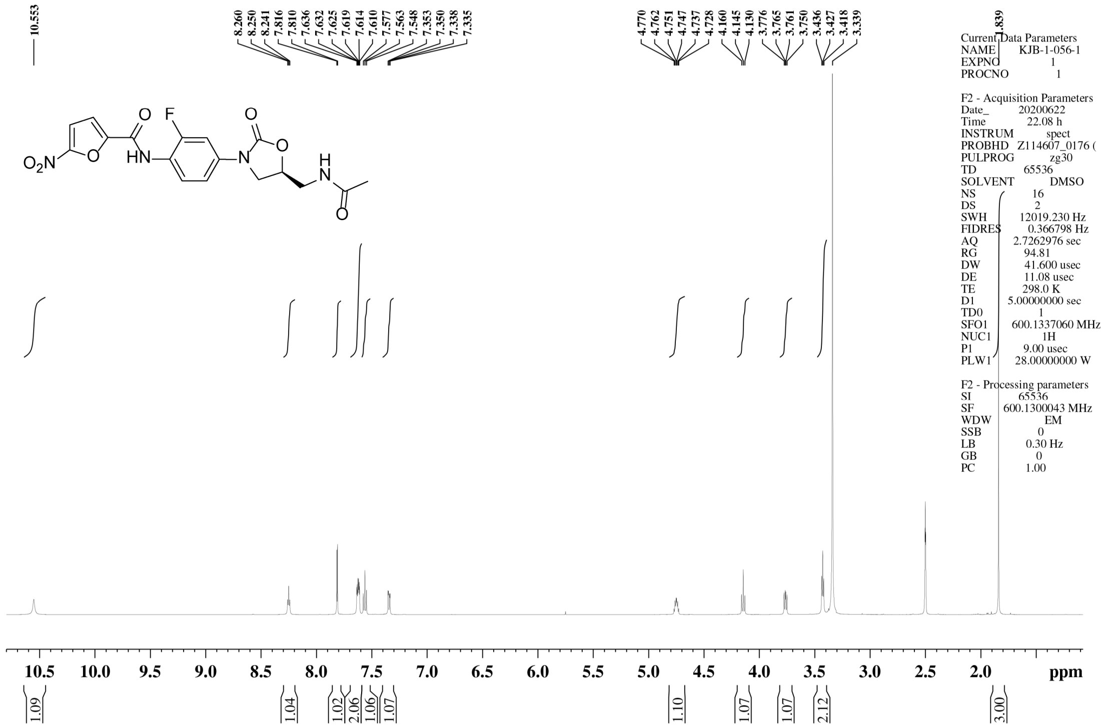

[13]C NMR spectrum of **45**

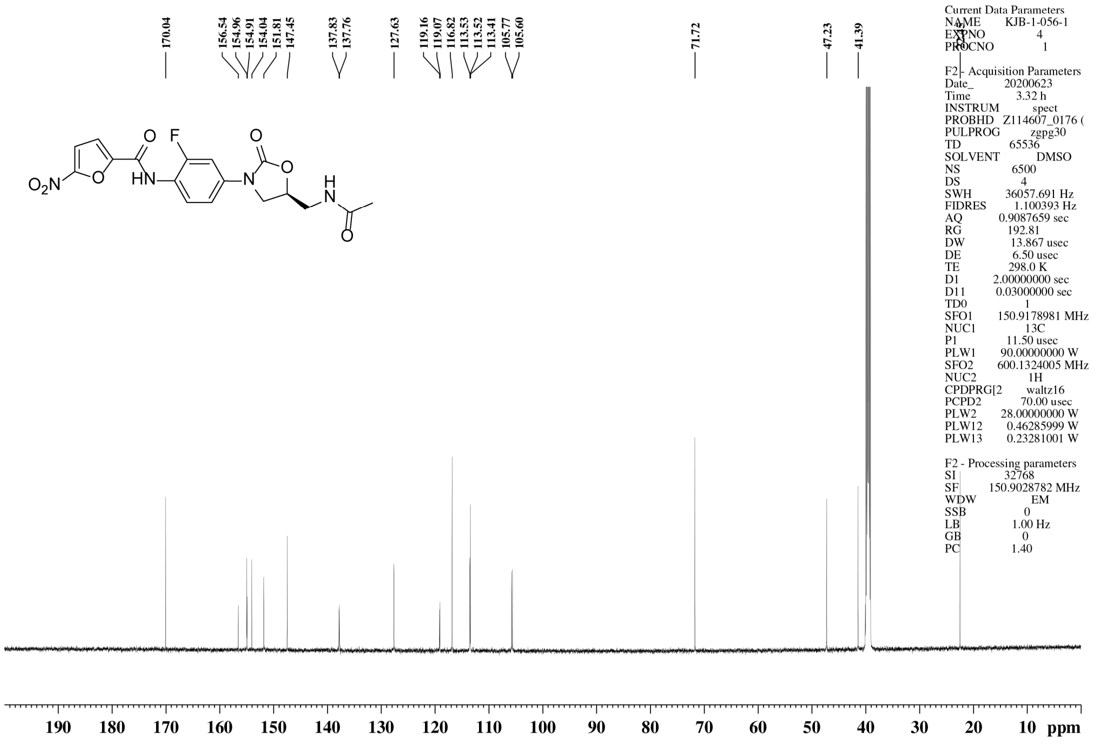

$^{1}$H NMR spectrum of **46**

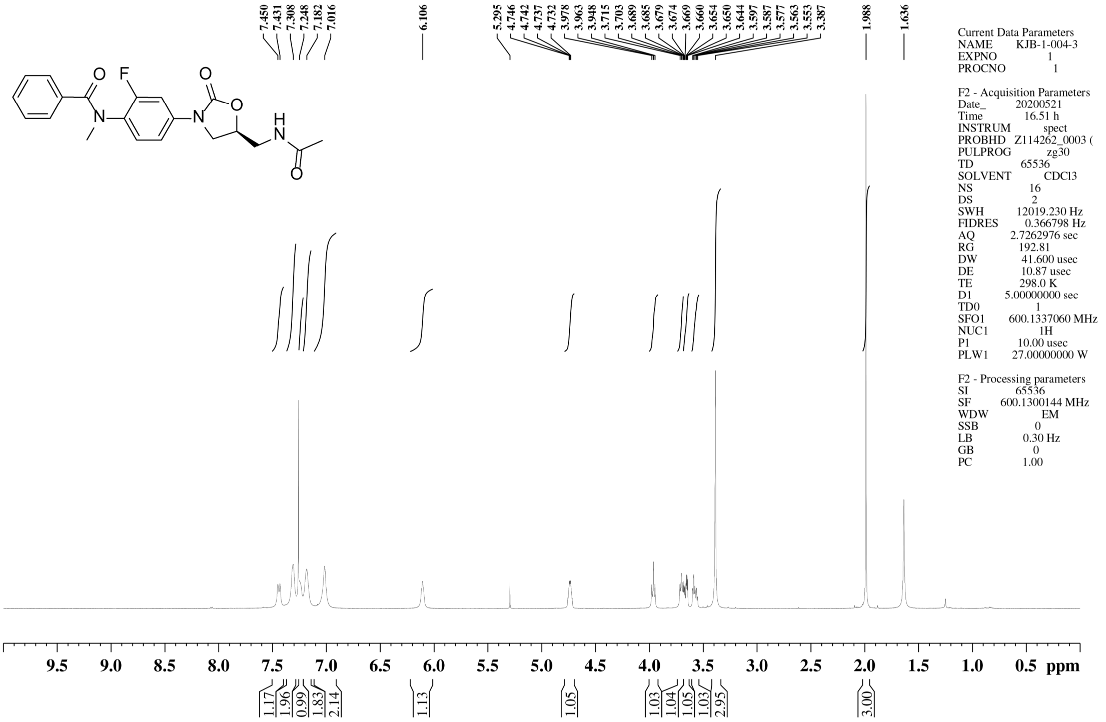


$^{13}$C NMR spectrum of **46**

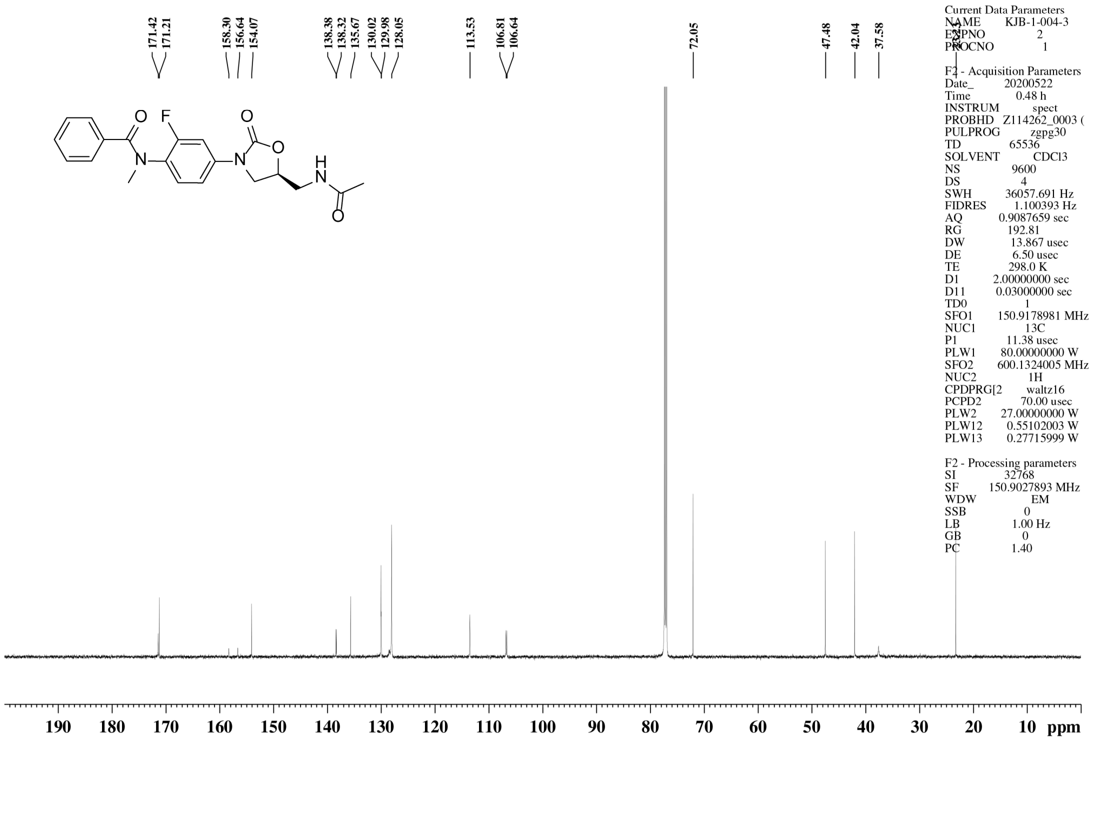

<sup></sup>

[1]H NMR spectrum of **47**

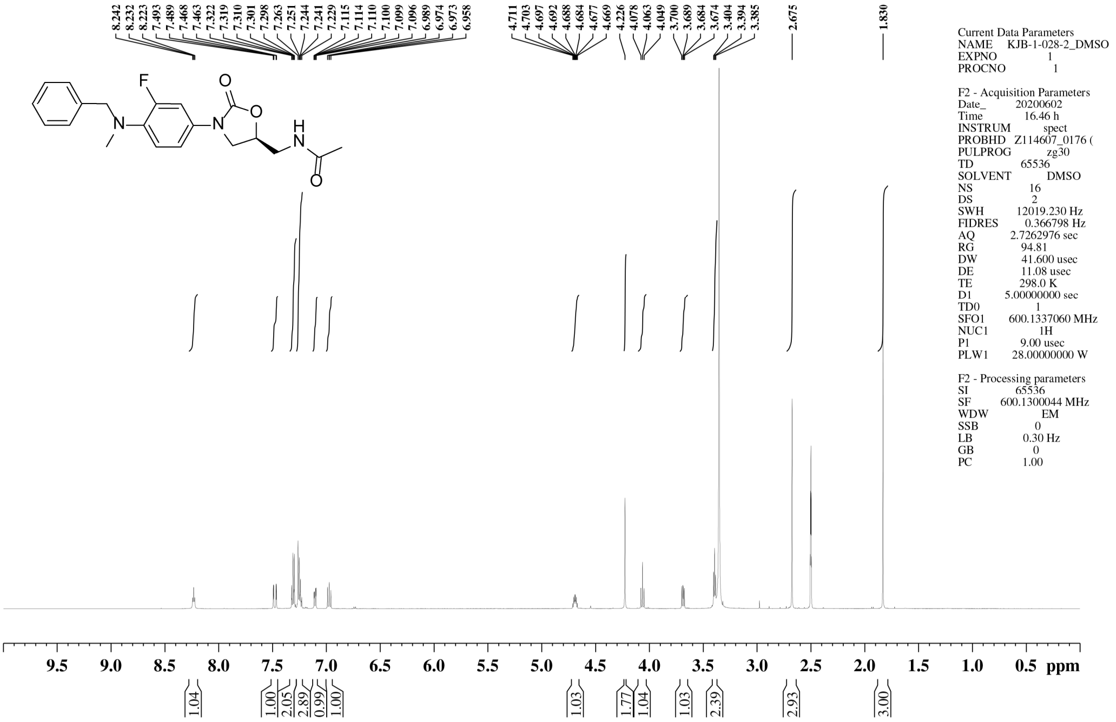

[13]C NMR spectrum of **47**

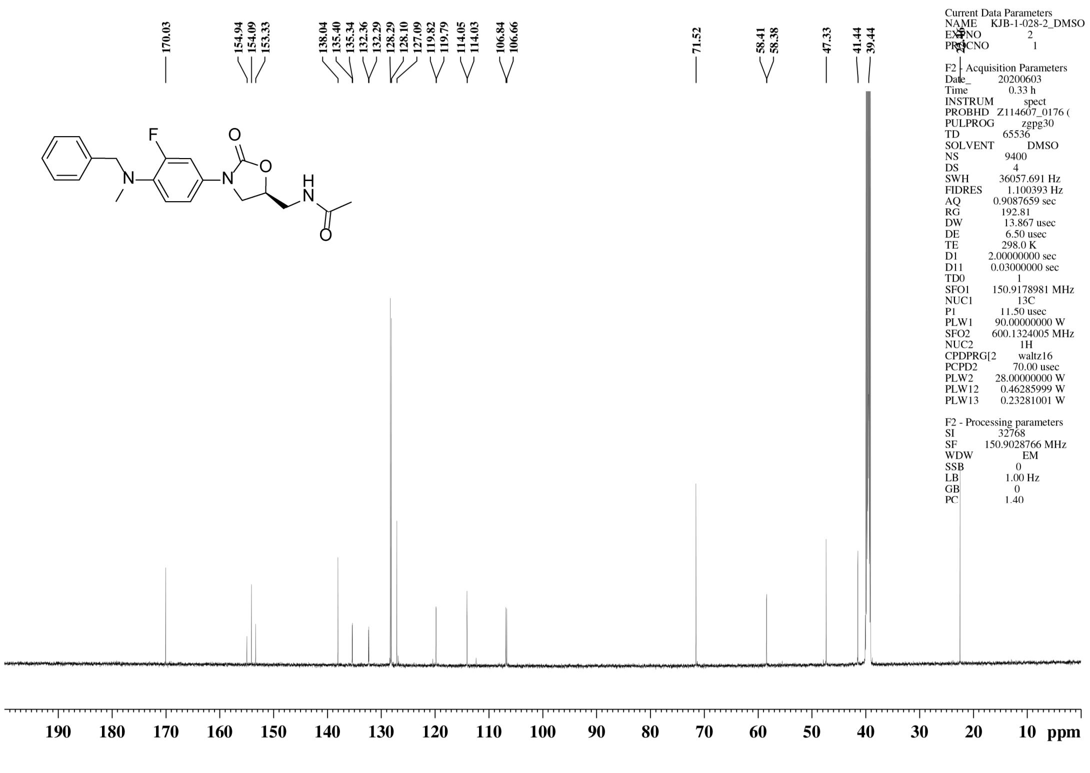

$^{1}$H NMR spectrum of **48**

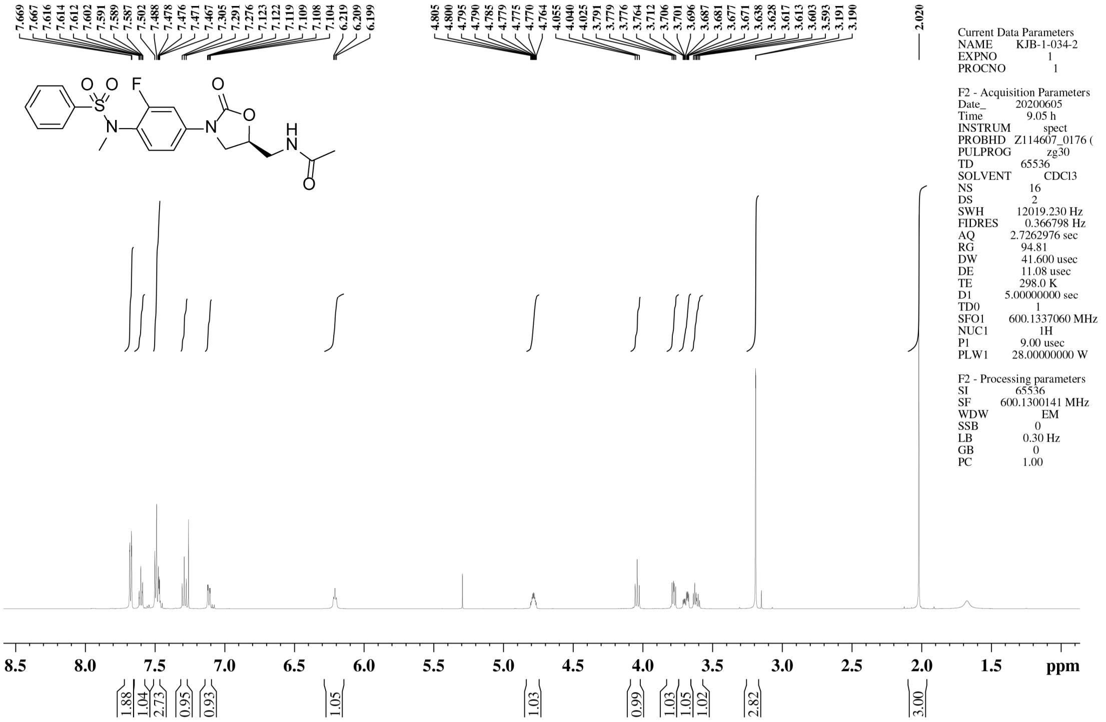


$^{13}$C NMR spectrum of **48**

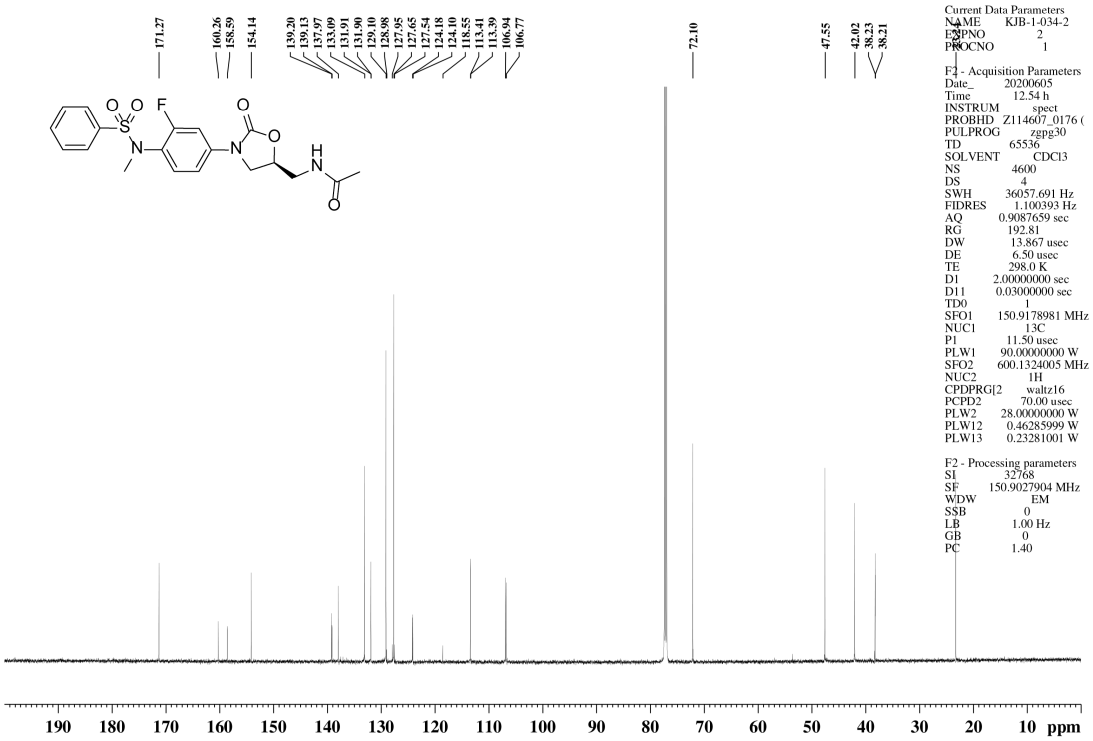

### $^{1}H$ NMR spectrum of **49**

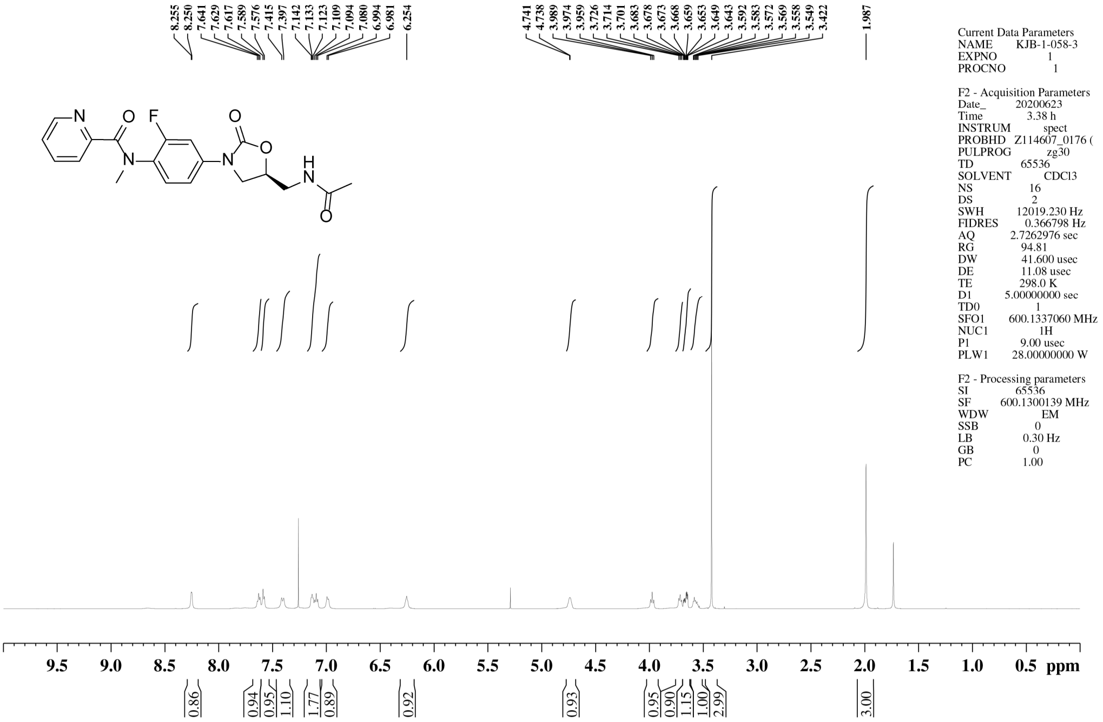


### $^{13}C$ NMR spectrum of **49**

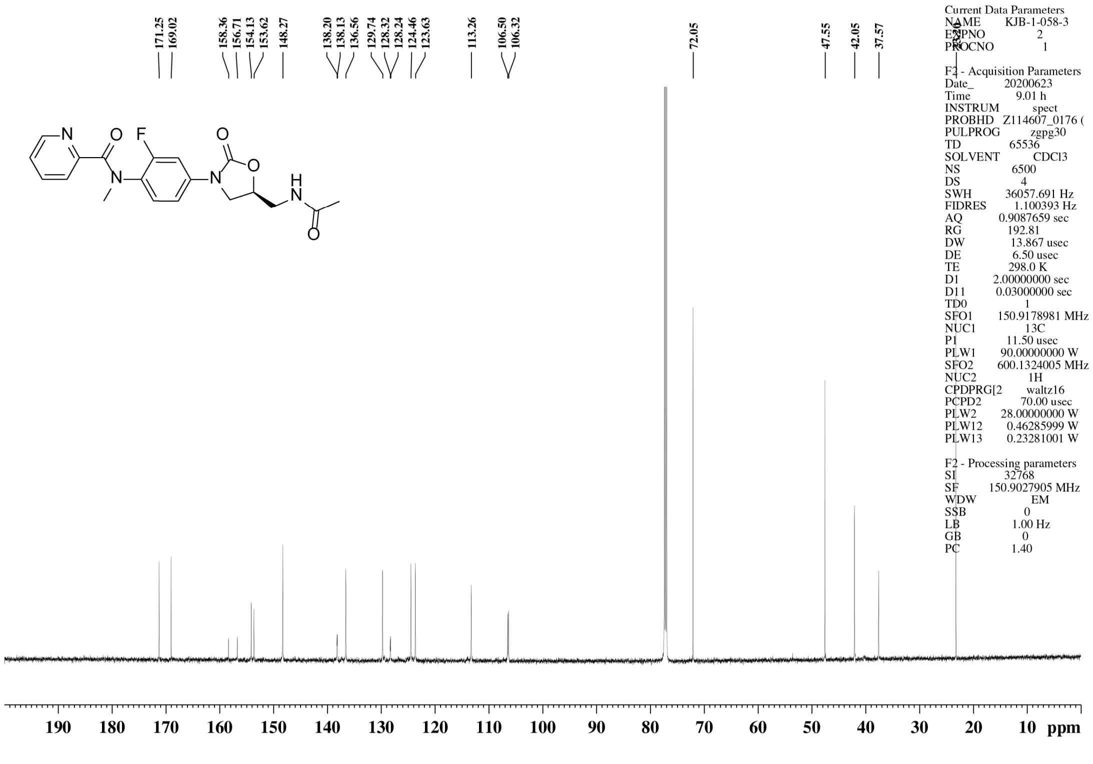

$^{1}$H NMR spectrum of **50**

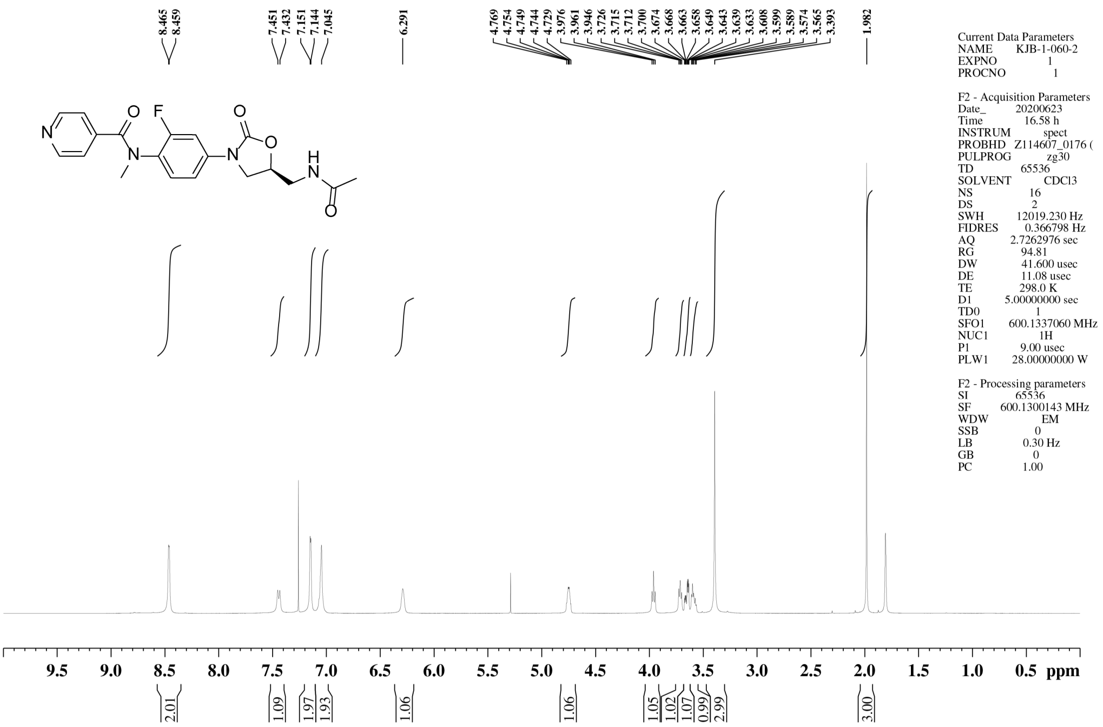


$^{13}$C NMR spectrum of **50**

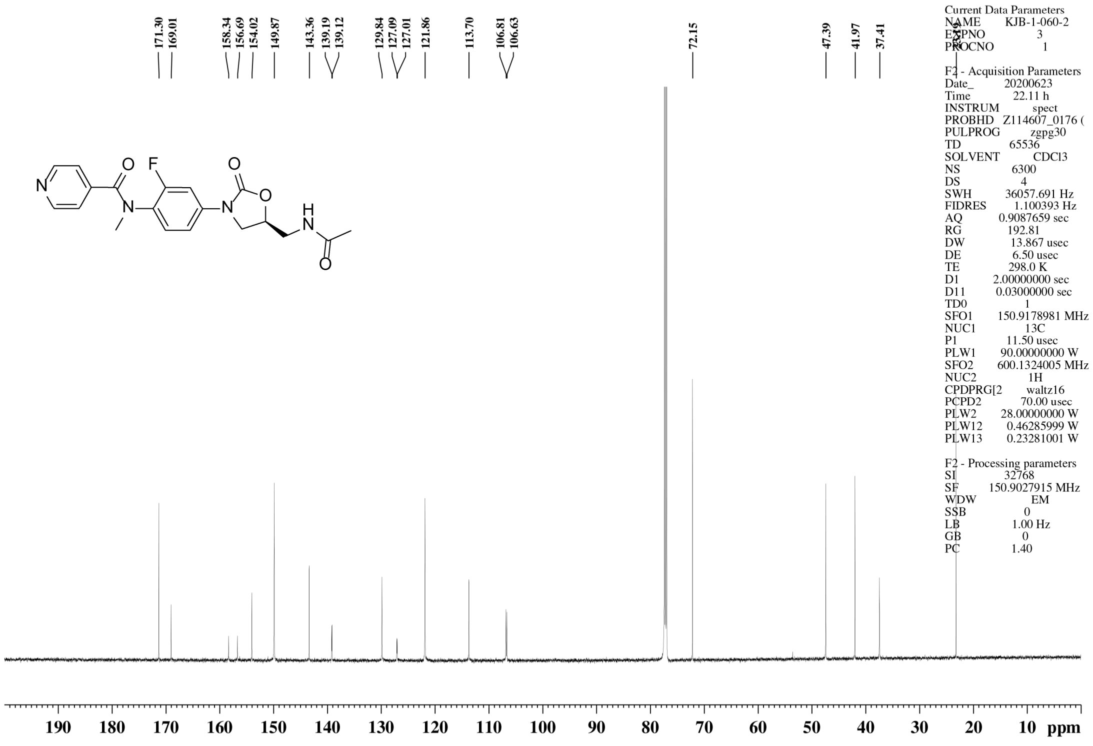

### $^{1}$H NMR spectrum of **51**

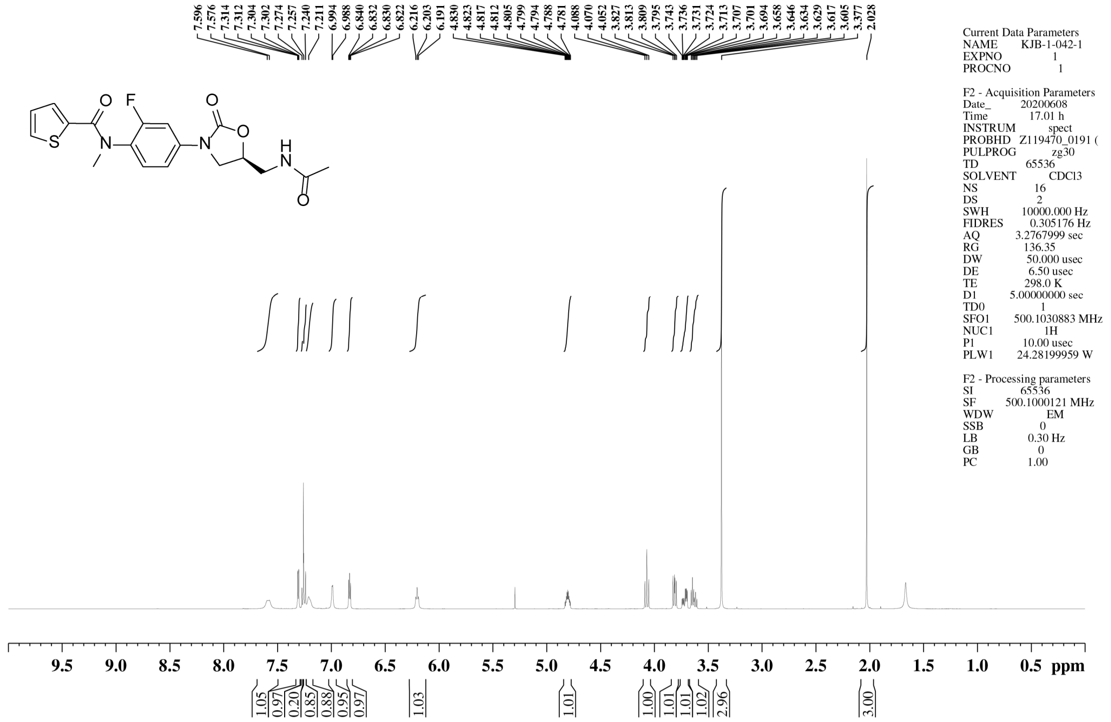


### $^{13}$C NMR spectrum of **51**

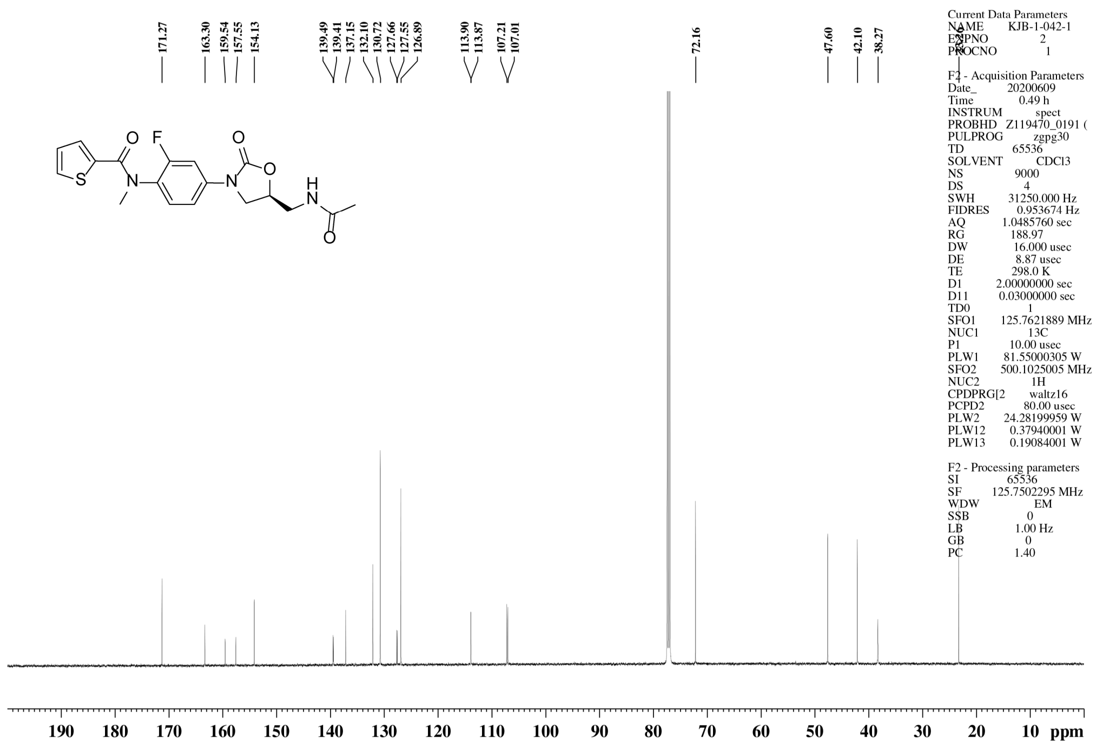

$^{1}$H NMR spectrum of **52**

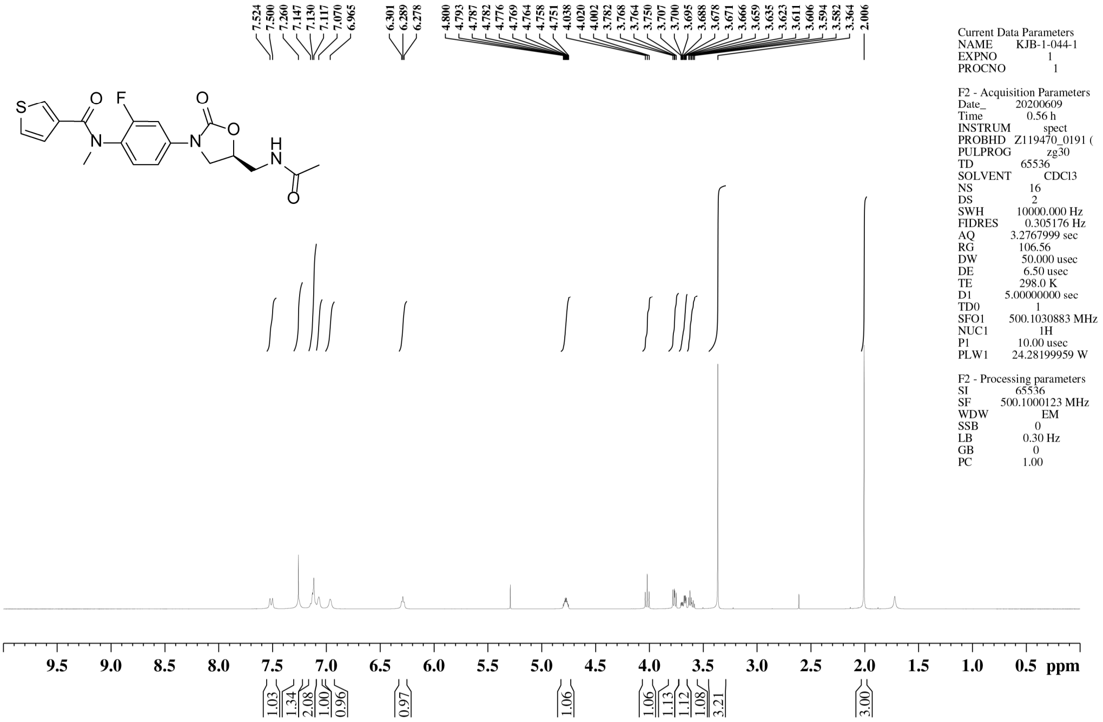


$^{13}$C NMR spectrum of **52**

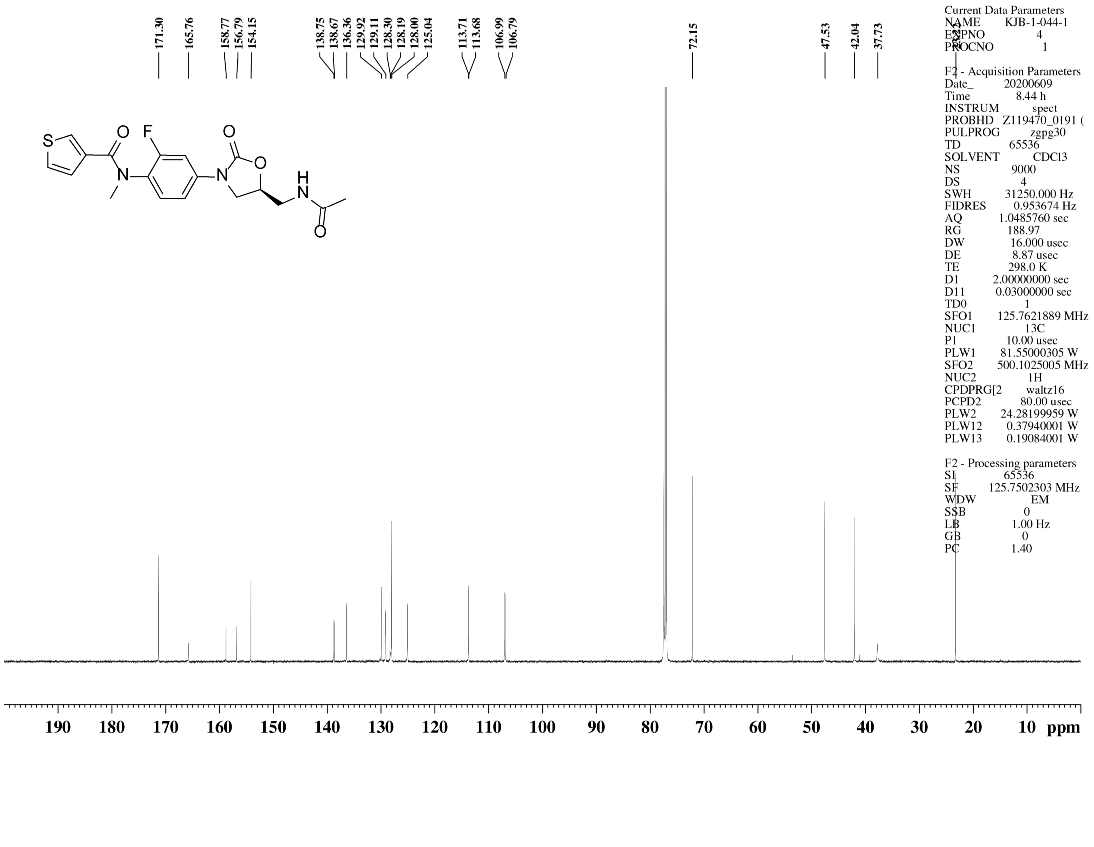

[1]H NMR spectrum of **53**

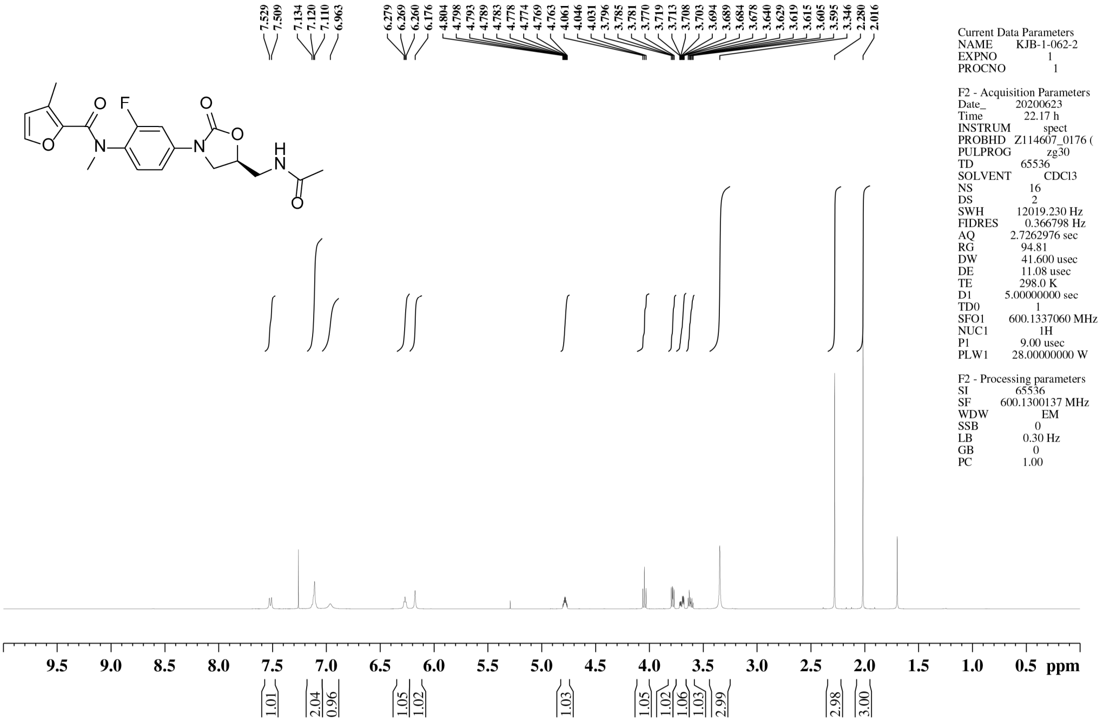


[13]C NMR spectrum of **53**

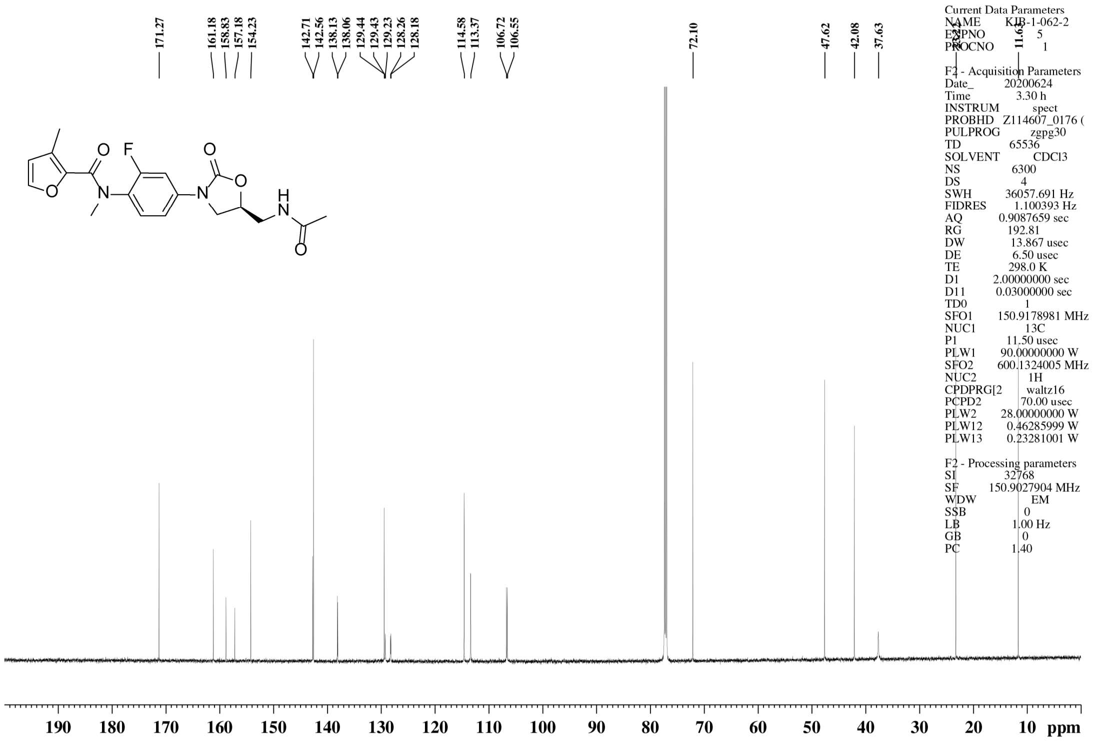

$^{1}$H NMR spectrum of **54**

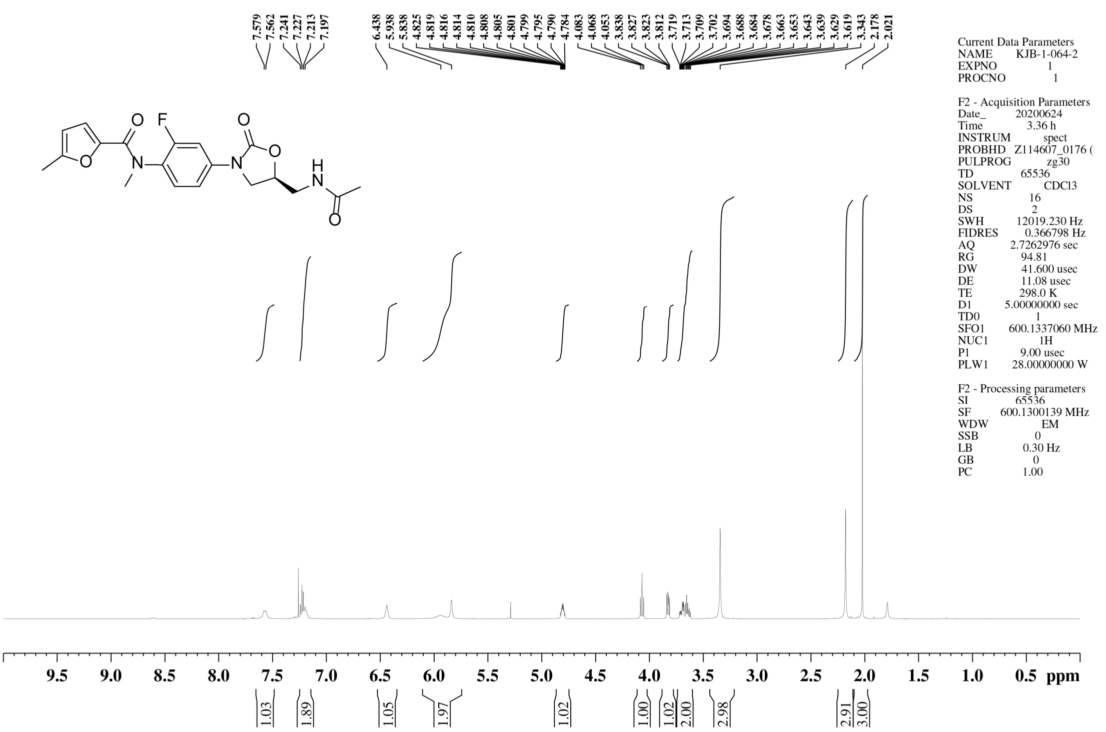


$^{13}$C NMR spectrum of **54**

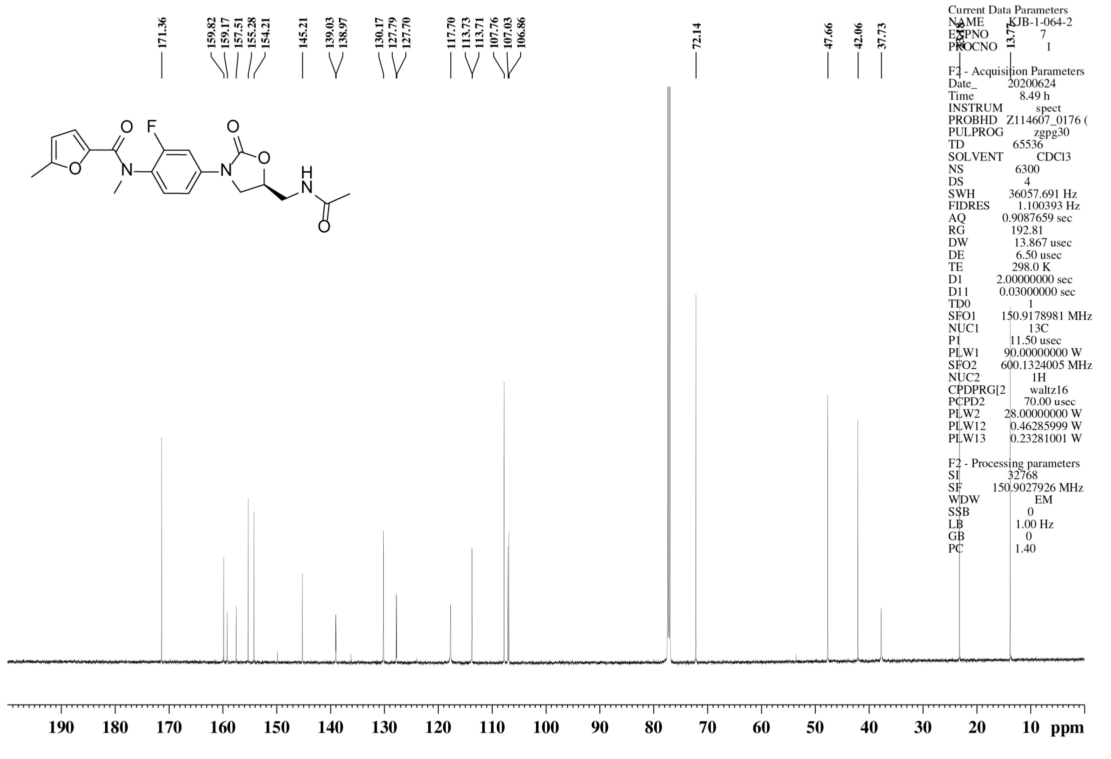

### $^{1}$H NMR spectrum of **g**

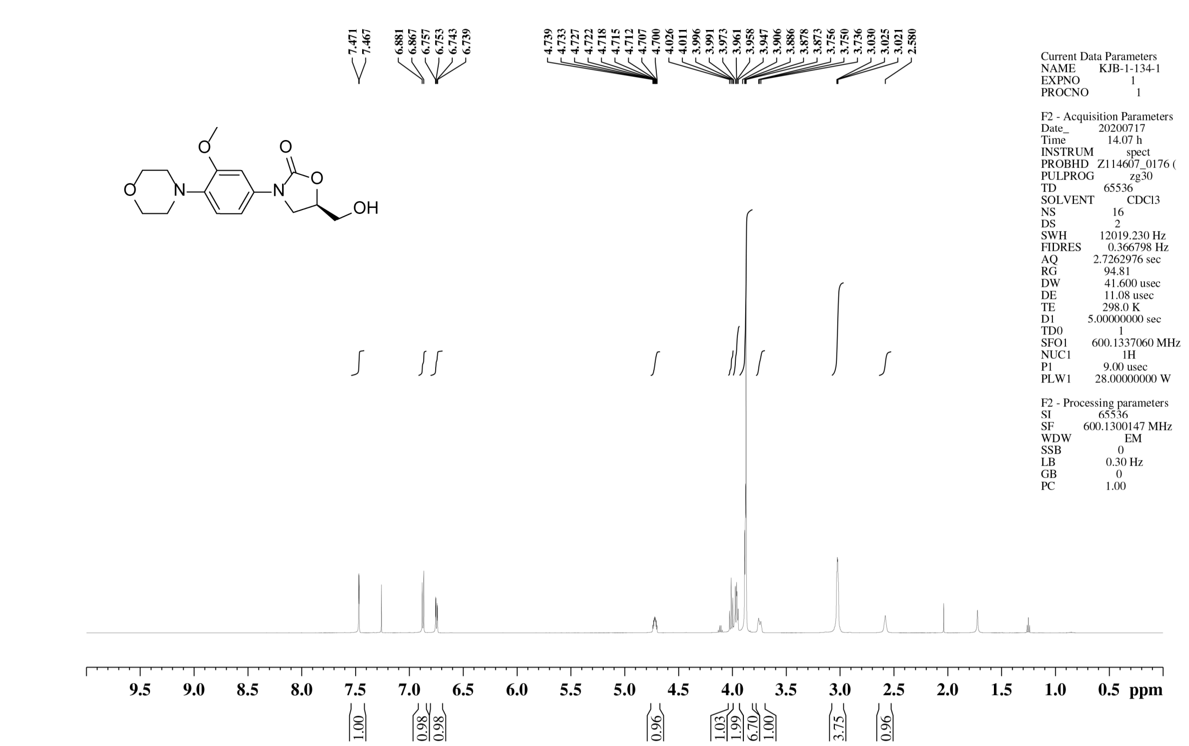


### $^{13}$C NMR spectrum of **g**

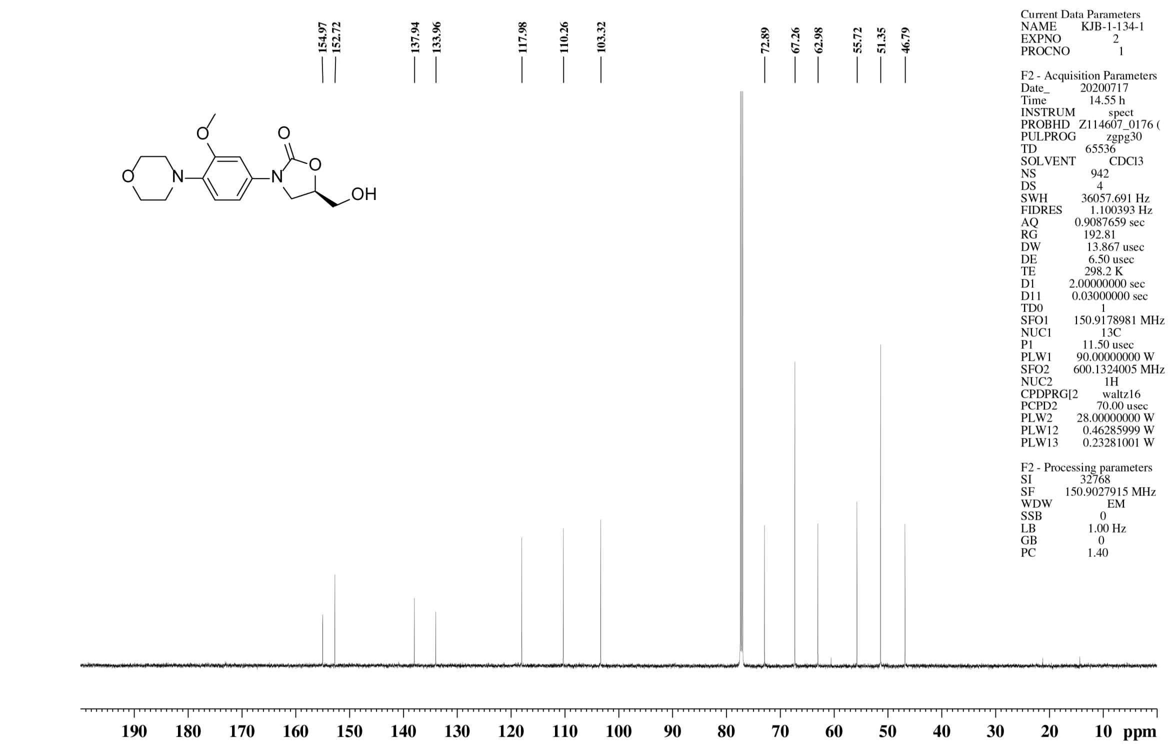

$^1$H NMR spectrum of **i**

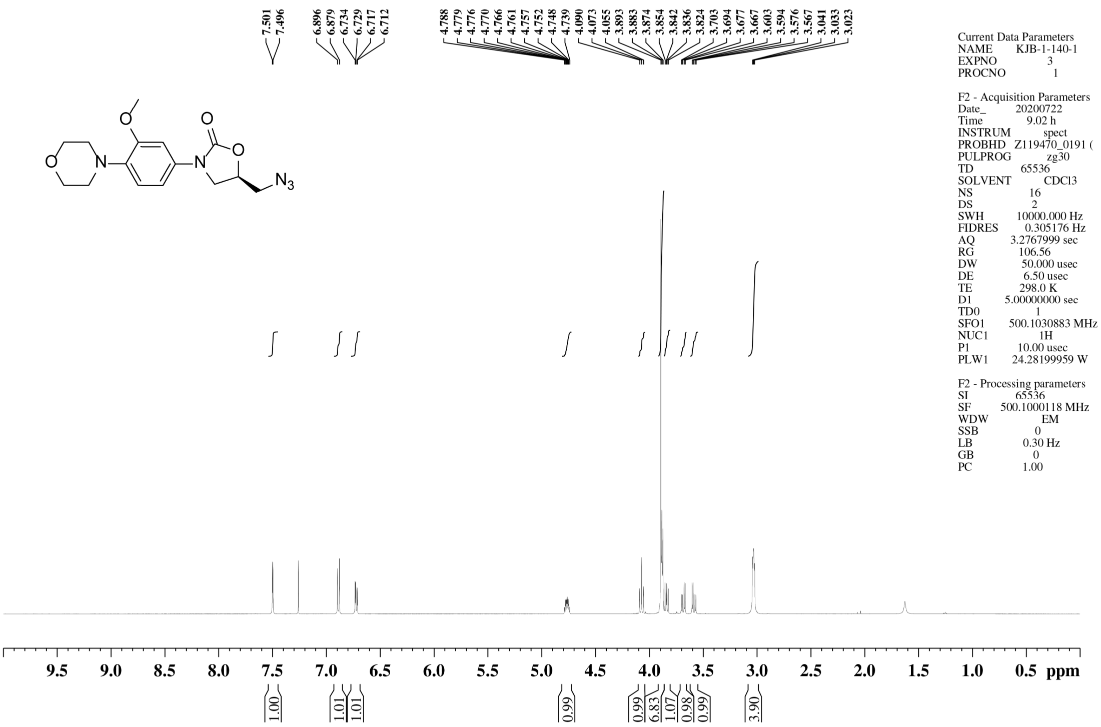

$^{13}$C NMR spectrum of **i**

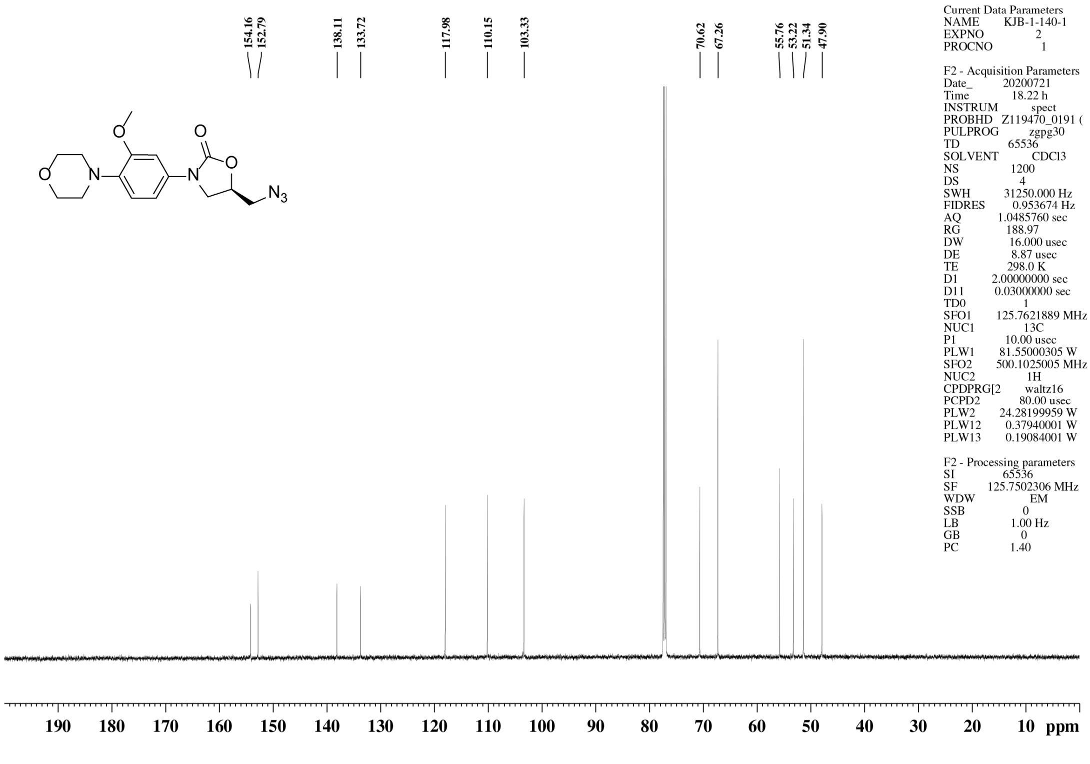

#### $^1$H NMR spectrum of **j**

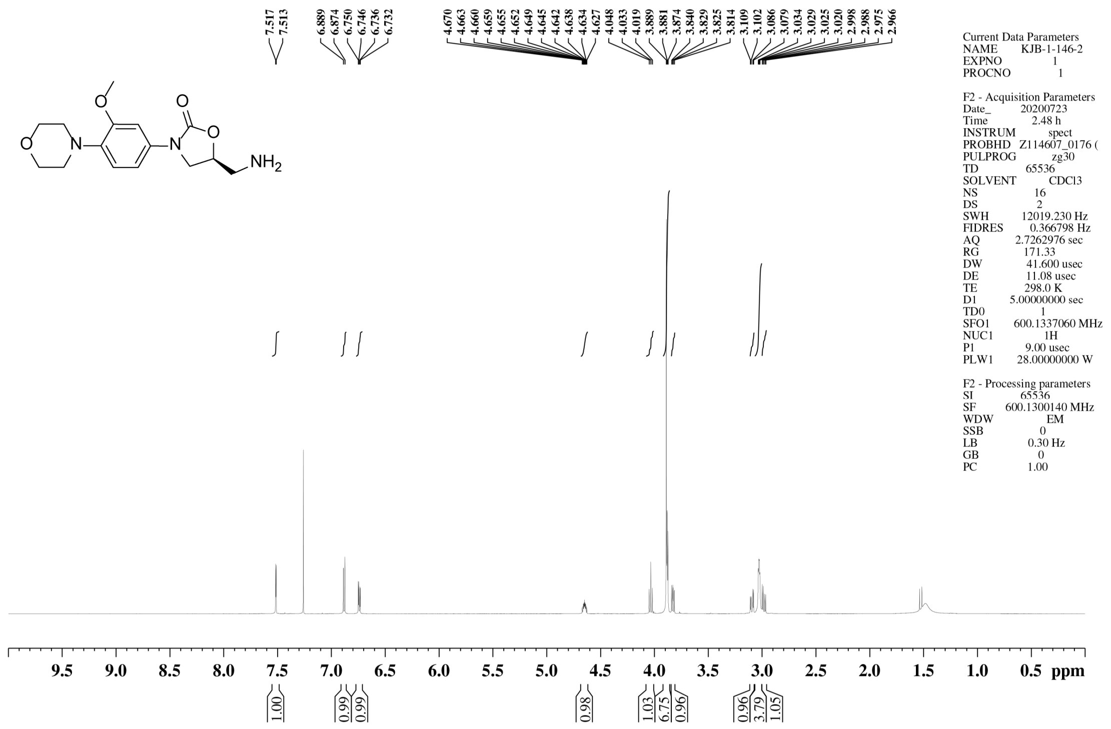


#### $^{13}$C NMR spectrum of **j**

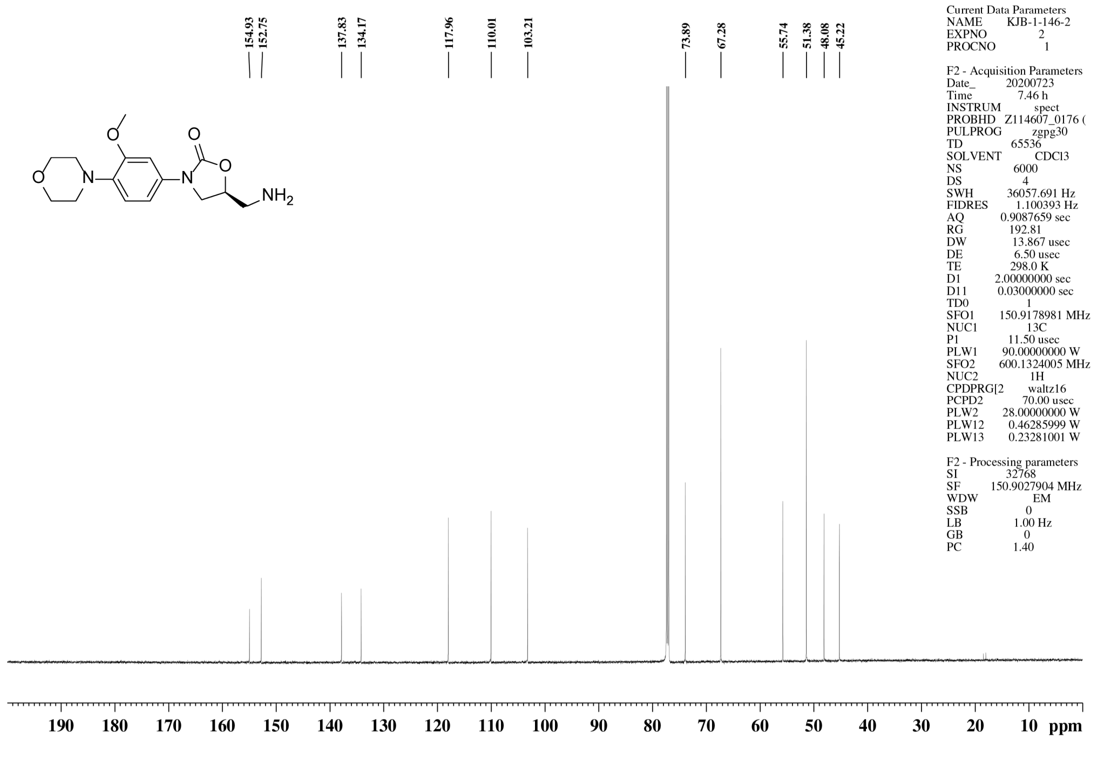

#### $^{1}$H NMR spectrum of **56**

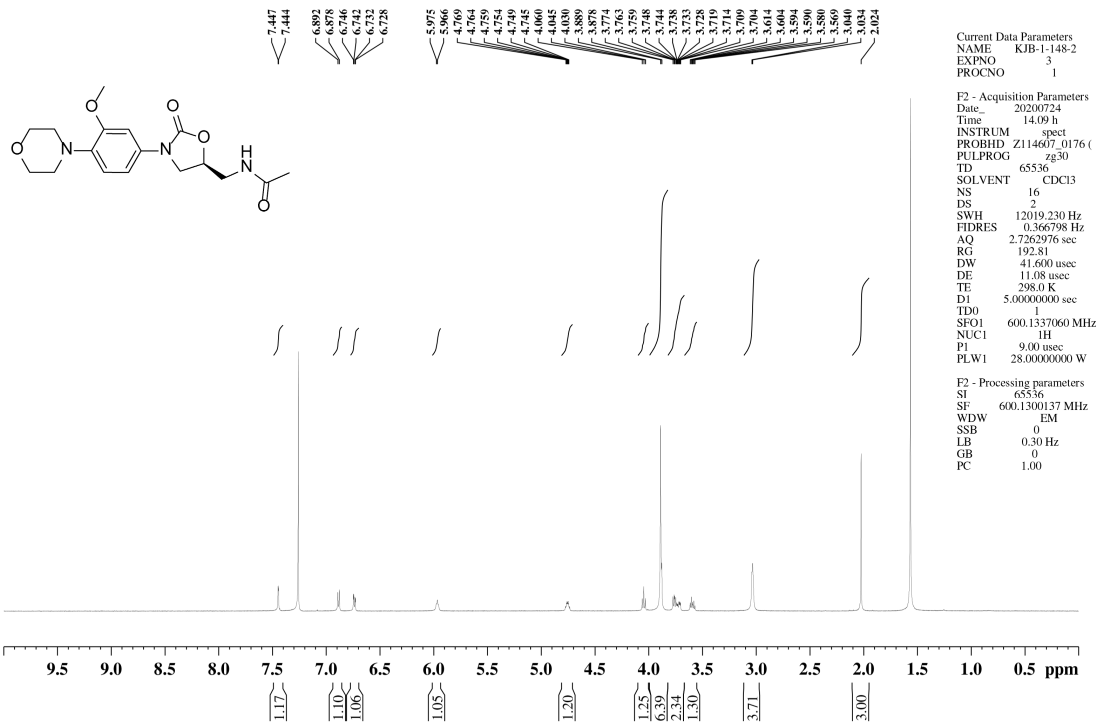


#### $^{13}$C NMR spectrum of **56**

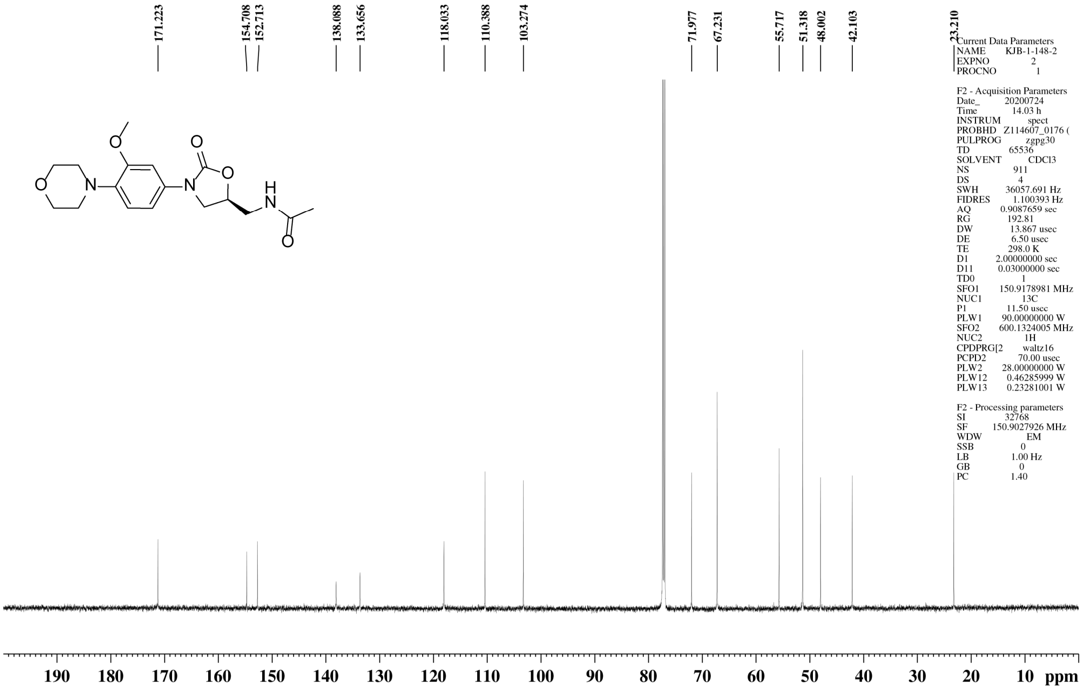

### $^{1}$H NMR spectrum of **57**

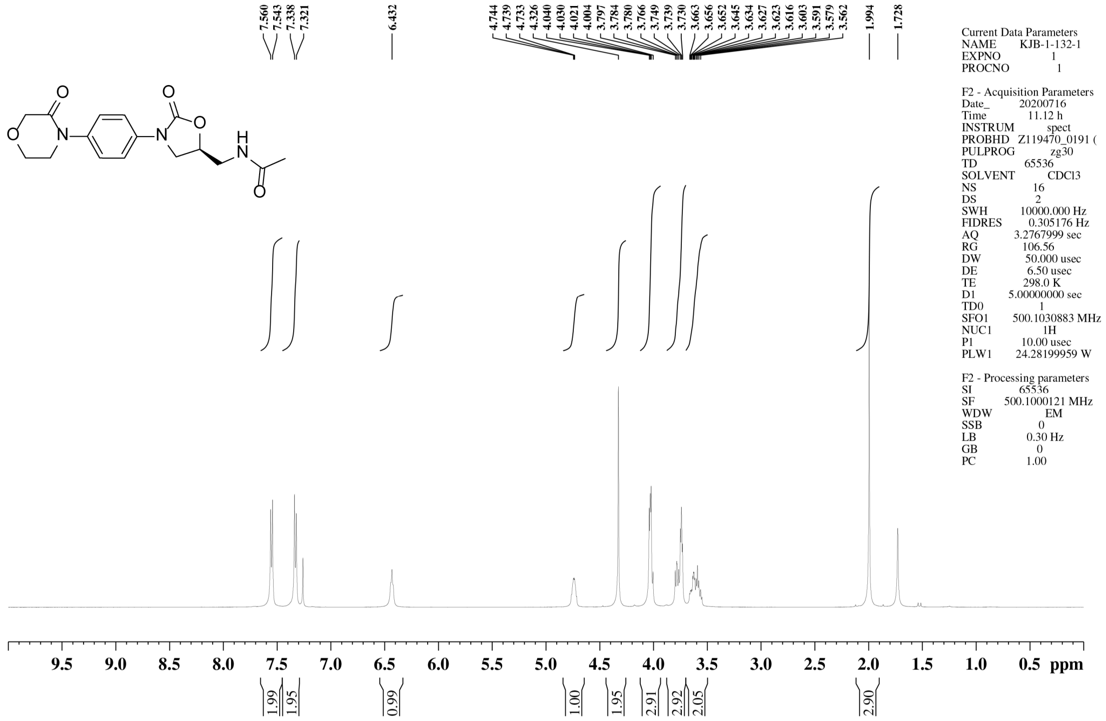


### $^{13}$C NMR spectrum of **57**

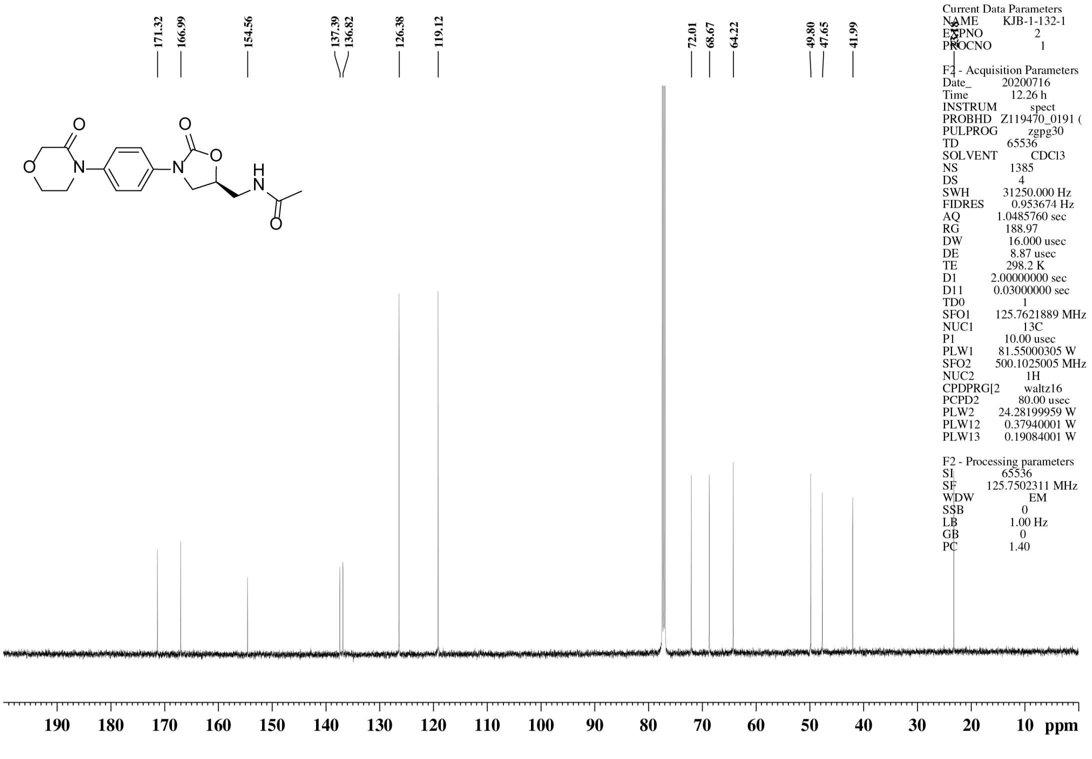
